# A high-throughput microbial glycomics platform for prebiotic development

**DOI:** 10.1101/2025.07.13.664583

**Authors:** Jennifer L. Modesto, Seth G. Kabonick, Jennifer E. Lausch, Kamalesh Verma, Kailyn M. Winokur, Jessica E. Gaydos, Asia Poudel, Gregory Yochum, Guy E. Townsend

## Abstract

The mammalian intestine contains diverse carbohydrate pools that govern the gut microbiome composition. Structurally distinct polysaccharides, also called glycans, are differentially consumed by gut microbial subsets and direct their abundance by controlling gene expression and metabolite production. Therefore, identifying gut microbial accessible carbohydrates (MACs) is necessary to develop new prebiotics that beneficially manipulate the gut microbiome. However, no methods exist to efficiently examine MACs in biologically-derived mixtures. Here, we present a high-throughput platform to detect MACs from various plant, animal, and microbial sources using a genome-wide library of engineered *Bacteroides thetaiotaomicron* (*Bt*) strains that harness their endogenous glycan detection machinery. We demonstrate that this platform exhibits specific and sensitive responses to glycan mixtures and use bacterially-encoded proteins to characterize a previously unknown MAC from yeast. Expanding this technology across gut *Bacteroides* species will generate a broadly applicable approach to characterize heterogeneous glycan mixtures and identify prebiotic substrates.

## Main

Commensal microbes harvest carbon from structurally diverse MACs, to achieve extraordinary cell densities in the mammalian intestine^1–5^. The glycosidic linkages that tether constituent monosaccharides into each MAC are frequently resistant to degradation in the gastrointestinal tract and therefore require substrate-specific enzymes to degrade^6–11^. Gut microbes can utilize available MACs by expressing carbohydrate-active enzymes that target specific glycosidic linkages^1, 8, 11, 12^. Characterizing MACs present in biologically-derived mixtures informs how and why the gut microbiota changes due to host dietary choices, genetics, or disease because chemically-distinct glycans support the expansion of defined microbial subsets. Additionally, identifying MACs that can influence microbial abundance and metabolism can promote engineered strain engraftment and gene expression to leverage the gut microbiome as a therapeutic target^13^. Glycan mixtures have traditionally been examined using tandem chromatography and mass spectrometry, arrayed glycan binding aptamers, or lectins that probe for a range of known carbohydrate structures^14–16^. Although these approaches can characterize glycan content in various sample types, they are unable to directly detect MACs.

Microbes access unique MACs by producing proteins that sequester, transport, degrade, and detect target molecules in their environment. For example, members of the dominant human intestinal phylum, *Bacteroidetes*, possess glycan sensors encoded within polysaccharide utilization genetic loci (PULs) that direct rapid, dramatic, and predictable increases to corresponding PUL gene transcripts following recognition of their target glycan structures^4, 6, 7, 17^. Ultimately, these sensors direct increased expression of surface glycan binding proteins (SGBPs), outer membrane transporters (SusCD), and glycolytic enzymes for consumption of available MACs. Different *Bacteroides* species encode vast and partially unique PUL repertoires, which endow each with the ability to access various target glycans that establish distinct intra-intestinal niches. We previously demonstrated that introduction of PUL promoters into a plasmid-borne *Bacteroides*-optimized *lux* cassette, p*Bolux,* converts these bacteria into biosensors of known and unknown target glycans present in biologically-derived mixtures^18^. We reasoned that PUL reporters could reflect the exquisite sensitivity and specificity of their cognate sensor proteins to detect MACs in glycan mixtures and be implemented to isolate and characterize those molecules in combination with corresponding PUL SGBPs and glycolytic enzymes.

Here, we generated a genome-wide *Bt* reporter library and arrayed these strains into a scalable and distributable high-throughput platform that sensitively indicates target glycans from various host-, microbial-, and plant-derived mixtures. We demonstrate that this approach distinguishes structurally distinct MACs according to monosaccharide composition, glycosidic linkages, and polysaccharide degree of polymerization. We use this technology to detect previously unknown PUL target glycans and employ a recombinant SGBP to isolate and characterize an unknown MAC in Baker’s yeast (*Saccharomyces cerevisiae).* Collectively, these results establish *Bacteroides* species as specific and sensitive toolkits to directly detect, isolate, and characterize unknown PUL target glycans that could be developed into prebiotics. Expanding these tools into additional *Bacteroides* species will support broad, untargeted glycomics applications to examine heterogenous glycan mixtures and overcome current detection limitations. Ultimately, our platform can be used to develop new prebiotics that manipulate gut microbial abundance and metabolism to benefit human health.

## Results

### Construction of an arrayed *Bt* PUL reporter library

We established that individual *Bt* PUL sensors direct predicted PUL transcript increases^19, 20^ because engineered strains expressing constitutively active sensor proteins from PULs 16, 19, 49, 53, and 57, exhibited dramatic increases in corresponding transcript abundances (Fig. 1a, Supplementary Table 1)^21, 22^. To examine whether sensor-directed PUL transcription can be faithfully reported by corresponding transcriptional reporters on a genome-wide scale, we generated a complete library of *Bt* PUL reporter plasmids by introducing putative promoters identified from RNAseq datasets^23^ and mapped transcription start sites^24, 25^ from annotated *Bt* PULs^26, 27^ into p*Bolux*^18^ (Supplementary Table 2). Although *Bt* possesses 88 PULs, we generated 91 total reporter strains because PULs 14, 52, 74, and 75 possess two divergently transcribed *susCD* pairs that were assigned unique reporters (denoted a and b) and PUL63 was omitted due to the absence of a *susD*-like gene (Fig. 1a, Supplementary Table 1, 2). Following cryopreservation of batch-produced 96-well plates, individual replicates were revived by the addition of rich media and incubated overnight before transfer to a 384-well plate and supplied 4 distinct carbohydrate preparations, one of which was always galactose as a negative control (Fig. 1b). Fold changes in growth-adjusted bioluminescence generated over 18 hours (Fig. 1c) were calculated from the area under each curve from strains supplied unique glycan preparations and normalized to a strain harboring the promoter-less p*Bolux* plasmid and cultures supplied galactose alone (Fig. 1d).

**Fig. 1.**
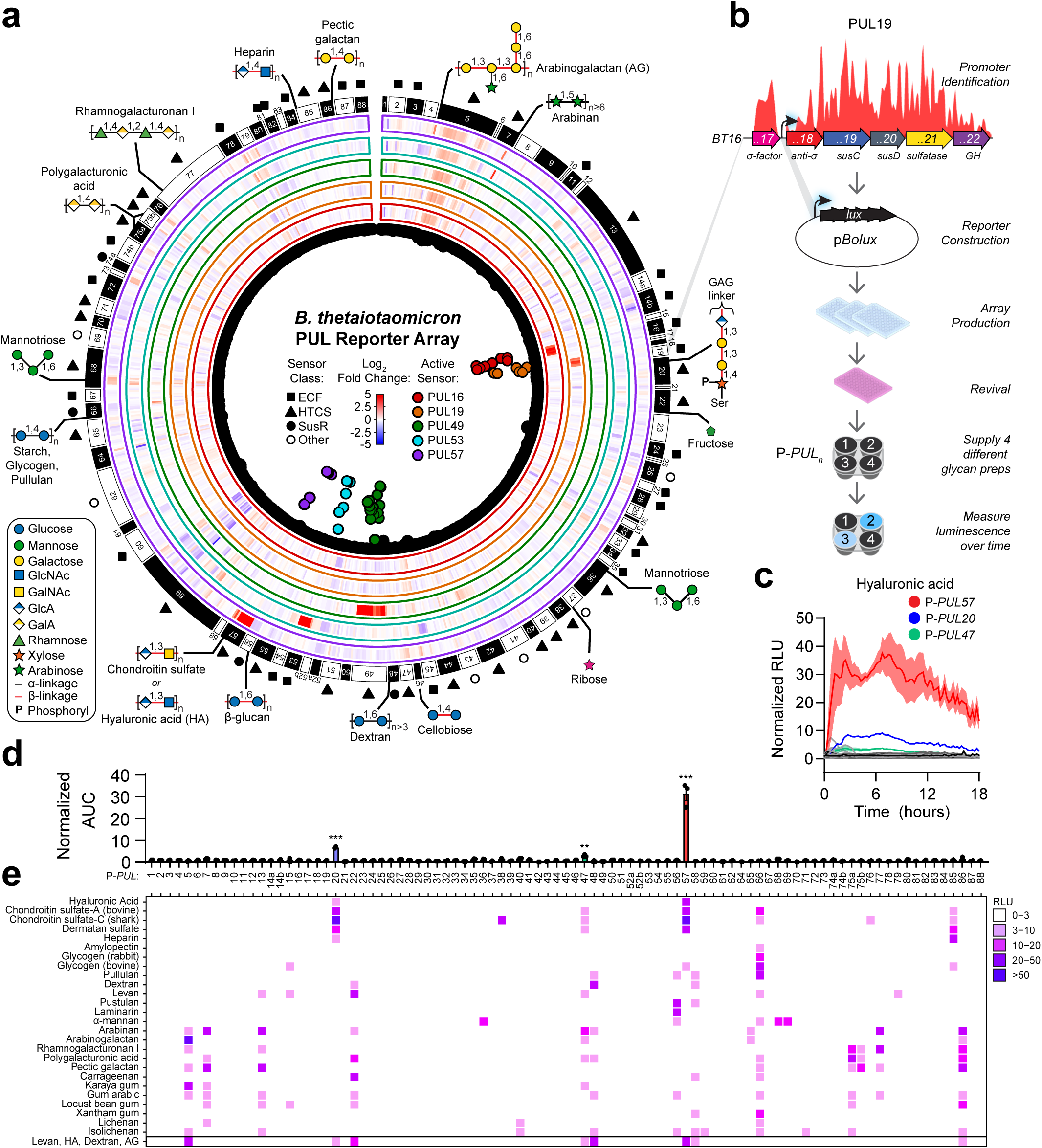
Construction of a genome-wide *Bt* PUL reporter library array. **a,** Cartoon depicting 88 *Bt* PULs denoted by individual genes (innermost black circles). Dots emanating inwards represent significantly increased PUL transcripts measured by RNA-seq from strains engineered to produce constitutively active sensor proteins from PUL16 (red), PUL19 (orange), PUL49 (green), PUL53 (cyan), or PUL57 (purple) and color-corresponding heatmaps are positioned centrally. The outermost symbols represent distinct classes of known or predicted PUL sensor proteins (filled square: ECF, extracytoplasmic sigma factor; filled triangle: HTCS, hybrid two-component system; filled circle: SusR, starch utilization system regular; open circle: other classes) and corresponding glycan structures that activate them emanate radially. **b,** schematic depicting PUL reporter construction and glycan analysis using the arrayed *Bt* library. **c-d,** Bioluminescence (**c**) from all 91 *Bt* reporters over 18 hours following introduction of hyaluronic acid and normalized by identical strains supplied galactose and corresponding (**d**) AUC normalized by responses from a strain containing p*Bolux*. All significantly increased reporters have corresponding colors and insignificant increases are displayed in grayscale. N=3, error is SEM. **e,** Heatmap of AUC over 18 hours from all reporters supplied mixtures containing galactose and various purified polysaccharides. N≥2. All statistics were calculated using 2-way ANOVA with Fisher’s LSD test and *** indicates *P*-values < 0.001, and ** < 0.01.

### *Bt* PUL reporters distinguish between glycosidic linkages and monosaccharide composition

To examine the *Bt* PUL reporter array functionality, we examined changes in bioluminescence from all strains following the introduction of various purified mono- and poly-saccharide preparations. As expected, the monosaccharides, fructose and ribose, increased bioluminescence from their respective PUL reporters, P-*PUL22* and P-*PUL37*, which agree with previous findings (Extended Data Fig. 1a)^18^. However, ribose also increased *PUL22* transcripts^28^ and corresponding bioluminescence from P-*PUL22* (Extended Data Fig. 1a, b). Ribose is likely contaminated with fructose because a *Bt* strain lacking *sensor*^PUL22^ (*BT1754*) harboring P-*PUL22* exhibited no bioluminescence increases when supplied ribose, whereas this strain harboring P-*PUL37* exhibited similar responses to *wild-type Bt* (Extended Data Fig. 1c). In contrast, a strain lacking *sensor*^PUL37^ (*BT2802*), which is unable to increase *PUL37* transcripts or utilize ribose as a sole carbon source^28^, did not display increased activity when harboring P-*PUL37* (Extended Data Fig. 1c). Similarly, P-*PUL22* also exhibited increased bioluminescence when supplied mannose (Extended Data Fig. 1a), that required *sensor*^PUL22^ (Extended Data Fig. 1b), suggesting that the mannose preparation examined here was also contaminated with fructose. These results indicate that the Sensor^PUL22^, which binds fructose with high affinity^7^, is responsible for increasing PUL22 expression when supplied ribose or mannose. Ultimately, these data highlight the specificity of PUL reporters for their target ligands and reflect their extraordinary sensitivity in mixtures.

We established that *Bt* PUL reporters can distinguish between MACs based on glycosidic linkages, monosaccharide composition, and degree of polymerization (d.o.p.). Distinct strains exhibited increased bioluminescence according to the administration of various purified plant, animal, and microbially-derived MACs consumed via known PULs^4, 6, 7, 11, 28, 29^. Strikingly, P-*PUL66*, P-*PUL48*, and P-*PUL56* were specifically activated by amylose, dextran, and pustulan, respectively, which are distinct glucose homopolymers differing by their α1,4-, α1,6-, and β1,6-glycosidic linkages (Fig. 1e). Each PUL is regulated by their respective SusR-like sensor proteins, encoded by BT3703 (Sensor^PUL66^)^30^, BT3091 (Sensor^PUL48^)^31^, and BT3309 (Sensor^PUL56^)^32^, to increase target PUL transcription and utilize these MACs as growth substrates. Sensor^PUL66^ and Sensor^PUL56^ differentiate between glycosidic linkage because their corresponding PUL reporters only exhibit increased bioluminescence when supplied α1,4- or β1,6-linked glucose disaccharides (Fig. 2a). However, P-*PUL48* can also distinguish between d.o.p. because bioluminescence from this strain only increased when supplied isomaltooligosaccharides greater than 3 α1,6-linked glucose units and requires at least 6 residues to reach activity similar to dextran (Fig. 2b). These data demonstrate that SusR-like PUL sensors detect distinct glycosidic linkages and d.o.p..

**Fig. 2.**
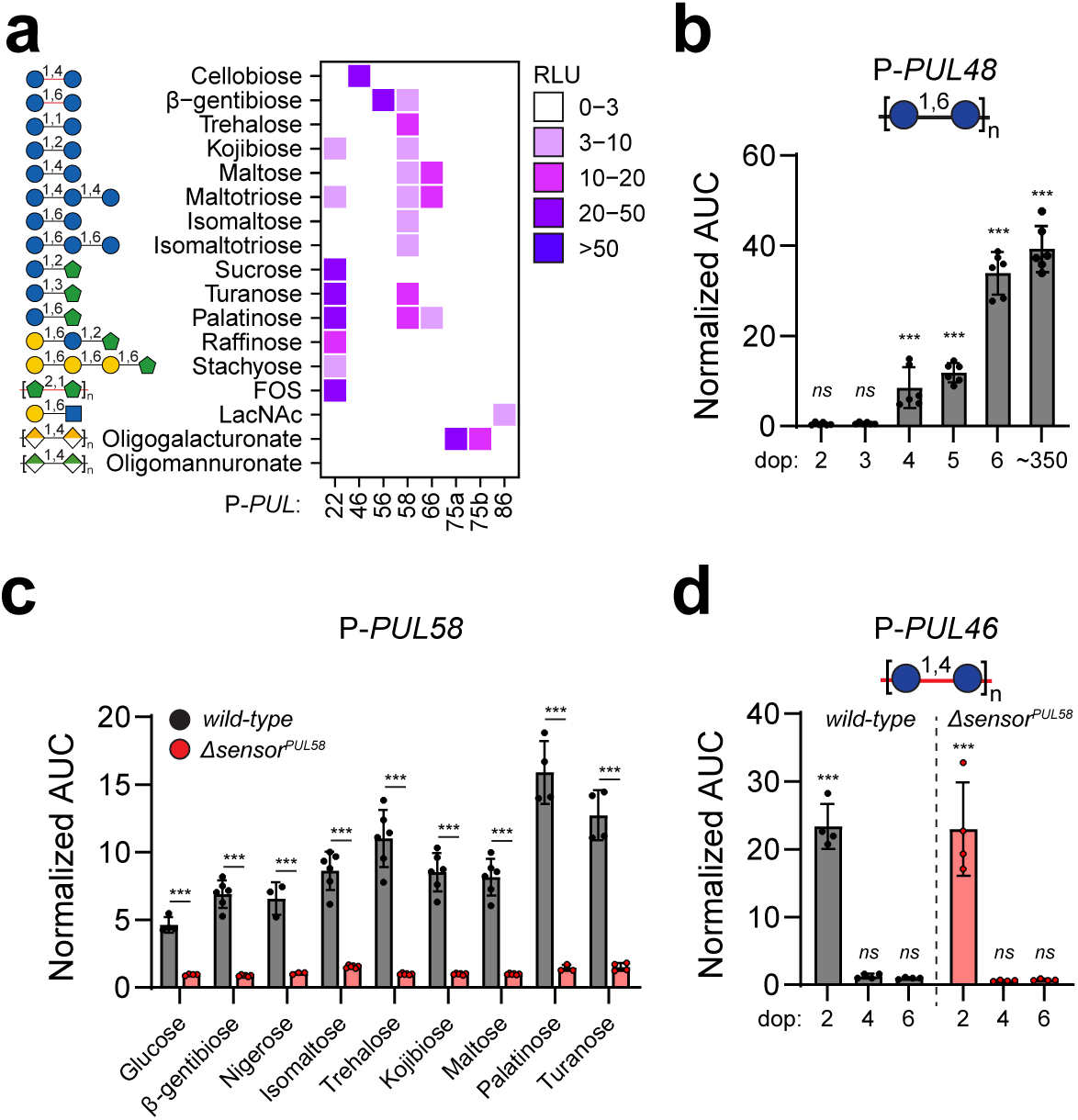
*Bt* PUL reporters distinguish between monosaccharide composition, d.o.p., and glycosidic linkage. **a,** A heatmap displaying normalized AUC following the introduction of various oligosaccharides to the *Bt* PUL reporter array. n=2. **b,** Normalized AUC from P-*PUL48* supplied isomaltooligosaccharides with increasing d.o.p. N=6, error is SEM. **c,** Normalized AUC from *wild-type* (black) or *sensor-^PUL^*^58^-deficient (*ΔBT2160*, red) strains harboring P-*PUL58* supplied the indicated di- and tri-saccharides. N≥3, error is SEM. **d,** Normalized AUC from *wild-type* (black) or *ΔBT2160* (red) strains harboring P-*PUL46* supplied the indicated cello-oligosaccha-rides. N=4, error is SEM. For panels **b-d**, *P*-values were computed using 2-way ANOVA. *** indicates values < 0.001 and *ns* indicates values > 0.05.

Further examination of purified oligosaccharides yielded several unexpected outcomes. First, monomeric glucose and various glucose-disaccharides, including maltose, isomaltose, trehalose, and β-gentibiose increased bioluminescence from P-*PUL58* (Fig. 2a). Similarly, supplying glucose-fructose heterodimers, turanose and palatinose, stimulated greater than 10-fold increases from this strain, whereas sucrose did not, suggesting that this PUL reporter responds to glucose- and fructose-containing disaccharides subsets. We hypothesized that P-*PUL58* could be governed by the SusR-like sensor, encoded by *BT2160*, that is an important *Bt* fitness determinant during growth in trehalose and palatinose^33^. Accordingly, *Δsensor^PUL^*^58^ harboring P-*PUL58* did not exhibit increased bioluminescence in any condition, indicating that this sensor facilitates PUL58 transcription in response to various disaccharides (Fig. 2c). Curiously, this sensor also responds to glucosinolates derived from cruciferous vegetables and is required for their conversion into isothiocyanates by increasing transcription of the linked genes *BT2159-56*^34^. A deeper examination of how and why this distally encoded sensor protein responds to these signals is necessary to understand this regulatory relationship. Unexpectedly, P-*PUL46* detected a *Bt*-inaccessible carbohydrate^35^ because introduction of the β1,4-linked glucose disaccharide, cellobiose, increased bioluminescence from this strain (Fig. 2a, d). This response was d.o.p.-specific because there was no change when supplied cellulose-derived tetra- or hexa-saccharides and this response did not require *sensor^PUL^*^58^ (Fig. 2d). Because *Bt* cannot utilize cellulose or its polysaccharide breakdown products including cellobiose^35^, these results suggest that cellobiose is a signal in the intestine.

### *Bt* PUL reporters detect discreet carbohydrate structures

The *Bt* PUL array can also distinguish between various microbial-, animal-, and plant glycans covalently tethered into polymeric complexes. For example, *Bt* utilization of yeast cell wall α-mannans (Extended Data Fig. 2a) requires three PULs: 36, 68, and 69^36, 37^, which increase activity from the reporter strains, P-*PUL36*, P-*PUL68,* and P-*PUL69* by the addition of purified α-mannan (Fig. 1e). The corresponding sensor proteins, BT2628 (Sensor^PUL36^) and BT3786 (Sensor^PUL68^) facilitate mannan-responsive PUL36/68 transcription because bioluminescence from strains harboring either P-*PUL36* or P-*PUL68* is not apparent in a strain lacking both sensors (*Δsensors*^PUL36/68^; Extended Data Fig. 2b), consistent with previous transcript measurements^36^. Alternatively, PUL69 transcription is facilitated by Sensor^PUL69^, encoded by *BT3853*^36^, which is required for increased bioluminescence from P-*PUL69* when supplied α-mannan (Extended Data Fig. 2b). Sensors^PUL36/68/69^ are collectively required for mannan utilization as a sole carbon source (Extended Data Fig. 2c) but detect discreet carbohydrate structures because the branched α1,3-[α1,6] trisaccharide increases P-*PUL36/68* bioluminescence (Extended Data Fig. 2d, e). Conversely, P-*PUL69* did not respond to any di- or tri-saccharide examined, indicating it responds to a different oligo-mannan structure (Extended Data Fig. 2f). These data highlight that PUL sensor proteins direct cognate PUL expression following recognition of distinct glycan moieties assembled as larger polymers.

*Bt* possess 3 PULs that respond to glycosaminoglycans (GAGs), which are intestinally abundant heteropolymers comprised of amine sugars and uronic acids, including previously characterized PULs 57^6, 10^ and 85^38^, and the uncharacterized PUL20 (Fig. 3a). Strains harboring P-*PUL57* exhibited increased bioluminescence when supplied chondroitin sulfate (CS) or dermatan sulfate (DS; Fig. 1e), which are variably sulfated linear polymers of repeating, disaccharide units comprised of glucuronic acid (GluA) and β1,3-linked N-acetylgalactosamine (GalNAc). P-*PUL57* also responds to hyaluronic acid^18^, an unsulfated GAG whose disaccharide unit is comprised of gluA and β1,3-linked N-acetylglucosamine (GlcNAc) because the Sensor^PUL57^ protein, BT3334, cannot distinguish between either CS- or HA-derived disaccharide subunits (Fig. 3b)^6^. Introduction of CS, DS, and HA also increased bioluminescence from P-*PUL20* (Fig. 1e), consistent with previous transcriptional measurements^3^, suggesting that all GAGs activate an additional PUL. However, Sensor^PUL57^ is not responsible for increasing bioluminescence from P-*PUL20* because: 1.) CS- and HA-derived disaccharides increased activity from P-*PUL57* but not P-*PUL20* (Fig. 3b), 2.) heterologous expression of a constitutively active Sensor^PUL57^ (BT3334*) increased *PUL57* but not *PUL20* transcripts (Fig. 1a), and 3.) BT3334* increased bioluminescence from strains harboring P-*PUL57* but not P-*PUL20* (Fig. 3c). Furthermore, a distinct GAG, heparin, also increased bioluminescence from only P-*PUL20* and P-*PUL85* (Fig. 1e), whereas the composite disaccharide comprised of alternating N-acetylglucosamine (GlcNAc) and uronic acid increased activity from only P-*PUL85* (Fig. 3b). We propose that PUL20 responds to the common GAG tetrasaccharide linker, GlcA-[β1-3]-Gal-[β1-3]-Galβ-[1-4]-Xyl2P-β-Ser (Fig. 1a), because CS from shark cartilage stimulates increased bioluminescence from P-*PUL20* and P-*PUL57* but no other contaminating glycans following co-incubation with SGBP^PUL57^ (Fig. 3d)^18^, indicating that the PUL20-target glycan is covalently tethered to CS.

**Fig. 3.**
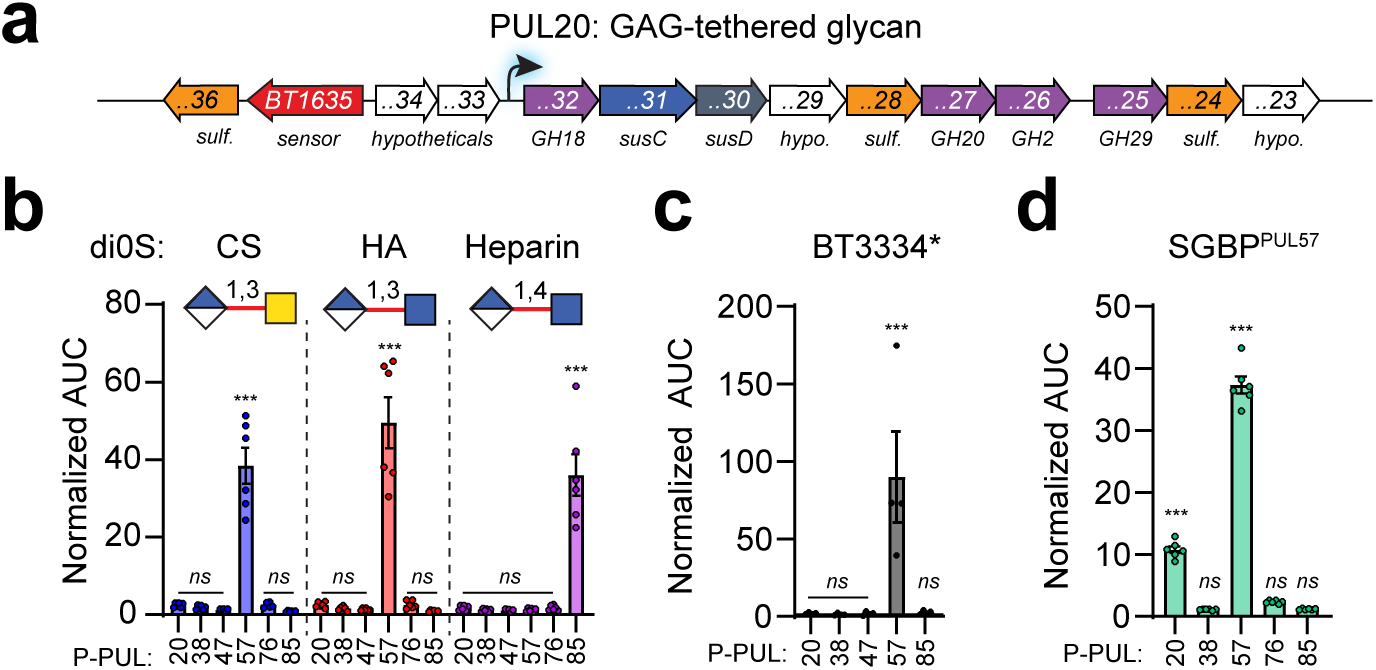
PUL 20 responds to a unique GAG-tethered glycan. **a,** cartoon representing PUL20 activated by GAGs. **b**, Normalized bioluminescence from *Bt* strains harboring P-*PUL20,* P-*PUL38,* P-*PUL47,* P-*PUL57,* or P-*PUL85* supplied unsaturated disaccharides (di0S) derived from CS (blue), HA (red), or heparin (purple) and normalized by responses from a strain harboring p*Bolux*. N=6. **c,** Normalized bioluminescence from a strain expressing BT3334* harboring the reporters described in (**b**) and normalized by identical strains harboring the empty vector cultured in galactose as a sole carbon source. N=4. **d,** Normalized bioluminescence from *Bt* strains harboring P-*PUL20,* P-*PUL38,* P-*PUL47,* P-*PUL57,* or P-*PUL85* supplied SGBP^PUL57^ co-purified material following incubation with shark CS and normalized by responses from a strain harboring p*Bolux*. N=6. For panels **b-d**, error is SEM and *P*-values were calculated using 2-way ANOVA with Fisher’s LSD test. *** indicates values < 0.001 and *ns* > 0.05.

Plant cell wall glycans called pectins and hemicelluloses, are frequently assembled into complex polymeric structures (Extended Data Fig. 3a) that are targeted by distinct *Bt* PULs^1, 4, 9, 11^. This is exemplified by P-*PUL7*, which exhibited increased bioluminescence when supplied arabinan (Fig. 1e). However, arabinan also elicited increases from strains harboring P-*PUL5*, P-*PUL13*, P-*PUL22*, P-*PUL47*, P-*PUL48*, P-*PUL65*, P-*PUL77*, and P-*PUL86*, indicating this preparation also contains arabinogalactan (PUL5), rhamnogalacturonan II (PULs 13 & 21), fructose (PUL22), dextran (PUL48), rhamnogalacturonan I (PUL77), and pectic galactan (PUL86) (Extended Data Fig. 3b). Reporter activity faithfully reflects sensor-dependent activity because the Sensor^PUL7^ (BT0366)^39^, which recognizes arabino-oligomers of at least 6 α1,5-linked arabinose residues^4^, was only required to stimulate P-*PUL7* when supplied arabinan (Extended Data Fig. 3b). Consistent with this notion, purified oligo-galacturonate increased bioluminescence from its corresponding reporter, P-*PUL75a&b* (Fig. 2a), whereas highly polymeric PGA also increased activity from P-*PUL7*, P-*PUL22*, P-*PUL47*, P-*PUL48*, and P-*PUL66*, and P-*PUL86* (Fig. 1e), demonstrating that PUL reporters respond predictably to complex MACs comprised of multiple ligands^1, 4, 7, 9, 11^. Finally, the ubiquitous food additives carrageenan, xantham gum, locust bean gum, karaya gum, and gum arabic are complex heteropolymers of largely *Bt*-inaccessible carbohydrates^9, 35^ but contain small amounts of detectable structures because each elicits increased bioluminescence from at least one PUL reporter (Fig. 1e). For example, P-*PUL5* is dramatically activated by purified arabinogalactan and exhibits significant increases when supplied karaya gum and gum arabic^9^ (Fig. 1e). Collectively, these data demonstrate the exquisite specificity of *Bt* PUL reporters for complex multi-valent MACs, facilitated by the cognate sensor proteins.

### The *Bt* PUL reporter array can faithfully survey glycan mixtures

*Bt* PUL reporters can simultaneously examine MACs present in heterogeneous glycan mixtures because P-*PUL5*, P-*PUL22*, P-*PUL57*, and P-*PUL48* exhibited increased bioluminescence when supplied an equal mixture of purified arabinogalactan (AG, PUL5), levan (PUL22), hyaluronan (HA, PUL57), and dextran (PUL48) (Fig. 1e). Additionally, this mixture increased activity from P-*PUL20* and P-*PUL47*, which were both increased by HA (Fig. 1c-e), displaying faithful reporting of lab-prepared mixtures. To characterize biologically derived mixtures, we measured reporter responses to PMOGs, a mixture of porcine mucosal-derived O-glycans that increases many *Bt* PUL transcripts^3, 18, 40^. When supplied PMOG, we detected increased activity in GAG-, and O-glycan-specific PUL reporters, consistent with previous transcript measurements^3, 18, 40^ indicating that the reporter strains can faithfully examine biologically-derived mixtures (Fig. 4a). To prepare other host-, diet-, and microbially-derived mixtures, we adapted the oxidative-release of natural glycans (ORNG) method that can liberate glycans from a wide variety of biological materials^41^. We determined that a preparation of porcine mucosal glycans using ORNG (PMG), yielded similar reporter profiles as PMOG (Fig. 4a), but did display a 2-fold decrease in bioluminescence from a strain harboring P-*PUL14b* and a 2-fold increase in signal from P-*PUL48*, indicating glycan subsets are differentially extracted or detected when this approach is applied (Extended Data Fig. 4a).

**Fig. 4.**
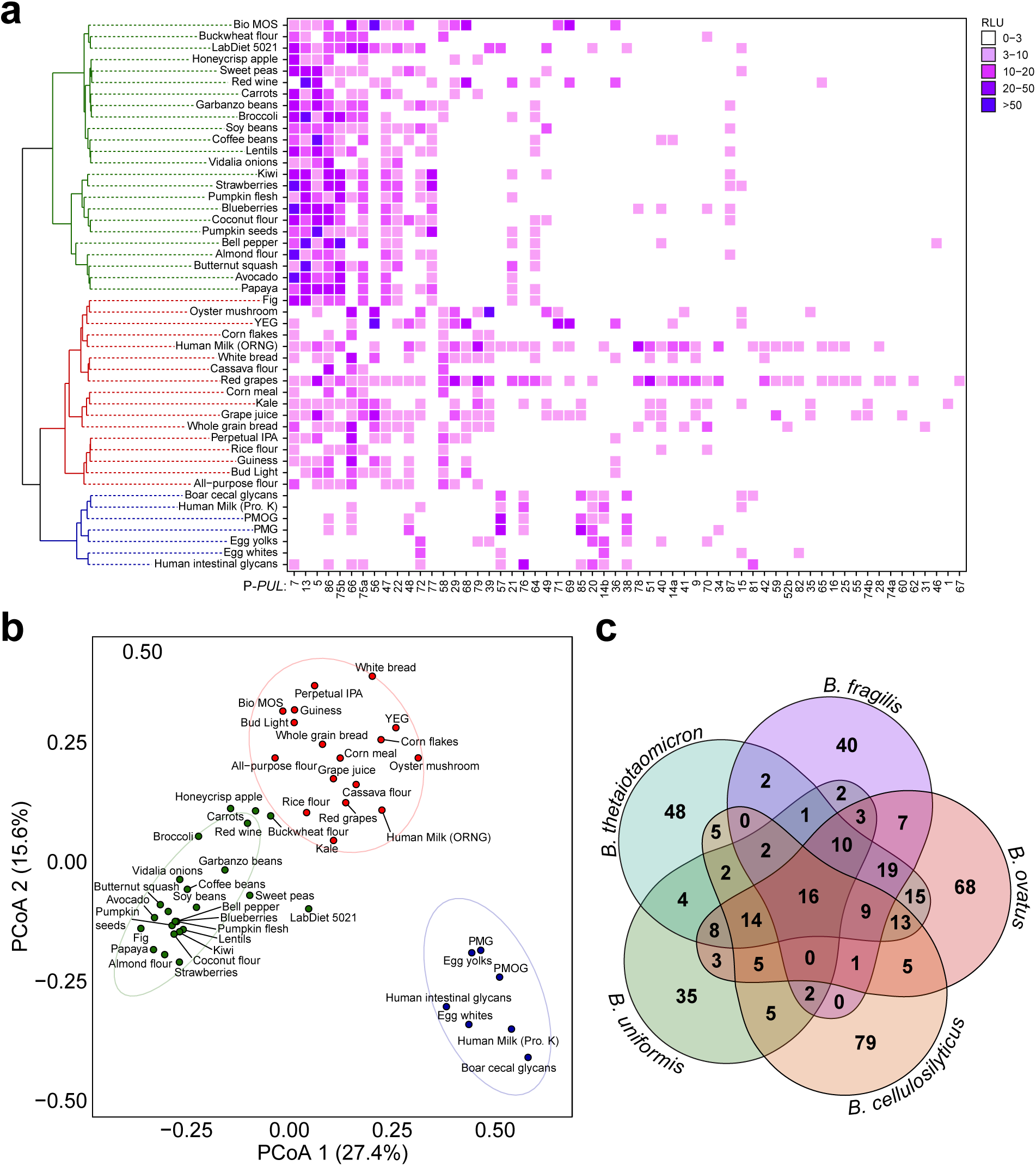
Arrayed *Bt* PUL reporter strains differentiate MAC sources. **a,** Heatmap representing *Bt* PUL array responses to the indicated biologically-derived mixtures clustered by reporter activation similarity. n≥2. **b,** PCoA of *Bt* reporter responses to different MACs from (**a**) using the Jaccard similarity index and Ward D2 distance calculations. **c,** Distribution of unique and shared predicted PULs across the indicated *Bacteroides* species.

To determine how gut microbial responses differ between animal and human mucosal material, we compared human-derived intestinal glycan mixtures collected from various patient intestinal samples recovered following surgery (HIG) and porcine cecal mixtures from freshly slaughtered boars (BCG). HIG elicited similar patterns of reporter activity as PMOG and fresh porcine cecal glycans, suggesting that mammalian hosts largely supply gut microbes with similar growth substrates (Fig. 4a). For example, HIG and BCG elicited similar bioluminescence from P-*PUL14*b, P-*PUL20*, P-*PUL38*, P-*PUL66*, and P-*PUL72* (Extended Data Fig. 4b). However, the following distinctions were readily apparent: on one hand, human material elicited reduced bioluminescence from the CS/HA reporter P-*PUL57* and heparin reporter P-*PUL85*, but on the other hand they elicited increased responses from P-*PUL76*, *P-PUL81, and P-PUL86* (Extended Data Fig. 4b). Furthermore, human milk glycans (HMG) elicited similar responses to HIG, however, signals were improved when HMG were prepared by defatting and proteinase K treatment (Extended Data Fig. 4c), suggesting that ORNG can result in glycan peeling from some source materials^42^.

We used ORNG to prepare various animal, plant, and fungal material that are abundant components of the human diet (Fig. 4a) and examined *Bt* reporter responses. As expected, plant-, animal-, and microbially-derived material occupied distinct space following PCoA analysis, indicating that MAC distribution reflects source phylogeny at the phylum level (Fig. 4b). For example, dietary plant products emanating from both mono- and di-cot species distinctly increased bioluminescence from strains harboring P-*PUL5* (arabinogalactan), P-*PUL7* (arabinan), P-*PUL13* and P-*PUL21* (RGII), P-*PUL22* (fructan), P-*PUL66* (starch/glycogen), P-*PUL75* (PGA), and P-*PUL86* (RGI), which is consistent with common plant fibers. Conversely, mammalian derived mixtures stimulated P-*PUL14b* (N-glycans), P-*PUL20* (GAG associated), P-*PUL38* (unknown mucin *O*-glycan), P-*PUL57* (CS/HA), P-*PUL66* (glycogen), and P-*PUL76* (unknown mucin *O*-glycan). Finally, fungal products consistently increased activity from strains containing P-*PUL29* (unknown), P-*PUL36/68* (α-mannan), P-*PUL56* (β-glucan), P-*PUL66* (starch/glycogen), and P-*PUL69* (α-mannan). Accordingly, processed foods containing both plant and fungal products, including breads and beer, contained reporter increases consistent with both grains and yeast (Fig. 4a).

Unexpectedly, several PUL reporters did not exhibit bioluminescence increases when supplied any purified carbohydrate or mixture examined in this study, suggesting either these target glycans are not present in the mixtures prepared for this work or the corresponding reporter plasmid may not contain the appropriate regulatory sequence to facilitate increased bioluminescence. However, we determined that the latter is an unlikely scenario because qPCR analysis for *susC* transcripts from PULs whose corresponding reporters failed to increase bioluminescence were not increased when supplied various mixtures (Extended Data Fig. 5a). Furthermore, expression of constitutively active sensor proteins from PULs 16, 19, 49, 53, and 57, which increased target PUL transcription (Fig. 1a)^21, 22^, stimulated bioluminescence increases when these strains contained the corresponding PUL reporter (Fig. 3c, Extended Data Fig. 5b). A notable exception is PUL2, which targets an unknown glycan, exhibits 180-fold increased transcripts when supplied yeast extract and red-wine derived glycan mixtures (Extended Data Fig. 5a), but whose corresponding reporter did not exhibit increased bioluminescence despite engineering an additional plasmid that included an internal transcription start site^25, 40^ (Extended Data Fig. 5c). Thus, the majority of PULs can be converted into glycan biosensors with a small minority that may require additional optimization to use p*Bolux*. Finally, establishing the functionality of various PUL reporters with unknown target glycans (Extended Data Fig. 5b) that are not activated across the common foods examined in this study, indicate that future investigations should focus on other glycan sources, including co-resident microbes and fermented products.

### Unique PULs across p*Bolux*-compatible *Bacteroides* species

This study demonstrates how *Bt* PUL reporters can be readily constructed and deployed to detect MAC content in various glycan preparations. However, other *Bacteroides* species possess unique PUL repertoires that putatively confer distinct glycan accessibility, indicating that they recognize a partially unique subset of carbohydrate structures (Fig. 4c, Supplementary Table 3). For example, *B. fragilis* PULs facilitate its growth in host-derived glycan preparations including those distinct from *Bt* target glycans^22^, whereas *B. ovatus*^43^ and *B. uniformis*^44^ possess PUL repertoires that target *Bt*-inaccessible hemicelluloses. We previously demonstrated that p*Bolux* can report promoter-dependent transcription in *B. ovatus*^18^ and *B. fragilis*^45^, and this is consistent in *B. uniformis* and *B. cellulosilyticus* because introduction of their corresponding *rpoD* promoters in this plasmid increase bioluminescence from all five species (Extended Data Fig. 6). Therefore, construction of future PUL reporter libraries in these species will expand the breadth of glycan detection capabilities to comprehensively characterize MAC content in host-, microbial-, and dietary-components.

### Isolation and characterization of a *Saccharomyces cerevisiae* glycan consumed by *Bt*

To explore the utility of PULs as glycomics tools for prebiotic development, we characterized an unknown glycan detected in a biologically derived mixture. We selected PUL71 (Fig. 5a) because yeast extract glycans (YEG) elicited 32-fold increased bioluminescence from strains harboring P-*PUL71*, which was also stimulated by preparations from yeast containing products like red wine and animal feeds, but not breads or beers (Fig. 4a). Furthermore, P-*PUL71* activity was increased in yeast-derived mannan prebiotic mannooligosaccharides (Bio MOS) but absent in host- and most diet-derived mixtures (Fig. 4a). Furthermore, this glycan is an excellent prebiotic candidate because a strain lacking PUL71 (*ΔBT3957-BT3965*) exhibited a 5-fold fitness defect relative to *wild-type Bt* in mice supplied a 0.5% YEG solution in place of water (Fig. 5b). *PUL71* includes a unique glycan sensor protein class, encoded by *BT3957*, that possesses a periplasmic glycan binding domain and a cytoplasmic DNA-binding domain similar to HTCSs (Extended Data Fig. 7a). We established that the putative sensor, BT3957, controls PUL71 transcription because a *sensor^PUL^*^71^-deficient strain (*Δsensor*^PUL71^) harboring P-*PUL71* failed to exhibit increased activity in response to YEG (Extended Data Fig. 7b). Conversely, *sensor*^PUL36/68^- or *sensor*^PUL69^-deficient strains harboring P-*PUL71* exhibited similar bioluminescence as *wild-type Bt* supplied YEG (Extended Data Fig. 7b). PUL71 responds to a glycan distinct from other well characterized yeast-derived glycans including α-mannan, which activates P-*PUL36/68/69* (Extended Data Fig. 2b)^36^ and β-1,6-glucan, which stimulates P-*PUL56* (Fig. 2a)^32^, because bioluminescence from P-*PUL71* was unchanged when supplied these purified glycans (Fig. 1e) and *Δsensor*^PUL71^ exhibited similar activity to *wild-type Bt* harboring P-*PUL36* or P-*PUL56* when supplied YEG (Extended Data Fig. 7b), indicating that Sensor^PUL71^ recognizes a unique yeast-derived glycan that increases *PUL71* expression.

**Fig. 5.**
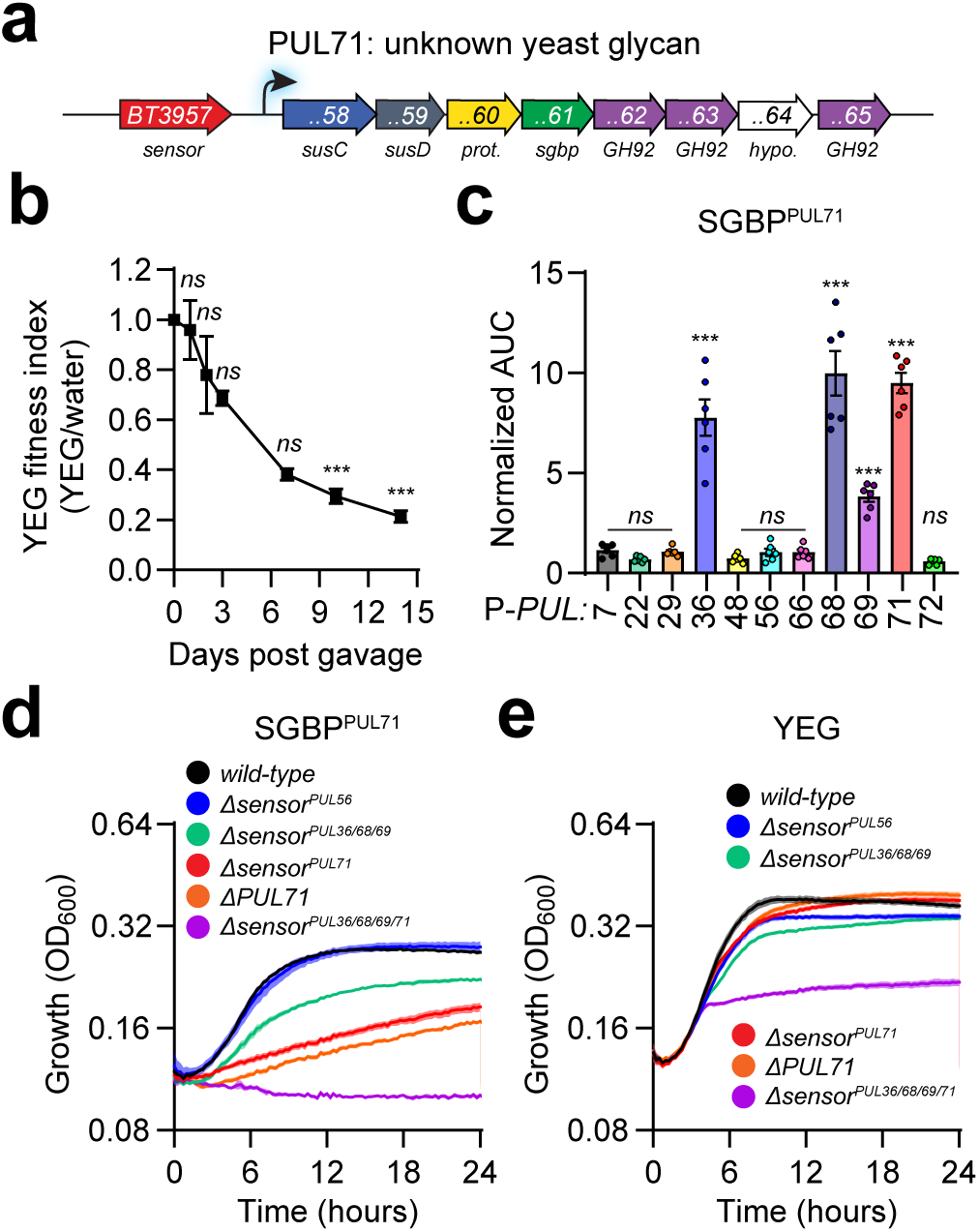
PUL71 consumes an unknown yeast cell wall glycan. **a,** Cartoon schematic of PUL71, which responds to yeast glycans. **b,** Fitness index of a *PUL71*-deficient strain relative to *wild-type Bt* in mice supplied a 0.5% YEG solution compared to those supplied water. N=5 and error is SEM. *P*-values were calculated using 2-way ANOVA with Bonferroni correction. *** indicates values < 0.001 and *ns* > 0.05. **c,** Normalized bioluminescence of YEG-induced reporter strains supplied SGBP^PUL71^-isolated glycans from GH30^PUL56^-treated YEG. N=6, error is SEM. *P*-values were calculated using 2-way ANOVA with Fisher’s LSD test. *** indicates values < 0.001 and *ns* > 0.05. **d&e,** Growth of *wild-type* and the indicated sensor-deficient *Bt* strains supplied (**d**) SGBP^PUL71^-co-purifying material following incubation with GH30^PUL56^-treated YEG or (**e**) untreated YEG. N=6, error is SEM.

To characterize the PUL71 target glycan, we co-incubated YEG with recombinant hexahistidine-tagged SGBP^PUL71^ (BT3961) and recovered protein-bound glycans using metal affinity chromatography^18^. The recovered glycans were enriched for the PUL71 target over other MACs present in YEG because the co-purifying material significantly increased activity from P-*PUL71,* but P-*PUL7* (α-arabinan), P-*PUL22* (fructans), P-*PUL29* (unknown), P-*PUL48* (dextran), P-*PUL66* (glycogen), and P-*PUL72* (high mannose N-glycan^36^) were no longer detectable (Extended Data Fig. 7c). However, P-*PUL36/68/69* and P-*PUL56* also exhibited increased activity indicating that α-mannan and β-glucan co-purified with the PUL71 target (Extended Data Fig. 7c). Significantly activated PULs contributed to accessing the SGBP^PUL71^ co-purifying material because the corresponding sensor-deficient strains exhibited reduced growth on this material as a sole carbon source, with *Δsensor*^PUL71^ exhibiting the largest reduction (Extended Data Fig. 7d). We hypothesized that the BT3961-purified material contained covalently tethered carbohydrates comprised of the PUL71-, PUL36/68/69-, and PUL56-target glycans because the yeast cell-wall is an interconnected matrix comprised of chitin, β1,3-, and β1,6-glucans decorated with mannoproteins through a GPI-anchor remnant glycan (Extended Data Fig. 2a). Consistent with this notion, material co-purifying with the SGBP^PUL56^, BT3313, incubated with YEG, also increased bioluminescence from strains harboring P-*PUL56*, P-*PUL36/68*, and P-*PUL71*, although bioluminescence from P-*PUL69* was not significantly increased possibly because the corresponding ligand was below the limit of detection (Extended Data Fig. 7e). Accordingly, only *Δsensor*^PUL56^, which is required for growth on β1,6-glucan (Extended Data Fig. 7f), exhibited substantially reduced growth when supplied SGBP^PUL56^-isolated material (Extended Data Fig. 7g), indicating the other target glycans are present in relatively low abundances. Collectively, these data demonstrate that the PUL56 and PUL71 SGBPs isolate their corresponding target glycans when assembled into polymeric structures.

To determine how the PUL71-target glycan is tethered to β1,6-glucan and α-mannan, we treated YEG with the PUL56-encoded β1,6-glucosidase (BT3312), referred to here as GH20^PUL56^, which liberated glucose and β-gentibiose from YEG (Extended Data Fig. 7h). We determined that GH^PUL56^ treatment removed β1,6-glucan, but not α-mannan, from the PUL71 target because SGBP^PUL71^-isolated material no longer increased bioluminescence from P-*PUL56* but exhibited significant increases from P-*PUL36/68/69* (Fig. 5c). Accordingly, *Δsensor*^PUL71^ and *Δsensors*^PUL36/68/69^ each exhibited reduced growth on this material as a sole carbon source and growth was completely abolished in a strain lacking all four sensors (*Δsensors*^PUL36/68/69/71^) (Fig. 5d), whereas these strains grew on YEG with reduced growth maxima (Fig. 5e). P-PUL71 likely responds to a distinct manno-oligosaccharide because introduction of various purified substrates were unable to increase bioluminescence (Extended Data Fig. 7i). Furthermore, compositional analysis demonstrated that SGBP^PUL71^-isolated material from untreated YEG contained glucose (Extended Data Fig. 8a), whereas this monosaccharide was no longer detectable when YEG was pre-treated with GH20^PUL56^ (Extended Data Fig. 8b). Consistent with this finding, HPAEC-PAD analysis revealed that SGBP^PUL56^-isolated glycans from untreated YEG contained mannose (Extended Data Fig. 8c), further suggesting that yeast β1,6-glucan and α-mannan are tethered in the absence of mannoproteins, which are eliminated by ORNG extraction. These data indicate that yeast α-mannan, β-glucan, and PUL71-target glycan are covalently assembled into a complex resistant to oxidation, and that the PUL71 target glycan decorates yeast-derived α-mannan prepared using ORNG (Fig. 4a) but not alkaline extraction, which was used to generate the α-mannan standard (Fig. 1e). PUL71 includes the mannosidases BT3962 and BT3965, which respectively convert α1,2- and α1,4-mannobioses into mannose (Extended Data Fig. 9a, b), and a putative mannosidase BT3963, which was unable to hydrolyze any mannobiose examined here (Extended Data Fig. 9c-f), in agreement with previous results^46^. Collectively, these data suggest that the PUL71-target glycan is an unknown yeast α-mannan decoration comprised of unique glycosidic linkages that can function as a prebiotic.

## Discussion

We have developed a microbial glycomics platform that harnesses *Bacteroides* PUL sensors to detect, isolate, and characterize MACs from biologically derived mixtures (Fig. 1). We demonstrated that a genome-wide *Bt* PUL-reporter library can indicate the monosaccharide composition, glycosidic linkages and d.o.p. of purified *Bt* accessible-(Fig. 1e, 2b) and - inaccessible carbohydrates (Fig. 2d). Furthermore, we demonstrated this toolkit can efficiently characterize MAC content in semi-purified preparations (Fig. 1e, Extended Data Figs. 1, 2) and various mixtures derived from the host mucosa and dietary material (Fig. 1e and Fig. 4a). We establish that reporter responses largely correspond to the biological source of each mixture (Fig. 4b), indicating that MAC content reflects phylogeny. The exquisite specificity and sensitivity of *Bacteroides* reporter strains highlight their utility as glycan biosensors that could be implemented in product quality control to detect carbohydrate contaminants (Extended Data Fig. 1b, c) and co-purifying moieties (Fig. 3). Furthermore, this approach could be applied to broader glycomics applications including surveillance of different carbohydrate structures from patient samples, plant cultivars, and microbial isolates (Fig. 4a). Finally, *Bt* PUL reporters can directly identify novel MACs that could be translated into prebiotics to manipulate the gut microbiome composition and products for therapeutic purposes (Fig. 5). Therefore, PUL reporters offer a powerful new toolkit that can be expanded using additional *Bacteroides* species that possess unique PUL repertoires (Fig. 4c, Extended Data Fig. 6).

## Materials and Methods

### Bacterial growth conditions

All bacteria were cultured as described previously^47^. Briefly, *Bacteroides* strains (described in Supplementary Table 5) were cultured on brain-heart infusion agar (MilliporeSigma) containing 5% horse blood (Hardy) under anaerobic conditions (85% N_2_, 12.5% CO_2_, 2.5% H_2_). Liquid cultures were inoculated from a single colony into TYG media and incubated under anaerobic conditions before sub-culture into Bacteroides minimal media containing 100 mM KH_2_PO_4_ (pH=7.2), 15 mM NaCl, 8.5 mM (NH_4_)_2_SO_4_, 0.5 μg/ml of L-cysteine, 1.9 μM hematin, 200 μM L-histidine, 100 μM MgCl_2_, 1.4 μM FeSO_4_, 50 μM CaCl_2_, 1 μg/ml of vitamin K_3_, 5 ng/ml of vitamin B_12_ and individual carbon sources described in Supplementary Table 4. All bacterial strains included the following antibiotics where appropriate: 100 μg/mL ampicillin, 200 μg/mL gentamicin, 2 μg/mL tetracycline, or 25 μg/mL erythromycin (MilliporeSigma).

### RNA-seq

Strains engineered to express constitutively active PUL sensors or corresponding vector control were cultured in triplicate in minimal media containing galactose as the sole carbon source to mid-exponential phase (OD_600_ = 0.45 - 0.7) before collection by centrifugation, frozen on dry ice, and stored at −80°C. Cell pellets were treated with RNAprotect Bacteria Reagent (Qiagen) and RNA was harvested using RNeasy Mini Kit with on-column DNase digestion (Qiagen) according to the manufacturer’s instructions. Purified RNA was treated with Turbo DNase (Invitrogen) according to the manufacturer’s instructions and re-purified using RNeasy Mini Kit with on-column DNase digestion. RNA-sequencing was performed by SeqCoast Genomics using a Nextseq 2000 (Illumina) following ribosomal RNA depletion using the Ribo-Zero Plus Microbiome rRNA Depletion Kit (Illumina) to generate 12 million 150bp paired-end reads per sample. Reads were aligned to the *B. thetaiotaomicron* genome (strain VPI-5482; GenBank accession number NC_004663) in Galaxy using Bowtie2 (v2.5.3). The mapped reads were quantified using featureCounts (v2.0.8) and differential gene expression was measured using DEseq2 (v2.11.40.8) and values are listed in Supplementary Table 1. Raw files are available on the NCBI Sequence Read Archive, PRJNA1298319.

### Reporter construction

Reporter plasmids were constructed by amplifying PUL promoter fragments (Supplementary Table 2) using oligonucleotides (Supplementary Table 6) as previously described^18^. Briefly, PUL promoters amplified using Q5 Master Mix (NEB) were combined with pBolux^18^ linearized with BamHI-HF and SpeI-HF (NEB) and NEBuilder Master Mix (NEB). Reactions were incubated for 1 hour at 50°C before dialyzed on a 0.025 µM dialysis membrane against water and subsequent electroporation into competent *E. coli* S17-1. All plasmids were verified by Sanger sequencing.

### Engineering Bt mutants

Indicated *Bt* genomic deletions were generated using pEXCHANGE-*tdk* plasmids (Supplementary Table 5) harboring flanking sequences, using amplicon-specific primers (Supplementary Table 6), as previously described^48^. In short, pEXCHANGE constructs were introduced via di-parental mating and chromosomal integration was validated by PCR. Parent strains were counter-selected on solid media containing 200ng/mL 5-fluoro-2-deoxyuridine (DOT Scientific). All strains were validated using linear amplicon sequencing.

### Bt reporter array production

All 91 reporter *Bt* strains described were cultured in TYG to stationary phase in a distinct position of a 96 deep-well plate (Supplementary Table 2). Each strain was combined with glycerol to a 10% final concentration, and 5 µL of the resulting mixture was dispensed into a sterile 96-well plate using a Mini-96 (Integra), covered with adhesive sterile foil, and placed on dry ice before ultimate storage at −80°C.

### Bt reporter assays

One replicate array plate was thawed and combined with pre-reduced TYG and cultured overnight at 37°C under anaerobic conditions. The following day, each strain was diluted 50-fold into minimal media containing 0.5% galactose and cultured to mid-logarithmic phase before centrifugation at 2,204 x *g* to pellet cells, resuspension into 2x minimal media containing 0.2% galactose and distribution into a sterile 384-well white, clear bottom microplate (Corning) containing equal amounts of 0.8% carbon. Absorbance and bioluminescence were measured using an Infinite M200 Plex (Tecan) every 15 minutes over 18 hours.

### Biologically derived glycan mixture preparation

Solid material was homogenized in 6% bleach (PureBright) and allowed to stir for 30 minutes at room temperature before addition of 1:100 volumes of formic acid were combined and stirred for 5 additional minutes at room temperature. Reactions were centrifuged at 7200 x *g* for 10 minutes and the resulting supernatant was combined with bleach to a final concentration of 0.2% and stirred for 16 hours at room temperature before addition of equal volumes of formic acid and stirring for an additional 5 minutes. Reactions were centrifuged at 7200 x *g* for 10 minutes and the resulting supernatant was sterilized using a 0.22 µm vacuum filter (Millipore). The resulting material was dialyzed with ultrapure H_2_O using 1KDa cutoff dialysis membrane for 3 days before lyophilization.

### Human material collection

The Pennsylvania State University College of Medicine Institutional Review Board approved this study (IRB Protocol No. HY98-057EP-A). Prior to colectomy, patients gave informed consent to have surgically resected tissue collected and banked into the Carlino Family Inflammatory Bowel and Colorectal Disease Biobank. Diagnoses were confirmed using colonoscopy and biopsy. Surgical specimens were immediately transferred from the operating room to the surgical pathology lab, where the tissue was examined by a pathologist and surgeon. The samples were then transported to the research laboratory on ice for further processing. Mucosa was scraped from approximately 1 inch tissue sections and stored at −80°C until samples were weighed, pooled, and subjected to glycan extraction.

### Protein expression and purification

SGBPs and GHs were recombinantly expressed as previously described^47^. Briefly, inserts encoding each enzyme were amplified from *B. thetaiotaomicron* VPI-5482 genomic DNA and cloned into pT7-7-N6H4A^18^ linearized with NotI-HF and HindIII-HF using NEBuilder Hi-Fi Master Mix (NEB). The resulting plasmids were introduced into *E. coli* S17-1 and verified by Sanger sequencing. Plasmids were introduced into BL21 and plated on selective media. A single colony was inoculated into Luria Bertani broth (BD) and cultured to mid-logarithmic phase before 50 µM IPTG addition and subsequent incubation for 4 hours at 30°C. Pelleted cells were lysed in Lysis Buffer (20 mM Tris (pH, 8) and 100 mM sodium chloride) and N-terminally hexa-histidine-tagged proteins were recovered following co-incubation with Ni^2+^-NTA resin (Thermo) and elution with Lysis Buffer containing 25 mM Histidine. Recovered proteins were buffer exchanged in 10 mM Tris (pH, 7.4) containing 10% glycerol using appropriate molecular weight cut-off centrifugal concentrators (MilliporeSigma). Protein concentrations were determined using a BCA protein quantification kit (Thermo).

### SGBP-mediated glycan isolation

Glycans were recovered from mixtures using recombinant SGBPs as previously described^18^. Briefly, an *Ec* strain BL21 cell lysate expressing N-terminal hexa-his-tagged SGBP was combined with 0.5% glycan and 1.0 mL Ni^2+^-NTA resin (Thermo) and incubated overnight at 4°C with rocking. The mixture was packed into a 3 mL gravity flow column and washed with 20 mM Tris (pH=8) and 100mM Sodium Chloride 8. Protein-glycan complexes were eluted with 25 mM histidine (MilliporeSigma), treated with proteinase K before incubation at 65°C for 2 hours. The reaction was combined with 3x volume of 100% ethanol incubated overnight at 4°C with rocking and centrifuged for 15 minutes at 21,291 x *g*. The pelleted material was dried under filtered air, resuspended in water and combined before lyophilization.

### qPCR

Stationary phase cultures were diluted 50-fold into pre-reduced minimal media containing the 0.5% galactose and incubated to mid-logarithmic phase (OD_600_ = 0.45 - 0.65). 1.0 mL of culture was pelleted by centrifugation, immediately placed on dry ice, and stored at −80°C Remaining cultures were pelleted by centrifugation at 7,200 x *g* for 3 minutes and resuspended in pre-reduced 2X minimal media containing no carbon. Equal amounts of resuspended culture and carbon were combined for a final concentration of 0.5% carbon and incubated for 60-minutes before 1.0 mL of culture was pelleted by centrifugation, immediately placed on dry ice, and stored at −80°C. RNA was isolated using the RNeasy Mini Kit (QIAGEN), quantified by absorbance, and 1 µg was converted to cDNA following addition of the Superscript IV VILO Master Mix with ezDNAse (Invitrogen) per the manufacturer’s directions. Transcript abundances were measured as previously described^47^ using a QuantStudio5 (ThermoFisher) and PowerUp SYBR Green Master Mix (ThermoFisher) with amplicon specific primers (Supplementary Table 3). The 16S *rRNA, rrs,* was used as the reference gene for all qPCR experiments.

### In vivo competitive fitness of B. thetaiotaomicron strains

All animal experiments were performed in accordance with protocols approved by Penn State Institutional Animal Care and Use Committee. 8-12 week old, mixed-gender, germ-free C57/BL6 mice were maintained in flexible plastic gnotobiotic isolators with a 12-hour light/dark cycle and provided a standard, autoclaved mouse chow (LabDiet, 5021) *ad libitum*^47^. Mice were randomly distributed into 2 groups, which were supplied either autoclaved drinking water or 0.5% yeast extract glycans, 2 days prior to gavage with 10^8^ CFU of each indicated strains suspended in 200 μL of phosphate-buffered saline. Input (day 0) abundance of each strain was determined by counting colony forming units following plating on solid media. Fresh fecal pellets were collected at the indicated days and genomic DNA was extracted as described previously^23^. The abundance of each strain was measured by qPCR, using barcode-specific primers (Supplemental Table 2) as described previously^47^.

### Monosaccharide Composition Analysis

Trifluoroacetic acid was added to the indicated glycan preparations to a final concentration of 2N and heated to 100°C for 4 hours. Reactions were cooled to room temperature, dried under filtered air, and washed twice with 50% isopropanol before being dried to completion. Reactions were resuspended in 0.2 mL of deionized water and analyzed by HPAEC-PAD.

### CAZyme activity

YEG was combined with 200 nM GH20^PUL56^ to a final concentration of 1.0% in 20 mM Tris (pH=8), incubated for 4 hours at 37 °C, and analyzed by HPAEC-PAD. Mannobioses were combined with 200 nM of each PUL71 GH in 20 mM Tris (pH=8) and 1 µM calcium chloride.

### Software and statistics

Data collection and curation was done in Microsoft Excel. All data was plotted in GraphPad Prism, except for heatmaps and PCoA, which were generated using ggplot2 in RStudio. Statistical analyses were calculated in Prism. For all experiments, N denotes individual biological replicates across at least two independent experiments.

### Resource Availability

All PUL reporter plasmids are available on addgene.com and a complete *Bt* reporter array will be supplied upon request. All other *Bt* and *Ec* strains used in this work are available upon request.

## Supplementary Information

**Supplementary Table 1.** RNA-seq analysis of *Bt* strains expressing constitutively active PUL sensors.

**Supplementary Table 2.** *Bt* PUL reporter plasmids.

**Supplementary Table 3.** PUL similarity across *Bacteroides* species.

**Supplementary Table 4. Carbohydrates examined in this study.**

**Supplementary Table 5. Strains and plasmids used in this study.**

**Supplementary Table 6. Oligonucleotides used in this study.**

## Acknowledgements

We thank Walter Koltun, Leonard Harris, Troy Ott, Wesley Raup-Konsavage, and Hannah Valensi for assistance producing glycan extracts. We thank Biswa Chourdry and Richard Helm for assistance with carbohydrate compositional assistance. Additionally, we thank Jordan Bisanz for assistance with animal experiments and useful discussions. This project is funded, in-part by National Institutes of Health Grants GM147178 and DK132711 to G.E.T. and a grant with the Pennsylvania Department of Health using Tobacco CURE Funds. The PA Department of Health specifically disclaims responsibility for any analyses, interpretations or conclusions. This work was supported by the Carlino fund to The Penn State Colorectal Disease Biobank.

---

**Extended Data Fig. 1.**
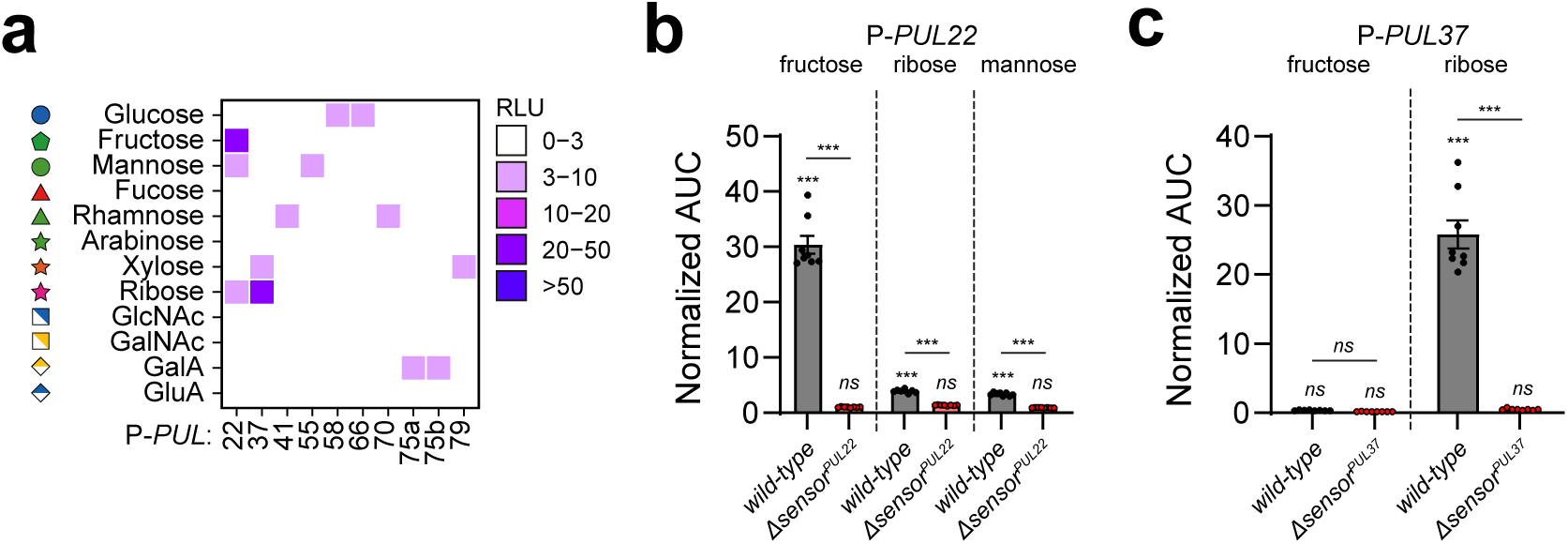
Monosaccharide-responsive PUL reporters exhibit extraordinary sensitivity. **a,** A heatmap displaying normalized bioluminescence responses to monosaccharides. N≥2. **b-c**, Normalized biolumi-nescence from *wild-type* (black) or corresponding *sensor*-deficient (red) *Bt* strains harboring (**b**) P-*PUL22* or (**c**) P-*PUL37* supplied the indicated monosaccharides. N=8, error is SEM. *P*-values were calculated using 2-way ANOVA with Fisher’s LSD test. *** indicates values < 0.001 and *ns* > 0.05.

**Extended Data Fig. 2.**
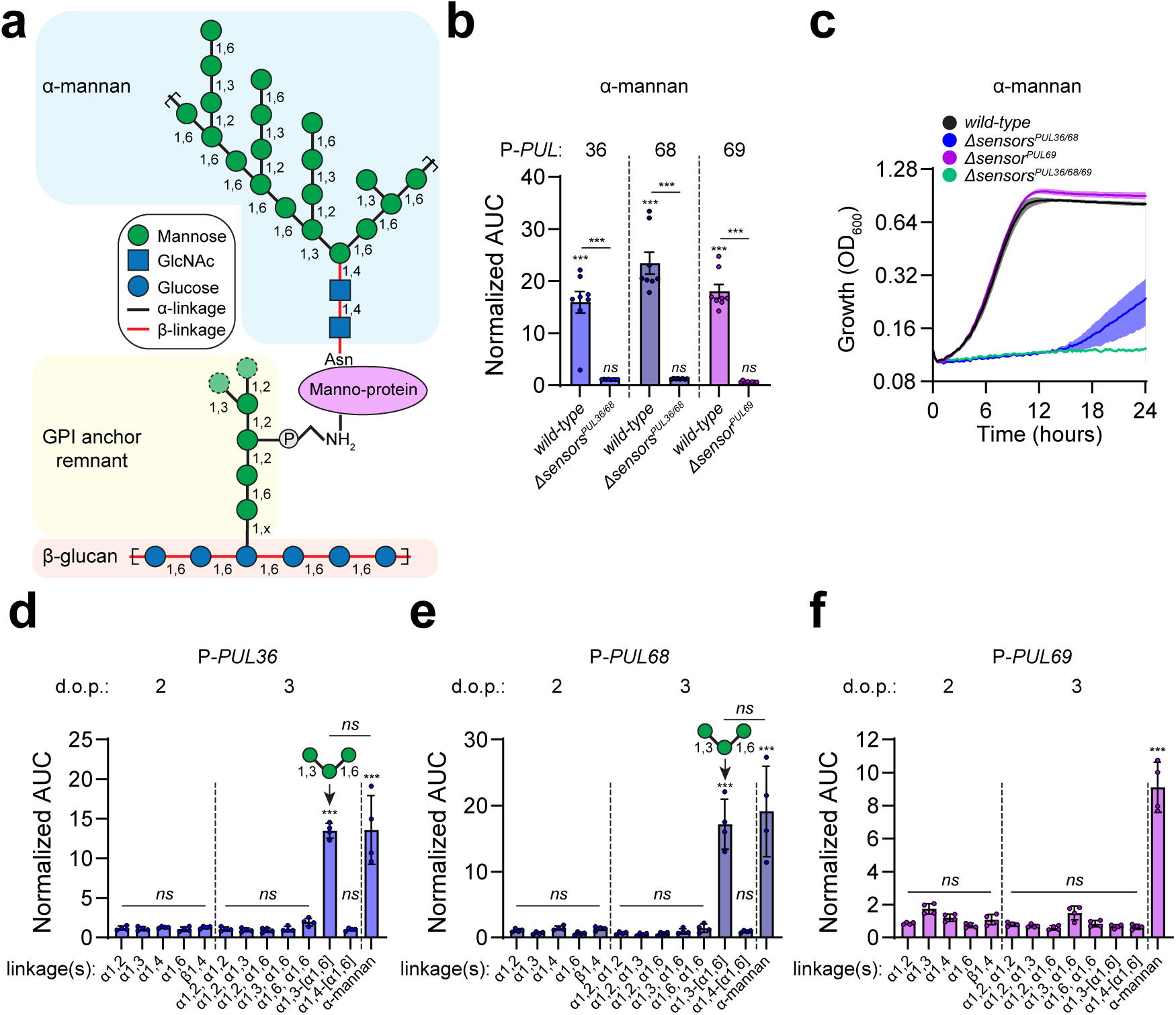
*Bt* mannan PUL reporters detect structurally distinct manno-oligosaccharides. **a,** Cartoon representation of yeast cell wall glycan structures containing mannoproteins decorated with α-mannan tethered to β 1,6-glucan by a GPI-anchor remnant glycan. **b**, Normalized bioluminescence from *wild-type*, *sensor*^PUL36/68^-, or *sensor-* ^PUL69^-deficient *Bt* strains harboring P-*PUL36, P-PUL68*, *or* P-*PUL69* supplied α-mannan and normalized by responses from isogenic strains harboring p*Bolux*. N=8, error is SEM. **c,** Growth of strains described in (**b**) supplied α-mannan as a sole carbon source. N=8, error is SEM. **d-f,** strains harboring (**d**) P-*PUL36*, (**e**) P-*PUL68*, or (**f**) P-*PUL69* were supplied the indicated manno-di-, tri-saccharides, or poly-saccharides. n=4, error is SD. For panels **b,d-f,** *P*-values were calculated using 2-way ANOVA with Fisher’s LSD test. *** indicates values < 0.001 and *ns* > 0.05.

**Extended Data Fig. 3.**
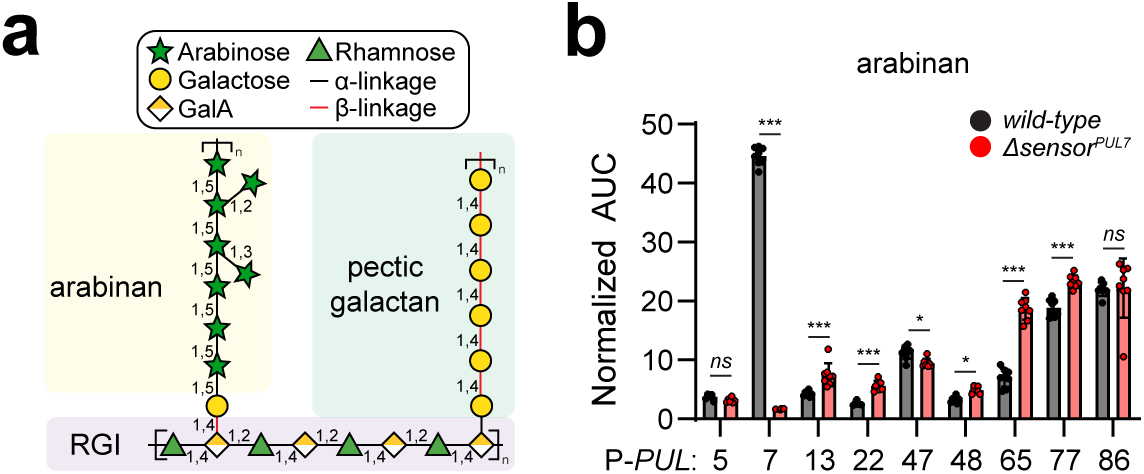
*Bt* PUL reporters recognize discreet glycan moieties from plant pectins. **a,** Cartoon representation of plant pectin structures including arabinan (PUL7), rhamnogalacturonan I (RGI, PUL77), and pectic galactan (PUL86). **b**, Normalized bioluminescence from *wild-type or Δsensor*^PUL7^ *Bt* strains harboring the indicated PUL reporters supplied arabinan and normalized by responses from a strain harboring p*Bolux*. n=8, error is SEM. *P*-values were calculated using 2-way ANOVA with Fisher’s LSD test and * indicates values <0.05, *** < 0.001, and *ns* > 0.05.

**Extended Data Fig. 4.**
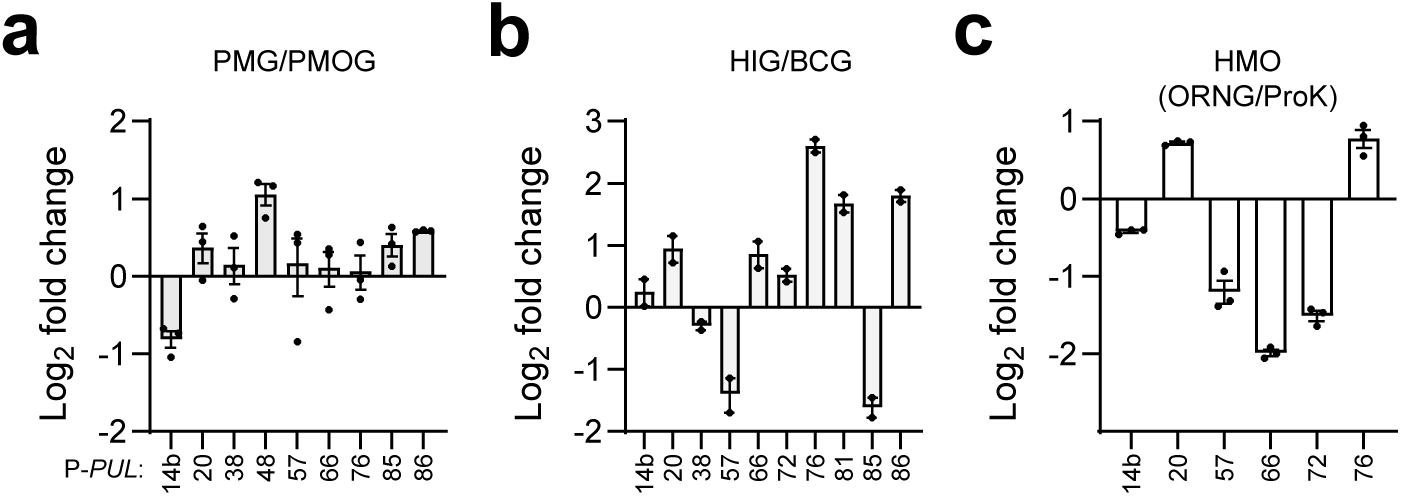
Extraction- and species-specific reporter responses. **a-c,** Log2 fold differences between reporter responses from relevant strains supplied (**a**) porcine gastric mucin prepared using ORNG (PMG) or β-elimination (PMOG), (**b**) fresh human (HIG) and boar (BCG), and (**c**) human milk oligosaccharides (HMO) prepared using ORNG or enzymatic treatment (ProK). N≥2, error is SD.

**Extended Data Fig. 5.**
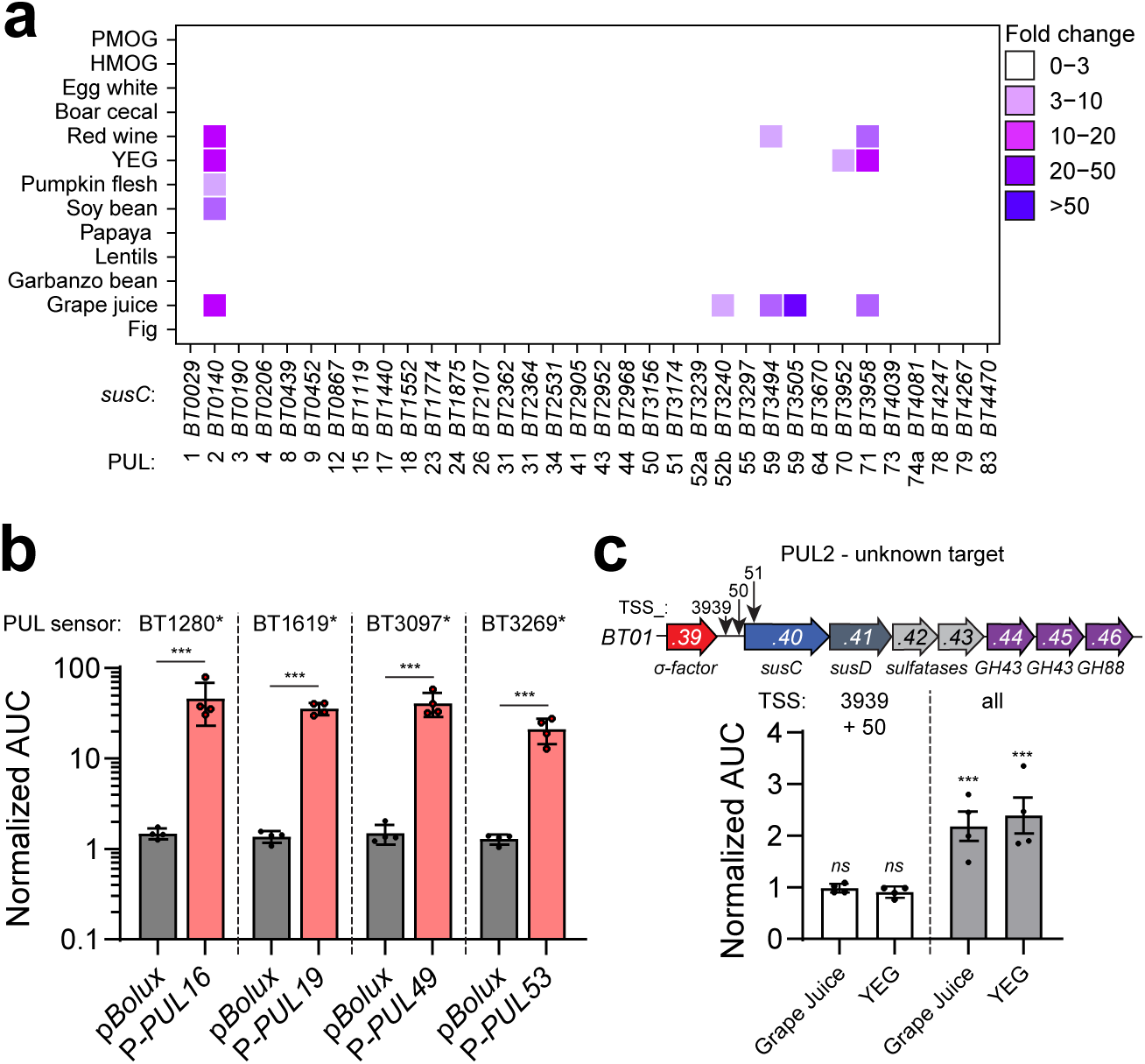
PUL expression is faithfully reported. **a,** Heatmap depicting the indicated *susC* transcript measurements from *Bt* cultures supplied the indicated glycan mixtures. N=3. **b,** Bioluminescence from strains expressing the indicated constitutively active PUL sensor and either p*Bolux* (black) or the corresponding PUL reporter (red) normalized by strains harboring an empty vector. N=4, error is SEM. **c,** Cartoon of PUL2 (top) and normalized bioluminescence (bottom) from 2 different PUL reporters containing distinct TSS combinations supplied grape juice or yeast extract glycans (YEG). N=4, error is SEM. For panels **b,c**, *P*-values were calculated using 2-way ANOVA with Fisher’s LSD test. *** indicates values < 0.001 and *ns* > 0.05.

**Extended Data Fig. 6.**
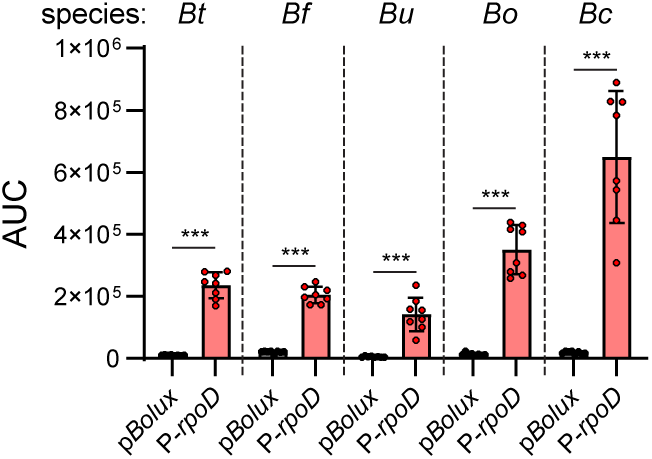
Various *Bacteroides* species can be converted into glycan biosensors. Normalized bioluminescence values from *Bt*, *B. fragilis* (*Bf*), *B. uniformis* (*Bu*), *B. ovatus* (*Bo*), and *B. cellulosilyticus* (*Bc*) type strains harboring either p*Bolux* (black) or a plasmid containing the corresponding *rpoD* promoter (red). N=8, error is SEM. *P*-values were calculated using 2-way ANOVA with Fisher’s LSD test. *** indicates values < 0.001.

**Extended Data Fig. 7.**
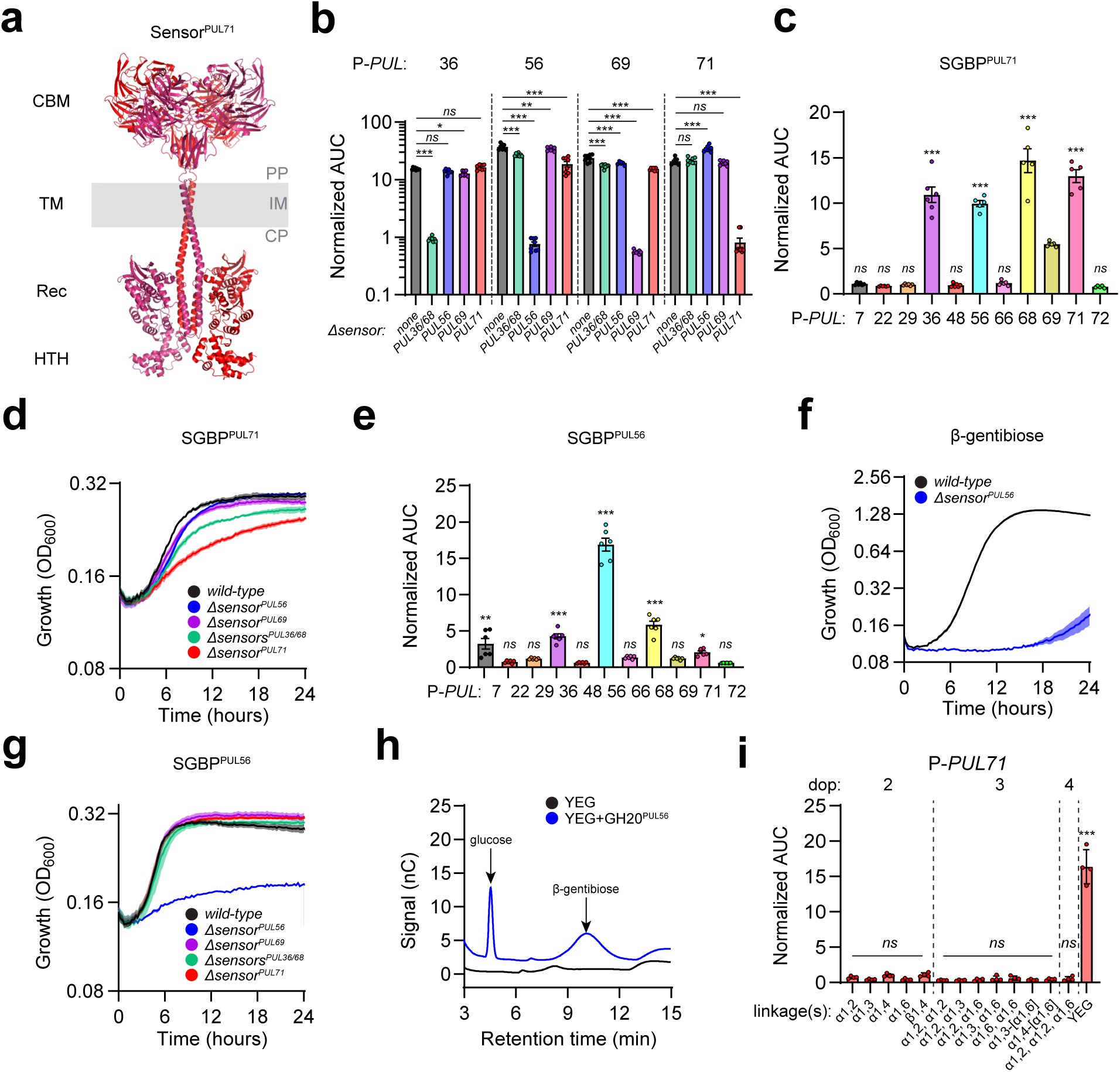
PUL reporters demonstrate a unique assemblage of distinct yeast glycans. **a,** Alpha-fold structure of Sensor^PUL71^ (BT3957) with labels indicating the periplasmic (PP) carbohydrate-binding module (CBM), the transmembrane (TM) region in the inner membrane (IM), and the cytoplasmic (CP) receiver (Rec) and helix-turn-helix (HTH) domains. **b,** Normalized AUC from *wild-type Bt* or the indicated sensor-deficient strains harboring P-*PUL36*, P-*PUL56*, P-*PUL69*, or P-*PUL71* supplied YEG. **c&e,** Normalized AUC from YEG-activated reporters supplied co-purified material following incubation of YEG with recombinant (**c**) SGBP^PUL71^ or (**e**) SGBP^PUL56^. N=6, error is SEM. **d&g,** Growth of *wild-type Bt* or the indicated sensor-deficient strains supplied co-purified material following incubation of YEG with recombinant (**d**) SGBP^PUL71^ or (**g**) SGBP^PUL56^. N=3, error is SEM. **f,** Growth of *wild-type Bt* or *Δsensor^PUL^*^56^ supplied β-gentibiose. N=8, error is SEM. **h,** HPAEC-PAD analysis of YEG treated with GH20^PUL56^. **i,** Normalized bioluminescence from a strain harboring P-*PUL71* supplied various di- and oligo-mannosaccha-rides or YEG. N=4, error is SEM. For panels **b**, **c**, **e**, and **i**, *P*-values were calculated using 2-way ANOVA with Fisher’s LSD test. * indicates values < 0.05, ** < 0.01, *** < 0.001 and *ns* > 0.05.

**Extended Data Fig. 8.**
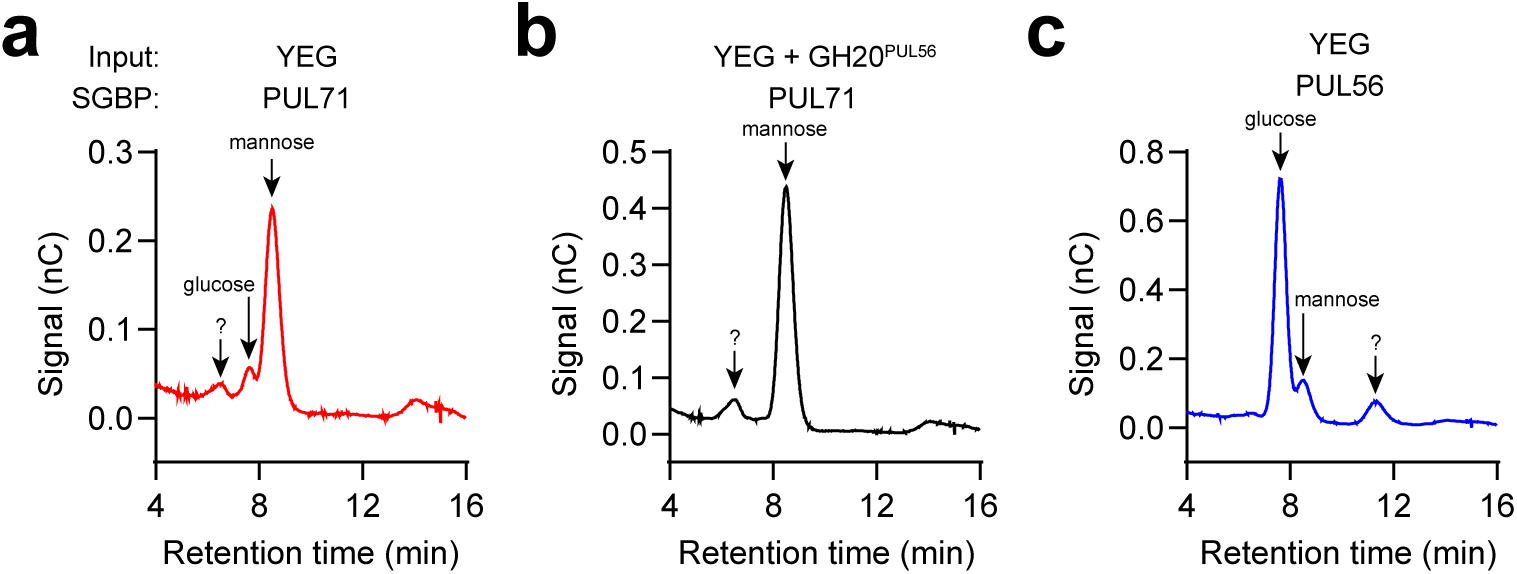
Monosaccharide composition analysis of SBGP-isolated glycans. **a-b,** HPAEC-PAD analysis of monosaccharides released from glycans that co-purified with SGBP^PUL71^ (**a**) before or (**b**) after YEG treatment with GH20^PUL56^. **c,** HPAEC-PAD analysis of monosaccharides released from glycans that co-purified with SGBP^PUL56^ following incubation with YEG.

**Extended Data Fig. 9.**
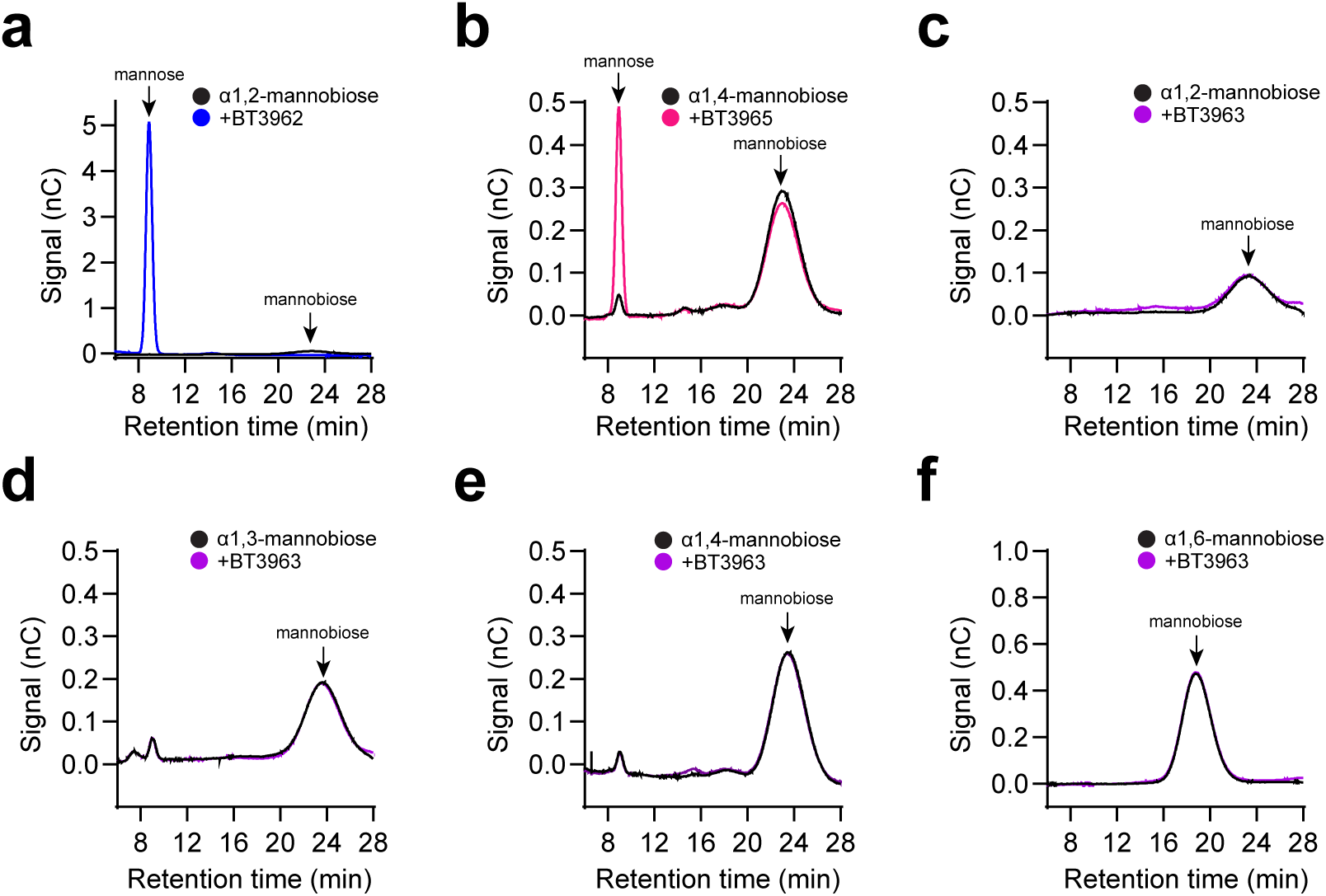
PUL71 GH specificity indicates novel glycan structure. **a-f,** HPAEC-PAD analysis of reactions containing (**a**) α1,2-mannobiose alone (black) or incubated with BT3962 (blue), (**b**) α1,4-mannobiose alone (black) or incubated with BT3965 (pink), or (**c**) α1,2-mannobiose, (**d**) α1,3-mannobiose, (**e**) α 1,4-mannobiose, or (**f**) α1,6-mannobiose alone (black) or incubated with BT3963 (purple).

