## Supplementary Table 1 for "A high-throughput microbial glycomics platform for prebiotic development"

**Supplementary Table 1. RNA-seq analysis of Bt strains expressing constitutively active PUL sensors**

| | | | Constitutively active sensor: | PUL16 | | PUL19 | | PUL49* | | PUL53* | | PUL57* | |
|---|---|---|---|---|---|---|---|---|---|---|---|---|---|
| PUL | Target glycan | Sensor Class | PUL gene (BTxxxx): | $Log_{10}$ *P*-value | Fold Change | LogP | Fold Change | LogP | Fold Change | LogP | Fold Change | LogP | Fold Change |
| 1 | Unknown | Unknown | 0028 | 0.240413297 | -0.597110129 | 0 | -0.325300311 | 0.000202759 | -0.091650795 | 0.033023423 | 0.183627938 | 0 | -1.2127687 |
| | | | 0029 | 2.903685204 | -0.911220866 | 0.13622255 | -0.294110283 | 0.004957298 | 0.17017593 | 0.167869959 | 0.17411654 | 0.726411954 | -0.419239313 |
| | | | 0030 | 1.321690196 | -0.684867528 | 0.02416949 | -0.093056341 | 0.000202759 | 0.06046375 | 0.052149942 | 0.080919871 | 0.442027766 | -0.352767312 |
| 2 | Unknown | ECF | 0139 | 1.16033676 | -2.82976726 | 0 | -1.753123975 | 0.101367857 | -2.118290848 | 0 | -3.010246716 | 0 | -2.369003009 |
| | | | 0140 | 0.598666251 | -0.277913126 | 0.258448928 | -0.422199505 | 0.000202759 | -0.083042988 | 0.157334119 | -0.171107499 | 0.294861869 | -0.268320483 |
| | | | 0141 | 0.391776843 | -0.286163411 | 0.001084382 | -0.011776619 | 0.024951395 | 0.333779399 | 0.278678395 | 0.332357832 | 0.14119382 | 0.202221247 |
| | | | 0142 | 0.011615989 | 0.013739384 | 0.039756938 | 0.217640233 | 0.101367857 | 0.629001709 | 0.027929047 | 0.066019815 | 0.195414364 | 0.318437842 |
| | | | 0143 | 0.228655727 | 0.153223083 | 0.062786228 | 0.240637886 | 0.006518177 | 0.245742497 | 0.356137358 | 0.330063418 | 0.111323577 | 0.167363399 |
| | | | 0144 | 0.167296875 | -0.693831391 | 0 | -2.286406141 | 0.12372202 | 1.739273393 | 0 | -1.483659685 | 0 | 0.639426409 |
| | | | 0145 | 0.320096047 | 0.587461865 | 0 | 0.452134019 | 0.075935742 | 0.9552435 | 0.285172467 | 0.737398006 | 0 | 0.913966511 |
| | | | 0146 | 1.321158351 | 0.711187958 | 2.5969E-05 | 0.005544619 | 0.022858763 | 0.390106216 | 0.0968992 | 0.161361765 | 0.913533851 | 0.681775473 |
| | | | 0147 | 3.885712689 | 0.52651739 | 0.047962043 | 0.120569024 | 0.000202759 | 0.13473916 | 4.590066877 | -0.503293397 | 0.717077397 | 0.181919761 |
| 3 | Unknown | ECF | 0188 | 0.064764735 | 0.037059153 | 0.131721647 | -0.241924155 | 0.000202759 | 0.004638509 | 0.145183382 | -0.138193417 | 0.041836818 | 0.049518469 |
| | | | 0189 | 0.218742502 | 0.190211054 | 0.057429347 | 0.257295445 | 0.000202759 | 0.172249149 | 0.095780646 | 0.173054274 | 0.09971995 | 0.199078496 |
| | | | 0190 | 0.090154279 | -0.044336149 | 0.022408738 | 0.067265258 | 0.000202759 | 0.03649599 | 0.489316487 | -0.28113793 | 0.310921151 | -0.206004685 |
| | | | 0191 | 0.272611676 | -0.225148127 | 0.091477514 | 0.314670636 | 0.000202759 | 0.169017398 | 0.04358834 | -0.103671751 | 0.052710278 | 0.104927988 |
| | | | 0192 | 0.041743616 | -0.029406165 | 0.097677999 | 0.266391952 | 0.000202759 | 0.101693833 | 0.115406393 | -0.135097975 | 0.278760936 | -0.297478823 |
| | | | 0193 | 0.089887015 | 0.096737367 | 0.246136177 | 0.572744578 | 0.000202759 | 0.183989588 | 0.011215592 | 0.028856758 | 0.005401152 | -0.011858318 |
| | | | 0194 | 0.174032665 | -0.142888862 | 0.010036094 | -0.071674673 | 0.019873149 | -0.338956832 | 0.369645869 | -0.454513617 | 0.262604346 | -0.35244258 |
| | | | 0195 | 0.213784238 | 0.136461894 | 0.002406745 | 0.025420062 | 0.000202759 | -0.167302412 | 0.114013956 | -0.145345994 | 0.087945687 | -0.128854777 |
| | | | 0196 | 1.109398722 | -1.180626691 | 0 | -0.77446506 | 0.040729453 | -0.777779691 | 0.065073717 | -0.187348702 | 0 | -0.643694367 |
| 4 | Unknown | Unknown | 0206 | 0.085213538 | -0.180466365 | 0.46350024 | 0.668615481 | 0.146615251 | -0.722586535 | 0.752958366 | -1.320706276 | 0.739983604 | 0.625179062 |
| | | | 0207 | 0.073076758 | -0.095476256 | 0.287580336 | 0.672311934 | 0.2279282 | -1.057191678 | 0.526711439 | -1.469696115 | 0.399601339 | 0.551136082 |
| | | | 0208 | 0.687717757 | -0.742694808 | 0.028406201 | 0.221568194 | 0.119882068 | -1.067415717 | 1.440775561 | -1.684894976 | 0.036998929 | -0.146150007 |
| | | | 0209 | 0.882588761 | -0.68674837 | 0.225652428 | 0.489412568 | 0.047316364 | -0.648636556 | 2.583970264 | -1.686991621 | 0.17002537 | 0.276401113 |
| | | | 0210 | 0.297070123 | -0.47818214 | 0.051612643 | 0.277461946 | 0.095493212 | -0.708804263 | 0.572034025 | -1.304620363 | 0.095958072 | 0.215823941 |
| | | | 0211 | 0.166016822 | -0.167294502 | 0.354319316 | 0.704431137 | 0.022858763 | -0.520953748 | 1.375190706 | -1.152529223 | 0.442491578 | 0.565561071 |
| | | | 0212 | 1.302919154 | -0.682646842 | 0.33083124 | 0.535950369 | 0.035411607 | -0.430326765 | 0.733315257 | -0.843899658 | 0.354873776 | 0.374210407 |
| | | | 0213 | 0.235810879 | -0.247462699 | 0.102007838 | 0.467940265 | 0.000202759 | -0.256779675 | 1.01647833 | -1.018999677 | 0.160925295 | 0.278457366 |
| | | | 0214 | 0.110642749 | -0.186069895 | 0 | 0.614609178 | 0.000202759 | -0.020041176 | 0.263265973 | -0.768301928 | 0.352833732 | 0.696062086 |
| 5 | Arabinogalactan | HTCS | 0262 | 0.121690298 | 1.269938144 | 0 | 1.159413094 | 0.000202759 | 1.044206356 | 0 | 1.11419199 | 0 | 3.206336626 |
| | | | 0263 | 0.21777142 | 0.481482534 | 0 | -1.539978261 | 0.000525887 | 0.474971384 | 0.002041448 | -0.015841592 | 0 | -0.062099666 |
| | | | 0264 | 0.031180252 | 0.058239606 | 0 | -0.362941326 | 0.000202759 | 0.054805368 | 0.04071404 | -0.16402019 | 0 | 0.174890839 |
| | | | 0265 | 0.032306892 | -0.072304679 | 0 | 0.003762755 | 0.000202759 | 0.394154857 | 0.003871216 | 0.019269711 | 0 | 0.692957718 |
| | | | 0266 | 0.043572169 | 0.075904234 | 0 | 0.245519036 | 0.000202759 | 0.251319281 | 0.456756804 | 0.76508352 | 0.276393065 | 0.613425472 |
| | | | 0267 | 8.139863988 | 1.426196351 | 2.083478675 | 1.275325052 | 4.725842151 | 1.450082152 | 0.677611064 | 1.159066624 | 7.677780705 | 1.637000602 |
| | | | 0268 | 10.08886662 | 1.555800141 | 5.484126156 | 1.566486264 | 15.20065945 | 1.853949145 | 4.436518915 | 1.540317362 | 15.75202673 | 1.823176655 |
| | | | 0269 | 11.20326885 | 1.991615996 | 6.194499142 | 1.933335191 | 11.75696195 | 2.224135041 | 3.196441834 | 1.797116571 | 8.37675071 | 2.054601046 |
| | | | 0270 | 10.67542418 | 2.420852071 | 9.245651664 | 2.506756782 | 9.213248578 | 2.331138468 | 2.548452415 | 2.007950734 | 5.343901798 | 1.977615752 |
| | | | 0271 | 6.549728101 | 1.836860688 | 4.363512104 | 1.892709784 | 4.844663963 | 2.039143228 | 3.654455722 | 1.797575832 | 4.815308569 | 1.89966288 |
| | | | 0272 | 23.89805197 | 1.941064033 | 11.70553377 | 1.965698176 | 16.16621563 | 1.980565947 | 1.330462606 | 1.516715978 | 12.71219827 | 1.64019241 |
| | | | 0273 | 18.18287311 | 2.277411851 | 10.86646109 | 2.288860902 | 18.57675413 | 2.428886955 | 4.596879479 | 1.688358846 | 9.100726813 | 1.812617837 |
| | | | 0274 | 5.287899193 | 1.490038274 | 1.521925112 | 1.182525502 | 2.527388632 | 1.359791014 | 1.138317512 | 1.094697743 | 1.952009157 | 1.259498674 |
| | | | 0275 | 16.0783953 | 1.701549364 | 14.50863831 | 1.835928239 | 11.76447155 | 1.641269785 | 0.856447673 | 1.254893791 | 6.342944147 | 1.279352811 |
| | | | 0276 | 15.60757278 | 1.790945289 | 14.15490196 | 1.923470262 | 15.47495519 | 1.922013125 | 4.206209615 | 1.478484742 | 8.42136079 | 1.415882241 |
| | | | 0277 | 12.06238453 | 1.665924511 | 15.45222529 | 2.080575044 | 9.484126156 | 1.701089883 | 1.072898849 | 1.378776378 | 5.311580178 | 1.471071951 |
| | | | 0278 | 7.503766004 | 1.551726769 | 9.007446482 | 1.887770285 | 8.140261434 | 1.671957135 | 3.92756995 | 1.424797699 | 5.41453927 | 1.380751141 |
| | | | 0279 | 0.815220127 | 0.743159566 | 1.502433582 | 1.550955921 | 0.050849732 | 0.700802966 | 0.204706381 | 0.500489914 | 0.603190053 | 0.880626718 |
| | | | 0280 | 0.472154639 | 0.302799545 | 0.039914557 | 0.197910623 | 0.00129528 | 0.209276167 | 0.239214212 | 0.289148544 | 0.201569628 | 0.287117457 |
| | | | 0281 | 0 | 0.11885581 | 0.002741709 | -0.028586588 | 0.000202759 | 0.075957054 | 0.125413292 | 0.156166055 | 0.017105142 | 0.029345554 |
| | | | 0282 | 0.562196726 | -0.928611874 | 0 | -0.697384922 | 0.008620635 | -0.762898074 | 0.024564735 | 0.096743815 | 0 | -0.245482635 |
| | | | 0283 | #NUM! | 0 | 0 | 0 | #NUM! | 0 | 0 | 0 | 0 | NA |
| | | | 0284 | 1.095427915 | 0.374127967 | 1.95089E-05 | 0.00067587 | 0.000202759 | 0.088580695 | 0.265623203 | -0.229845671 | 0.10215342 | 0.113488639 |
| | | | 0285 | 4.063586693 | 0.76098515 | 2.435829456 | 0.864292635 | 4.725842151 | 1.061616199 | 0.404800447 | 0.370304703 | 3.86439672 | 0.865736203 |
| | | | 0286 | 2.80368672 | 0.950420539 | 0.978450448 | 0.876157724 | 1.734446412 | 1.056578573 | 0.094922799 | 0.168392041 | 0.816282972 | 0.676673527 |
| | | | 0287 | 3.394523641 | 1.414608538 | 1.985331432 | 1.476735973 | 3.076915114 | 1.747233566 | 0.810736735 | 0.845196925 | 4.588380294 | 1.757468946 |
| | | | 0288 | 11.97245749 | 1.187955115 | 6.585026652 | 1.109558142 | 7.671620397 | 1.494400653 | 3.418016227 | 0.961448822 | 11.1857524 | 1.245791634 |
| | | | 0289 | 0.23552848 | -0.126411754 | 0.029046553 | 0.120810451 | 0.000202759 | 0.05107626 | 3.509701527 | 0.802500441 | 0.4589945 | 0.316295348 |
| | | | 0290 | 1.331626665 | 0.595135789 | 0.473842342 | 0.643317528 | 0.257485603 | 0.645480095 | 0.369341852 | 0.469438605 | 1.050915586 | 0.737656463 |
| | | | 0291 | 0.016664219 | 0.01551808 | 0.088475439 | 0.261345788 | 0.000202759 | 0.141095231 | 0.078589419 | 0.114085725 | 0.705121341 | 0.505664919 |
| 6 | Unknown | Unknown | 0317 | 4.171307846 | -0.622751293 | 2.989924459 | -1.061853717 | 2.911309432 | -0.762162841 | 15 | -1.337274666 | 3.906494266 | -0.523925688 |
| | | | 0318 | 2.999549436 | -0.675068558 | 2.358441815 | -1.010594003 | 0.061220885 | -0.348134696 | 6.906578315 | -0.962450798 | 1.283487102 | -0.419607558 |
| | | | 0319 | 1.094484667 | -0.512211789 | 1.328883777 | -0.971473449 | 0.047316364 | -0.388606897 | 2.98053226 | -1.07227526 | 0.101361563 | -0.128892784 |
| 7 | Arabinan | HTCS | 0360 | 0.13060476 | 0.097227493 | 0.026534872 | 0.124266847 | 0.002723978 | -0.218574671 | 0.139717915 | -0.198738101 | 0.011742117 | -0.026816618 |
| | | | 0361 | 0.058894326 | 0.054737546 | 0.065951221 | 0.258233177 | 0.000202759 | 0.01464003 | 0.396143447 | 0.40595633 | 0.011742117 | -0.029190718 |
| | | | 0362 | 0.174479798 | 0.110114173 | 0.007352477 | 0.048118778 | 0.000202759 | 0.129502776 | 0.043252873 | 0.063834458 | 0.088462562 | -0.128904035 |
| | | | 0363 | 0.858272176 | 0.487373079 | 0.17353122 | 0.436206066 | 0.16705161 | 0.602021074 | 0.127197608 | 0.199995676 | 0.275216104 | 0.373714018 |
| | | | 0364 | 0.826704615 | -0.43378428 | 0.029046553 | -0.146794142 | 0.000202759 | 0.059498014 | 0.09593723 | -0.133296268 | 0.177860026 | -0.235226018 |
| | | | 0365 | 0.670976019 | -0.433713779 | 0.05128535 | -0.234066832 | 0.001985343 | 0.220141549 | 0.078375293 | -0.132243826 | 0.105044758 | -0.183866732 |
| | | | 0366 | 0.78040425 | -0.209343615 | 0.437865006 | -0.401449053 | 0.000202759 | -0.070890682 | 0.941195408 | -0.362458506 | 0.47334291 | -0.267456482 |
| | | | 0367 | 0.617409991 | 2.549733579 | 0 | 1.455311534 | 0.000202759 | 0.082435062 | 2.010399985 | 4.731888091 | 0 | 3.435117542 |
| | | | 0368 | 3.0486858 | -0.500941338 | 0.783359121 | -0.489181217 | 0.008620635 | -0.200850777 | 0.508567886 | -0.227641483 | 1.294106683 | -0.359477811 |
| | | | 0369 | 1.194844445 | -0.349462995 | 0.542370502 | -0.477504035 | 0.000202759 | -0.034341031 | 0.007313381 | 0.010258742 | 0.180140181 | -0.150183054 |
| 8 | Unknown | Cytoplasmic | 0434 | 0.630502818 | -0.25457015 | 0.632183689 | -0.479205187 | 0.000202759 | -0.14997078 | 0.868728917 | -0.382131765 | 0.55678459 | -0.286465825 |
| | | | 0435 | 1.345504451 | -0.451091241 | 0.365151682 | -0.418376263 | 0.000202759 | -0.090816777 | 0.028528329 | -0.039192106 | 0.027517133 | -0.03947996 |
| | | | 0436 | 1.478381437 | -0.329250098 | 1.134696094 | -0.523525861 | 0.018081257 | -0.192744323 | 0.221651769 | -0.144638574 | 0.116662935 | -0.099969222 |
| | | | 0437 | 0.847700321 | -0.456872705 | 0.37354928 | -0.591102813 | 0.000202759 | -0.082248198 | 0.16540968 | -0.230117242 | 0.010760479 | -0.021524811 |
| | | | 0438 | 0.839721297 | -0.37769832 | 0.189002949 | -0.402785044 | 0.000202759 | -0.047768598 | 0.010460478 | 0.021611081 | 0.116673872 | 0.202882213 |
| | | | 0439 | 0.199914468 | -0.191273209 | 0.638258136 | -0.470276013 | 0.000202759 | 0.063544782 | 0.001663652 | -0.004751291 | 0.048032659 | 0.059280875 |
| | | | 0440 | 0.170950113 | -0.097297909 | 0.328790824 | -0.318684149 | 0.040729453 | 0.287046058 | 0.212640599 | 0.20560358 | 0.083643803 | 0.089090477 |
| | | | 0441 | 0.24549773 | 0.141370373 | 0.15722954 | -0.23470291 | 0.035411607 | 0.339106368 | 0.337968051 | 0.291725332 | 0.244143755 | 0.162388078 |
| | | | 0442 | 0.146251653 | 0.096118614 | 0.004858317 | -0.030443649 | 0.030771831 | 0.305635076 | 0.029153899 | 0.043362458 | 0.063506855 | 0.077762348 |
| | | | 0443 | 0.261769596 | 0.312765246 | 0 | -0.155124483 | 0.021897661 | 0.54919225 | 0.196602118 | 0.393033621 | 0.484378751 | 0.72820438 |
| | | | 0444 | 0.139526671 | 0.080172042 | 0.002406745 | -0.022292023 | 0.009622678 | 0.191802468 | 0.066148547 | 0.069063711 | 0.041871035 | -0.054011331 |
| | | | 0445 | 0.344028537 | 0.176067852 | 0.246136177 | -0.341483006 | 0.003466816 | 0.169159534 | 0.326024022 | 0.220382756 | 0.149325501 | 0.126707145 |
| 9 | Unknown | Unknown | 0446 | 0.042203527 | -0.066079382 | 0.031693841 | -0.187409043 | 0.062040053 | -0.511792492 | 0.167376431 | -0.239475853 | 0.013646459 | 0.033689287 |
| | | | 0447 | 0.134821585 | -0.100160877 | 0.001084382 | -0.008969899 | 0.005522707 | -0.243804775 | 0.056277448 | 0.091995885 | 0.03084631 | 0.054107611 |
| | | | 0448 | 0.158141343 | -0.122514998 | 0.047218354 | 0.216701921 | 0.000202759 | 0.031673812 | 0.454627723 | 0.405347118 | 0.170306694 | 0.221051326 |
| | | | 0449 | 0.140408035 | -0.094965148 | 0.001084382 | -0.009744574 | 0.000202759 | -0.057632628 | 0.099250163 | 0.205947016 | 0.171142454 | 0.171547145 |
| | | | 0450 | 0.629742129 | -0.288865917 | 0.03044408 | -0.157975088 | 0.000202759 | -0.154161228 | 0.404070781 | 0.261522996 | 0.1588045 | 0.156947602 |
| | | | 0451 | 1.834407131 | -0.699291887 | 0.123120774 | -0.382068363 | 0.001985343 | -0.215322626 | 0.065073717 | -0.09084597 | 0.081706947 | -0.112644325 |
| | | | 0452 | 0.826277744 | -0.638385184 | 0.46350024 | -0.581753029 | 0.101367857 | -0.376631289 | 0.038173492 | -0.047869504 | 0.153483196 | -0.137027301 |
| | | | 0453 | 1.876369906 | -0.556304268 | 0.423626118 | -0.587917315 | 0.022858763 | -0.328637066 | 0.289087113 | -0.229531362 | 0.023623375 | -0.036615282 |
| | | | 0454 | 0.348855364 | -0.205386525 | 0.23299351 | -0.431215159 | 0.132099897 | -0.409869767 | 0.112928853 | -0.111175222 | 0.432373223 | -0.295444239 |
| | | | 0455 | 1.061984174 | -0.268701024 | 0.036192153 | 0.09100996 | 0.437579198 | -0.43430117 | 1.166150957 | -0.316100908 | 0.551850996 | 0.227002119 |
| | | | 0456 | 1.301209553 | -0.384273882 | 0.029046553 | 0.091942167 | 0.144280548 | -0.366231034 | 0.325222429 | -0.180965786 | 1.301405875 | 0.412365113 |
| | | | 0457 | 0.756109791 | -0.249893356 | 0.136903401 | 0.219410784 | 0.123624592 | -0.322178049 | 0.200331567 | -0.140714434 | 1.55496868 | 0.484187499 |
| | | | 0458 | 0.468577399 | -0.124398728 | 0.086396171 | 0.151066102 | 0.055661433 | -0.23217244 | 1.12430639 | -0.278051461 | 0.841111284 | 0.235000697 |
| | | | 0459 | 0.614212883 | -0.392076149 | 0.081905528 | 0.177705387 | 0.573176206 | -0.615896458 | 1.399402044 | -0.444532756 | 0.851805558 | 0.362813871 |

| | | | | | | | | | | | | | |
|---|---|---|---|---|---|---|---|---|---|---|---|---|---|
| | | | 0460 | 1.515999685 | -0.417313912 | 0.129213774 | 0.233269358 | 0.573176206 | -0.600976887 | 1.504854689 | -0.458298365 | 0.561597811 | 0.280933765 |
| | | | 0461 | 1.300264194 | -0.394119403 | 0.378442142 | 0.393088036 | 0.554104282 | -0.574134032 | 0.908984723 | -0.398504979 | 0.646921808 | 0.312699926 |
| 10 | Unknown | Unknown | 0483 | 0.120884723 | -0.039357633 | 1.358116032 | -0.553639732 | 0.211160447 | -0.308542938 | 2.290076629 | -0.555900691 | 7.974694135 | -0.680752244 |
| | | | 0484 | 1.759386062 | 0.268392892 | 0.33083124 | -0.27812852 | 0.000202759 | 0.010818474 | 1.430947388 | -0.374760667 | 3.502353114 | -0.439177785 |
| 11 | Unknown | ECF | 0752 | 0.866854776 | 0.226019626 | 0.02416949 | 0.068540895 | 0.000202759 | 0.01427645 | 0.001689286 | 0.001902109 | 0.082030473 | 0.066611274 |
| | | | 0753 | 0.665248041 | -0.423272757 | 0.081674782 | -0.325781534 | 0.000202759 | -0.133385534 | 0.040160385 | 0.07335871 | 0.015119444 | -0.035276747 |
| | | | 0754 | 0.172215756 | -0.078429553 | 0.390763326 | -0.386462519 | 0.002964065 | -0.14531477 | 1.128846735 | -0.428630842 | 1.61252295 | -0.529146347 |
| | | | 0755 | 0.046782802 | -0.028646707 | 0.116104328 | -0.24459941 | 0.000202759 | -0.128809093 | 1.183746434 | -0.522076619 | 1.359054136 | -0.596320369 |
| | | | 0756 | 0.513220019 | -0.221325579 | 0.018104878 | -0.059773636 | 0.000202759 | -0.078197002 | 1.499176581 | -0.565775148 | 0.477437415 | -0.302634384 |
| | | | 0757 | 0.230436653 | 0.0809778 | 0.011969108 | 0.039831575 | 0.004537328 | 0.130817687 | 0.017575703 | 0.019020911 | 0.39826892 | -0.218702336 |
| | | | 0758 | 0.043982896 | -0.045692887 | 0.05128535 | 0.215980913 | 0.062704852 | 0.429046207 | 0.099101049 | 0.166602912 | 0.551674538 | 0.497169797 |
| 12 | Unknown | Unknown | 0865 | 0 | -3.041972859 | 0 | -3.427892202 | 0 | -2.883264831 | 0 | -2.916788183 | 0 | -1.975056132 |
| | | | 0866 | 0 | -3.165148162 | 0 | -3.549503167 | 0 | -3.08282009 | 0 | -3.266822345 | 0 | -2.015466364 |
| | | | 0867 | 0 | -3.156528777 | 0 | -3.576689555 | 0 | -3.199302099 | 0 | -3.642126947 | 0 | -2.379136257 |
| 13 | Rhamnogalacturonan II | HTCS | 0981 | 0.005522287 | -0.001557509 | 0.047962043 | -0.081955822 | 0.000202759 | -0.054895305 | 0.908984723 | -0.193142604 | 0.401917211 | -0.123318585 |
| | | | 0982 | 0.408239031 | 0.656507259 | 0 | -0.970067747 | 0.000202759 | -0.277592717 | 0.007965004 | 0.038102266 | 0 | 0.037908348 |
| | | | 0983 | 0.394579357 | 0.175156371 | 0.305428651 | 0.355900582 | 0.000202759 | -0.165708531 | 0.133195081 | 0.142210549 | 0.164561955 | -0.17163091 |
| | | | 0984 | 1.37597377 | 0.6771449 | 0.530857432 | 0.717762163 | 0.056322103 | 0.4696629 | 2.583970264 | 1.162515349 | 0.352265817 | 0.476052124 |
| | | | 0985 | 1.019299652 | 0.669007023 | 0.978450448 | 1.05794414 | 0.112152493 | 0.654038265 | 2.503727962 | 1.195689521 | 1.031287902 | 0.898930921 |
| | | | 0986 | 0.313952847 | 0.184863421 | 0.031054033 | 0.133387635 | 0.024985102 | 0.309466018 | 3.654455722 | 1.157065832 | 2.057334798 | 0.952484538 |
| | | | 0987 | 4.811188414 | -0.608502854 | 1.626563296 | -0.569061268 | 0.000202759 | -0.074994688 | 2.222284855 | 0.540950304 | 0.800851737 | 0.348657103 |
| | | | 0988 | 12.06462592 | -0.75244191 | 4.216096421 | -0.698025401 | 0.003466816 | -0.170897967 | 0.040668836 | -0.038946502 | 0.021069188 | -0.021946805 |
| | | | 0989 | 0.659082617 | 0.23699347 | 0.029046553 | 0.092268665 | 0.000202759 | 0.028537488 | 0.011474642 | -0.014800088 | 0.012267406 | 0.016836337 |
| | | | 0990 | 0.33502976 | -0.228462357 | 0.008774443 | -0.058650678 | 0.000202759 | -0.037039314 | 0.158120009 | -0.208835945 | 0.018666274 | -0.037559318 |
| | | | 0991 | 0.011865028 | -0.011175584 | 0.015997086 | -0.099583299 | 0.002964065 | -0.250575205 | 0.052149942 | -0.096436706 | 0.170119655 | -0.320829761 |
| | | | 0992 | 0.049427986 | 0.045722589 | 0.005185202 | 0.052506045 | 0.000202759 | -0.104125419 | 0.101525844 | 0.158139122 | 0.03525235 | 0.076066303 |
| | | | 0993 | 0.304662605 | 0.505927018 | 0 | -0.147810195 | 4.81016E-05 | 0.000102407 | 0.016009812 | -0.077668443 | 0 | 0.071540461 |
| | | | 0994 | 0.044073496 | 0.083593746 | 0 | -0.372756694 | 0.000202759 | 0.085057035 | 0.295527296 | -0.783077771 | 0 | -0.566460617 |
| | | | 0995 | 0.147844662 | 0.129980272 | 0.097677999 | -0.418701645 | 0.145883736 | -0.678608546 | 0.424515129 | -0.512799198 | 0.082030473 | 0.169702421 |
| | | | 0996 | 0.297882593 | -0.55996799 | 0 | 0.662530314 | 0.021870103 | 0.670517467 | 0.030268172 | 0.137428748 | 0 | 0.344418507 |
| | | | 0997 | 0.290072794 | -0.448474709 | 0 | -0.654455241 | 0.000202759 | -0.232417976 | 0.193238482 | -0.521732617 | 0 | -0.779664045 |
| | | | 0998 | 0.150322567 | 0.036427157 | 0.131721647 | -0.148761892 | 0.001687904 | -0.087685982 | 1.048538963 | -0.20010127 | 0.629921161 | 0.166106611 |
| | | | 0999 | 0.012332144 | 0.004325376 | 0.117461091 | -0.159165335 | 0.000202759 | -0.074226737 | 0.108627099 | -0.068994708 | 0.772122675 | 0.238042414 |
| | | | 1000 | 0.627176182 | -0.674619632 | 0 | -0.573967685 | 0.000202759 | -0.112710195 | 0.63597808 | 0.856329965 | 0.216053366 | 0.423521815 |
| | | | 1001 | 0.305745724 | 0.444419532 | 0 | 0.385819052 | 0.260665694 | 1.033223987 | 1.19888847 | 1.317945476 | 0 | 0.727516096 |
| | | | 1002 | 0.385854234 | -0.738213439 | 0 | 0.067453437 | 0.00129528 | 0.469931365 | 0.157450595 | 0.547347935 | 0 | 0.62919939 |
| | | | 1003 | 0.071021657 | 0.061123704 | 0.029723943 | 0.144661306 | 0.000202759 | 0.067377628 | 0.166355564 | 0.229844494 | 0.384244122 | 0.460971073 |
| | | | 1004 | 1.289247286 | 0.802607161 | 0.372306225 | 0.752472835 | 0.000202759 | 0.017055008 | 0.100353145 | 0.213889573 | 0.588984702 | 0.74130118 |
| | | | 1005 | 0.945774966 | 0.807892228 | 0 | -0.029212614 | 0.000202759 | 0.29326924 | 0.158225503 | 0.352928922 | 0.75697334 | 0.978215999 |
| | | | 1006 | 0.426362005 | 0.170611277 | 0.297992593 | -0.346435558 | 0.130683092 | -0.332550327 | 2.947367995 | -0.703026977 | 0.021069188 | -0.030279111 |
| | | | 1007 | 1.229624575 | 0.200158074 | 0.262602829 | 0.269162066 | 0.101367857 | 0.225135683 | 0.243881587 | -0.122677911 | 0.344691592 | -0.132788879 |
| | | | 1008 | 3.519062192 | 0.666498227 | 0.039222445 | 0.106893997 | 0.000202759 | 0.063454652 | 3.544504947 | -0.775134716 | 1.129000759 | 0.477001423 |
| | | | 1009 | 0.044329694 | 0.047628347 | 0.030941433 | 0.180688147 | 0.000202759 | -0.113576094 | 0.096746354 | -0.180201559 | 0.175797449 | 0.329288117 |
| | | | 1010 | 0.001158306 | 0.002161806 | 0 | -1.11947714 | 0.000202759 | 0.071230254 | 0.505896464 | -1.042096834 | 0 | -0.313471449 |
| | | | 1011 | 0.471979482 | -0.808634822 | 0 | -0.917907078 | 0.000525887 | -0.476114634 | 0.409085414 | -1.072539078 | 0 | -0.743813621 |
| | | | 1012 | 0.189279682 | 0.362314009 | 0 | 1.074980643 | 0.020797091 | 0.756314317 | 0.148001628 | 0.489293332 | 0 | 0.379596131 |
| | | | 1013 | 0.448401856 | 0.424642449 | 0.004534747 | 0.069082806 | 0.023506576 | 0.450008668 | 0.182162699 | 0.328035468 | 0.013520434 | -0.04586404 |
| | | | 1014 | 0.497856029 | -1.055994299 | 0 | -0.022612451 | 6.90027E-05 | 0.003164125 | 0.024564735 | 0.14520378 | 0 | 0.226145055 |
| | | | 1015 | 0.089299187 | 0.032912915 | 0.271004767 | -0.274094206 | 0.000202759 | -0.028581382 | 0.841615592 | -0.273018451 | 0.087072133 | -0.062250981 |
| | | | 1016 | 1.275152083 | 0.32756718 | 0.066030681 | -0.135276943 | 0.000202759 | 0.05595094 | 0.575182706 | -0.275743904 | 0.394738715 | 0.230883226 |
| | | | 1017 | 0.402133513 | -0.219383213 | 0.022170845 | -0.089405844 | 0.000202759 | 0.018279269 | 0.145183382 | -0.162881088 | 0.099455291 | 0.124018374 |
| | | | 1018 | 0.021467582 | 0.02112586 | 0.073836009 | -0.293323057 | 0.000202759 | 0.080067867 | 0.028528329 | 0.057976598 | 0.069457134 | 0.130254537 |
| | | | 1019 | 0.019289031 | 0.019746534 | 0.009142892 | -0.081828931 | 0.000202759 | -0.132615538 | 0.04358834 | -0.08930054 | 0.037099704 | 0.086897619 |
| | | | 1020 | 0.056943218 | 0.086814177 | 0 | -0.370316484 | 0.004851778 | 0.379400398 | 0.149854007 | 0.319551052 | 0.166431005 | 0.387555514 |
| | | | 1021 | 1.152764395 | -1.51571727 | 0 | -1.949019678 | 0.0219888 | -0.68872503 | 0.168342665 | -0.47991565 | 0 | -0.770954583 |
| | | | 1022 | 0.108129748 | -0.105636299 | 0.049454347 | -0.25599757 | 0.000202759 | 0.11603447 | 0.171528111 | -0.386342647 | 0.036008543 | 0.085038438 |
| | | | 1023 | 0.043345191 | 0.032839599 | 0.022460809 | -0.089955679 | 0.000202759 | 0.11211377 | 0.380503592 | -0.303387285 | 0.1097465 | 0.123071964 |
| | | | 1024 | 0.359314498 | -0.546229326 | 0 | -0.101820856 | 0.000202759 | -0.026961574 | 0.176050725 | -0.465925543 | 0.006776078 | 0.019978625 |
| | | | 1025 | 0.165116825 | 0.193609467 | 0 | -0.864627417 | 0.000202759 | -0.171964858 | 0.156453695 | -0.333337454 | 0.175400579 | 0.358839645 |
| | | | 1026 | 0.438156371 | -0.763297407 | 0 | -0.985894444 | 0.000525887 | -0.462187831 | 0.516865724 | -1.175802848 | 0 | -1.431934297 |
| | | | 1027 | 0 | 0 | 0 | 0 | 0 | 0 | 0 | 0 | 0 | 1.706052383 |
| | | | 1028 | 0.204873887 | 0.210026476 | 0.029883833 | 0.210037751 | 0.030771831 | 0.464966911 | 0.137149118 | -0.288395771 | 0.208876433 | 0.40500965 |
| | | | 1029 | 0.510365553 | -0.321572824 | 0.111803586 | -0.303899999 | 0.000202759 | -0.156406474 | 0.635841052 | -0.4936719 | 0.375310889 | -0.353990486 |
| | | | 1030 | 0.330241734 | -0.312626682 | 0.088109783 | -0.390057665 | 0.004537328 | -0.307303837 | 1.764843489 | -1.379664538 | 0.734338328 | -0.855963004 |
| 14a | N-glycans | ECF | 1032 | 1.896166769 | 0.382456398 | 0.129213774 | -0.165316414 | 0.000202759 | -0.035718389 | 0.034759066 | -0.026529207 | 0.009328765 | -0.006666737 |
| | | | 1033 | 0.778772844 | 0.223329743 | 0.578428436 | -0.400666823 | 0.109659506 | -0.250796116 | 0.48530913 | -0.220333896 | 1.084287954 | -0.344383908 |
| | | | 1034 | 3.630409401 | 0.966543853 | 0.623528953 | -0.914556242 | 0.101367857 | -0.474533172 | 0.771230271 | -0.571580735 | 0.313517768 | -0.382170899 |
| | | | 1035 | 0.167603019 | 0.149990793 | 0.039756938 | -0.205545977 | 0.004851778 | 0.281144824 | 0.252840848 | -0.360381478 | 0.166443664 | -0.292190213 |
| | | | 1036 | 0.049964432 | -0.043780101 | 0.062212259 | 0.232320922 | 0.000525887 | -0.261030116 | 0.662204629 | -0.605794499 | 1.732844329 | -1.196513184 |
| | | | 1037 | 0.01920307 | -0.022077951 | 0.002741709 | -0.042279603 | 0.006518177 | -0.30815828 | 0.404265598 | -0.573713247 | 0.772122675 | -0.92297406 |
| | | | 1038 | 0.038958351 | -0.042542186 | 0.057501259 | 0.257115253 | 0.004537328 | -0.281709302 | 0.166829583 | -0.32015727 | 1.268847884 | -1.014828667 |
| | | | 1039 | 0.051375166 | -0.056934537 | 0.031693841 | 0.140735536 | 0.001985343 | -0.225931214 | 0.411724325 | -0.425817625 | 2.553945547 | -1.152244103 |
| | | | 1040 | 0.564278857 | -0.289531862 | 1.95089E-05 | -0.002185227 | 0.360623717 | -0.499978842 | 1.190173677 | -0.731177411 | 4.598599459 | -1.40488659 |
| 14b | N-glycans | ECF | 1042 | 0.308995935 | 0.169937442 | 1.01829094 | -0.674501557 | 0.000202759 | 0.038002991 | 0.614850748 | -0.36458765 | 0.105538171 | -0.221558042 |
| | | | 1043 | 0.361243663 | 0.336047231 | 0.939398157 | -1.030981365 | 0.000202759 | -0.01030333 | 0.777237447 | -0.689023419 | 0.118715289 | 0.328077877 |
| | | | 1044 | 0.725679658 | 0.653028118 | 0.073836009 | -0.454329857 | 0.071879253 | 0.698598407 | 0.305709629 | 0.486487248 | 0.4188138 | 0.792365309 |
| | | | 1045 | 1.331073472 | 0.939804933 | 0.048778119 | -0.276675446 | 0.002964065 | 0.381934576 | 0.008838497 | -0.024694997 | 0.252372021 | 0.58180002 |
| | | | 1046 | 0.800227054 | -0.642360494 | 0.248157356 | -0.664140092 | 0.019865857 | -0.553713902 | 0.037954975 | 0.187016309 | 1.479536198 | -1.269514182 |
| | | | 1047 | 1.394046658 | -0.880999789 | 0.422749717 | -0.803185975 | 0.0823077 | -0.628407279 | 0.07315814 | 0.21803086 | 2.439247743 | -1.513679155 |
| | | | 1048 | 0.605119606 | -0.616826219 | 0.390763326 | -0.916748068 | 0.257485603 | -1.006235456 | 0.001689286 | 0.008083363 | 5.152427341 | -3.516011873 |
| | | | 1049 | 2.083094462 | -1.262727936 | 1.004070651 | -1.231535127 | 0.118753099 | -0.735992872 | 0.029069294 | 0.166605527 | 3.583391109 | -2.000106388 |
| | | | 1050 | 0.404508733 | -0.454976909 | 0.00230855 | -0.037378242 | 0.000202759 | 0.200581028 | 0.398924245 | 0.690694037 | 0.241653152 | -0.522499002 |
| | | | 1051 | 0.926808407 | -0.230926267 | 0.195073704 | -0.226620175 | 0.000202759 | -0.068821517 | 0.118206908 | -0.079852981 | 1.280450899 | -0.354502726 |
| | | | 1052 | 1.367782223 | 0.31228237 | 0.086396171 | 0.174887555 | 0.722809134 | 0.505526164 | 1.968668988 | 0.455705334 | 0.8517503 | 0.360444471 |
| | | | 1053 | 0.10964578 | -0.041842183 | 0.012450752 | -0.040622234 | 0.000202759 | -0.014291637 | 0.02569838 | -0.024988442 | 0.102312713 | 0.083746191 |
| 15 | Unknown | Unknown | 1117 | 0.657517375 | -0.27774419 | 0.46350024 | -0.537141992 | 0.000202759 | 0.056855391 | 0.427317043 | 0.284218957 | 0.650381752 | 0.362879972 |
| | | | 1118 | 1.285075195 | -0.344319259 | 0.105628234 | -0.231124232 | 0.000202759 | -0.072196971 | 0.011474642 | 0.010392588 | 0.240565097 | 0.128663639 |
| | | | 1119 | 0.296189522 | -0.078542012 | 0.064227735 | -0.118170068 | 0.024985102 | 0.154634457 | 0.038713413 | -0.025071915 | 0.05693114 | 0.036916386 |
| 16 | Unknown | ECF | 1278 | 157.5793631 | 3.709630335 | 0.036192153 | -0.118690369 | 0.007393621 | 0.199151235 | 0.065073717 | -0.082699857 | 0.137446329 | -0.139535739 |
| | | | 1279 | 128.2196823 | 3.971005426 | 0.019048619 | -0.086240626 | 0.000202759 | 0.01607169 | 0.005777132 | 0.009565667 | 0.250335122 | -0.27987206 |
| | | | 1280 | 280.2422928 | 5.274354767 | 0.476127375 | -0.543673956 | 0.1208687 | -0.349160683 | 1.583495893 | -0.550775559 | 1.591918513 | -0.590810214 |
| | | | 1281 | 203.0165594 | 5.26374098 | 0.202576826 | -0.481948487 | 0.000202759 | -0.088480584 | 0.06969121 | -0.117602681 | 0.258644131 | -0.307128881 |
| | | | 1282 | 130.4323261 | 4.837831259 | 0.151191457 | -0.465206695 | 0.000202759 | -0.130836065 | 0.040307982 | -0.089268948 | 0.113183059 | -0.200616978 |
| | | | 1283 | 239.984774 | 4.983651739 | 0.016868642 | -0.085677488 | 0.000202759 | 0.156358239 | 0.207378924 | -0.227804337 | 0.245352221 | -0.284249724 |
| | | | 1284 | 264.167037 | 4.859363289 | 0.07037195 | -0.259987815 | 0.002964065 | -0.197821711 | 0.297669221 | -0.262776522 | 0.137446329 | -0.162983515 |
| | | | 1285 | 279.9835882 | 4.389197896 | 0.16086148 | -0.2444187 | 0.614060977 | -0.480013711 | 2.508892513 | -0.602100756 | 2.960133578 | -0.640118471 |
| 17 | Unknown | Unknown | 1439 | 2.883447044 | 0.434729398 | 0.005565012 | -0.026215776 | 0.000525887 | 0.113029724 | 0.835520774 | -0.274509009 | 0.038339279 | -0.034288569 |
| | | | 1440 | 2.031998545 | 0.340342228 | 0.057429347 | -0.11101943 | 0.000202759 | 0.025319642 | 1.349243151 | -0.372643307 | 0.287765596 | -0.136851772 |
| 18 | Unknown | Unknown | 1551 | 0.297982201 | 0.065793799 | 0.207458029 | 0.178465066 | 0.000525887 | 0.102720913 | 0.500658269 | 0.137035914 | 0.113699157 | -0.048760806 |
| | | | 1552 | 0.488217116 | -0.093529578 | 0.019109392 | 0.039768647 | 0.019865857 | -0.151217738 | 0.338130364 | -0.105590798 | 0.808502554 | -0.203649707 |
| | | | 1553 | 0.002473918 | 0.000706347 | 0.205046956 | 0.191020649 | 0.000202759 | -0.058059359 | 0.101525844 | -0.050108407 | 0.523857859 | -0.195196136 |
| | | | 1617 | 0.243803096 | 0.09779375 | 94.02045163 | 3.657040396 | 0.000525887 | 0.114754825 | 0.101525844 | -0.092245743 | 0.471049065 | 0.25980034 |
| | | | 1618 | 0.037615015 | 0.028040144 | 44.40450378 | 3.066552238 | 0.000202759 | -0.092407508 | 0.173567334 | -0.181074335 | 0.229566737 | 0.221014144 |

| | | | | | | | | | | | | | |
|---|---|---|---|---|---|---|---|---|---|---|---|---|---|
| 19 | Unknown | ECF | 1619 | 0.598913014 | -0.163626163 | 203.2403322 | 4.669813716 | 0.491403646 | -0.439999521 | 2.370741207 | -0.529776241 | 0.465252407 | -0.207966536 |
| | | | 1620 | 0.530061088 | -0.164863204 | 188.1420647 | 4.475126341 | 0.101367857 | -0.312582445 | 4.707743929 | -0.835079784 | 0.647791738 | -0.279804725 |
| | | | 1621 | 0.783677191 | -0.261332322 | 83.51712642 | 3.330813789 | 0.761257185 | -0.571314696 | 1.256880343 | -0.437787821 | 0.469237984 | -0.254895147 |
| | | | 1622 | 0.182143888 | -0.088081628 | 68.70114692 | 2.968688989 | 0.005314538 | -0.16266198 | 1.030762987 | -0.398218347 | 0.03546796 | -0.041076824 |
| 20 | GAG-associated glycan | HTCS | 1623 | 1.266507996 | 0.266612934 | 0.133625486 | 0.174514101 | 0.000202759 | -0.061986576 | 1.63860343 | -0.442748117 | 0.245352221 | 0.143699862 |
| | | | 1624 | 1.254024638 | -0.211582271 | 0.053599877 | -0.101861463 | 0.008620635 | -0.165031583 | 2.468015935 | -0.400815639 | 0.496889966 | -0.172368857 |
| | | | 1625 | 1.19371325 | -0.233068808 | 0.148628592 | -0.18906527 | 0.381796523 | -0.325583534 | 4.013228266 | -0.49631151 | 1.466003788 | -0.402897181 |
| | | | 1626 | 0.558841134 | -0.150427149 | 0.019109392 | -0.055247117 | 0.013169173 | -0.156615672 | 0.792017704 | -0.283651047 | 0.201569628 | 0.134656232 |
| | | | 1627 | 1.798274766 | -0.333250676 | 0.763282073 | -0.472579614 | 0.057021257 | -0.250930885 | 3.583885624 | -0.573923895 | 0.212559314 | -0.136248019 |
| | | | 1628 | 1.27628031 | -0.277413765 | 1.664551066 | -0.537512287 | 0.733793748 | -0.428183056 | 5.331614083 | -0.600247457 | 2.136028087 | -0.443482378 |
| | | | 1629 | 0.940820298 | -0.402004252 | 0.115702589 | -0.273581287 | 0.006518177 | -0.209697081 | 0.280969775 | -0.244883152 | 0.583454219 | 0.379797578 |
| | | | 1630 | 0.673381745 | -0.303021103 | 0.029046553 | 0.102995857 | 0.000202759 | -0.048536206 | 0.085177082 | -0.103687287 | 1.405789682 | 0.555786081 |
| | | | 1631 | 2.928430576 | -0.54443464 | 0.961893977 | -0.50745464 | 0.360336337 | -0.346601577 | 2.611883872 | -0.550473108 | 0.552798143 | 0.26153036 |
| | | | 1632 | 0.93959755 | -0.313932226 | 0.14022317 | -0.287124558 | 0.024479244 | -0.215705684 | 0.698364915 | -0.349834831 | 1.899547326 | 0.64884381 |
| | | | 1633 | 0.451940314 | 0.106048684 | 0.001084382 | 0.006298436 | 0.002964065 | 0.100907605 | 0.793350553 | 0.229719202 | 0.672085898 | -0.229573102 |
| | | | 1634 | 0.310856317 | -0.07274496 | 0.057429347 | -0.108826612 | 0.090128477 | -0.197598129 | 0.298701901 | -0.111126279 | 1.798616014 | -0.3101626 |
| | | | 1635 | 0.104956562 | -0.028945015 | 0.340398208 | -0.26587496 | 0.000202759 | -0.070980788 | 0.11339643 | 0.051228843 | 0.448384338 | -0.179231681 |
| | | | 1636 | 0.344964121 | -0.148738998 | 0.342738351 | 0.344341393 | 0.228672682 | -0.395879583 | 0.115132936 | -0.098368678 | 0.760595735 | 0.360166502 |
| 21 | Unknown | Unknown | 1682 | 0.405934008 | 0.330273185 | 1.518039136 | -1.627773827 | 0.024985102 | -0.517583747 | 0.467064485 | -0.659423859 | 0.463211421 | -0.841518085 |
| | | | 1683 | 0.564404339 | 0.483739855 | 0.329595114 | -0.886785053 | 0.000202759 | 0.163854798 | 0.506318872 | -0.767145671 | 0.198369392 | -0.501852417 |
| 22 | Fructose | HTCS | 1754 | 0.25813692 | -0.06790073 | 0.022170845 | -0.052925234 | 0.000202759 | -0.067251944 | 0.218677522 | -0.089231095 | 0.446443449 | -0.144711706 |
| | | | 1755 | 1.167593842 | -0.642310625 | 0.029046553 | 0.153968366 | 0.004537328 | -0.253584969 | 0.989607465 | 0.822411579 | 0.527189548 | 0.473000844 |
| | | | 1756 | 0.122950821 | 0.14274374 | 0.03140915 | -0.234032558 | 0.000202759 | -0.135844521 | 0.098394466 | 0.244937269 | 0.405192703 | 0.54863253 |
| | | | 1757 | 0.137908668 | -0.059657874 | 0.036987217 | -0.102184079 | 0.001687904 | -0.124789941 | 0.102816913 | -0.084762498 | 0.160925295 | -0.120440862 |
| | | | 1758 | 0.234459633 | -0.141990581 | 0.070919568 | -0.229948973 | 0.004851778 | -0.207839603 | 0.0253957 | -0.039840353 | 0.02822287 | 0.047927364 |
| | | | 1759 | 0.058835449 | -0.033692601 | 0.439998695 | -0.521034905 | 0.006518177 | -0.202429035 | 0.173567334 | -0.192695617 | 0.381139923 | -0.320622994 |
| | | | 1760 | 0.441300982 | -0.2496391 | 0.095347032 | -0.263209272 | 0.000202759 | -0.132197244 | 0.056277448 | -0.090049899 | 0.031350432 | 0.050611884 |
| | | | 1761 | 0.338712221 | -0.296875704 | 0.068720957 | -0.299440001 | 0.000202759 | -0.040252383 | 0.244712345 | -0.370364742 | 0.076014776 | -0.152916471 |
| | | | 1762 | 1.36947716 | -0.678472821 | 0.343595974 | -0.588860269 | 0.004189015 | -0.24295348 | 1.024536321 | -0.741941791 | 0.510448775 | -0.505962976 |
| | | | 1763 | 0.526813726 | -0.320345169 | 0.533751903 | -0.74299203 | 0.197069787 | -0.652971818 | 1.128389684 | -0.760553731 | 1.005908827 | -0.80621365 |
| | | | 1764 | 0.683042083 | -0.202990366 | 0.096829484 | -0.183906864 | 0.000202759 | -0.081571639 | 0.564317165 | -0.256295615 | 0.362300023 | -0.206632994 |
| | | | 1765 | 0.951031481 | -0.573449742 | 0.141843698 | -0.442460134 | 0.173484833 | -0.681389789 | 0.106982117 | 0.209991843 | 0.478587226 | -0.537658313 |
| 23 | Unknown | Unknown | 1768 | 0.654164373 | 0.141198363 | 0.240279901 | 0.216448469 | 0.000202759 | 0.056206339 | 0.297471338 | 0.116692297 | 0.178984529 | 0.086673177 |
| | | | 1769 | 1.158575328 | -0.218676329 | 0.019048619 | -0.042514495 | 0.024985102 | -0.148163782 | 1.8116598 | 0.333097167 | 0.695506687 | -0.263662761 |
| | | | 1770 | 0.017334774 | -0.005601845 | 0.019656129 | -0.040604289 | 0.007237187 | -0.098317075 | 0.01626963 | -0.008528545 | 0.588984702 | -0.144904396 |
| | | | 1771 | 0.10447303 | -0.066050713 | 0.001776386 | -0.013521094 | 0.000202759 | 0.11026506 | 1.028069181 | -0.590509221 | 0.36480768 | -0.314518026 |
| | | | 1772 | 0.093510826 | 0.081254233 | 0.022170845 | -0.109601039 | 0.000202759 | 0.106475119 | 0.639936755 | -0.56349515 | 0.208876433 | -0.305357783 |
| | | | 1773 | 0.118841486 | 0.095892422 | 0.029046553 | -0.121693199 | 0.000202759 | 0.044299223 | 0.56139397 | -0.46020062 | 0.702885223 | -0.59303965 |
| | | | 1774 | 1.203561124 | -0.398587184 | 0.739620379 | -0.550910091 | 0.268813884 | -0.441911728 | 5.244125144 | -0.98317841 | 3.351531673 | -0.861335126 |
| | | | 1775 | 0.564387948 | 0.396662805 | 0.056699829 | 0.261136635 | 0.047316364 | 0.43148442 | 0.300757166 | -0.43798679 | 0.071526193 | 0.164278252 |
| | | | 1776 | 0.754962466 | -0.232173575 | 0.128189685 | -0.211693896 | 0.004537328 | -0.139257627 | 0.424515129 | -0.256096119 | 0.576075636 | -0.282331647 |
| | | | 1777 | 0.825046515 | -0.196323805 | 0.39685192 | -0.306061157 | 0.033695882 | -0.179459758 | 3.29700628 | -0.486505674 | 3.099283391 | -0.494515758 |
| | | | 1778 | 0.386618481 | -0.361123992 | 0.047801115 | -0.250330738 | 0.006934664 | -0.335699984 | 0.398924245 | -0.512401606 | 0.115442824 | -0.22464918 |
| | | | 1779 | 0.035855397 | 0.052119758 | 0.02416949 | -0.186128371 | 0.022858763 | -0.505983751 | 0.065753763 | -0.16468292 | 0.072654149 | -0.196058434 |
| | | | 1780 | 0.02014491 | -0.023139727 | 0.015574976 | 0.113930233 | 0.000202759 | -0.02714605 | 0.145947007 | -0.279204214 | 0.53417938 | -0.71426015 |
| | | | 1781 | 0.175325383 | 0.196504409 | 0.029046553 | 0.200348505 | 0.000202759 | -0.032191671 | 0.036556433 | -0.097042028 | 0.058690566 | -0.167230024 |
| 24 | Unknown | ECF | 1871 | 0.175404497 | -0.632552773 | 0.108227129 | -0.206696776 | 0.000202759 | 0.025434116 | 0 | -1.698881431 | 1.50494397 | -0.234319664 |
| | | | 1872 | 0.130401536 | -0.53508123 | 0.002741709 | -0.022150329 | 0.000202759 | 0.001789264 | 0 | -1.518155004 | 0.462959377 | -0.126831019 |
| | | | 1873 | 0.127277097 | -0.087748467 | 0.344922117 | -0.526777362 | 0.000202759 | 0.01593462 | 0.230124902 | -0.245838592 | 0.149295702 | -0.202581939 |
| | | | 1874 | 0.291952955 | -0.205283095 | 0.024194763 | 0.115782439 | 0.000202759 | 0.122401843 | 0.024114989 | -0.041835836 | 0.340903953 | 0.374822478 |
| | | | 1875 | 0.306451345 | -0.186599344 | 0.091477514 | -0.232455531 | 0.000202759 | 0.074360727 | 0.143005629 | -0.184705268 | 0.151952698 | -0.180234964 |
| | | | 1876 | 0.076033933 | 0.056446059 | 0.00230855 | -0.021370817 | 0.000202759 | 0.003504891 | 0.119594328 | 0.154108206 | 0.036008543 | 0.065175481 |
| | | | 1877 | 0.124947629 | -0.079324466 | 0.022460809 | -0.100041202 | 0.024985102 | -0.256159092 | 0.424515129 | -0.306490759 | 0.272890113 | -0.24335218 |
| | | | 1878 | 0.041380501 | 0.012575837 | 0.437865006 | 0.34394877 | 0.344680472 | -0.288211558 | 1.581852081 | -0.275166499 | 0.536659002 | -0.170503842 |
| 25 | Unknown | Unknown | 2032 | 0.471305385 | -0.18660159 | 0.029046553 | -0.063504771 | 0.25395283 | -0.356410006 | 11.04720756 | -1.065468459 | 28.32148162 | -1.339470578 |
| | | | 2033 | 1.106256769 | -0.366784924 | 0.074888922 | -0.134827555 | 0.491403646 | -0.389262607 | 13.59176003 | -1.155161933 | 27.54821356 | -1.450838646 |
| 26 | Unknown | Other | 2103 | 0.500845579 | 0.142355716 | 0.056878987 | 0.109306923 | 0.131882496 | 0.265713251 | 1.02108785 | 0.278677858 | 0.427069436 | 0.2115121 |
| | | | 2104 | 0.243346936 | -0.227272941 | 0.00230855 | 0.029370278 | 0.000202759 | 0.123206445 | 0.543897851 | -0.629646486 | 0.087391607 | -0.185390341 |
| | | | 2105 | 1.049814943 | -0.707569174 | 0.441074219 | -0.899782736 | 0.000202759 | 0.123377702 | 0.013966822 | -0.038170043 | 0.512981095 | -0.632504595 |
| | | | 2106 | 0.050384919 | 0.088954619 | 0 | -0.483782002 | 0.000202759 | -0.209075748 | 0.075000064 | -0.251390671 | 0 | 0.155159378 |
| | | | 2107 | 0.486648238 | -0.316496792 | 0.743180606 | -0.877119477 | 0.000202759 | 0.024120937 | 0.904569097 | -0.644188248 | 0.854576415 | -0.683427163 |
| | | | 2108 | 0.221309761 | -0.224261207 | 0.174347966 | -0.550678319 | 0.000202759 | 0.126967863 | 0.131102287 | -0.234487237 | 0.361900214 | -0.514666198 |
| | | | 2109 | 0.092498124 | -0.167104559 | 0 | -0.418830276 | 0.004537328 | 0.508013452 | 0.425042206 | 0.811078048 | 0 | 0.703158967 |
| | | | 2110 | 0.315330206 | -0.32727519 | 0.057501259 | -0.391847473 | 0.000202759 | -0.052698928 | 0.079247556 | 0.170177315 | 0.005770189 | 0.014599875 |
| | | | 2111 | 0.020227698 | 0.02234404 | 0.036192153 | 0.18705609 | 0.000202759 | -0.070153542 | 0.157098882 | 0.229950022 | 0.04762567 | 0.099060608 |
| | | | 2112 | 0.306465897 | -0.535888386 | 0 | 0.101516438 | 0.033990745 | 0.621278289 | 0.154440678 | 0.37296492 | 0.009091692 | 0.03418709 |
| | | | 2113 | 0.192127681 | -0.164032261 | 0.00230855 | -0.021516858 | 0.004851778 | 0.222115684 | 0.398924245 | 0.389468051 | 0.616396085 | 0.494946979 |
| 27 | Unknown | ECF | 2169 | 0.001586769 | 0.000851978 | 2.5969E-05 | -0.002569723 | 0.000202759 | -0.08118373 | 0.09184558 | 0.083113843 | 0.057526879 | -0.062893976 |
| | | | 2170 | 0.149991835 | -0.069631582 | 0.039756938 | 0.136477231 | 0.004537328 | 0.144963565 | 2.335281325 | 0.576631174 | 2.638221562 | 0.76047232 |
| | | | 2171 | 0.282391544 | 0.109058965 | 0.022503699 | 0.071026857 | 0.000202759 | -0.0786355 | 1.498201683 | 0.490830315 | 4.568636236 | 0.779686436 |
| | | | 2172 | 0.178683448 | 0.075926359 | 0.133625486 | 0.207212299 | 0.001687904 | 0.120843477 | 3.292687923 | 0.599801578 | 5.910094889 | 0.849237729 |
| | | | 2173 | 0.898658473 | 0.431673894 | 1.048657312 | 0.800689294 | 0.133650453 | 0.488055796 | 2.010399985 | 0.76885467 | 3.202412571 | 1.018583418 |
| 28 | Unknown | ECF | 2190 | 0.369061794 | 0.333205789 | 0.097677999 | 0.432865613 | 0.000202759 | 0.228349821 | 0.279510526 | 0.399641891 | 0.164168464 | 0.330943691 |
| | | | 2191 | 0.027647587 | 0.017546593 | 0.06947662 | -0.188342526 | 0.000202759 | -0.070831463 | 0.173567334 | -0.167573178 | 0.002300483 | -0.002611986 |
| | | | 2192 | 0.211643332 | -0.172841791 | 0.001082813 | 0.008088761 | 0.000202759 | 0.070518866 | 0.00614555 | 0.012404755 | 0.081390339 | 0.163098413 |
| | | | 2193 | 0.095611387 | -0.084035055 | 0.041750309 | -0.193696906 | 0.000202759 | -0.081727553 | 0.265623203 | -0.347927719 | 0.031350432 | -0.06681144 |
| | | | 2194 | 0.508403045 | 0.70289113 | 0 | 0.424563475 | 0.000202759 | 0.254564173 | 0.409549224 | 0.731883499 | 0 | 0.712505665 |
| | | | 2195 | 0.113121368 | -0.101664596 | 0.025030963 | 0.131467566 | 0.000202759 | -0.0419316 | 0.182307088 | -0.284603647 | 0.038254705 | -0.086276248 |
| | | | 2196 | 1.122517036 | -0.41905255 | 0.116104328 | -0.254867611 | 0.236070184 | -0.477824461 | 1.399402044 | -0.600790754 | 0.867964555 | -0.494668708 |
| | | | 2197 | 0.026895109 | -0.031583463 | 2.5969E-05 | -0.004691451 | 0.000202759 | -0.058228525 | 0.088644925 | -0.189479532 | 0.002522079 | 0.00510025 |
| | | | 2198 | 0.36708148 | 0.137924909 | 0.081515547 | 0.186201555 | 0.001985343 | 0.126342174 | 0.040307982 | -0.044323903 | 0.168668256 | 0.139107369 |
| 29 | Unknown | Other | 2199 | 0.248138193 | -0.090272901 | 0.00230855 | 0.015803479 | 0.000202759 | 0.059539985 | 0.042158373 | -0.036821074 | 0.170968951 | -0.115254346 |
| | | | 2200 | 0.138242868 | -0.085454039 | 0.004660803 | 0.037787818 | 0.024951395 | 0.275842187 | 0.513343105 | 0.351655435 | 0.0168207 | 0.029958732 |
| | | | 2201 | 1.780319903 | -0.434219217 | 0.513153888 | -0.441043297 | 0.000202759 | -0.08301462 | 0.408405689 | -0.237821975 | 0.622314668 | -0.321377037 |
| | | | 2202 | 4.30034021 | -0.668406057 | 0.909820752 | -0.52232122 | 0.006518177 | -0.173968186 | 2.412890281 | -0.523318716 | 1.316143217 | -0.421299902 |
| | | | 2203 | 1.032593202 | -0.38004397 | 0.554797283 | -0.579441738 | 0.022858763 | -0.244146351 | 1.377498598 | -0.547241035 | 1.433196947 | -0.603010589 |
| | | | 2204 | 0.237089635 | -0.06444215 | 0.141392593 | -0.180828948 | 0.001687904 | -0.105080377 | 3.418141327 | -0.472766814 | 0.962101911 | -0.23494626 |
| 30 | Unknown | Unknown | 2268 | 0.882138154 | 0.762088759 | 0.001084382 | 0.024509831 | 0.000202759 | -0.298374792 | 0.056370013 | 0.249098074 | 0.03546796 | 0.115579199 |
| | | | 2269 | 0.691837265 | 0.812072338 | 0.001776386 | 0.054667341 | 0.000202759 | -0.300928558 | 0.056181935 | 0.303168484 | 0.035561327 | 0.155941096 |
| 31 | Unknown | Unknown | 2362 | 0.302566833 | 0.288006362 | 1.626563296 | -1.533059127 | 1.78282275 | -1.485565643 | 1.399402044 | -0.986478217 | 0.762930714 | -0.685956251 |
| | | | 2363 | 0.171722385 | 0.167761376 | 0.403577471 | -0.847666979 | 0.64039358 | -1.163493756 | 0.477600697 | -0.588239957 | 0.258644131 | -0.446306539 |
| | | | 2364 | 1.852145234 | -1.712889507 | 0.02416949 | -0.256421865 | 0.004537328 | -0.536547461 | 0.024114989 | 0.09887112 | 0.39826892 | -0.907266859 |
| | | | 2365 | 2.951296067 | -2.173436503 | 0.029046553 | -0.327021077 | 0.007046896 | -0.639679438 | 0.064361882 | 0.259681304 | 0.27004935 | -0.783226798 |
| 32 | Unknown | HTCS | 2391 | 0.325298318 | -0.446654764 | 0.060901795 | 0.238976522 | 0.000202759 | -0.035122708 | 0.265025474 | -0.778545461 | 0.060902928 | -0.183028841 |
| | | | 2392 | 0.016419492 | -0.029448158 | 0 | -0.144732034 | 0.084278937 | -1.026663254 | 0.095991126 | -0.288628961 | 0 | -0.983510296 |
| | | | 2393 | 0.334248231 | -0.290631938 | 0.136903401 | -0.436679047 | 0.002964065 | -0.216463352 | 0.923909635 | -0.801907734 | 0.275570889 | -0.376243366 |
| | | | 2394 | 0.265630328 | -0.283191453 | 0.056878987 | -0.323728873 | 0.001687904 | -0.31849444 | 0.186101623 | -0.340385676 | 0.014185709 | -0.04627305 |
| | | | 2395 | 0.012155292 | 0.016733028 | 0.055096551 | -0.327960628 | 0.030771831 | 0.465092094 | 0.145183382 | -0.299386851 | 0.023651477 | -0.079793371 |
| 33 | Unknown | ECF | 2457 | 1.45261604 | 1.028673837 | 0.295120544 | 0.810703321 | 0.380270052 | 1.077636883 | 1.334665454 | 1.256494106 | 1.332295267 | 1.263275097 |
| | | | 2458 | 0.671326652 | 0.54327391 | 0.124108142 | 0.460401419 | 0.203356203 | 0.778149736 | 1.654125463 | 1.057286096 | 1.463586161 | 1.080201408 |
| | | | 2459 | 1.470189855 | 0.714092954 | 0.121551038 | 0.399666248 | 0.64039358 | 0.905106114 | 4.199970641 | 1.360035047 | 2.110935964 | 1.071940865 |
| | | | 2460 | 0.037885263 | -0.038803684 | 0.029046553 | -0.161223927 | 0.017081821 | 0.354552855 | 0.599737064 | 0.606446465 | 0.292857238 | 0.429609585 |
| | | | 2461 | 0.210370708 | 0.147326132 | 0.005185202 | 0.051282613 | 0.006518177 | 0.262800745 | 0.483062902 | 0.433338838 | 0.382601203 | 0.386042249 |

| | | | | | | | | | | | | | |
|---|---|---|---|---|---|---|---|---|---|---|---|---|---|
| | | | 2462 | 0.019322191 | -0.019777527 | 0.002741709 | 0.04170784 | 0.004537328 | 0.283847936 | 0.182359462 | 0.26574537 | 0.085844925 | 0.159031696 |
| | | | 2463 | 1.480574538 | 0.658503681 | 0.155592224 | 0.418979209 | 0.038757573 | 0.423265729 | 0.986932622 | 0.684481288 | 0.413109667 | 0.46003924 |
| 34 | Unknown | Unknown | 2529 | 0.130477541 | -0.222172067 | 0 | -0.26821365 | 0.000202759 | 0.160427232 | 0.068817223 | 0.231942381 | 0.22564787 | 0.54853305 |
| | | | 2530 | 0.700596317 | -0.623741229 | 0.022460809 | -0.144424547 | 0.000202759 | -0.077473225 | 0.231800683 | -0.371885882 | 0.090430788 | -0.198576681 |
| | | | 2531 | 0.141912946 | -0.132747682 | 0.026534872 | -0.14935702 | 0.006518177 | -0.387785706 | 0.083685199 | -0.163033575 | 0.187395103 | -0.320772637 |
| | | | 2532 | 0.108698188 | 0.153216244 | 0.131721647 | 0.557339271 | 0.022858763 | 0.51705486 | 0.100629809 | 0.248496536 | 0.01465377 | 0.057575833 |
| | | | 2533 | 0.630370867 | 1.102651243 | 0 | 1.604279775 | 0.019318782 | 0.902783859 | 0.284243364 | 0.954622847 | 0 | 0.524423834 |
| 35 | Unknown | ECF | 2559 | 0.14392303 | -0.080008879 | 0.097353676 | -0.252565811 | 0.024985102 | -0.248618085 | 0.088644925 | -0.095491204 | 0.011820122 | -0.017098497 |
| | | | 2560 | 0.24004915 | -0.098442478 | 0.700434042 | -0.545253814 | 0.130683092 | -0.311522907 | 0.982230775 | -0.40589366 | 1.86172674 | -0.496932877 |
| | | | 2561 | 0.135917783 | 0.061783249 | 0.019048619 | -0.06577019 | 0.000202759 | 0.047195703 | 0.65985513 | 0.29188594 | 0.110861317 | 0.098305398 |
| | | | 2562 | 0.093581172 | -0.042444886 | 0.029883833 | -0.094005809 | 0.00129528 | -0.147563673 | 0.155411899 | -0.114051378 | 0.374641456 | -0.219508541 |
| 36 | α-mannan | HTCS | 2615 | 0.026001695 | 0.025387708 | 0.061719218 | -0.29829319 | 0.000202759 | -0.136357577 | 0.333860038 | -0.37740046 | 0.113699157 | -0.189738068 |
| | | | 2616 | 0.063762107 | -0.127431659 | 0 | 0.502626975 | 0.000202759 | 0.229934464 | 0.267386307 | 0.614423723 | 0 | 0.002797574 |
| | | | 2617 | 0.085452411 | 0.0597649 | 0.06947662 | -0.224903345 | 0.024985102 | -0.265261056 | 0.648256565 | -0.420889937 | 0.867358587 | -0.594687914 |
| | | | 2618 | 0.795601716 | 0.499138102 | 0.272852563 | 0.549502499 | 0.000202759 | 0.135621794 | 0.207378924 | 0.290420769 | 0.172156349 | 0.293305709 |
| | | | 2619 | 1.087469003 | 0.365031646 | 0.154217493 | 0.278736673 | 0.000202759 | 0.124968768 | 0.403080353 | 0.264568942 | 0.043580897 | -0.061932736 |
| | | | 2620 | 0.143187138 | 0.09755401 | 0.057501259 | 0.229887836 | 0.000202759 | 0.087158113 | 0.016581476 | -0.029425541 | 0.138536034 | -0.182688936 |
| | | | 2621 | 0.552944992 | 0.266466136 | 0.098306198 | 0.25210821 | 0.000202759 | 0.051772169 | 0.08239121 | -0.107104783 | 0.352925226 | -0.320998669 |
| | | | 2622 | 0.326425175 | 0.18531244 | 0.015664147 | 0.080031218 | 0.003466816 | 0.208684405 | 0.207617972 | 0.206840362 | 0.085844925 | -0.133627941 |
| | | | 2623 | 0.060934899 | -0.036923789 | 0.042376758 | -0.177289978 | 0.000202759 | -0.042353654 | 0.030510593 | -0.043058687 | 1.024157966 | -0.531548851 |
| | | | 2624 | 0.215788003 | -0.140585804 | 0.029883833 | -0.14701335 | 0.022858763 | -0.307178286 | 0.286880075 | -0.282844394 | 0.742033686 | -0.580381108 |
| | | | 2625 | 0.327428551 | -0.205281537 | 0.081692542 | -0.283728201 | 0.000202759 | -0.132874093 | 0.152663351 | -0.177915958 | 0.696835924 | -0.482229729 |
| | | | 2626 | 1.109948026 | -0.421655055 | 0.612111244 | -0.763778234 | 0.006793048 | -0.225427348 | 0.157334119 | -0.159374793 | 0.439833481 | -0.369305004 |
| | | | 2627 | 0.146705113 | -0.090377795 | 0.736144676 | -0.702343319 | 0.006518177 | -0.234960617 | 0.797544322 | -0.496179848 | 0.824408589 | -0.541607328 |
| | | | 2628 | 0.914501792 | -0.372458278 | 0.748519366 | -0.662825308 | 0.024479244 | -0.283356412 | 0.764774647 | -0.426407741 | 0.857610396 | -0.509241621 |
| | | | 2629 | 0.147945829 | -0.072178529 | 0.4373438 | -0.437192621 | 0.030771831 | -0.249088218 | 2.377552827 | -0.626126187 | 3.459595438 | -0.865371352 |
| | | | 2630 | 1.176349076 | -0.278804945 | 0.745461223 | -0.444958126 | 0.25395283 | -0.363989342 | 0.935018223 | -0.332983221 | 2.983777543 | -0.584227153 |
| | | | 2631 | 1.270291694 | -0.311106425 | 0.854464855 | -0.546782777 | 0.006518177 | -0.174415525 | 0.217385408 | -0.159597871 | 1.041868209 | -0.376065384 |
| | | | 2632 | 1.178634851 | -0.332575465 | 0.763282073 | -0.595183558 | 0.314929789 | -0.377298723 | 0.401130206 | -0.20085653 | 1.581043468 | -0.378836831 |
| | | | 2633 | 0.868917173 | -0.472272838 | 0.258448928 | -0.623121141 | 0.051341828 | -0.378002206 | 0.056277448 | -0.091171095 | 0.26059884 | -0.301172421 |
| 37 | Ribose | Other | 2802 | 0.256714337 | -0.072752053 | 0.46350024 | -0.343244265 | 0.195903958 | -0.291977542 | 2.352039235 | -0.466278668 | 2.533952235 | -0.45631716 |
| | | | 2803 | 0.438988195 | -0.193213626 | 0.018104878 | -0.064496389 | 0.000202759 | 0.110249409 | 0.173567334 | 0.159371299 | 0.109140466 | -0.122618677 |
| | | | 2804 | 1.358057501 | -0.367588247 | 0.4922517 | -0.392434687 | 0.101367857 | -0.257518108 | 0.021537641 | -0.02604404 | 0.557874327 | -0.293350727 |
| | | | 2805 | 1.494531529 | -0.273211622 | 0.372751145 | -0.308425808 | 0.015529428 | -0.136165559 | 0.148694653 | 0.104944702 | 1.456840195 | -0.367623217 |
| | | | 2806 | 2.627078612 | -0.453845742 | 0.446763368 | -0.3740717 | 0.017081821 | -0.165231438 | 0.357692496 | 0.216075633 | 1.031813804 | -0.334698996 |
| | | | 2807 | 0.410069616 | -0.154558994 | 0.029046553 | -0.081235311 | 0.004537328 | -0.153149306 | 0.492048258 | 0.25133861 | 0.969114147 | -0.377007613 |
| | | | 2808 | 0.015313664 | -0.007275544 | 0.00230855 | 0.015434945 | 0.006518177 | -0.16390586 | 0.30519768 | 0.169661851 | 0.729324089 | -0.346394653 |
| | | | 2809 | 0.030013472 | -0.013620094 | 0.036987217 | -0.114426145 | 0.037591022 | -0.221479224 | 0.028528329 | 0.030447923 | 0.767729409 | -0.354218168 |
| 38 | Unknown mucin *O*-glycan | 38 | 2818 | 0.32687066 | -0.161913838 | 0.763085102 | -0.621561865 | 0.047316364 | -0.299699454 | 0.157998581 | -0.18747548 | 0.772197657 | -0.506951776 |
| | | | 2819 | 0.118327677 | -0.10404115 | 0.002406745 | 0.031026174 | 0.024985102 | -0.356601353 | 0.016120144 | -0.034638173 | 0.383100755 | -0.411756673 |
| | | | 2820 | 0.564544111 | 0.328793285 | 0.018130683 | -0.097326636 | 0.000202759 | 0.07628522 | 0.594083282 | 0.502934525 | 0.173390284 | 0.244785709 |
| | | | 2821 | 0.29947405 | 0.224036489 | 0.003720629 | 0.043788698 | 0.006518177 | -0.327329855 | 0.104792928 | -0.198626793 | 0.305314473 | -0.405750883 |
| | | | 2822 | 0.163400898 | -0.146147594 | 0.019916802 | -0.126773256 | 0.017081821 | 0.316067827 | 0.017800411 | 0.044251934 | 0.1097465 | -0.22040897 |
| | | | 2823 | 0.747527017 | 0.481550415 | 0.118210922 | 0.402672351 | 0.090042064 | 0.56108701 | 0.318943264 | 0.398823287 | 0.162007648 | 0.280470143 |
| | | | 2824 | 0.978697005 | 0.590795165 | 0.150627663 | 0.465416425 | 0.000202759 | 0.130571926 | 0.244039621 | 0.358269216 | 0.001055555 | 0.002589262 |
| | | | 2825 | 0.265838403 | 0.458006132 | 0 | 0.167791454 | 0.000202759 | 0.099232919 | 0.039363221 | 0.17918293 | 0 | 0.369899023 |
| | | | 2826 | 0.540127581 | 0.146849089 | 0.086687612 | 0.145819878 | 0.000202759 | 0.090507029 | 0.045748256 | -0.042549916 | 0.102809241 | -0.075462554 |
| 39 | Unknown | HTCS | 2851 | 0.937172861 | -0.19964357 | 0.088109783 | -0.132693998 | 0.097494468 | -0.211571199 | 0.657202105 | -0.238662007 | 3.182324253 | -0.450178133 |
| | | | 2852 | 1.097581131 | -0.247970482 | 0.32717508 | -0.295545419 | 0.145883736 | -0.271305553 | 0.506318872 | -0.207885972 | 1.885189326 | -0.432322043 |
| | | | 2853 | 0.194851675 | 0.183965143 | 0.136903401 | -0.508512537 | 0.000202759 | 0.133321967 | 0.209757604 | -0.350183494 | 0.055704297 | -0.142270421 |
| | | | 2854 | 0.543634067 | -0.157728415 | 0.108695747 | -0.179351657 | 0.019865857 | -0.168585918 | 0.99276002 | -0.324929332 | 1.283873643 | -0.398352559 |
| | | | 2855 | 0.150860905 | -0.079516613 | 0.05562578 | 0.161051956 | 0.000202759 | -0.08474942 | 0.177383853 | 0.158058135 | 0.264190685 | -0.218773002 |
| | | | 2856 | 0.133633233 | 0.13866327 | 0.047218354 | 0.246672179 | 4.81016E-05 | -0.000467292 | 0.408508132 | 0.473499973 | 0.014321959 | 0.043795813 |
| | | | 2857 | 0.208437013 | -0.197251132 | 0.074888922 | 0.329940573 | 0.000202759 | 0.054637369 | 0.718431223 | 0.597506204 | 0.020734582 | 0.053312514 |
| | | | 2858 | 1.215993402 | -0.573158757 | 0.051405557 | -0.187269787 | 0.000202759 | -0.119476743 | 0.422199904 | 0.323253913 | 0.161535022 | -0.203639886 |
| | | | 2859 | 2.757229559 | -0.828784792 | 0.792044503 | -0.740631027 | 0.159950897 | -0.380464724 | 0.007313381 | -0.010999032 | 1.172392002 | -0.459891307 |
| | | | 2860 | 1.451542537 | -0.25983333 | 0.036987217 | -0.084426526 | 0.000202759 | -0.096450122 | 0.102298813 | 0.060550295 | 0.111323255 | -0.069225565 |
| 40 | Unknown | HTCS | 2892 | 0.529549545 | 0.439759011 | 0.200367529 | 0.554284909 | 0.209550615 | 0.770124077 | 0.243391785 | 0.389459561 | 0.406143676 | 0.546680173 |
| | | | 2893 | 0.184673663 | -0.136052789 | 0.009717219 | -0.064462891 | 0.000525887 | -0.188189707 | 0.575652019 | -0.505834694 | 0.425847307 | -0.426197148 |
| | | | 2894 | 0.537062553 | -0.369980838 | 0.28949292 | -0.570316023 | 0.000202759 | 0.059149566 | 0.512360021 | -0.47750869 | 0.11946286 | -0.190349782 |
| | | | 2895 | 0.607724424 | 0.697575817 | 0 | 0.149407532 | 0.000202759 | 0.23557564 | 0.064361882 | -0.236214488 | 0 | 0.648700826 |
| | | | 2896 | 0.768333031 | -0.633135944 | 0.136757135 | -0.551155934 | 0.024062964 | -0.521855046 | 0.933261348 | -0.952261047 | 0.081390339 | -0.190602183 |
| | | | 2897 | 0.220725302 | -0.14866589 | 0.086396171 | -0.258805453 | 0.000202759 | -0.077721287 | 0.267386307 | -0.294637427 | 0.445884716 | 0.367528833 |
| 41 | Unknown | Other | 2898 | 0.970777493 | -0.348998564 | 0.136903401 | -0.2861216 | 0.000525887 | -0.12972191 | 0.162066127 | -0.147068166 | 0.245352221 | -0.189421624 |
| | | | 2899 | 0.903558332 | -0.204742598 | 0.023875995 | -0.058055083 | 0.000525887 | -0.096901518 | 1.181769196 | -0.312416269 | 3.009179603 | -0.513591456 |
| | | | 2900 | 0.053856452 | 0.078154938 | 0 | 0.183124165 | 0.004851778 | 0.383670958 | 0.03370229 | -0.109818645 | 0.021069188 | -0.080054347 |
| | | | 2901 | 0.217676553 | -0.145845604 | 0.001600848 | -0.012783156 | 0.000202759 | -0.060196157 | 0.828996495 | -0.555601949 | 0.939572518 | -0.639798802 |
| | | | 2902 | 0.64171782 | -0.487706588 | 0.039729294 | -0.227290799 | 0.097494468 | -0.621905439 | 0.348706042 | -0.46133028 | 0.135762502 | -0.261291205 |
| | | | 2903 | 0.153050998 | 0.157019467 | 0.023875995 | -0.160635863 | 0.02375206 | 0.402570745 | 0.074106006 | -0.177348358 | 0.112934157 | -0.249178693 |
| | | | 2904 | 1.323691785 | -0.445617089 | 0.724764987 | -0.635805345 | 0.195903958 | -0.440541576 | 2.184026188 | -0.754164464 | 3.267236443 | -0.955441407 |
| | | | 2905 | 0.322385161 | -0.145521581 | 0.069868405 | -0.181716208 | 0.006518177 | -0.198160546 | 1.356796917 | -0.523029445 | 1.84634465 | -0.712156246 |
| | | | 2906 | 1.355320951 | -0.508212305 | 0.002406745 | -0.023914582 | 0.003182526 | -0.186906686 | 0.903370385 | -0.508961121 | 1.797645043 | -0.763668537 |
| | | | 2907 | 0.42863126 | -0.158041535 | 0.094162752 | -0.19712061 | 0.004537328 | -0.165999531 | 0.362278361 | -0.264761963 | 1.194860586 | -0.503282384 |
| | | | 2908 | 0.222349651 | -0.160250746 | 0.422749717 | -0.62336385 | 0.040729453 | -0.363899047 | 0.434364772 | -0.381787599 | 0.536659002 | -0.450801147 |
| | | | 2909 | 1.100543719 | -0.382289079 | 0.623528953 | -0.587978579 | 0.15711585 | -0.406812477 | 1.354072602 | -0.55104987 | 0.510448775 | -0.344553742 |
| | | | 2910 | 0.97078738 | -0.220232063 | 0.540444706 | -0.406344404 | 0.228940066 | -0.367408075 | 2.753823567 | -0.533746321 | 2.467160935 | -0.498390016 |
| 42 | Unknown | HTCS | 2911 | 0.100795774 | -0.078199767 | 0.057501259 | 0.228756019 | 0.000202759 | 0.132327363 | 0.219030786 | -0.268333289 | 0.245352221 | 0.277415519 |
| | | | 2912 | 1.117728662 | 0.381969814 | 0.202251719 | 0.318052822 | 0.89421298 | 0.647426587 | 0.68965487 | 0.377558611 | 1.000530084 | 0.460592429 |
| | | | 2913 | 0.178669617 | 0.131218917 | 2.5969E-05 | 0.004313871 | 0.164880931 | 0.52355776 | 0.698364915 | 0.583831009 | 0.229566737 | 0.271273443 |
| | | | 2914 | 0.419803627 | -0.402161074 | 0.036987217 | -0.240315602 | 0.071879253 | 0.602450912 | 0.103939395 | 0.201083415 | 0.077591052 | 0.18436164 |
| | | | 2915 | 0.350609911 | -0.365587178 | 0.013460281 | -0.122784266 | 0.097494468 | 0.640744782 | 0.304550745 | 0.456551606 | 0.227491412 | 0.423508871 |
| | | | 2916 | 0.170526748 | -0.179032454 | 0.029046553 | 0.170988164 | 0.505675327 | 0.867081993 | 0.011759532 | -0.032374504 | 0.022448779 | -0.065863486 |
| | | | 2917 | 0.40460264 | -0.443203777 | 0.131721647 | -0.549132981 | 0.097494468 | 0.622104881 | 0.197690467 | -0.369240975 | 0.141315032 | -0.307252038 |
| | | | 2918 | 0.156430227 | 0.184529544 | 0.00536243 | -0.085378499 | 0.276049897 | 0.934067161 | 0.223910648 | -0.443249338 | 0.036008543 | -0.118609152 |
| | | | 2919 | 0.610708509 | -0.523929011 | 0.039914557 | -0.256880811 | 2.481918991 | 1.305118328 | 0.021871765 | -0.055153858 | 0.545073933 | -0.703508199 |
| | | | 2920 | 0.797609274 | -0.386120936 | 0.011269086 | -0.061304026 | 5.598599459 | 1.316706146 | 0.297209389 | -0.28115077 | 0.444293423 | -0.439712865 |
| | | | 2921 | 0.289540768 | -0.324251579 | 0 | -0.761994862 | 0.573176206 | 1.116468868 | 0.144919586 | -0.289847315 | 0.399601339 | -0.690561649 |
| | | | 2922 | 0.330450276 | 0.422817997 | 0 | 0.512537851 | 4.517126416 | 2.118717047 | 0.064175148 | 0.204118662 | 0 | 0.272515499 |
| | | | 2923 | 0.211514406 | -0.084547177 | 0.096829484 | -0.19800571 | 9.782516056 | 1.219596035 | 0.224145349 | 0.144012037 | 0.457408727 | 0.251492181 |
| 43 | Unknown | Other | 2948 | 0.631896478 | 0.235120225 | 0.035284985 | 0.104053075 | 0.047316364 | 0.301219619 | 2.096810191 | 0.514647771 | 0.119931401 | 0.09879495 |
| | | | 2949 | 0.163086648 | -0.147638827 | 0.009717219 | -0.070789338 | 0.000202759 | 0.119670128 | 0.179575827 | -0.272506397 | 0.489354185 | -0.553443736 |
| | | | 2950 | 0.399170634 | -0.406474802 | 0.009717219 | 0.095815238 | 0.000202759 | 0.070700582 | 0.132193619 | -0.274997672 | 0.055433963 | -0.162974774 |
| | | | 2951 | 0.264953775 | -0.251531942 | 0.273541724 | 0.608073652 | 0.000202759 | 0.212735724 | 0.165626811 | 0.285438666 | 0.030539411 | -0.077446557 |
| | | | 2952 | 0.109838314 | 0.072711417 | 0.011667305 | 0.056001123 | 0.054272408 | 0.323865435 | 0.010733949 | -0.016499248 | 0.138536034 | -0.179705646 |
| | | | 2953 | 0.059712225 | 0.100287517 | 0 | 0.560290828 | 0.000202759 | -0.196285791 | 0.211447536 | 0.481735603 | 0 | 0.35529098 |
| | | | 2954 | 0.125720959 | 0.624384604 | 0 | -0.393561011 | 0.000202759 | -0.919286514 | 0 | 0.372081801 | 0 | 0.582675432 |
| | | | 2955 | 0.701085917 | -0.934997401 | 0 | -0.565106812 | 0.000202759 | -0.234345624 | 0.04009803 | -0.144990009 | 0 | -0.422756478 |
| | | | 2956 | 0.190243054 | 0.843971631 | 0 | 0.860401964 | 0.006518177 | 1.213175932 | 0 | 0.64765965 | 0 | 0.799453001 |
| | | | 2957 | 0.005971218 | -0.005390726 | 0.08125053 | -0.24678853 | 0.006518177 | 0.225568294 | 0.174967974 | 0.200884221 | 0.53417938 | 0.398945319 |
| | | | 2961 | 0.177968563 | -0.079699118 | 0.050460707 | 0.129715863 | 0.017081821 | 0.190609949 | 0.056268098 | 0.054969977 | 0.072973959 | -0.079162979 |
| | | | 2962 | 0.179728365 | 1.195926743 | 0 | 0.476153479 | 0.000202759 | 1.306046974 | 0 | 0.912060909 | 0 | -0.238358975 |
| | | | 2963 | 0.262664128 | -0.118395899 | 0.032272961 | -0.101994604 | 0.063456331 | -0.300549836 | 0.104922088 | -0.088834657 | 0.095958072 | -0.09266951 |
| | | | 2964 | 0.415798896 | -0.239461245 | 0.05128535 | -0.188960274 | 0.055500432 | -0.364376745 | 0.215070163 | -0.231731164 | 0.078141122 | -0.113052526 |

| | | | | | | | | | | | | | |
|---|---|---|---|---|---|---|---|---|---|---|---|---|---|
| 44 | Unknown | HTCS | 2965 | 0.453318001 | -0.426804373 | 0.002741709 | -0.053326007 | 0.024479244 | 0.415140776 | 0.001689286 | -0.005117113 | 0.022729476 | 0.065553749 |
| | | | 2966 | 1.088138584 | -0.58478704 | 0.061697555 | -0.23628805 | 0.017081821 | -0.285907995 | 0.209959925 | -0.246226002 | 0.081390339 | -0.131620078 |
| | | | 2967 | 0.745571335 | -0.323463083 | 0.046998121 | 0.133822323 | 0.035394591 | -0.313854953 | 0.160498548 | -0.152576695 | 0.272420415 | -0.287610923 |
| | | | 2968 | 2.537858162 | -0.584082294 | 0.446763368 | -0.389607134 | 0.494215315 | -0.494147453 | 1.610911706 | -0.553634152 | 1.556601221 | -0.465627528 |
| | | | 2969 | 0.161452322 | -0.119202799 | 0.129852334 | -0.34175141 | 0.000202759 | 0.053189154 | 0.038029568 | -0.068802944 | 0.031350432 | 0.058108279 |
| | | | 2970 | 0.769595422 | -0.382528299 | 0.396273723 | -0.631055508 | 0.000202759 | -0.183209917 | 0.043191226 | -0.068150676 | 0.068054388 | -0.104541463 |
| | | | 2971 | 2.253951637 | -0.308739195 | 0.340692504 | -0.280367979 | 0.145994578 | -0.237205406 | 0.434364772 | -0.152189828 | 0.693069803 | -0.199278837 |
| 45 | Unknown mucin O-glycan | ECF | 3010 | 0.049507285 | 0.031912128 | 1.95089E-05 | 0.001266689 | 0.145469171 | 0.429391595 | 0.221651769 | 0.212165787 | 0.318333475 | 0.261926883 |
| | | | 3011 | 0.100661369 | 0.113992911 | 0.029046553 | -0.192139873 | 0.000202759 | 0.081660376 | 1.24352999 | -1.261393646 | 0.950467546 | 0.848958557 |
| | | | 3012 | 0.194957164 | -0.081724525 | 1.055844075 | -0.93420424 | 0.15711585 | -0.340301132 | 25.03385827 | -2.142795799 | 0.229566737 | -0.151774788 |
| | | | 3013 | 0.263565214 | 0.128642656 | 0.105628234 | -0.345269722 | 0.000202759 | -0.123553541 | 16 | -2.301051828 | 0.125919411 | -0.14286772 |
| | | | 3014 | 0.047026022 | -0.036118137 | 0.129213774 | -0.496879117 | 0.000202759 | -0.122272613 | 6.649751982 | -1.915990532 | 0.03546796 | -0.074096795 |
| | | | 3015 | 0.039123418 | 0.023417498 | 0.133625486 | -0.338847214 | 0.017805709 | -0.232689684 | 11.24565166 | -1.877103315 | 0.924137542 | 0.421561292 |
| | | | 3016 | 0.994332102 | 1.640893501 | 0 | 1.262731309 | 0.018081257 | 0.975898053 | 0.197690467 | 0.875984229 | 0 | 1.844866677 |
| | | | 3017 | 0.729823956 | 1.159146634 | 0 | 0.783367869 | 0.076819953 | 1.182182342 | 0.077466615 | 0.420676721 | 0 | 1.066028156 |
| 46 | Cellobiose | Unknown | 3024 | 1.207217908 | 0.596792414 | 0.097166039 | 0.328536638 | 0.047316364 | 0.462400553 | 0.112494207 | -0.399349088 | 0.130217829 | 0.217683025 |
| | | | 3025 | 0.446479811 | 0.337259179 | 0.767879512 | 0.881675344 | 0.065636162 | 0.502666691 | 0.108627099 | -0.27112866 | 0.068862828 | 0.152429607 |
| | | | 3026 | 1.171382227 | 0.538180239 | 0.295146693 | 0.505489115 | 0.06599851 | 0.438674522 | 0.015588425 | -0.045298067 | 0.317659847 | 0.372984206 |
| | | | 3027 | 0.609093868 | 0.457686383 | 0.459912311 | 0.795836877 | 0.15711585 | 0.699477576 | 0.04358834 | -0.13532431 | 0.215166247 | 0.370161379 |
| 47 | Unknown | HTCS | 3043 | 0.129757209 | 0.114466223 | 0.086396171 | 0.314831642 | 0.000202759 | 0.03476562 | 1.448157862 | 0.827252437 | 0.158063137 | 0.250726481 |
| | | | 3044 | 0.174831905 | -0.259851816 | 0.001600848 | -0.022507552 | 0.000202759 | -0.149674173 | 0.30475453 | 0.501121384 | 2.42245834 | 1.471963054 |
| | | | 3045 | 0.263605066 | -0.234629797 | 0.056699829 | -0.218048252 | 0.017796183 | -0.34903568 | 0.206056337 | 0.240132544 | 4.496209317 | 1.245004843 |
| | | | 3046 | 0.454720385 | -0.327720823 | 0.222995506 | -0.506986795 | 0.06599851 | -0.480725443 | 0.016581476 | 0.0309976 | 2.030718684 | 0.848336901 |
| | | | 3047 | 1.194251508 | -0.83135468 | 0.280317844 | -0.710876712 | 0.229231935 | -1.013884492 | 0.349545106 | -0.452697479 | 0.3405279 | 0.435284242 |
| | | | 3048 | 0.068646497 | 0.056524155 | 0.022170845 | 0.104808879 | 0.004720284 | -0.242588512 | 0.010631097 | -0.022601735 | 0.409098484 | 0.417333258 |
| | | | 3049 | 0.442161733 | -0.148875381 | 0.437865006 | -0.374538772 | 0.006518177 | -0.202628223 | 0.550367877 | -0.243447484 | 0.050922736 | -0.050821838 |
| 48 | Dextran | SusR | 3086 | 0.107734603 | -0.071653601 | 0.021295324 | 0.077905978 | 2.407055188 | 0.887494664 | 0.751125701 | 0.369951997 | 0.210337489 | 0.208539689 |
| | | | 3087 | 1.00568653 | -0.397414146 | 0.053599877 | 0.167417182 | 1.760592324 | 0.851938544 | 1.399402044 | 0.542208963 | 0.026020025 | 0.053563455 |
| | | | 3088 | 1.505318239 | -0.597488847 | 0.409751247 | -0.567868564 | 0.036400646 | 0.29796395 | 0.04358834 | -0.067213647 | 0.278201654 | -0.300732036 |
| | | | 3089 | 0.580944117 | -0.28857428 | 0.123120774 | -0.295659704 | 0.61982549 | 0.62807314 | 0.040060676 | 0.057179556 | 0.110293557 | -0.134022433 |
| | | | 3090 | 2.500200235 | -0.59080812 | 0.446763368 | -0.437522697 | 1.41844935 | 0.537833292 | 0.158050257 | -0.146570725 | 2.612533645 | -0.625334479 |
| | | | 3091 | 0.311064685 | -0.102651238 | 0.046998121 | -0.110115059 | 39.13076828 | 2.117838362 | 0.31878892 | -0.181873598 | 1.232506436 | -0.38484122 |
| 49 | Unknown | HTCS | 3092 | 0.800870029 | -0.431560975 | 0.295120544 | -0.514961488 | 295.4259687 | 6.305264833 | 0.173030098 | 0.178247825 | 0.088309791 | -0.121435292 |
| | | | 3093 | 1.669265211 | -0.619000768 | 0.175126877 | -0.34283944 | 300 | 6.464576323 | 0.221651769 | 0.204669416 | 0.764487148 | -0.47273378 |
| | | | 3094 | 0.720166525 | -0.4434387 | 0.256084944 | -0.63971735 | 208.2848326 | 6.273219008 | 0.118541348 | 0.195031788 | 0.273859344 | -0.343075126 |
| | | | 3095 | 3.563926448 | -0.871624458 | 0.916621967 | -0.780865572 | 148.0385789 | 4.367135734 | 0.116754347 | -0.128588962 | 0.761339988 | -0.451172691 |
| | | | 3096 | 2.052519714 | -0.58346102 | 0.191448537 | -0.312862058 | 172.7619539 | 4.594486559 | 0.07314107 | 0.075235001 | 2.612533645 | -0.784453716 |
| | | | 3097 | 0.218834412 | -0.090644802 | 0.027178702 | -0.074526387 | 258.3595186 | 4.561134578 | 1.658111514 | 0.449182771 | 1.223297167 | -0.447919853 |
| | | | 3098 | 1.408136081 | -1.385356308 | 0 | -0.539258344 | 21.9788107 | 3.464364517 | 0.229884473 | 0.441462826 | 0.003328382 | -0.009719401 |
| | | | 3099 | 0.549095815 | -0.471284034 | 0.027163795 | -0.169679779 | 44.01637371 | 3.881221351 | 0.511824295 | 0.608799222 | 0.014185709 | -0.044204356 |
| | | | 3100 | 0.158207367 | 0.218613024 | 0 | -0.472029532 | 163.7212464 | 6.952908577 | 0.014601605 | 0.047761191 | 0.095926871 | 0.257396378 |
| | | | 3101 | 0.447743895 | -0.345106932 | 0.07028867 | -0.316171606 | 200.0395292 | 7.494916411 | 0.064062447 | -0.132569833 | 0.142769778 | -0.245942874 |
| | | | 3102 | 0.061888695 | -0.053859313 | 0.004534747 | -0.058841812 | 196.6055483 | 7.994725833 | 0.184849338 | 0.246379162 | 0.146431127 | -0.231703353 |
| | | | 3103 | 0.495200354 | -0.280313764 | 0.070919568 | -0.28979992 | 182.3575355 | 7.918533504 | 0.027418213 | -0.046306272 | 0.826356322 | -0.606079143 |
| | | | 3104 | 0.24406576 | -0.275099706 | 0.118210922 | -0.527432074 | 143.0629839 | 7.890043675 | 0.149854007 | -0.296198482 | 0.444293423 | -0.651654288 |
| | | | 3105 | 0.50688538 | -0.500255705 | 0.247805688 | -0.904667697 | 183.2182446 | 7.960925293 | 0.25130561 | -0.431728468 | 0.142530882 | -0.381267386 |
| | | | 3106 | 1.905257304 | -1.001340766 | 0.538002238 | -1.120052213 | 201.2240257 | 7.626934868 | 0.28471488 | -0.384884045 | 0.378875277 | -0.567133901 |
| | | | 3107 | 0.573142718 | -0.515852122 | 0.029046553 | -0.213936811 | 229.0757207 | 7.956609151 | 0.064175148 | -0.156112688 | 0.453432094 | -0.694540586 |
| | | | 3108 | 0.273973565 | -0.392539998 | 0 | -0.577607167 | 217.7670039 | 8.244252988 | 0.017575703 | 0.063075118 | 0 | -0.456728691 |
| | | | 3109 | 0.068956171 | 0.095565048 | 0 | -0.207906123 | 222.0204516 | 8.064044123 | 0.234697157 | -0.608317366 | 0.087606517 | -0.303011336 |
| 50 | Unknown | Unknown | 3155 | 0.799259249 | 0.4462311 | 0.208875627 | 0.476028612 | 0.024985102 | 0.374952848 | 0.51698233 | 0.5661924 | 0.459123886 | 0.479709196 |
| | | | 3156 | 0 | -0.013153069 | 0.02416949 | -0.072968679 | 0.064022577 | -0.253397576 | 0.810706233 | -0.425244601 | 2.849736757 | -0.667150873 |
| | | | 3157 | 0.382040303 | -0.172474269 | 0.003600372 | -0.027675161 | 0.024985102 | -0.241538024 | 1.021822004 | -0.498481066 | 0.924413768 | -0.497212628 |
| | | | 3158 | 1.172432787 | -0.488091154 | 0.039224445 | -0.139759315 | 0.040729453 | -0.308718233 | 0.115132936 | -0.131114945 | 0.890533628 | -0.556468063 |
| | | | 3159 | 0.419411197 | -0.29851485 | 0.17999386 | -0.463118334 | 0.097494468 | -0.516116625 | 0.797544322 | -0.628968616 | 0.450028322 | -0.497263457 |
| 51 | Unknown | HTCS | 3171 | 0.257860445 | -0.134079545 | 0.069670121 | -0.200016836 | 0.000202759 | -0.015354755 | 0.040776683 | -0.053101144 | 0.055433963 | -0.078267912 |
| | | | 3172 | 0.921533673 | -0.199090978 | 0.343595974 | -0.301664165 | 107.0004345 | 2.716932872 | 3.292687923 | -0.506906379 | 0 | 3.52841716 |
| | | | 3173 | 0.800098903 | -0.242804255 | 0.343595974 | -0.335006772 | 0.529484679 | -0.412653206 | 1.806455536 | -0.486413583 | 4.185752404 | -0.780184056 |
| | | | 3174 | 0.040377087 | 0.043117149 | 0.081674782 | -0.411550311 | 0.000202759 | 0.02925514 | 0.008103241 | -0.022671681 | 0.154956655 | 0.266602911 |
| | | | 3175 | 0.467898585 | -0.592782524 | 0 | -0.736039046 | 0.1208687 | -0.997528884 | 0.723667269 | -1.087370571 | 0 | -0.689370367 |
| | | | 3176 | 0.147668394 | -0.214446183 | 0 | -0.003598161 | 0.000202759 | -0.172542916 | 0.100936037 | 0.268548168 | 0 | -0.693338642 |
| | | | 3177 | 0.055076464 | -0.131039953 | 0 | -0.59253628 | 0.000202759 | 0.2252282 | 0.201789649 | -0.651284853 | 0 | -0.310331351 |
| | | | 3178 | 1.229065694 | 0.973980556 | 0 | 0.67852715 | 0.097494468 | 0.851275955 | 0.168342665 | 0.412114453 | 0.964818934 | 1.097894303 |
| | | | 3179 | 0.344426133 | 0.115128241 | 0.001084382 | 0.005760258 | 0.006518177 | 0.157298441 | 0.146742039 | 0.09797851 | 0.02467993 | 0.03129395 |
| | | | 3180 | 1.001286421 | -0.390650513 | 0.704001573 | -0.583565534 | 0.06381626 | -0.337916694 | 1.249910334 | -0.506352885 | 0.352833732 | -0.30899991 |
| 52a | Unknown | Unknown | 3235 | 3.436393154 | -1.432730716 | 0.11329862 | -0.577111582 | 0.000202759 | -0.093065086 | 1.845666304 | -1.257881828 | 0.327972145 | -0.755433752 |
| | | | 3236 | 3.775518735 | -1.463631977 | 0.119183439 | -0.693049095 | 0.000202759 | -0.230080739 | 1.796579931 | -1.226465846 | 0.341069718 | -0.824276529 |
| | | | 3237 | 4.025032417 | -1.699489125 | 0.2914418 | -0.957766693 | 0.001985343 | -0.427740344 | 2.207548424 | -1.493525074 | 0.574291609 | -1.08696274 |
| | | | 3238 | 3.830299121 | -1.547854698 | 0.136903401 | -0.769227963 | 0.004851778 | -0.400034689 | 2.181123021 | -1.409172378 | 0.526288112 | -1.077515145 |
| | | | 3239 | 3.690785166 | -1.560601257 | 0.159486293 | -0.801022107 | 0.002964065 | -0.372207724 | 2.096542864 | -1.409895467 | 0.469148494 | -1.02338488 |
| 52b | Unknown | Unknown | 3240 | 0.504303624 | 0.449855498 | 0.069670121 | 0.400063323 | 0.000202759 | -0.190544163 | 0.235285934 | 0.478440192 | 0.073337048 | -0.263826956 |
| | | | 3241 | 0.286364589 | 0.354903695 | 0.036192153 | 0.323144864 | 0.000202759 | -0.202501964 | 0.173567334 | 0.454423196 | 0.068215878 | -0.26676095 |
| | | | 3242 | 0.183219996 | 0.254560486 | 0.033490544 | 0.306114417 | 0.000202759 | -0.227566869 | 0.138398255 | 0.403588279 | 0.056408353 | -0.259651684 |
| | | | 3243 | 0.178961921 | 0.259729782 | 0.047218354 | 0.381160214 | 0.000202759 | -0.098904497 | 0.161559399 | 0.480089829 | 0.03546796 | -0.17281077 |
| | | | 3244 | 0.337641708 | 0.439362069 | 0.05562578 | 0.444558421 | 0.000202759 | -0.053507431 | 0.216817162 | 0.585976387 | 0.007189956 | -0.0348756 |
| 53 | Unknown | ECF | 3269 | 1.008744734 | -0.265952795 | 0.062208833 | -0.154832209 | 0.062040053 | -0.227404053 | 68.99567863 | 2.425129988 | 0.060038884 | -0.052159629 |
| | | | 3270 | 0.071124802 | -0.082188068 | 0.136903401 | 0.483963935 | 0.001687904 | 0.277622593 | 175.7423214 | 5.428526779 | 0.249885411 | 0.394803872 |
| | | | 3271 | 0.070479862 | 0.036199016 | 0.042083336 | -0.129436944 | 0.000202759 | 0.004285507 | 300 | 5.846606829 | 0.052056869 | 0.066884189 |
| | | | 3272 | 0.450536413 | -0.359298951 | 0.097677999 | -0.391015718 | 0.000202759 | 0.013033854 | 241.4329736 | 5.934346347 | 0.048710444 | 0.129676108 |
| | | | 3273 | 1.03075295 | -0.832125343 | 0.086396171 | -0.420141794 | 0.356160488 | -1.084316058 | 147.89279 | 5.475255829 | 0.064603237 | -0.162683583 |
| | | | 3274 | 0.048430989 | -0.039292439 | 0.07028867 | 0.275784242 | 0.000202759 | -0.066357722 | 300 | 5.960249125 | 0.340382573 | 0.30863701 |
| | | | 3275 | 0.275606126 | -0.117891977 | 0.081674782 | 0.166305951 | 0.001687904 | -0.130050076 | 300 | 5.421502228 | 0.004297005 | 0.003594601 |
| | | | 3276 | 0.358011417 | -0.1650568 | 0.0879532 | 0.162140618 | 0.000202759 | 0.004758061 | 300 | 5.476162851 | 0.010760479 | 0.011269793 |
| 54 | Unknown | ECF | 3277 | 0.469933461 | 0.266066733 | 0.005185202 | -0.043627817 | 0.000202759 | 0.114110797 | 0.29400234 | 0.298291335 | 0.046072041 | -0.082152922 |
| | | | 3278 | 0.578356582 | 0.715376319 | 0 | 0.0354606 | 0.001687904 | 0.426321406 | 0.182359462 | 0.501768513 | 0 | 0.279106272 |
| | | | 3279 | 0.041433036 | 0.037836483 | 0.00230855 | -0.027827889 | 0.002964065 | 0.23259586 | 2.583970264 | 0.913482121 | 0.522154261 | 0.508816474 |
| | | | 3280 | 0.017584133 | -0.02219399 | 0.002741709 | 0.04900646 | 0.002964065 | 0.283341657 | 1.36208725 | 0.911877066 | 0.246366528 | 0.400852858 |
| | | | 3281 | 0.020027056 | 0.035203813 | 0 | 0.029489863 | 0.000202759 | 0.05799948 | 0.841615592 | 0.971638906 | 0.157638318 | 0.414997576 |
| | | | 3282 | 0.10178288 | -0.047334575 | 1.95089E-05 | 0.00088138 | 0.024985102 | -0.203455786 | 0.209633625 | 0.150579861 | 0.021069188 | 0.024986651 |
| 55 | Unknown | HTCS | 3292 | 1.342269143 | -0.595265735 | 0.961893977 | -0.890456788 | 0.123624592 | -0.556751553 | 0.356008544 | -0.334604309 | 1.369702809 | -0.805513996 |
| | | | 3293 | 1.787208693 | -0.255132767 | 0.268476964 | -0.223287539 | 0.024985102 | -0.157759077 | 0.065753763 | -0.041619553 | 1.941409514 | -0.319920879 |
| | | | 3294 | 0.227142927 | -0.048821031 | 0.004858317 | 0.023203603 | 0.000202759 | -0.060563757 | 0.676275337 | 0.146811842 | 0.773496728 | -0.196711471 |
| | | | 3295 | 0.789694937 | -0.663200136 | 0.002406745 | -0.043414459 | 0.024985102 | -0.600637358 | 0.031989336 | -0.096292384 | 0.229566737 | -0.485541874 |
| | | | 3296 | 1.357818991 | -0.686950905 | 0.003499596 | 0.04164197 | 0.064022577 | -0.513156469 | 0.244255663 | -0.435967802 | 0.478306439 | -0.634527611 |
| | | | 3297 | 0.19714804 | -0.155741772 | 0.005185202 | -0.047808969 | 0.02375206 | -0.433175366 | 0.155411899 | -0.453141124 | 1.285855442 | -1.014902927 |
| | | | 3298 | 0.341985778 | -0.296180913 | 0.057429347 | 0.249169044 | 0.018781022 | -0.346586645 | 0.234222743 | -0.496261514 | 0.306316693 | -0.598999606 |
| | | | 3299 | 0.724534995 | -0.547134968 | 0.039756938 | 0.179914972 | 0.000202759 | -0.035521623 | 0.07661626 | -0.188516101 | 0.036008543 | -0.111921685 |
| | | | 3300 | 0.682789404 | -0.654168004 | 0.069307866 | 0.339084479 | 0.000202759 | -0.2269688 | 0.000386725 | 0.001294442 | 0.021069188 | 0.06899298 |
| | | | 3301 | 0.506521584 | 0.416020852 | 0.119183439 | 0.436181803 | 0.000202759 | 0.162105904 | 0.151544967 | 0.28892963 | 0.104493603 | 0.221925124 |
| | | | 3302 | 1.105258078 | -0.262906865 | 0.766370655 | -0.461113125 | 0.000202759 | 0.003183038 | 0.541325401 | -0.24555216 | 0.575751984 | -0.251972228 |
| 56 | β1,6-Glucan | SusR | 3309 | 0.461682647 | 0.121320312 | 0.001776386 | -0.009450899 | 0.004371729 | 0.110962885 | 0.076330004 | 0.055216917 | 0.037255298 | 0.031753415 |
| | | | 3310 | 1.183885566 | -0.295879375 | 0.205046956 | -0.284688532 | 0.004537328 | -0.130016361 | 1.159414852 | -0.409882448 | 0.560305009 | -0.898095159 |
| | | | 3311 | 0.10525514 | -0.055143878 | 0.008774443 | 0.039938446 | 0.024985102 | -0.258669701 | 0.381089661 | -0.273366997 | 1.250167092 | -1.069304253 |

| | | | | | | | | | | | | | |
|---|---|---|---|---|---|---|---|---|---|---|---|---|---|
| [illegible] | [illegible] | [illegible] | 3312 | 0.342565881 | 0.148729532 | 0.033337935 | 0.106897837 | 0.000202759 | -0.063919948 | 0.408405689 | -0.281777012 | 1.232506436 | -0.981304794 |
| | | | 3313 | 0.20895831 | -0.115897922 | 0.261186257 | 0.388938118 | 0.000202759 | 0.046231685 | 0.415140626 | -0.380398622 | 1.551411945 | -1.152486547 |
| | | | 3314 | 0.019247705 | -0.008198717 | 0.07028867 | 0.140954852 | 0.004851778 | -0.139023794 | 0.582635984 | -0.302243088 | 0.804946785 | -1.085124366 |
| 57 | Chondroitin Sulfate, Hyaluronic Acid | HTCS | 3324 | 0.237053367 | -0.106418344 | 0.031117471 | 0.102606702 | 0.000202759 | 0.1077209 | 0.427855468 | 0.290937645 | 300 | 5.386349354 |
| | | | 3328 | 0.010024447 | -0.006650309 | 0.004858317 | 0.037493501 | 0.000202759 | 0.042094898 | 0.197046982 | 0.169494592 | 300 | 7.833368216 |
| | | | 3329 | 0.134391539 | -0.067230892 | 0.012450752 | -0.056326909 | 0.001985343 | 0.142860709 | 0.02584005 | 0.028486777 | 300 | 7.952988778 |
| | | | 3330 | 0.790100995 | -0.400195735 | 0.213833626 | -0.427359433 | 0.004189015 | -0.208491785 | 0.030268172 | 0.046913795 | 300 | 7.593254565 |
| | | | 3331 | 2.107826786 | -0.662165407 | 0.396273723 | -0.502849423 | 0.006518177 | -0.198348357 | 0.336375824 | -0.255326684 | 300 | 7.554482006 |
| | | | 3332 | 1.78978256 | -0.419213986 | 0.790079675 | -0.56251732 | 0.120677539 | -0.283989343 | 0.781401248 | -0.304820474 | 300 | 7.561822205 |
| | | | 3333 | 0.783622778 | 0.407121535 | 0.081674782 | 0.272665038 | 0.000202759 | 0.121329024 | 0.002851763 | 0.006476151 | 151.6695862 | 4.388911256 |
| | | | 3334 | 0 | 0 | 0 | 0 | 0.006518177 | 0.120150394 | 0 | 0 | 112.025028 | 2.36926289 |
| | | | 3348 | 0.626014905 | -0.142385643 | 0.008774443 | 0.02929808 | 0.047316364 | -0.174595437 | 0.326024022 | -0.143741038 | 300 | 5.80959991 |
| | | | 3349 | 0.548649195 | -0.127691897 | 0.02416949 | -0.056636199 | 0.06381626 | -0.204147609 | 0.101525844 | -0.062313447 | 300 | 5.804582233 |
| | | | 3350 | 0.174488895 | -0.067204159 | 0.00230855 | 0.01527116 | 0.000202759 | -0.10208786 | 0.184816022 | 0.113554552 | 300 | 5.851026022 |
| | | | 4410 | 0.506102103 | -0.172931738 | 0.00123049 | -0.007391146 | 0.000202759 | -0.099767208 | 0.056268098 | -0.057775364 | 300 | 7.370277168 |
| | | | 4411 | 0.432160235 | -0.339006557 | 0.040641621 | 0.191857493 | 0.000202759 | 0.119320893 | 0.066531274 | 0.122601915 | 300 | 7.258724909 |
| 58 | Unknown | Unknown | 3344 | 0.160900553 | 0.261467679 | 0 | -0.340400647 | 0.000202759 | -0.084242498 | 0.075024279 | 0.271806409 | 0 | -0.003856398 |
| | | | 3345 | 0.27994573 | -0.448192531 | 0 | -0.944600507 | 0.004537328 | -0.530606503 | 0.098820464 | -0.347413454 | 0.032893916 | -0.152182543 |
| | | | 3346 | 0.153788463 | 0.179542456 | 0.388667317 | -1.01279991 | 0.008810859 | -0.445682888 | 0.063802018 | 0.183885436 | 0.133027292 | -0.338795778 |
| | | | 3347 | 0.346108084 | 0.351041569 | 0.021295324 | -0.17163627 | 0.097494468 | -0.89100391 | 0.103455492 | -0.274726914 | 0.021069188 | -0.075580729 |
| 59 | Unknown | HTCS | 3461 | 0.007621322 | -0.003042132 | 0.195073704 | 0.25868792 | 0.000202759 | -0.01267537 | 0.177383853 | -0.112700244 | 0.158418836 | -0.103251988 |
| | | | 3462 | 0.019891398 | -0.006354744 | 0.05128535 | 0.097135732 | 0.000202759 | 0.036268526 | 0.145183382 | -0.074130382 | 0.092597872 | 0.059728595 |
| | | | 3463 | 0.585939313 | -0.136046002 | 0.025030963 | -0.066566944 | 4.81016E-05 | 0.0000553 | 0.221896723 | -0.106888243 | 0.118230181 | 0.068062906 |
| | | | 3464 | 0.757930487 | 0.217034836 | 0.437865006 | 0.404168553 | 0.293418619 | 0.41087733 | 0.435184807 | 0.230663214 | 1.243277209 | 0.402737201 |
| | | | 3465 | 0.05713929 | -0.01713019 | 0.02416949 | 0.056228256 | 0.022858763 | 0.169107133 | 0.02584005 | 0.017873867 | 0.168668256 | 0.088024286 |
| | | | 3466 | 0.134545834 | -0.38032894 | 0 | -1.277366373 | 0.004851778 | -0.819851296 | 0.173567334 | 0.660173452 | 0 | 1.021106445 |
| | | | 3467 | 0.182544856 | -0.057873164 | 0.258448928 | -0.252911141 | 0.024985102 | -0.171767835 | 1.503724651 | -0.403051503 | 2.069130147 | -0.414094649 |
| | | | 3468 | 0.571426544 | -0.172706782 | 0.305657373 | -0.350762343 | 0.000202759 | -0.102722036 | 0.435679977 | -0.216419747 | 0.472306766 | -0.22613732 |
| | | | 3469 | 0.01484016 | -0.00513591 | 0.002406745 | 0.014514341 | 0.227998176 | 0.265323651 | 1.596986057 | 0.338268501 | 0.589084403 | 0.187009148 |
| | | | 3470 | 1.807559146 | -0.313577271 | 0.098306198 | -0.166097092 | 0.000202759 | 0.043970982 | 0.287934158 | 0.147332638 | 0.065153495 | -0.048491428 |
| | | | 3471 | 1.823805623 | -0.603491242 | 0.289638968 | -0.691031778 | 0.101367857 | -0.464656946 | 0.357541066 | 0.314424245 | 0.053692678 | -0.082747661 |
| | | | 3472 | 0.86763624 | -0.191215567 | 0.138893965 | -0.193326076 | 0.000202759 | -0.046841709 | 0.29819014 | 0.136628831 | 0.186434536 | -0.108343001 |
| | | | 3473 | 1.795674319 | 1.643243879 | 0 | 1.585301918 | 0.113478827 | 1.07039033 | 2.7795786 | 2.337444276 | 0 | 1.561671954 |
| | | | 3474 | 1.034667066 | 0.935589449 | 0.043712581 | 1.150265657 | 0.037279308 | 0.671249503 | 3.830943067 | 1.961165355 | 0.540928449 | 0.868945372 |
| | | | 3475 | 0.395940604 | 0.469943847 | 0.027187628 | 0.583781684 | 0.00129528 | 0.376903417 | 0.957943514 | 1.148941796 | 0.185141848 | 0.480987272 |
| | | | 3476 | 0.909129072 | 1.038701657 | 0 | 0.973926655 | 0.056730599 | 0.890352962 | 1.400478776 | 1.534703826 | 0 | 0.224592162 |
| | | | 3477 | 0.369949281 | -0.247476831 | 0.216990326 | -0.445570455 | 0.000202759 | -0.036829267 | 0.108103464 | -0.153664582 | 0.478587226 | -0.431583554 |
| | | | 3478 | 0.689135824 | 0.156952828 | 0.078947475 | 0.142796432 | 0.287954616 | 0.312031979 | 1.738439248 | 0.34241994 | 2.796044809 | 0.460881422 |
| | | | 3479 | 0.200450019 | -0.483947101 | 0.162665201 | -0.277130756 | 0.000202759 | 0.008640673 | 0.418549057 | -1.659535695 | 0.560305009 | -0.169758944 |
| | | | 3480 | 0.229041142 | -0.421393252 | 0.058008143 | -0.178064648 | 0.006518177 | 0.161746751 | 0.418821466 | -1.320927464 | 0.031631871 | 0.032520757 |
| | | | 3481 | 0.49773548 | 1.063749934 | 0 | -0.667775227 | 0.030771831 | -1.943356921 | 0.131865197 | 0.757288943 | 0 | -0.524065406 |
| | | | 3482 | 0.053075597 | 0.17846921 | 0.047218354 | -0.512177253 | 0.023373218 | -0.954149795 | 0.053967688 | 0.500986081 | 0.336267584 | -1.083520894 |
| | | | 3483 | 0.281948997 | 0.3533739 | 0.002741709 | -0.118043657 | 0.075866865 | -0.955370352 | 0.052663102 | 0.371076345 | 0.239049707 | -0.621363119 |
| | | | 3484 | 1.19657276 | 0.863159813 | 0.017461642 | -0.15566387 | 0.000202759 | -0.092926368 | 0.07314107 | 0.429562146 | 0.097232212 | -0.281019867 |
| | | | 3485 | 1.321417811 | 1.530913812 | 0 | 0.371004022 | 0.000202759 | 0.262909389 | 0.173567334 | 1.190519099 | 0 | 0.666947033 |
| | | | 3486 | 0.626006292 | 0.359744154 | 0.047801115 | -0.211850563 | 0.000202759 | -0.171128886 | 0.030510593 | 0.06781894 | 0.245352221 | -0.344069373 |
| | | | 3487 | 0.701447907 | 0.431466138 | 0.039756938 | -0.189999954 | 0.000202759 | -0.135047458 | 0.036145046 | -0.076605128 | 0.502178158 | -0.481400168 |
| | | | 3488 | 0.705199831 | 0.527450567 | 0.004534747 | -0.063771749 | 0.000202759 | 0.157024001 | 0.26137023 | 0.440971343 | 0.03546796 | 0.102667727 |
| | | | 3489 | 0.012902017 | -0.011462774 | 0.074888922 | -0.276210061 | 0.001985343 | -0.21987495 | 0.263265973 | -0.307834068 | 0.017996334 | -0.037779584 |
| | | | 3490 | 0.160493249 | -0.06173219 | 0.15722954 | -0.253611566 | 0.04531731 | -0.23213799 | 0.361161153 | -0.18350938 | 0.239140483 | -0.165616575 |
| | | | 3491 | 1.060402691 | -2.574152383 | 0 | -3.122227393 | 0.560801483 | -4.672387323 | 0.51698233 | -2.180628295 | 0 | -2.5832612 |
| | | | 3492 | 2.688937959 | -3.448882114 | 0 | -2.713275297 | 0.051118142 | -1.274738468 | 0.520488978 | -2.259796857 | 0.286011586 | -1.167673572 |
| | | | 3493 | 0.391040926 | -0.617875133 | 0 | -1.846056692 | 0.047316364 | -1.064617274 | 0.613113143 | -1.294100687 | 0.068653947 | -0.29588761 |
| | | | 3494 | 2.598315365 | -2.01867365 | 0.632183689 | -1.652178996 | 0.16571248 | -1.179127664 | 0.941708438 | -1.396610228 | 0.21488881 | -0.650276533 |
| | | | 3495 | 0.965377775 | -0.981491348 | 0.151191457 | -0.696916587 | 0.000525887 | -0.343834998 | 0.49192631 | -0.82947102 | 0.012272811 | -0.054690511 |
| | | | 3496 | 1.729121062 | -1.663631852 | 0.343595974 | -1.168416689 | 0.004851778 | -0.490616301 | 0.921273775 | -1.357947813 | 0.052710278 | -0.2087277 |
| | | | 3497 | 0.319844899 | -2.546134966 | 0 | -0.25707142 | 0.000202759 | -0.839442587 | 0 | 1.065169614 | 0 | 0.788650591 |
| | | | 3498 | 0.004260548 | 0.020525319 | 0 | -0.459346052 | 0.000202759 | 0.110756652 | 0 | -0.435264187 | 0 | -0.768354771 |
| | | | 3499 | 0.326785803 | -1.292546114 | 0 | -1.938505368 | 0.000202759 | 0.163764783 | 0 | -1.004433593 | 0 | -0.62543286 |
| | | | 3500 | 0.097934949 | 0.283067548 | 0 | 0.464370519 | 0.019865857 | 0.949216422 | 0.056241538 | -0.383828273 | 0 | 0.160529368 |
| | | | 3501 | 0.301844457 | -0.575205044 | 0 | -0.884948861 | 0.006793048 | -0.687669378 | 0.268330382 | -0.798627856 | 0 | 0.079788529 |
| | | | 3502 | 0.059115962 | -0.653615208 | 0 | -0.764140258 | 0.000202759 | -0.879346996 | 0 | -0.809361362 | 0 | -1.219077385 |
| | | | 3503 | 0.854042083 | -2.902246172 | 0 | -0.556668929 | 0.000202759 | 0.219579584 | 0 | -1.008706479 | 0 | -0.158620659 |
| | | | 3504 | 0.713426942 | -1.235427927 | 0 | -0.017345523 | 0.411258989 | 1.423123998 | 0.560237662 | -1.739144756 | 0 | 0.567322561 |
| | | | 3505 | 5.543340925 | -2.841010652 | 0.231650901 | -0.763985136 | 0.133352832 | 0.79494481 | 2.868954251 | -2.048555633 | 0.161855361 | 0.369804262 |
| | | | 3506 | 3.091957544 | -2.250370599 | 0.025030963 | -0.219582169 | 0.372914409 | 1.163199651 | 1.051471252 | -1.345816286 | 0.592225055 | 0.894029798 |
| | | | 3507 | 2.141955724 | -2.522889143 | 0 | 0.151007444 | 0.236933987 | 1.165604299 | 0.108769832 | -0.360062933 | 1.25347164 | 1.377066512 |
| 60 | Unknown | ECF | 3517 | 0.175027478 | -0.068160467 | 0.173473728 | -0.235852585 | 0.000202759 | -0.065406613 | 0.231727198 | -0.137436738 | 0.105538171 | -0.081648406 |
| | | | 3518 | 0.094249382 | -0.044545654 | 0.070919568 | -0.190852358 | 0.000202759 | 0.062265643 | 0.041150305 | 0.043838328 | 0.437149567 | 0.266922004 |
| | | | 3519 | 0.850748286 | -0.211682833 | 0.133625486 | -0.198681152 | 0.02474203 | -0.185176327 | 0.206056337 | -0.125470702 | 1.327963613 | -0.4210899 |
| | | | 3520 | 0.478756373 | -0.153240482 | 0.012450752 | 0.041305213 | 0.004851778 | -0.131861185 | 0.468258441 | -0.221444951 | 1.162552472 | -0.414862247 |
| | | | 3521 | 0.382427823 | -0.156147211 | 0.017736303 | 0.055881897 | 0.000202759 | -0.053671705 | 0.165626811 | -0.134595964 | 1.194164925 | -0.460408152 |
| | | | 3522 | 0.135388924 | -0.086467628 | 0.029046553 | 0.116737661 | 0.006518177 | 0.220622032 | 0.28471488 | 0.241043503 | 0.329580799 | -0.286025325 |
| | | | 3523 | 0.250240423 | 0.140721702 | 0.364990929 | 0.452384169 | 0.004851778 | 0.210156386 | 0.007918642 | -0.012022724 | 0.195235922 | -0.213196036 |
| | | | 3524 | 0.148918655 | 0.131970952 | 0.002406745 | 0.037664019 | 0.000202759 | -0.07398783 | 0.16647838 | -0.270999613 | 0.115221175 | -0.24364543 |
| | | | 3525 | 0.124434219 | -0.081665879 | 0.012450752 | -0.064911264 | 0.000202759 | -0.146400977 | 0.035584994 | -0.054009702 | 0.509408764 | -0.408637007 |
| | | | 3526 | 0.64312627 | -0.244252592 | 0.053599877 | 0.13731389 | 0.006518177 | -0.168497321 | 0.217385408 | -0.195240681 | 0.06663155 | -0.079022611 |
| | | | 3527 | 2.29703468 | -0.382576577 | 0.839290822 | -0.431148078 | 0.049606671 | -0.224227316 | 0.160498548 | -0.123491763 | 2.125607486 | -0.421689818 |
| | | | 3528 | 0.703939861 | -0.245559317 | 0.231537557 | -0.315827882 | 0.000202759 | -0.10489229 | 0.040307982 | 0.044707087 | 0.245352221 | -0.176277763 |
| | | | 3529 | 0.797319383 | -0.229435242 | 0.22460127 | -0.259549825 | 0.000202759 | -0.018036875 | 0.007965004 | 0.008287268 | 0.403810517 | -0.186586524 |
| | | | 3530 | 0.833927804 | -0.250598367 | 0.15722954 | -0.216718965 | 0.000202759 | -0.047289886 | 0.046395686 | 0.049129679 | 0.293487858 | -0.157443188 |
| | | | 3531 | 0.725789426 | -0.217432241 | 0.186074744 | -0.286048202 | 0.000202759 | -0.076178615 | 0.201350477 | 0.120944753 | 0.152993699 | -0.105108672 |
| | | | 3532 | 0.201623726 | -0.101109503 | 0.133074163 | -0.25153461 | 0.000202759 | 0.035366012 | 0.465558545 | -0.288993564 | 0.484378751 | -0.295871148 |
| 61 | Unknown | Unknown | 3566 | 4.45525475 | 0.56686299 | 0.000400818 | 0.002619272 | 0.000202759 | -0.053691862 | 10.30627305 | -0.758530852 | 0.53417938 | -0.259394021 |
| | | | 3567 | 2.326716106 | 0.436439816 | 0.085350226 | -0.149794113 | 0.000202759 | -0.099863609 | 9.313363731 | -0.838066599 | 0.464683954 | -0.284955599 |
| | | | 3568 | 3.170723894 | 0.419600635 | 0.05562578 | -0.097425651 | 0.000202759 | -0.013532279 | 15.97061622 | -0.803337089 | 0.647791738 | -0.283846278 |
| | | | 3569 | 1.8916336 | 0.340234349 | 0.273541724 | -0.249658756 | 0.000202759 | -0.132521678 | 14.86646109 | -0.889050521 | 1.048475085 | -0.371481391 |
| | | | 3570 | 0.192235326 | -0.078420439 | 0.108681784 | -0.227344563 | 0.020797091 | -0.196185586 | 0.017575703 | -0.019830542 | 3.283837166 | 0.607226662 |
| | | | 3581 | 1.163702829 | -0.220041013 | 0.172447079 | -0.198943424 | 0.000202759 | -0.032210648 | 0.252840848 | 0.133387986 | 0.02467993 | -0.023640705 |
| | | | 3582 | 0.280772923 | -0.188154332 | 0.063161503 | -0.232467957 | 0.000202759 | -0.079753038 | 0.065073717 | 0.104731333 | 0.044397487 | 0.08582913 |
| | | | 3583 | 0.27488184 | -0.090849788 | 0.008774443 | -0.035104052 | 0.000202759 | -0.066389668 | 0.155097962 | 0.13229264 | 0.011742117 | 0.012383097 |
| | | | 3584 | 0.124414711 | -0.061310746 | 0.133625486 | -0.2590998 | 0.000202759 | -0.107369471 | 0.008085793 | -0.011463728 | 0.405358102 | -0.290922655 |
| | | | 3585 | 0.135647851 | -0.050517886 | 0.01875079 | -0.04839894 | 0.000525887 | -0.1078009 | 0.291101598 | -0.174474355 | 0.400605803 | -0.229543702 |
| | | | 3586 | 0.100630827 | -0.043760849 | 0.029046553 | -0.086545114 | 0.000202759 | -0.082775954 | 0.76505073 | -0.30773497 | 0.363571276 | -0.21091906 |
| | | | 3587 | 0.123409057 | 0.101784563 | 0.32457274 | 0.560288642 | 0.006518177 | -0.289423238 | 0.135325575 | 0.225955274 | 0.03546796 | 0.077013021 |
| | | | 3588 | 0.082150854 | -0.058436193 | 0.142377104 | 0.311138807 | 0.000202759 | -0.067329632 | 0.055162202 | -0.119342486 | 0.304519889 | -0.357699842 |
| | | | 3589 | 0.354807687 | 0.193900359 | 0.004534747 | -0.039267293 | 0.000202759 | -0.152630265 | 0.001085528 | -0.002275112 | 0.496312802 | -0.439752345 |
| | | | 3590 | 0.237771518 | 0.122835268 | 0.13622255 | 0.274367709 | 0.000202759 | 0.028042021 | 0.076469374 | -0.125222738 | 0.196458646 | -0.212455056 |
| | | | 3591 | 1.035957887 | 0.373169015 | 0.287580336 | 0.454464996 | 0.000202759 | 0.076167822 | 0.022093468 | 0.042621224 | 0.229566737 | -0.26443676 |
| | | | 3592 | 0.030403209 | -0.023977739 | 0.157869687 | 0.374901341 | 0.007237187 | -0.278166584 | 0.021794295 | -0.078112205 | 0.474758046 | -0.50231563 |
| | | | 3593 | 0.543678021 | 0.321441669 | 0.118636734 | 0.338752932 | 0.000202759 | -0.034656132 | 0.042158373 | 0.094030546 | 0.168668256 | -0.319954053 |
| | | | 3594 | 0.783989101 | 0.419337825 | 0.149252365 | 0.397968979 | 0.000202759 | -0.125668994 | 0.007965004 | 0.034558865 | 0.276844537 | -0.431035203 |
| 62 | Unknown | Other | 3595 | 1.186080294 | 0.49455074 | 0.564955016 | 0.669556909 | 0.000202759 | -0.200214712 | 0.038173492 | 0.087954364 | 0.409419514 | -0.526062398 |

| | | | | | | | | | | | | | |
|---|---|---|---|---|---|---|---|---|---|---|---|---|---|
| | | | 3596 | 0.361822498 | 0.495665841 | 0.07028867 | 0.353743274 | 0.000202759 | -0.087811775 | 0.008982821 | 0.044124586 | 0.083555031 | -0.346041913 |
| | | | 3597 | 0.63221037 | 0.412042076 | 0.201050693 | 0.505818874 | 0.000202759 | -0.071662174 | 0.038173492 | 0.184318951 | 0.038390496 | -0.106623833 |
| | | | 3598 | 1.082414697 | 0.57323115 | 0.623528953 | 0.783046513 | 0.000202759 | 0.256781064 | 0.127197608 | 0.369819199 | 0.141861057 | 0.282372084 |
| | | | 3599 | 0.603756551 | 0.412634004 | 0.159486293 | 0.489500619 | 0.000202759 | 0.219344399 | 0.145183382 | 0.515256922 | 0.06893846 | 0.175876446 |
| | | | 3600 | 1.209250291 | 0.800454455 | 0.342796909 | 0.804596648 | 0.037591022 | 0.623303356 | 0.623569551 | 0.90760264 | 0.285540285 | 0.587486458 |
| | | | 3601 | 0.100486214 | 0.095612549 | 0.123120774 | 0.44253248 | 0.008620635 | 0.392016147 | 0.168095185 | 0.558104493 | 0.109140466 | 0.267911137 |
| | | | 3602 | 0.178108503 | 0.268480003 | 0.02416949 | 0.234681498 | 0.000202759 | -0.118928771 | 0.05556034 | 0.295785417 | 0.065929575 | -0.31329933 |
| | | | 3603 | 0.245851346 | 0.269652502 | 0.17241522 | 0.414348635 | 0.000202759 | 0.211607582 | 0.139717915 | 0.50177674 | 0.02467993 | -0.114307459 |
| | | | 3604 | 0.060291172 | 0.083343404 | 0.061645108 | 0.253488916 | 0.000202759 | 0.095057761 | 0.07314107 | 0.323248683 | 0.088309791 | -0.344602611 |
| | | | 3605 | 0.069629611 | -0.063517832 | 0.021295324 | -0.114325388 | 0.000202759 | -0.011245937 | 0.035584994 | 0.175768074 | 0.0168207 | -0.089681552 |
| | | | 3606 | 0.205251956 | -0.28786568 | 0.067014811 | -0.308978415 | 0.000202759 | -0.108287649 | 0.004263331 | 0.025708654 | 0.107064324 | -0.416496609 |
| | | | 3607 | 0.031974447 | -0.026119247 | 0.02416949 | 0.11382964 | 0.000202759 | 0.079608911 | 0.027271487 | 0.150753101 | 0.013646459 | -0.071808163 |
| | | | 3608 | 0.357519532 | -0.160024919 | 0.070919568 | -0.178543521 | 0.001985343 | -0.145878196 | 0.100353145 | -0.09582669 | 0.032188006 | -0.044748817 |
| | | | 3609 | 0.675181736 | -0.187774038 | 0.047801115 | -0.10940831 | 0.002964065 | -0.106789549 | 0.343012128 | -0.152544824 | 0.452631958 | -0.212376466 |
| 64 | Rhamnogalacturonan II | Unknown | 3662 | 1.622344104 | -0.512795937 | 0.552535981 | -0.553685586 | 0.104072316 | -0.391248815 | 0.68965487 | -0.380122561 | 0.516638448 | -0.331947163 |
| | | | 3663 | 0.051892939 | -0.050486164 | 0.032272961 | -0.176877118 | 0.000202759 | 0.119070514 | 0.187491453 | 0.28301356 | 0.042088494 | 0.099980811 |
| | | | 3664 | 0.265130515 | -0.186705616 | 0.003600372 | 0.040700848 | 0.004537328 | 0.220729344 | 0.170367402 | 0.205296068 | 0.060038884 | 0.108056312 |
| | | | 3665 | 0.391387388 | -0.153955723 | 0.248157356 | -0.328657254 | 0.000202759 | 0.027555046 | 0.378161598 | -0.234469456 | 0.453768389 | -0.268498058 |
| | | | 3666 | 0.546784713 | -0.31950945 | 0.295120544 | -0.521444302 | 0.039312497 | -0.366677423 | 1.332439085 | -0.761129486 | 1.321843374 | -0.83023401 |
| | | | 3667 | 0.757125021 | -0.378403348 | 0.371545701 | -0.553780212 | 0.024951395 | -0.30537894 | 0.522464037 | -0.496131881 | 0.480144141 | -0.425938508 |
| | | | 3668 | 0.141615459 | -0.340575966 | 0 | 0.08885723 | 0.000202759 | -0.236975518 | 0 | -4.13421921 | 0 | -0.082852958 |
| | | | 3669 | 0.437023623 | 0.984336767 | 0 | 1.134937944 | 0.013169173 | 0.943421688 | 0.215295641 | 0.917052889 | 0 | 1.474532741 |
| | | | 3670 | 0.281468427 | -0.331137516 | 0.05794275 | -0.370741791 | 0.100424356 | -0.854929114 | 0.018338495 | 0.057240362 | 0.021069188 | -0.074955581 |
| | | | 3671 | 0.566200237 | -0.685534931 | 0.029046553 | -0.21458834 | 0.084278937 | -0.824849657 | 0.062016042 | -0.169804677 | 0.003328382 | 0.00823959 |
| | | | 3672 | 0.483604854 | 0.930672854 | 0 | -0.394684394 | 0.000202759 | 0.451465027 | 0.105657817 | 0.436726619 | 0 | -0.177873805 |
| 65 | Rhamnogalacturonan II | HTCS | 3674 | 0.120318167 | 0.063941576 | 0.004858317 | 0.032629834 | 0.006518177 | -0.180003639 | 0.027484529 | -0.033230197 | 0.856232317 | -0.399018049 |
| | | | 3675 | 0.173418029 | -0.135913571 | 0.028406201 | 0.140267428 | 0.000202759 | 0.167284967 | 0.267386307 | -0.340747029 | 0.224540654 | -0.321905646 |
| | | | 3676 | 0.250336645 | -0.171585413 | 0.123599436 | -0.334466068 | 0.000202759 | 0.026336722 | 0.088644925 | -0.14059978 | 0.053692678 | 0.095144698 |
| | | | 3677 | 0.547905541 | -0.262161398 | 0.089640486 | -0.220315809 | 0.000202759 | 0.014910885 | 0.028528329 | 0.041135826 | 0.046927295 | 0.069033284 |
| | | | 3678 | 1.048717359 | -0.710712742 | 0.113740599 | -0.268739175 | 0.804307694 | -0.94803373 | 9.950781977 | -2.064261945 | 0.359803952 | -0.508165486 |
| | | | 3679 | 0.007493729 | 0.018999481 | 0 | -0.557365019 | 4.81016E-05 | -0.000147509 | 0.14217371 | -0.595479176 | 0 | -0.305555487 |
| | | | 3680 | 0.219166128 | -0.257406845 | 0.004534747 | -0.076198925 | 0.010532763 | -0.47233432 | 0.218524188 | -0.462489333 | 0.279364108 | -0.562009583 |
| | | | 3681 | 0.610238392 | -0.588214456 | 2.5969E-05 | 0.00513157 | 0.006518177 | -0.364374724 | 1.082429771 | -1.136334627 | 0.299969002 | -0.595846516 |
| | | | 3682 | 0.046664265 | -0.091274663 | 0 | -0.290963899 | 0.000202759 | -0.12505976 | 0.213341119 | -0.644061652 | 0 | -1.038203517 |
| | | | 3683 | 1.147767265 | 0.653906465 | 0.286676738 | 0.595344511 | 0.05625593 | 0.479188205 | 0.046395686 | 0.109697423 | 0.274399919 | 0.420248041 |
| | | | 3684 | 0.052828477 | -0.378457159 | 0 | 1.046720915 | 0.001985343 | -2.365676175 | 0 | -1.332242058 | 0 | -0.299962486 |
| | | | 3685 | 0.383783133 | 0.142826344 | 0.340692504 | 0.380612129 | 0.311600082 | 0.457558081 | 0.211364251 | 0.163827836 | 0.400605803 | 0.217801272 |
| | | | 3686 | 0.771166879 | 0.264396976 | 0.21432977 | 0.342918601 | 0.145883736 | 0.401161088 | 0.259322232 | 0.219574466 | 1.51814589 | 0.57588063 |
| | | | 3687 | 1.591736929 | 0.294236452 | 0.096977493 | 0.157122057 | 0.467401413 | 0.415996855 | 2.010399985 | 0.420343483 | 4.153662888 | 0.570371278 |
| 66 | Amylopectin, Glycogen, Pullulan | SusR | 3698 | 0.002763426 | -0.002057456 | 0.001921284 | -0.016174929 | 0.000202759 | 0.028335249 | 0.038173492 | 0.054995777 | 0.071057962 | 0.092310179 |
| | | | 3699 | 0.234112702 | 0.176993386 | 0.05128535 | 0.204850387 | 0.000202759 | 0.11561237 | 0.34072836 | 0.304690492 | 0.210337489 | -0.277803402 |
| | | | 3700 | 0.170627148 | 0.125345899 | 0.012450752 | 0.07600147 | 0.000202759 | -0.030857464 | 0.010733949 | -0.019778917 | 0.245352221 | -0.311073388 |
| | | | 3701 | 0.110072075 | -0.079096126 | 0.282272956 | -0.494020886 | 0.000202759 | -0.090581159 | 1.19877401 | -0.696090403 | 0.878711739 | -0.638407003 |
| | | | 3702 | 1.69265383 | -0.443434135 | 0.632183689 | -0.570307562 | 0.091379945 | -0.320575988 | 1.944540437 | -0.575212628 | 3.870932355 | -0.903407486 |
| | | | 3703 | 0.247124532 | -0.14666917 | 0.07037195 | -0.216498977 | 0.058901468 | 1.00825009 | 0.341831888 | -0.291660355 | 2.081010764 | -0.875354558 |
| | | | 3704 | 0.108681238 | -0.125029333 | 0.358586521 | -0.835926305 | 0.506287619 | 2.867714327 | 0.09549372 | -0.188805856 | 0.719921547 | -0.867395878 |
| | | | 3705 | 0.052778908 | 0.021067284 | 0.050460707 | -0.119337895 | 0.000202759 | -0.047689461 | 0.272920718 | -0.155311068 | 0.03546796 | -0.035208549 |
| 67 | Unknown | ECF | 3748 | 1.296369042 | 0.447313024 | 0.036192153 | 0.131240663 | 0.043157144 | 0.309761292 | 0.030779173 | 0.045320169 | 0.163566245 | 0.167275887 |
| | | | 3749 | 0.366029928 | -0.218277061 | 0.468808804 | -0.635298671 | 0.024951395 | -0.324202759 | 0.433108492 | -0.47153397 | 0.085374999 | -0.128942413 |
| | | | 3750 | 0.569996628 | 0.198680586 | 0.089640486 | 0.1815273 | 0.000202759 | 0.083470466 | 0.043252873 | -0.057100382 | 0.258644131 | -0.190780092 |
| | | | 3751 | 0.276975731 | 0.280798605 | 0.091477514 | 0.386298014 | 0.000202759 | 0.127520982 | 0.01749368 | 0.051564019 | 0.133027292 | -0.273634697 |
| | | | 3752 | 0.080817855 | -0.058888144 | 0.018104878 | 0.072778887 | 0.024985102 | -0.281807967 | 0.067640727 | -0.0913911 | 0.442491578 | -0.372263659 |
| | | | 3753 | 0.109206196 | -0.118606322 | 0.014195748 | 0.096425797 | 0.000202759 | 0.269922007 | 0.286405588 | 0.402499381 | 0.151952698 | -0.308235338 |
| | | | 3754 | 0.00354992 | -0.002629232 | 0.017461642 | 0.069346608 | 0.000202759 | -0.008442498 | 0.832515349 | -0.517388049 | 0.047935946 | -0.071497229 |
| 68 | α-mannan | HTCS | 3773 | 0.420863327 | -0.115198637 | 0.258448928 | 0.234677935 | 0.000202759 | 0.052191064 | 0.564517116 | -0.206189681 | 0.015961708 | -0.0131528 |
| | | | 3774 | 1.45048521 | -0.438145545 | 1.055844075 | -0.55701136 | 0.000202759 | -0.043080603 | 0.198603873 | 0.133915054 | 0.109140466 | -0.10430768 |
| | | | 3775 | 0.336321272 | 1.112756676 | 0 | 0.272625477 | 0.000202759 | 0.436505701 | 0 | 1.913997052 | 0 | 1.410195142 |
| | | | 3776 | 0.167944941 | -0.650776748 | 0 | 1.702559094 | 0.000525887 | 0.737100978 | 0.435450452 | 1.367252887 | 0 | 1.392395834 |
| | | | 3777 | 0.693423802 | 1.338963068 | 0 | 0.686473387 | 0.000202759 | 0.256817248 | 0 | 0.236567207 | 0 | 1.676715258 |
| | | | 3778 | 0.658008248 | 1.118140691 | 0 | 1.026932794 | 0.000202759 | -0.20864751 | 0.41768881 | 1.129627418 | 0 | 1.877861101 |
| | | | 3779 | 0.158053521 | 0.269781 | 0 | -0.959656986 | 0.000202759 | -0.335739503 | 0.034076797 | -0.152762312 | 0 | 0.500440612 |
| | | | 3780 | 0.047343301 | 0.022786599 | 0.047801115 | -0.127039332 | 0.000202759 | 0.025151391 | 0.841615592 | 0.289047045 | 0.175681484 | -0.125419428 |
| | | | 3781 | 2.000614962 | -0.355861461 | 0.724764987 | -0.373398931 | 0.037279308 | -0.177214197 | 0.06355942 | -0.039520817 | 0.91870498 | -0.26982203 |
| | | | 3782 | 1.103299769 | -0.223936602 | 0.22460127 | -0.243599209 | 0.004537328 | -0.106400559 | 0.028528329 | 0.021931999 | 1.416971519 | -0.311419736 |
| | | | 3783 | 0.383032537 | -0.094706203 | 0.121551038 | -0.165092075 | 0.069316676 | -0.196803181 | 0.170436508 | -0.08965281 | 3.459595438 | -0.477443305 |
| | | | 3784 | 0.258355122 | 0.107540725 | 0.029046553 | -0.083504738 | 0.004537328 | -0.139117687 | 1.755859202 | -0.520803443 | 5.364516253 | -0.926363858 |
| | | | 3785 | 0.152749162 | -1.537302843 | 0 | -0.682813503 | 0.000202759 | -1.760680723 | 0 | -0.031920753 | 0 | -0.150294341 |
| | | | 3786 | 1.416501723 | -0.359174314 | 0.612111244 | -0.439891175 | 0.006518177 | -0.156676046 | 1.256880343 | -0.402801049 | 2.521995999 | -0.62853231 |
| | | | 3787 | 0.959451183 | -0.441967422 | 0.659731745 | -0.627348514 | 0.314795922 | -0.542914262 | 0.630867801 | -0.393020638 | 1.732844329 | -0.716122148 |
| | | | 3788 | 1.537781583 | -0.341979961 | 0.694990811 | -0.505840483 | 0.180575102 | -0.343213529 | 0.149389978 | -0.112755947 | 3.385594279 | -0.663920798 |
| | | | 3789 | 1.876941556 | -0.462210138 | 0.43852202 | -0.45598913 | 0.218410058 | -0.380741256 | 0.258337948 | -0.203539161 | 1.054746932 | -0.427864577 |
| | | | 3790 | 0.305460682 | -0.237253674 | 0.121551038 | -0.439477321 | 0.000202759 | -0.02020994 | 0.235285934 | -0.325247002 | 0.014321959 | -0.037800217 |
| | | | 3791 | 0.121507246 | -0.066489384 | 0.107867327 | -0.264284249 | 0.000202759 | -0.005270578 | 0.272940587 | 0.205947557 | 0.563688913 | -0.400446223 |
| | | | 3792 | 0.908631884 | 0.278751381 | 0.01706425 | -0.052554662 | 0.000202759 | -0.018697382 | 0.241985483 | 0.160787843 | 0.277191349 | -0.171938229 |
| 69 | α-mannan | Other | 3853 | 0.057429692 | 0.018047512 | 0.074217884 | -0.131935272 | 0.000525887 | -0.085566574 | 0.344901487 | -0.128361751 | 0.382601203 | -0.143016782 |
| | | | 3854 | 1.412150756 | -0.419959599 | 0.285456893 | -0.356379945 | 0.119882068 | -0.376130942 | 0.291966355 | -0.209718932 | 2.184679186 | -0.643298759 |
| | | | 3855 | 1.935971313 | -0.666885893 | 0.228817236 | -0.399502109 | 0.001985343 | -0.168513742 | 0.111016255 | -0.135390802 | 0.629791117 | -0.455450039 |
| | | | 3856 | 0.289952485 | -0.447516678 | 0 | 0.253776202 | 0.047316364 | 0.706224287 | 0.08290293 | 0.242180961 | 0.06162748 | 0.213716907 |
| | | | 3857 | 0.603404685 | -0.319252447 | 0.029046553 | -0.118837982 | 0.000202759 | -0.141510806 | 0.044854925 | -0.066598692 | 0.121450853 | -0.166984242 |
| | | | 3858 | 0.092990603 | -0.078844133 | 0.021928238 | 0.101425343 | 0.000202759 | -0.025032161 | 0.128834957 | -0.172243603 | 0.068215878 | -0.125090555 |
| | | | 3859 | 0.637037942 | -0.417602558 | 0.04436107 | -0.215296618 | 0.008620635 | -0.285860677 | 0.01969829 | -0.043938694 | 0.023253432 | -0.053777579 |
| | | | 3860 | 0.777237631 | -0.518319176 | 0.029640605 | -0.162737263 | 0.000202759 | -0.075872405 | 0.323027273 | -0.424231117 | 0.313136231 | -0.422164679 |
| | | | 3861 | 0.181746766 | 0.169439211 | 0.005185202 | 0.06082513 | 0.000202759 | 0.183051788 | 0.046395686 | 0.105054532 | 0.042415157 | -0.105582721 |
| | | | 3862 | 0.456368882 | 0.348947192 | 0.17241522 | 0.463902591 | 0.000202759 | 0.170226764 | 0.174938376 | 0.25497338 | 0.312328512 | 0.385753117 |
| 70 | Unknown | HTCS | 3951 | 0.645752889 | -0.129889532 | 0.180998028 | -0.189156262 | 0.000202759 | -0.031577118 | 0.263265973 | -0.102938205 | 0.274537035 | -0.101658313 |
| | | | 3952 | 3.919424015 | -0.760479224 | 1.521925112 | -0.700930252 | 0.145469171 | -0.319442367 | 0.318170032 | -0.204915619 | 1.645284626 | -0.556654853 |
| | | | 3953 | 0.962559003 | -0.455140215 | 0.093402382 | -0.274447042 | 0.000202759 | -0.019102803 | 0.007260769 | -0.011775393 | 0.289043232 | -0.308398681 |
| | | | 3954 | 0.220945156 | -0.199396327 | 0.00230855 | 0.027570786 | 0.000202759 | 0.116606682 | 0.381640113 | 0.381848233 | 0.021069188 | -0.047325106 |
| | | | 3955 | 0.171696775 | 0.132562873 | 0.029046553 | 0.139420596 | 0.470149552 | 0.677852223 | 0.471265545 | 0.410669852 | 0.289043232 | 0.321915815 |
| | | | 3956 | 0.303032765 | 0.144250795 | 0.029046553 | 0.103399452 | 0.000202759 | 0.068188352 | 0.55857929 | 0.30801735 | 0.199805352 | -0.172431124 |
| 71 | Unknown yeast Glycan | HTCS | 3957 | 0.289700139 | -0.072699854 | 0.021866066 | -0.04831618 | 0.000202759 | 0.069019941 | 0.167376431 | 0.083480787 | 0.362527683 | 0.136992555 |
| | | | 3958 | 0.96566385 | 0.39944887 | 0.002406745 | -0.027933565 | 0.064661442 | 0.36197114 | 0.038713413 | -0.06331292 | 2.075222027 | -1.568595508 |
| | | | 3959 | 0.83884563 | 0.463452452 | 0.039729294 | 0.189683755 | 0.000202759 | 0.077819798 | 0.019975729 | -0.047673771 | 1.779052495 | -1.32509258 |
| | | | 3960 | 1.083182034 | 0.815253537 | 0.082777732 | 0.446057398 | 0.024985102 | 0.555662078 | 0.08310815 | 0.224874897 | 0.253228355 | -0.685294737 |
| | | | 3961 | 1.007023492 | 0.769257596 | 0.069868405 | 0.395574514 | 0.047316364 | 0.615679081 | 0.024114989 | 0.071201366 | 0.457408727 | -0.820673748 |
| | | | 3962 | 0.065227416 | -0.038770328 | 0.062212259 | -0.175582033 | 0.000202759 | -0.096999195 | 0.046395686 | -0.070492996 | 2.521857473 | -0.81746608 |
| | | | 3963 | 0.24903793 | -0.106674072 | 0.02416949 | -0.075299179 | 0.000202759 | -0.02418534 | 0.048814202 | -0.05923582 | 1.540999278 | -0.536423419 |
| | | | 3964 | 0.153244523 | -0.120801546 | 0.017461642 | -0.091664788 | 0.000202759 | 0.037397732 | 0.190850301 | -0.233984666 | 0.645048083 | -0.553305885 |
| | | | 3965 | 0.063419532 | -0.03658116 | 0.036987217 | 0.10874444 | 0.024985102 | 0.211835249 | 0.100936037 | 0.088377478 | 1.159761616 | -0.459199013 |
| | | | 3983 | 2.073367692 | -0.464646762 | 0.754458894 | -0.538630371 | 0.164826789 | -0.341629845 | 3.397031113 | -0.779960215 | 2.307489988 | -0.770923092 |
| | | | 3984 | 0.099047306 | -0.056837448 | 0.195073704 | -0.334514006 | 0.084278937 | -0.339771784 | 0.504328119 | -0.4281728 | 1.922887804 | -0.770083057 |
| | | | 3985 | 0.308642849 | -0.209511702 | 0.057429347 | -0.234951168 | 0.000202759 | -0.092670173 | 0.867544214 | -0.622631691 | 0.427342255 | -0.447133364 |
| | | | 3986 | 0.083527049 | -0.048712115 | 0.057501259 | -0.180667604 | 0.000202759 | -0.120015436 | 0.548953981 | -0.378663885 | 0.772197657 | -0.451595566 |

| | | | | | | | | | | | | | |
|---|---|---|---|---|---|---|---|---|---|---|---|---|---|
| 72 | High mannose N-glycan | ECF | 3987 | 0.342596258 | 0.200500939 | 0.004534747 | 0.036815119 | 0.000202759 | 0.112982132 | 0.602765601 | -0.468242106 | 0.427069436 | -0.384581819 |
| | | | 3988 | 0.778150074 | 0.320928246 | 0.056277223 | -0.17527325 | 0.000202759 | -0.102968773 | 0.308316603 | -0.263599983 | 0.087391607 | -0.116674804 |
| | | | 3989 | 0.072598264 | -0.035257591 | 0.057501259 | 0.180073076 | 0.024985102 | 0.243245274 | 0.903837373 | 0.342615905 | 0.468761955 | 0.279822108 |
| | | | 3990 | 1.070942011 | -0.222146021 | 0.297362226 | -0.237975805 | 0.781562372 | -0.365106095 | 2.205776863 | -0.281505073 | 1.271616135 | -0.275983087 |
| | | | 3991 | 0.117697305 | -0.031637321 | 0.115372276 | -0.143064656 | 0.183105341 | -0.2475441 | 1.128833144 | -0.211821986 | 0.389758712 | -0.123090246 |
| | | | 3992 | 1.132084053 | 0.248392815 | 0.00230855 | 0.014049537 | 0.003566779 | 0.119184428 | 0.417395854 | 0.179159076 | 0.140910441 | 0.093719962 |
| | | | 3993 | 0.937474414 | -0.212955818 | 0.245785282 | -0.251956707 | 0.000202759 | -0.097515602 | 0.757251067 | -0.243632049 | 0.588984702 | -0.211146701 |
| | | | 3994 | 0.135136462 | -0.035096813 | 0.129348331 | -0.160447733 | 0.144280548 | -0.209569039 | 1.197998802 | -0.216084987 | 1.919492528 | -0.323572771 |
| 73 | Unknown | Unknown | 4038 | 1.064504902 | -1.027828782 | 0.124866445 | 1.00928061 | 0.085269724 | -0.943012541 | 0.005254972 | 0.039267417 | 0.675434979 | 1.81917945 |
| | | | 4039 | 1.691405539 | -1.249947248 | 0.088475439 | 0.814147477 | 0.16705161 | -1.108245401 | 0 | -0.079095638 | 0.589084403 | 1.57721035 |
| | | | 4040 | 1.425799865 | -1.133550641 | 0.105796805 | 0.912855227 | 0.16571248 | -1.075974397 | 0 | -0.137495726 | 0.72101945 | 1.689945955 |
| 74a | Unknown | ECF & SusR | 4080 | 0.014711147 | 0.013916632 | 0.036192153 | -0.17286843 | 0.004851778 | -0.255776348 | 0.469335062 | -0.635538299 | 0.179957537 | 0.247772149 |
| | | | 4081 | 0.388623742 | -0.193935918 | 0.133764089 | -0.310459591 | 0.000202759 | -0.064955998 | 0.159553902 | -0.453273037 | 0.024025283 | -0.038435281 |
| | | | 4082 | 0.026751621 | -0.021215751 | 0.067049149 | -0.243662888 | 0.004851778 | -0.228410874 | 0.395303295 | -0.454997436 | 0.004712492 | -0.006662754 |
| | | | 4083 | 0.15909255 | 0.147144835 | 0.057501259 | 0.302990336 | 0.000202759 | 0.105681781 | 0.001085528 | 0.002771682 | 0.487381977 | 0.593474153 |
| | | | 4084 | 0.664283349 | 0.471009729 | 0.010036094 | -0.102647403 | 0.035394591 | 0.459576776 | 0.056241538 | 0.168354618 | 0.257482896 | 0.413148318 |
| 74b | Unknown | ECF & SusR | 4085 | 0.811507649 | 0.61949718 | 1.95089E-05 | -0.00302727 | 0.000202759 | 0.005288289 | 0.018151806 | -0.063811671 | 0.008676452 | 0.03065701 |
| | | | 4086 | 0.229538027 | -2.150643722 | 0 | -1.301215289 | 0.000202759 | -0.436883889 | 0 | -1.339469324 | 0 | -0.884350687 |
| | | | 4087 | 0.321042152 | 0.359015001 | 0 | -0.642091681 | 0.040729453 | -0.770098228 | 0.219195752 | -0.53367363 | 0.132873726 | -0.441206298 |
| | | | 4088 | 0.286745776 | 0.23250502 | 0.009717219 | 0.087388472 | 0.000202759 | 0.047298257 | 0.313393876 | -0.613591623 | 0.143108967 | -0.280933104 |
| | | | 4089 | 0.485218105 | 0.397778651 | 0.039756938 | 0.319655383 | 0.000202759 | 0.133059333 | 0.13690367 | -0.285512389 | 0.208435831 | 0.39818581 |
| | | | 4090 | 1.125950361 | 0.611521625 | 0.064917876 | 0.315202306 | 0.000202759 | 0.089135271 | 0.02584005 | -0.062213093 | 0.116399865 | 0.211510667 |
| | | | 4091 | 0.916427927 | -0.540530603 | 0.22460127 | -0.476083189 | 0.024479244 | -0.325283055 | 0.978752174 | -0.7011738 | 0.399601339 | -0.417704773 |
| | | | 4092 | 0.033502614 | -0.020139003 | 0.001622876 | -0.011094473 | 0.02746524 | 0.243593646 | 0.326024022 | 0.232696918 | 0.148768453 | 0.140665671 |
| | | | 4093 | 0.802446011 | -0.410526135 | 0.011969108 | -0.068023241 | 0.000525887 | -0.200493912 | 0.080739349 | 0.115454855 | 0.018666274 | -0.036360764 |
| | | | 4094 | 0.175123483 | 0.132303115 | 0.108217711 | 0.342059344 | 0.047316364 | 0.369758506 | 1.26928649 | 0.668158746 | 0.795753333 | 0.560584107 |
| | | | 4095 | 0.012006955 | -0.011209292 | 0.019048619 | 0.096510676 | 0.003466816 | 0.229235883 | 0.871823558 | 0.603904797 | 1.047129391 | 0.642420963 |
| | | | 4096 | 1.289266543 | 0.795135077 | 0.330973559 | 0.749134291 | 0.500622816 | 1.031793591 | 1.765061154 | 1.203675693 | 1.08698086 | 0.994527397 |
| 75a | Polygalacturonic acid | HTCS | 4108 | 0.200392094 | 0.285008526 | 0 | -0.206221645 | 0.001879041 | 0.431126257 | 0.021751441 | 0.080125848 | 0.031893531 | 0.12677861 |
| | | | 4109 | 0.40210191 | -0.627253655 | 0 | -0.479019526 | 0.000202759 | -0.080510491 | 0.008712114 | 0.033735244 | 0 | 0.003043231 |
| | | | 4110 | 0.761600422 | -0.467673904 | 0.00536243 | -0.055146869 | 0.000202759 | 0.201402947 | 0.146358683 | -0.212793325 | 0.245352221 | 0.28376731 |
| | | | 4111 | 0.696729488 | 0.187493933 | 0.078440958 | 0.154028421 | 0.000202759 | 0.073814306 | 0.444367274 | -0.211188435 | 0.45839809 | -0.218764758 |
| | | | 4112 | 1.40613125 | -0.427799697 | 1.134696094 | -0.759466269 | 0.145883736 | -0.401210745 | 1.8179486 | -0.619566011 | 1.247335391 | -0.522279383 |
| | | | 4113 | 0.755900786 | -0.270725126 | 0.538002238 | -0.496187214 | 0.001687904 | -0.166075871 | 0.343012128 | -0.261342675 | 1.772635739 | -0.645304855 |
| | | | 4114 | 3.233167191 | -0.532322038 | 1.358572068 | -0.546508528 | 0.123624592 | -0.256633489 | 3.292687923 | -0.540188542 | 3.920153346 | -0.697647257 |
| | | | 4115 | 0.476601697 | -0.211804197 | 0.047218354 | -0.137948959 | 0.123624592 | -0.366466388 | 0.371644327 | -0.255565723 | 0.767878156 | -0.453624126 |
| | | | 4116 | 0.093594634 | -0.17129872 | 0 | 0.268285637 | 0.006518177 | -0.645656928 | 0.748595288 | -1.543458388 | 0 | -0.846995016 |
| 75b | Polygalacturonic acid | HTCS | 4117 | 0.86331847 | -0.297521318 | 0.564955016 | -0.507075442 | 0.1208687 | -0.345806217 | 0.151576783 | -0.137841768 | 1.046924094 | -0.463850832 |
| | | | 4118 | 1.276276937 | -0.417800516 | 0.592395737 | -0.536668731 | 0.09041073 | -0.358753552 | 0.83323199 | -0.39199866 | 1.771112053 | -0.618147766 |
| | | | 4119 | 0.078068889 | 0.312796053 | 0 | 0.586616481 | 0.000202759 | 0.393340564 | 0.315031421 | 1.37833298 | 0 | 1.586044411 |
| | | | 4120 | 0.191583931 | -0.344564945 | 0 | -0.045880136 | 0.000202759 | -0.353013992 | 0.016581476 | 0.069642501 | 0 | 0.061358232 |
| | | | 4121 | 0.303125277 | -0.278351713 | 1.95089E-05 | 0.000539391 | 0.000202759 | 0.025063164 | 0.041150305 | -0.097038072 | 0.201442297 | -0.358634736 |
| | | | 4122 | 0.00788269 | 0.010734948 | 0.002406745 | 0.045062719 | 0.000202759 | -0.04334619 | 0.138509665 | -0.322025832 | 0.131260899 | 0.275642377 |
| | | | 4123 | 0.202162872 | 0.401484444 | 0 | 0.257712591 | 0.000202759 | 0.240123871 | 0.34759364 | 0.820989986 | 0 | 0.363104109 |
| | | | 4124 | 1.569204761 | -0.475222289 | 0.790758095 | -0.594035615 | 0.15711585 | -0.399850549 | 0.107843231 | 0.097639863 | 0.143108967 | -0.14549175 |
| 76 | Uknown mucin & milk O-glycan | HTCS | 4132 | 0.552043273 | 1.096116692 | 0 | 0.84179951 | 0.122319465 | 1.426120345 | 0.470713645 | 1.244988065 | 0 | 1.414511491 |
| | | | 4133 | 0.373498737 | 0.841834167 | 0 | 0.833190485 | 0.270942869 | 1.858140427 | 0.188591954 | 0.816837708 | 0 | 1.7961992 |
| | | | 4134 | 0.005028648 | -0.008422807 | 0.02416949 | 0.208083508 | 0.000202759 | 0.241837225 | 0.039589372 | 0.133435136 | 0.208876433 | 0.499339069 |
| | | | 4135 | 0.172512599 | -0.173848066 | 0.015558764 | 0.099155072 | 0.000202759 | 0.043955177 | 0.038173492 | -0.098930713 | 0.036008543 | 0.091736714 |
| | | | 4136 | 0.428035162 | -0.60738303 | 0 | 0.05345202 | 0.004537328 | -0.489066193 | 0.04358834 | -0.168430166 | 0 | 0.364278639 |
| | | | 4137 | 1.048602204 | -0.226950532 | 0.530857432 | -0.353840098 | 0.410119657 | -0.359599913 | 0.810837901 | -0.246233027 | 0.010004078 | 0.007795481 |
| 77 | Pectic galactan | HTCS | 4145 | 0.141844702 | 0.048105913 | 0.424332294 | -0.332828719 | 0.000202759 | 0.006646741 | 0.041150305 | 0.036621438 | 0.135688876 | -0.092447515 |
| | | | 4146 | 0.293314886 | 0.120715343 | 0.022503699 | -0.071688589 | 0.000202759 | -0.07468593 | 0.043252873 | -0.046259453 | 0.215760137 | -0.182401682 |
| | | | 4147 | 0.500223682 | 0.349928682 | 0.055022197 | 0.244805482 | 0.017081821 | 0.32784711 | 0.338777704 | 0.40659209 | 0.005770189 | 0.012910472 |
| | | | 4148 | 0.08039125 | 0.127721074 | 0 | -0.168485213 | 0.000202759 | 0.106846331 | 0.127298302 | -0.376087906 | 0 | 0.023503834 |
| | | | 4149 | 0.311559226 | 0.204840661 | 0.06746313 | -0.245307349 | 0.000202759 | 0.171555327 | 0.085359576 | 0.123314996 | 0.040169348 | 0.077453687 |
| | | | 4150 | 0.784002547 | -0.368066416 | 0.390763326 | -0.516614145 | 0.027287597 | -0.276207659 | 0.144967034 | -0.178196504 | 0.12950405 | -0.147788706 |
| | | | 4151 | 0.362450194 | -0.133922709 | 0.21432977 | -0.317949658 | 0.000525887 | -0.110634356 | 0.679417847 | -0.286630078 | 0.607796423 | -0.278105584 |
| | | | 4152 | 0.976656739 | -0.252315114 | 0.850409817 | -0.489546125 | 0.001687904 | -0.144255858 | 1.526833183 | -0.457455911 | 2.184679186 | -0.497629332 |
| | | | 4153 | 0.092066196 | -0.078359642 | 0.129213774 | -0.417844585 | 0.004851778 | 0.239550302 | 0.40559111 | 0.404859134 | 0.341069718 | 0.353020307 |
| | | | 4154 | 0.247947468 | 0.330677406 | 0 | 0.68184348 | 0.170641083 | 0.912528152 | 0.718966992 | 0.912173378 | 0.305511249 | 0.608981523 |
| | | | 4155 | 0.112950187 | -0.042854376 | 0.022287946 | 0.058141044 | 0.040729453 | 0.206312835 | 0.12037287 | 0.079668518 | 0.03546796 | -0.032864965 |
| | | | 4156 | 2.810397969 | -0.466348504 | 0.909820752 | -0.453484042 | 0.057021257 | -0.220381618 | 0.381831871 | -0.209385156 | 0.575591545 | -0.234123313 |
| | | | 4157 | 0.030757547 | 0.019518374 | 0.00230855 | 0.021165935 | 0.004851778 | 0.205457254 | 0.154879137 | -0.150628168 | 0.174995883 | -0.21727445 |
| | | | 4158 | 0.080393652 | 0.106937745 | 0 | -0.095969703 | 0.000202759 | -0.02694028 | 0.0253957 | -0.072885722 | 0.027517133 | -0.101516274 |
| | | | 4159 | 0.556828701 | -0.240363662 | 0.763083891 | -0.638343076 | 0.017081821 | -0.219441296 | 0.793350553 | -0.426960388 | 0.533778411 | -0.347509233 |
| | | | 4160 | 0.275005841 | -0.16822771 | 0.156202505 | -0.305907662 | 0.000202759 | -0.035466881 | 0.16238594 | -0.166268088 | 0.021069188 | -0.033415776 |
| | | | 4161 | 0.073136953 | -0.059851887 | 0.021772152 | 0.10247806 | 0.000202759 | 0.056166282 | 0.20492201 | 0.239567352 | 0.444293423 | 0.393043241 |
| | | | 4162 | 0.032969307 | -0.018076642 | 0.305657373 | 0.365173668 | 0.000202759 | 0.108393443 | 0.146941016 | 0.114714453 | 0.151751566 | -0.14245168 |
| | | | 4163 | 2.391150826 | -0.435975489 | 1.457601586 | -0.620666809 | 0.506287619 | -0.380782525 | 3.832744712 | -0.565572636 | 4.995678626 | -0.711730435 |
| | | | 4164 | 3.192669991 | -0.574694177 | 1.626563296 | -0.646420983 | 0.008215329 | -0.181638979 | 1.959798361 | -0.47260328 | 1.42331712 | -0.444510391 |
| | | | 4165 | 2.028306298 | -0.689811864 | 0.518454807 | -0.631249825 | 0.001687904 | -0.187987669 | 0.664863191 | -0.441847676 | 0.385286055 | -0.339869197 |
| | | | 4166 | 0.398003476 | -0.185593337 | 0.136903401 | -0.282406088 | 0.000202759 | -0.04268872 | 0.030083468 | -0.045538647 | 0.293668358 | -0.244344614 |
| | | | 4167 | 0.783062552 | -0.248080759 | 0.124866445 | -0.255460532 | 0.000202759 | 0.035802845 | 0.356137358 | -0.213358454 | 0.425847307 | -0.246694524 |
| | | | 4168 | 0.909138658 | -0.244059556 | 0.009973103 | 0.041411428 | 0.000202759 | -0.014050358 | 0.08239121 | -0.06633028 | 0.695100779 | -0.276822649 |
| | | | 4169 | 0.192862943 | -0.087815081 | 0.007352477 | 0.047199489 | 0.000202759 | 0.04531082 | 0.055146266 | -0.057831183 | 0.23577071 | -0.178382722 |
| | | | 4170 | 0.013722628 | 0.012297999 | 0.056878987 | 0.217097151 | 0.000202759 | 0.093712851 | 0.100067012 | -0.15150372 | 0.102485566 | -0.161923589 |
| | | | 4171 | 0.622288212 | 0.44599827 | 0.156296213 | 0.400240908 | 0.000202759 | 0.172280872 | 0.155411899 | -0.25661076 | 0.163566245 | 0.248968588 |
| | | | 4172 | 0.191812221 | -0.083170983 | 0.295120544 | -0.351550419 | 0.000202759 | -0.100400674 | 0.277912369 | -0.169051454 | 0.490217046 | -0.246627233 |
| | | | 4173 | 0.459070526 | -0.156064274 | 0.07028867 | 0.160125876 | 0.024951395 | 0.195011551 | 0.100936037 | 0.076640433 | 0.105399547 | -0.099171324 |
| | | | 4174 | 0.330963456 | 0.132196895 | 0.053599877 | 0.140418859 | 0.000202759 | 0.02487479 | 0.082791484 | -0.080820414 | 0.383426441 | -0.250659261 |
| | | | 4175 | 0.425865398 | -0.185373635 | 0.145832241 | -0.298330412 | 0.000202759 | 0.06358931 | 0.378201309 | 0.242569942 | 0.055433963 | -0.068888078 |
| | | | 4176 | 0.348115374 | -0.227875771 | 0.295146693 | -0.534145097 | 0.000525887 | 0.193693956 | 0.318505886 | 0.288919255 | 0.009331348 | -0.015527682 |
| | | | 4177 | 0.206855312 | -0.467581477 | 0 | 0.495740134 | 0.02541983 | 0.781441002 | 0.111016255 | 0.45515107 | 0 | 0.647538253 |
| | | | 4178 | 0.482872203 | -0.118858484 | 0.396414188 | -0.311114739 | 0.000202759 | -0.067047885 | 0.435184807 | -0.155659634 | 0.201389847 | -0.098200961 |
| | | | 4179 | 0.859066334 | -0.35737139 | 0.121250097 | -0.320588484 | 0.000202759 | -0.051892963 | 0.024564735 | -0.03335604 | 0.260640408 | 0.249695187 |
| | | | 4180 | 0.553493404 | -0.229464994 | 0.258448928 | -0.353638299 | 0.000202759 | -0.093330026 | 0.001689286 | 0.00280534 | 0.036008543 | 0.047967584 |
| | | | 4181 | 0.012829743 | 0.006476647 | 0.131721647 | 0.210234959 | 0.000202759 | 0.089258184 | 0.008838497 | -0.009947025 | 2.092734055 | 0.511981177 |
| | | | 4182 | 0.285431059 | -0.086528334 | 0.145832241 | -0.200500539 | 0.000202759 | 0.010642022 | 0.464610984 | -0.190179906 | 0.748745194 | -0.250258566 |
| | | | 4183 | 0.347680051 | 0.355847077 | 0.15722954 | 0.548859097 | 0.506287619 | 0.931470734 | 0.280964037 | 0.425184758 | 1.059994994 | 0.92590612 |
| 78 | Unknown | ECF | 4241 | 0.603558478 | -0.191975843 | 0.142377104 | 0.215103886 | 0.372015876 | -0.437715539 | 1.172962701 | -0.341096602 | 0.133260664 | 0.099953526 |
| | | | 4242 | 0.73950829 | -0.27279589 | 0.059693545 | 0.14503981 | 0.360623717 | -0.523058422 | 1.250455674 | -0.401486834 | 0.329789542 | 0.195257015 |
| | | | 4243 | 0.474748947 | -0.158577221 | 0.131721647 | 0.188031899 | 0.858347237 | -0.561911801 | 2.267790935 | -0.483129589 | 0.24557737 | 0.145432729 |
| | | | 4244 | 0.546839315 | 0.813076967 | 1.004839639 | -1.018207954 | 0.024985102 | -0.404085146 | 0.060392205 | 0.27874386 | 0.065153495 | 0.316350003 |
| | | | 4245 | 0.469787883 | 0.759027088 | 1.034380434 | -1.09457336 | 0.035411607 | -0.458303974 | 0.04358834 | 0.20819123 | 0.062901523 | 0.329794448 |
| | | | 4246 | 0.374426087 | 0.634938046 | 1.977363481 | -1.338880778 | 0.101367857 | -0.583349064 | 0.0253957 | 0.114507167 | 0.036008543 | 0.200282731 |
| | | | 4247 | 0.297897987 | 0.520964247 | 3.926853756 | -1.632050116 | 0.12372202 | -0.898562385 | 0.036111348 | -0.165205972 | 0.027019588 | -0.151352267 |
| | | | 4248 | 1.707473487 | 1.14938479 | 0.004858317 | -0.070901395 | 0.000202759 | -0.042814747 | 1.015535117 | 1.167610273 | 0.94812125 | 1.035933162 |
| | | | 4249 | 0.408124392 | 1.171728955 | 0 | 0.619511778 | 0.000202759 | 0.249260349 | 0.981648791 | 2.406670101 | 0 | 2.109294472 |
| | | | 4250 | 0.227816331 | 0.080065614 | 0.096829484 | -0.163277535 | 0.000202759 | 0.023879554 | 0.141887534 | 0.11818361 | 0.247781557 | 0.157098948 |
| | | | 4266 | 0.238515759 | -0.577481337 | 0 | -0.200141902 | 0.000202759 | 0.1618951 | 0.433108492 | 1.347297795 | 0 | 1.216134356 |
| | | | 4267 | 0.437910581 | -0.482700993 | 0 | -1.164216277 | 0.174206705 | -1.030510393 | 0.219195752 | 0.595057372 | 0.024631114 | -0.085212036 |
| | | | 4268 | 0.129130927 | -0.220170792 | 0 | -0.441518544 | 0.004851778 | -0.638190109 | 0.216512389 | 0.715123335 | 0.39413038 | 0.747320387 |

| | | | | | | | | | | | | | |
|---|---|---|---|---|---|---|---|---|---|---|---|---|---|
| 79 | Unknown | Unknown | 4269 | 0.269000906 | -0.598893483 | 0 | 0.217848046 | 0.000202759 | -0.031827485 | 0.234697157 | 0.838380413 | 0 | 0.926434488 |
| | | | 4270 | 0.400440127 | -0.736018453 | 0 | -0.453984319 | 0.001687904 | -0.534928541 | 0.544674774 | 1.144453313 | 0 | 0.574746089 |
| | | | 4271 | 1.139787106 | 1.93071675 | 0 | 1.943964546 | 0.024985102 | 1.188334843 | 0 | 1.015034071 | 0 | 1.925276539 |
| | | | 4272 | 0.330997438 | -0.216647529 | 0.056699829 | -0.207212748 | 0.000525887 | -0.187958612 | 0.173567334 | 0.208618173 | 0.387528143 | -0.381042326 |
| 80 | Unknown | Unknown | 4294 | 0.214562806 | -0.33793643 | 0.004858317 | 0.097303916 | 0.06599851 | -0.834694749 | 0.082023162 | -0.300665219 | 0.149325501 | -0.438775264 |
| | | | 4295 | 0.360173541 | -0.452417808 | 0.009717219 | -0.117689862 | 0.049606671 | -0.738041295 | 0.131260699 | -0.390489136 | 0.16294715 | -0.481210745 |
| | | | 4296 | 0.409391152 | -0.52259906 | 0.029046553 | -0.291833165 | 0.057854626 | -0.803533585 | 0.184256983 | -0.462705046 | 0.245352221 | -0.601726957 |
| | | | 4297 | 0.616870974 | -0.733401815 | 0.057501259 | -0.455630492 | 0.153503746 | -1.019839471 | 0.381869837 | -0.669464033 | 0.484378751 | -0.839457622 |
| | | | 4298 | 0.770027291 | -0.829167315 | 0.17949869 | -0.732073114 | 0.370147349 | -1.148383306 | 0.832086142 | -0.858949404 | 0.829455302 | -0.961714542 |
| | | | 4299 | 0.798145878 | -0.829718025 | 0.15722954 | -0.647679172 | 0.337575026 | -1.102222229 | 0.989607465 | -0.92104561 | 0.766243626 | -0.927788659 |
| | | | 4300 | 0.396840319 | 0.119189309 | 0.02416949 | -0.067717595 | 0.000202759 | -0.047780109 | 0.239473889 | -0.142035644 | 0.11293464 | -0.091401575 |
| 81 | Unknown mucin O-glycan | ECF | 4355 | 0.244954048 | -0.134890164 | 0.126157731 | -0.247524657 | 0.012288344 | -0.262456236 | 0.038173492 | -0.047043719 | 0.113390923 | -0.120033842 |
| | | | 4356 | 0.101611145 | -0.175319292 | 0 | -0.557414314 | 0.000202759 | -0.212264265 | 0.008047175 | -0.029347797 | 0.214311032 | 0.509757476 |
| | | | 4357 | 0.875070766 | 0.359255725 | 0.022170845 | 0.091162438 | 0.000202759 | 0.023220506 | 0.100221098 | 0.134388786 | 0.03757404 | -0.063978546 |
| | | | 4358 | 0.562769237 | 0.400718235 | 0.022170845 | -0.130294774 | 0.000202759 | 0.112698546 | 0.096961565 | 0.193328344 | 0.129669379 | -0.276578477 |
| | | | 4359 | 1.212260278 | -0.37398797 | 0.840859298 | -0.646116589 | 0.047316364 | -0.363598846 | 1.221460107 | -0.548870009 | 1.223297167 | -0.536959034 |
| 82 | Unknown | ECF | 4402 | 0.03960509 | 0.022109051 | 0.118210922 | -0.235043751 | 0.636121945 | -0.827868775 | 0.254481596 | -0.214425834 | 1.71718405 | -0.674332988 |
| | | | 4403 | 0.603483439 | -0.296249286 | 0.082198137 | -0.229453453 | 0.384989329 | -0.699250244 | 0.046840248 | -0.072002378 | 0.088482468 | -0.102877916 |
| | | | 4404 | 1.158222825 | -0.420169168 | 2.083478675 | -0.890782771 | 1.195008095 | -0.805259467 | 0.814933739 | -0.314066956 | 0.094823289 | -0.093378733 |
| | | | 4405 | 0.387028311 | -0.231392528 | 0.006595511 | -0.060449622 | 0.000202759 | -0.070439678 | 0.36338463 | 0.3293637 | 0.591310542 | 0.461522557 |
| | | | 4406 | 0.020495305 | 0.016852271 | 0.142377104 | -0.376753592 | 0.006603479 | -0.294722212 | 0.032723141 | 0.055986423 | 0.11946286 | 0.163507248 |
| | | | 4407 | 0.204271395 | 0.119860764 | 0.008774443 | 0.058132504 | 0.022858763 | -0.355857181 | 0.016581476 | 0.02564215 | 0.681069085 | 0.434858824 |
| | | | 4408 | 0.139337628 | 0.108551208 | 0.033337935 | -0.155436587 | 0.000202759 | -0.105095286 | 0.07315814 | 0.11578356 | 3.182999514 | -1.321490043 |
| | | | 4409 | 0.090324479 | 0.038234683 | 0.02368151 | -0.067654841 | 0.000202759 | 0.094259018 | 0.077466615 | 0.063977688 | 0.968419557 | -0.369846914 |
| 83 | Unknown | Unknown | 4470 | 0.274315806 | -0.06011057 | 0.005565012 | -0.023917929 | 0.000202759 | 0.02985762 | 0.135325575 | -0.06122439 | 0.091930185 | -0.047889122 |
| | | | 4471 | 0.006185299 | 0.00216668 | 0.008774443 | 0.026803637 | 0.100476028 | 0.200232235 | 0.176287162 | 0.092153636 | 0.088167793 | 0.053429467 |
| | | | 4472 | 0.855158104 | -0.150565772 | 0.017536276 | 0.035848059 | 0.065590272 | -0.171252582 | 0.582255798 | -0.203965555 | 2.612533645 | -0.336275299 |
| | | | 4473 | 0.214878805 | -0.063411651 | 0.029883833 | -0.08733321 | 0.024951395 | -0.173881182 | 0.592120204 | -0.21995958 | 1.553380083 | -0.366928029 |
| 84 | Unknown | ECF | 4631 | 0.431238449 | 0.149432752 | 0.036987217 | -0.094726595 | 0.000202759 | -0.031485271 | 0.014601605 | 0.014454691 | 0.002676958 | -0.002086885 |
| | | | 4632 | 0.246861298 | -0.137110689 | 0.043712581 | -0.156862626 | 0.000202759 | -0.134363131 | 0.009272856 | -0.011801656 | 0.314543154 | -0.2439926 |
| | | | 4633 | 0.013124216 | 0.007094795 | 0.105796805 | -0.216894921 | 0.003466816 | -0.137531252 | 0.02584005 | -0.026980091 | 0.201569628 | -0.178874152 |
| | | | 4634 | 0.13262853 | -0.048004098 | 0.235627399 | -0.278025573 | 0.000202759 | -0.068597816 | 0.207378924 | -0.130209347 | 0.499790543 | -0.208872609 |
| | | | 4635 | 0.033322466 | -0.017983411 | 0.057501259 | 0.162384672 | 0.002964065 | 0.150526023 | 0.941195408 | 0.392237744 | 0.853592316 | 0.362539878 |
| | | | 4636 | 0.081991396 | 0.058296158 | 0.029046553 | 0.123984188 | 0.000202759 | 0.043598679 | 0.011726286 | 0.020834942 | 0.149803213 | 0.188985504 |
| 85 | Heparin Sulfate | HTCS | 4652 | 0.076418353 | -0.038402677 | 0.00230855 | 0.017495951 | 0.042165743 | -0.298278192 | 0.017575703 | 0.02253265 | 0.548403925 | -0.3100162 |
| | | | 4653 | 1.455059194 | -0.51118727 | 0.132716777 | -0.290907167 | 0.372015876 | -0.553530616 | 1.528410928 | -0.666543119 | 1.187589533 | -0.588017885 |
| | | | 4654 | 0.812481617 | -0.37829676 | 0.046998121 | -0.160927195 | 0.024985102 | -0.299332289 | 0.925268314 | -0.578108267 | 1.247773566 | -0.668632596 |
| | | | 4655 | 0.835640228 | -0.271174755 | 0.074888922 | 0.162132315 | 0.000202759 | -0.135763465 | 0.2029784 | -0.156893635 | 1.301499974 | -0.494747274 |
| | | | 4656 | 0.257700294 | -0.132376409 | 0.025030963 | 0.090475674 | 0.000202759 | -0.128321495 | 0.308316603 | 0.221153133 | 0.27362822 | -0.233743405 |
| | | | 4657 | 0.951372558 | -0.294095296 | 0.022460809 | -0.076050274 | 0.000202759 | -0.106236412 | 0.057191685 | 0.059094238 | 0.581606719 | -0.324624213 |
| | | | 4658 | 0.366020549 | -0.137386449 | 0.005185202 | -0.036653449 | 0.000525887 | -0.133360494 | 0.221651769 | 0.173310595 | 0.475012074 | -0.273801953 |
| | | | 4659 | 0.366619647 | -0.153673184 | 0.007352477 | -0.039870989 | 0.000202759 | -0.089418871 | 0.001919287 | 0.003412628 | 0.652789622 | -0.364552549 |
| | | | 4660 | 0.833258005 | -0.217482478 | 1.95089E-05 | -0.00162723 | 0.008215329 | -0.151920565 | 0.238778415 | 0.138580396 | 0.390632101 | -0.18216225 |
| | | | 4661 | 0.645200234 | -0.209477604 | 0.021866066 | 0.05900839 | 0.000202759 | -0.038029033 | 0.280964037 | 0.150575929 | 0.263933288 | -0.162939586 |
| | | | 4662 | 2.025844409 | -0.507103914 | 0.304437538 | -0.339284995 | 0.000202759 | 0.068859855 | 0.171088358 | -0.129423595 | 0.595541055 | -0.319271166 |
| | | | 4663 | 0.582933504 | 0.099360373 | 0.019048619 | 0.038511393 | 0.000202759 | 0.035552161 | 0.048343434 | -0.024172207 | 0.006776078 | -0.00405994 |
| 86 | Pectic galactan | HTCS | 4667 | 0.390723824 | 0.168361878 | 0.136903401 | 0.257641317 | 0.000202759 | -0.102725727 | 0.005872299 | -0.007496656 | 0.434374046 | -0.287707558 |
| | | | 4668 | 0.010639736 | 0.01319339 | 0.018394107 | -0.115413642 | 0.000202759 | -0.236498739 | 0.269039044 | -0.417209561 | 0.478166095 | -0.646166138 |
| | | | 4669 | 0.117837794 | 0.093953582 | 0.039756938 | 0.169008934 | 0.000202759 | -0.016131252 | 0.07314107 | 0.135979183 | 0.162835097 | -0.279644647 |
| | | | 4670 | 0.020766642 | 0.017828184 | 0.004182751 | 0.042430989 | 0.000202759 | -0.095464872 | 0.33141656 | -0.357893913 | 0.644126381 | -0.603123143 |
| | | | 4671 | 0.72600143 | -0.291370064 | 0.202251719 | -0.319577531 | 0.051134087 | -0.290372276 | 0.842464869 | -0.428068684 | 1.648084047 | -0.65043823 |
| | | | 4672 | 0.321856665 | -0.780244603 | 0 | 0.005343383 | 0.000202759 | -0.019091709 | 0.386455471 | 0.89436549 | 0 | 0.236886474 |
| | | | 4673 | 0.512443282 | -0.133903956 | 0.022460809 | -0.054615224 | 0.000202759 | 0.008093132 | 0.221651769 | -0.111034188 | 0.052245628 | -0.038100492 |
| | | | 4674 | 0.205681205 | 0.069556647 | 0.016333115 | 0.050513545 | 0.012335956 | 0.166566094 | 0.018859178 | 0.017031959 | 0.091936434 | -0.061126591 |
| | | | 4675 | 0.368578667 | -0.2097864 | 0.033490544 | 0.114030532 | 0.000202759 | 0.010717378 | 0.08691967 | -0.135789149 | 0.588984702 | -0.367089673 |
| 87 | Unknown | ECF | 4705 | 0.06433252 | 0.033791755 | 0.097677999 | -0.235701091 | 0.000202759 | -0.011137995 | 0.257415065 | -0.21786689 | 0.031631871 | -0.040055268 |
| | | | 4706 | 0.192858356 | -0.164152015 | 0.374514015 | -0.650339842 | 0.000202759 | -0.096397091 | 0.021537641 | 0.041061304 | 0.137179314 | 0.192262484 |
| | | | 4707 | 0.163695674 | -0.089970427 | 0.322879592 | -0.44798992 | 0.000202759 | -0.077514713 | 0.167277072 | -0.156924163 | 0.021069188 | 0.030041193 |
| | | | 4708 | 0.325103791 | -0.221871411 | 0.856913372 | -0.95661458 | 0.051341828 | -0.421599587 | 0.619428944 | -0.531397296 | 0.141442807 | -0.228411364 |
| | | | 4709 | 0.198516125 | -0.260268392 | 0.011969108 | -0.115264987 | 0.000525887 | 0.309850923 | 0.108635487 | -0.271348266 | 0.055433963 | 0.191955075 |
| | | | 4710 | 0.051297616 | -0.064328445 | 0.057501259 | -0.336037908 | 0.000202759 | 0.006484766 | 0.126490449 | 0.237272018 | 0.120051111 | 0.245740172 |
| | | | 4711 | 0.032328151 | 0.029466822 | 0.204732375 | -0.518158252 | 0.00129528 | -0.228152014 | 0.398265892 | -0.457116114 | 0.161893478 | -0.260173526 |
| | | | 4712 | 0.347065427 | 0.336780878 | 0.205763151 | -0.665108096 | 0.004851778 | -0.360214071 | 0.038029568 | -0.091035327 | 0.195235922 | 0.329335953 |
| | | | 4713 | 1.772204687 | -0.305099806 | 1.540120064 | -0.541836477 | 0.136665917 | -0.364661349 | 2.189262421 | -0.491529974 | 1.974933349 | -0.408891247 |
| | | | 4714 | 2.545213355 | -0.655413538 | 3.742242252 | -0.976819768 | 0.000202759 | -0.088557066 | 0.078589419 | -0.077041687 | 0.013043269 | 0.014737756 |
| 88 | ECF | 88 | 4722 | 0.6775221 | -0.336119648 | 0.272852563 | -0.439910612 | 0.020630121 | -0.258370225 | 0.274843534 | -0.295043004 | 0.332732982 | -0.334498785 |
| | | | 4723 | 0.530994726 | -0.260154297 | 0.173473728 | -0.40034817 | 0.035099601 | -0.30033094 | 1.347062771 | -0.65621321 | 0.997190117 | -0.561141091 |
| | | | 4724 | 1.155848039 | -0.410516971 | 0.860932411 | -0.605096099 | 0.034635382 | -0.243705919 | 0.958021732 | -0.410551271 | 1.730295285 | -0.613521206 |
| | | | 4725 | 0.580609818 | -0.328957479 | 0.053229012 | -0.201704595 | 0.000202759 | -0.171522416 | 0.440511159 | -0.43350813 | 0.695506687 | -0.55764094 |
| | | | 4726 | 0.106025512 | -0.082227931 | 1.95089E-05 | 0.000140644 | 0.000202759 | -0.028077832 | 0.017575703 | -0.034371194 | 0.196928973 | -0.26371712 |
| | | | 4727 | 0.231497714 | -0.198631206 | 0.005185202 | 0.055618314 | 0.000202759 | 0.203101527 | 0.030779173 | 0.065402406 | 0.055433963 | -0.118227326 |
| | | | 4728 | 0.260399998 | -0.201746261 | 0.081674782 | -0.296666683 | 0.000202759 | 0.089982938 | 0.376401931 | -0.426709515 | 0.039798179 | -0.087293246 |
