## Supplementary Table 2 for "A high-throughput microbial glycomics platform for prebiotic development"

| Supplementary Table 2. *Bt* PUL reporter plasmids | | | |
|---|---|---|---|
| **P-*PUL*** | **Promoter** | **Transcription Start Site (PMCID: PMC10994844)** | **Promoter sequence** |
| 1 | BT0028 | N/A | aggacgaaaatgtaactttgccataacaaaaaactttatgtctaaagatatagaaactttggataataacaaagcccgggcttgcgaagtctgggctttgcgcataaaaaagggtgtgccaacgttgaagtgcacctcaaaagttagataaaaaacttttggggtcatttcacatttgacacaatctctttcttttcaactttcttatacttttatgaccaaagagaaagaaaaagattatgtgcgtttgaatatgttaaaatatgacccgatttgtcacgtttaggtggtatttctttttgtcgaactaggaattttgtttttgcgacctacagattttccatttcctgttttcagactgcaaatttacgaataatccatatttttatttctatatatgtaataataacaaattaaagaatttaataattccttaatattcccgaaagaaatgtgaaataaaatatatggaattttgcagcgagaaaaaatagtaagt |
| 2 | BT0140 | TSS_3939, TSS_50 | cttccaatgatactgaagagaaaatcattgctgatgatttgatggcgatggaacggacacggttggcagccatgcccgaacagcgacggctgatctatacgttaaatcgttttgaggataaatcttctccggagatagcctccgaattagaattgagttgccgtacagtggaaaaccatctttttctcggacgccgtgatatgcgtgacttcttccgaaactgtatataaaattattcatataataaacaacctttggtgcttatggaaaaggaatcaatactgagtgagaggtatctttatgcctcaacaaattttagagaacaagaatttaatactatttttaaacctcaaatttaatgtt |
| 3 | BT0189 | TSS_98 | tctggagtccgtacttaaatccatcgaatttatcagtgatatcaattataagaaagaaggcaggaatatacttctttataaaaaatgagaattaaaaaatagatcaatgtttaacccgttaataaggacagaatgaaagcatcaccctcatagaaactccctaaaaagatagaagcactcctaaggaatgcttctatccgcaacccggacggatattcattcgcttcgacccaaatgaataacgtatcgcacgttgcttgtttttaataactttagtcaattactaaaacgcacaaattt |
| 4 | BT0206 | N/A | cgtgtggcacggctgcagctacttttacgaatccgtagttcatcttgtacctttttatatatagtgggtacaaaaataggaaatagttttggatttttgcccttttgactctgaatcctcaaaatgaggtgtcttatgtcgttttaatgactattgcataagcaactaataagaatgtcaaatggaaagtaatatggtcttattttgacggtttatatgggatatatctatttgtttattaatatttgttaattatgcaataaaaagacatttgaggaaatttggtataaaaaatgagactttttgtttttctatttaactattataattatatttgtagccaaattaaagaattttataggcaatttacacctaaacaaacgataagcacttattaatatctataattttagagagatggacatacactttattagtattgcagcgacacaagacatttatttagacatatggcgtaatattaaaaatcagccaagctaatttccctattctatatattgtatagactatttgatagtctgtcctggaacgatgataaaagtatacttttcaaagtcgctccaaggggagaagtatattactgtttcatgcgtgaagcgtaatatgctaagttttcgttttgtagataaatctgtaaaaataaatcaattcattaatttaaattcaagtatcaaag |
| 5 | BT0268 | N/A | gtctaagtcttctctttataggaaaatgaaattagcaacaggactgtcacccattgagtttattcgaaatataagattaaaacatggaagtcaattattgaaggataaatcaatatccgtagcagaggtagcttatgaatgcggattttctaatcctaaatattttgcaacctgttttaaagaagaatttggcgtaactccaaaggaatatcaaaaatcttgttaggataaaaataatatgaacttaaaatacacaagtacgcgtcttaatggaagatgcgtacttttccatatatcaatgatctatcccatttgaatgattcctgaacttatattgaacgatttttagacctgttatagttaatagcgattatggtccaattttggaagtttttgaatgattagagaacttctttctactggataactcgcacttttgtgacgcatttgatgcacaactaatacttatttggtctaaataactttataaatctaatagt |
| 6 | BT0317 | TSS_183 | ttcaatttcaataataacttttgtgtgttagagtccctggaatgctgagtgtctacagcagttccagggatttctttattatcttttcatttaatgccaaagaaggttcattctgttttgaatattgtaattttgttagccgatataattattctttttgtttaaagaaaaaaaactacttattttcttgttatttaaaataaattcatatttttgtaatcgtagtcagaaccaaactatgaaatattgacgtcctctctacacacagacaatttttattaaccttaaaagagcagtatt |
| 7 | BT0365 | N/A | tcaatgtgacaccaagcgcactgctgaataacgaagaataactgatataagggttgctattatgaaacaatgatagcagcccttcgtcttaaaatgagctcattttaagagacatccgtccaagattatactcaaagtgtaaaaaagacacttatataaccgtttatccacctcgaaaattcatttgtccacctttaatatactcatcggttgtatttttgcgacactacaacgagtagaattattgttgaaccattaatatcaatatctatgaagacaaggaaatggccgttactactt |
| 8 | BT0439 | N/A | ttcattatccggccagaagtttgatgaaaaacaattttatcaatcaattactgacttcgaatatgagtggacattatccaaagagcaccatcctataatttctggagagaacccgatattgttggccaagacattatctgagaaatatatgcaatatttttattaagtacatacctcaaaataattataaattataaaacaaaacagcctatgaatataataccgaacttataattaaggaaattatcaaatctaaagaagaaaaaaaatacagaattacttaactttaaattattaatgt |
| 9 | BT0454 | N/A | aaaataggaatttgcgtctgatgtacaaaaacggtgtctgaagctttcacaaagatagagcaccctaacaagagaataagggtaaataaataatgatttcttttcataaatggggggtattagttaatttaacgaaggcaaagataagattttctctgcatatatttaacagtcaaagcgacaaacttgcgtttttttaccaagttctcgtaggactagtcaacactctgaccaccgttcttgctcttgtgatcattgacagggtcggccgcaagcaactggtctattacggagtttcgggg |
| 10 | BT0483 | TSS_278, TSS_279 | gctatggctacggatacggttacggacaggaaaaaggtgctaaatcgtaatccggttctaagaataaacctgagaataaacgatagcccgtttcaatctgattgaccacagtatatcccccaagatatagtcagtcagatatagccgaccgcttgattttccccaatcaagcggtttgttgtattcatctacgttgacaatataatttttatatcccatatcaaagaaaagcgggtatccggacaaccagatgcccgtttttttgttatatttgtattatcccttaaaaaatgataatt |
| 11 | BT0753 | TSS_491 | tattgcccatcgtctggaacgtaaattccaggtgaagatcagcatagaaagcgaacgtctgaaggccgaacacttctccggttctttcgacagcaatcagaatatttatgatatattacacgaaatcaacgttgagaaacaatacacgtggaaagtcagtggtgacaccatctttattaccgataaaagaaagggggtaagataaaaggaaggagaaagaaaatggacacagccgatagatataccggcagtccgcagctacgataaataactatatgttaacctttaatagtgataaaa |
| 12 | BT0867 | N/A | ttgatgcttttttaaaagttatttcttcgtttaccctatcttttcctttataattctgataaagtaatatttcaggaggtatttgaaattaatatgtactcataatactcatcccaccaagttctccatgaacttataccgccttagagtttactaaattctatatttcacaacctaatttttaatcaaaaatcaacacacaatcaatgcctatggaaatattacagtattatgatttaccatccctatcgtttacaaacaaaaccggtaaactagttattattttaaaatcaaaatt |
| 13 | BT1030 | N/A | ggtggattacaatgatcttatccattcgtcctttatatgaatcggttaaagtggaagttaaaacgatgtgaagatgtggagaagaacgatgattggaatataaaatatttaaaagaaaattttcaacaataatggggaggggaattatgatatagtttaataaaactaagagaaaaaacttcagattctatttttttattctttttaagagattacgattcttttagttgtatctttttatttatgtgaatagatgtctgatgatgtaatatatcattttatataaattaaaatttaatct |
| 14a | BT1040 | N/A | tggtgtttcctttaaaaccaaaatgccgttccttttaaaccaacgactgctccgttcgtatattttaaagctctttcatagaattgaagatggcatttttatgcactaagcaattgttttatgcaaacacatcgcataatcttgcaataaattatacattttttttgctatttattagatttcatagactttgactacctggctattcttaaaaatagaacataaagaagtatgaaaacagctataaattcattactcccaatcaatcacgcttgccaacgcccaacccctaaaagtcaca |
| 14b | BT1042 | TSS_700, TSS_701 | tagtttttcagaactacttaatgctctatttatcaaatgattatgtatttatttttcaaaaaaatgcttagtctaaccattttttacgaaaaaaaatcttattttcgtttagtatggttgcagtatcaaatgtattactgatgaataagcgcttatgaaaaatcatttttagtgtaacctttaaatttataaatatgagaaacagagatacatagatcctttgtatgagtggcggttatatttattacccgcacatgtttgttaactacctaaaaactatttctaattaaaaagaataag |
| 15 | BT1119 | TSS_755 | acaaaaggagataaatgaccagatatataacagtagtattacattaaggtaaagaacaaccatggtatagtattattgcaatttaactatttaaagtttgtttataatcaaatcatttaaagatttttgtccttctatgatttttttgtgtttgataaatgtcgtaattaaataaaaaaatatcacgcataatgactttttctttttttctaaatttcttcaaatgacaaattattgttacctttgtaatgtaatgtagttaatggttgcgcttataaagtccttggaaaggatttatgtataaaagaggtagatgtttactctgttcttgtttggattatcagcatcactgatatgaaataacaaaaaaaatatacaatggctaagcggacggacaggcagggaaatagtacagattcccaatccccctctttataatacattgcgtataagatttattaactgtctggtctggggaatctacagagatttaacgaatcaag |
| 16 | BT1279 | N/A | agaccaggcaccactatctgaggtaatggatgtgatagtaaatgtagtaggtcaaatgaactataagataaaaggagaccgctgctacttaatcccaaagtaatgggtatcatcccaaaagaaaaacagtaaaaaaacaacaatataataacctttttaaccactaaagattcatgcaaaaaagtatccggaacggagccctaactctatcccggatatgaaatcaagattctcaatattagagttttactctattcaattaccgtgaaataatttagttaataacctaatactaagaaa |
| 17 | BT1440 | TSS_1022 | tcttaaaagcgcttggtttggccggtatcacagaatcagtttgcagactatcgctcataagggtatctctcctttcagcctgggcctgtaaggagagtgacagaaacagaaaaagagatagaacaatttgcttcatcctgtgatttttcgttgttatgattgatgctgttcgtttcacgaatgtactacctttcgttgatttatactaataatcgtctaaaaaaaatgtttcgtttcctaaacaaatgagatatgtattctgttagttagaacaagcaaataaatcctaagccacaaaatt |
| 18 | BT1551 | TSS_1121 | aaggggaaagtcgaaaggtggcacaaaggcagcctgctttttttcctgtgcatacccctccacgcaacaggccaataataaagcagcaatatatcttttcattatcttcttcactttttttactcctgcaaatgtacattattccaactagattacgaaatatccgggaagaatgatagcagatgaacctagttaataattccttttattattatattcgcaactaaactgtaatcattatattattaatattattacttttgcggctgtttttaataataattaaaagtaaaaggactta |
| 19 | BT1618 | N/A | catacacatcatcgtccctccgcatctggctaccaccgatgtatataccatcaagttcaaacaaaaagcccctatcaaggagatcatgaatatcgtgaccaaaaccataggaaacatagactataaagtcgaagacgagaacattctcttaatttattcacccctaaataagaaaggaggccgatagacagaaaaaagggagtcgatatttcgcggataccaactcccccaaatcaaaattacttcatttaaattagttgtgtacacttaaattagtattaatgattaactcacaaaagt |

| | | | |
|---|---|---|---|
| 20 | BT1632 | TSS_1190 | gggcggaagagtttttaagagagaatatagtatcatgagtatattcgggtaattagagttaatgaaaggcggagatattcggaaacggatattgccgcctttctttttttgtctcgaaagttgtctgataatgtctgttgacctcgaaaaatgcgataaaatggactattttagacatttttggaagaatcatagccaaaatcgaacatcttaaatcttagatataacctaaatttgtggcagtgaatgtcaggaaagacttcgctacttatataaaaggtaatcttaattaataaactt |
| 21 | BT1683 | N/A | gtactatgcagagagttacggacacgcagccagttgaccggaaaaaagtcgttctttcggaagaaacggtaaccgaacatggaattatccagattgcggaattcaataagaatgcaggtgcgtgatgcttcttgtgtgagatgctccagcatttcaccgaaaatttcttcctccttctctttacttgccggtagagaagtgtccagaaattcacctgcaccatataccacacattgtttgacaagagaggagggaagccactttttggtcttgcgtatggcggcaagcaggcgtgctacaggtctgccttcttctgttgccactatcagtagcggggtatatcccggagtcgattcatagatctgaaacaactccttggaatgaaaagtattggttcccggtaaatcaggaatgtcttttccgcgataatatgttgttagtttaagtggcatccaaatacaaatttacactttttattcttcattcaacaataaaaaagagaatttgttggcttagcctaaaatttgtcttttatcctttgcatatatttgaggaaaaaaaggcttggaaaagcgaatttttgtattattgtcgtcctttgtgagtacagaagtaaaatggaaacgtctatataaaagattatgctaattgtattttactaaacaaaggaaaattaatattaactttaaattaaaacga |
| 22 | BT1763 | TSS_1290, TSS_1291, TSS_1292 | tatcattcagttttctgttggttactttgagtgattatatgcttgttatatgaataatatcccttttttacaaggacttaaattactatacaaaggtagtattttctatgattgagagagaatataatcccgactttcttatgatggagaattattcttataaaccctataaaatagcgttctatgcaacatatcttacagcatttgaaacatttttttcagtcggttgacttaaatcaaccgtaatttgcatcatcgaaaagagaaaattcgtacttttaaattaataacattaaacta |
| 23 | BT1775 | TSS_1306 | gcttaattttcgcttaattgattattaaatatcccttaaatgtagaatttatgcaaataagtatgctgaagaggggataaaagttagctgtagttataaatattcctgttattccttattatataggaaacttatattatatgagaaatagggaatagtacaacttaaagcttatatcatggggagcgctttgaaaatctatcaggaataacgccattgtaatactaaaaatccacatcttatataatcggataagtcactaattggatatctcctaatttgtgttatgtttgtatagtatttgccttatttaccttttttagcctttctacttgaaatattcttcttttttgcaggaagtgaatactcatttagataaagaaatagggtgtggacagcatactacaggatataaaacttccccctttttcctgaaattaatacctaagattgcgaaaaattataagaagatagtattagttttaaaattaattagaaattaatcagagtttaatcgtattttataaaaaatcttaagcaggctcttgttactttgcatcagaactcaaaaataaccttaaatattaagtttgttcagtcgacgctttcttaaataatagataaaaagggtgttttcgaggtgtaacaaatagcaaatgtataatataaaaaagagaccgtgaaaaattatttgaaaatcacg |
| 24 | BT1876 | N/A | tggaacgtgaacttgaaaacgaaatatataaggctatcgataaattgccggacgaatgccgtcgggtttttgctaaaagtcgctttgaaggaaaatcatacgaagaaatttccggtgaactgggaatatctattaatacagttaagtatcatataaagaatgctttagcttctttacacgctcatttgagtaagtacctgatttctctattactatttttttttcaggtaaagcgaaaaaaaatcattttctgactacccttaattaaaaaacaactgtctcctttatagaaagcataaat |
| 25 | BT2032 | N/A | tttagcagcatttaaacattattacctaacagaaagaaagtgttaaaaccgctattttcataaaaaaaaacatcagatattcgttcgaaatggtatctttttatcattttatgccttaaatagccccaaaaacacttaacgcttcccgaaagggcaactccccaccgtaccaagttcttctaaaactttgcgcgtttcctatgtaggaatgactccgcaaactgttacgatcacacaaggcgttatcaaagtagtattaaaaagtgacgccaagtcactagacgaagtcgtagtaacggcc |
| 26 | BT2107 | N/A | tgtaatctatctaattttatgttcgtttcataaatttaaaaccatacacacaaagccgcattaacatttattctcaaaacaaccattcatatataaatttatattaaataaataaatctttctagttttcctgctttaaatatatatatttgcaatatgtatatacatatagacattatataatacatatatcataagcaattaatactatgtagattaaaatatacagcctttatatacaagaaatcacattaaatctaatataaaacccttatgacttactttttaaaatcacagagt |
| 27 | BT2170 | N/A | cgaccggcttccggaacgttgccgggaaatattcatccgtatccgtgaagaaaaacaaagttatgcccaagtagcagaagaattggatatcagcataaagacagttgatgcccagctacagaaagctgtctcccgactgaaagagatgatttcgtccagcggataaccggagatcacccgaaccgttcagcacttctttttttatattaacttttatttgcacaaaaggcgtagggagtttctgtcaatctccggtctttatcagcaaataaaccgattaatcactaaaaaagattgaatg |
| 28 | BT2197 | N/A | ttgcttataaatctattcgtctgacagttgttgaactttcgtctgacagttgttgaactttcgtctaacaactgtcagacgaaagttcaacaactgtcagacgattaaatttaagtataatgaatactaaatctgtgtacatcaagaattaaatcttagagtaaatccgaacagattagttcgtttcaaaacagttcttagcttaaacttattggattttacttcccttccttgatttttttttcattttcctttggtggtaaatgtaagttgctgtgtcttcttattaaaagcgcgcaat |
| 29 | BT2203 | TSS_1700, TSS_1701 | gctggcactggcctatggtatgcatgcccttaaccatacaggttcgtcacaagaagcaattctgcaatattccctgtttgtgcgtgagtacaagcaaatgatgaatgaagaatattctacttcttatgcagagctgatgagcaaaaatcctccttttcatcgataaaaacatatcggttaatgatatttattcgcttattaatccgttattaacccatcactaataagttgtttaaccttgctacattatgtttgcagcgatgttatttacttattgttttatttaaaaacctaatacgt |
| 30 | BT2268 | TSS_1759 | caaaaagaaacgtttgcagggtagttttatatccaaaaacaagcaaatagaatgttaattactacactttgcattcacaaagtgacatttcatgcttgataaatcacaatatttatagcaaataaagatgtttatcagcatattttcgatgttataaataacaatatgttatatatacaaccttttttttatcaaattaattgttagtttcatatttattaataattttgcaacttgttagagtatagagacaattaatgtaaagttgaactttaagagagaatttatgaaaagaaaatta |
| 31 | BT2362 | N/A | aaagatagtaaaaatcaacccattccgattcaaagtcaactttattttttttcaatccacaacgctctatgcctatttttgtccaatcagtctaaataaactgatattgatagtggcgccacatcaaatacatacactcagagattgaccgtcaattcacggtacaatcgcagttttaaccaataaaatctaatagcctatggagtaaatcaaatcggtataatcggaacatagatttaaatcattcagagatttaactattcaaaaaccaattagtgaacatacaattaaacaaaaaac |
| 32 | BT2392 | N/A | gcaaatgataaaaaaagcaattttgaatgaaaagatatacatggatgaataaaagcatatgctcctgattccttgaaataccctaagtttgtatcctgaaaaccgaacggaaagtgtgcttcacaattatatatagccatttacacctttgaatctataatatttaaaacaattaatcacatgagagaaaaagccaaaagcccttaagtcaaattgaaatcaattagaatctattaaaccttatttaaatagaaagcctaaaagacctttttttcaaactaataaaatcttatagaaaaac |
| 33 | BT2462 | N/A | ttgcaatcattacctgaacaaacctcccgtatatttactctcagccgatttggaaataaaacaaatcgggaaattgcctgcgaattaaatatttcaatcaaagatgtggaatatcatatatccaaatcacttaaggctcttagaaaaacgctaaaagattacctgcctttattttatttcttttttaccatatgtaacaccgattactacgcacgttcctttatcttttgaaaaaaatattctttttcgtttagggtaaaccgaagatgaatcgttatataaatgaaaacgataaaaag |
| 34 | BT2529 | N/A | gaaacgccagacgaatcaggactttaccgaaaagcaatacgagcacatccacttcatgatgaagctgaccgatgatgccctcgcacaaatgattgtggtggtagagaaaccggaacaccaaagcattgatatcaacaagtctttcaacatcgaaaacgaaatcaataactaccgtaaccaactgaaaaatcagaatattctggatgtcaacaacaaggaatatgattatcagatgggagtctattatatggatattatcgccgaatgtgagaaactgggcgactatgtagtgaatgtggtagaagccagcagtgatgtaaaagagaagaaagcctcttaaataaagaggaactcaatcaaagaaatggatagccaatttatgcgttgcagcataaaattgactatccatttcttttttatttccacacagacaaaacaactcatgtccgtttacaaggacactcacactttacaaagacgttacatttttttacagacgttacgattacatttcttttccgtgaagttcccgttaagcttttactgcgcctattgcaggtctttttttttgcatatacatttgcgccaaaatacacaggggcctatgtttagtaccattttccaatgaccggacaaaggcattccaaaagaaaaccaaatctctaatttaaagcaaaagaaatgaaaattaaaaaagtattg |
| 35 | BT2561 | N/A | aagtgcaggagcatgagctaatccagtatgaatcccgttccatcacccgaagtctatcttcgcccaaagagcagacttttaaaacagaacgtatcattttatggtttctgaataaaagaaatattcccatcctgaaaaaagacagagaagcattttgggaaaagttgtctcctgagattcatgaattatataataataaatatgaaagtcagcttttacgcctctttgactttacggcgtggatggaatccaaaattcggaaagaaaaactgtctgaagtcctcagagcacgtgctagcgcaaaagaatgttaaagagtaaggtgtttgattcgtatttcagacgataaaaacactcttttcaaatctccattttctctttgtaattatataaattcgcttcacattacctctatattctctcctctatacaaggaaaatccaatataagatatcatttttgagattatataaaccaattattaatgtagaaaactatgcatgattattgctttcatttgcatagtcttaatttttaatact |
| 36 | BT2627 | TSS_2064 | agatagatattgggagtgcatatgtttgtttttacaagatgataaagtgtattatagcgacaatatagtaaatatttgtgccaagagaaagagactgattaaatattctgagggaatggaagtttgaaaacttacatttgcatataattgacttgtgatttttattattagaatagttatatttgttatacttttaaattaaaatgcttatgaatattatagaaagacatccaccatgattaatgatatccgctgtttaatcatataatagaaactaatcaaatataaagaatatagaa |
| 37 | BT2803 | TSS_2203 | tatgaaagtcagcttttacgcctctttgactttacggcgtggatggaatccaaaattcggaaagaaaaactgtctgaagtcctcagagcacgtgctagcgcaaaagaatgttaaagagtaaggtgtttgattcgtatttcagacgataaaaacactcttttcaaatctccattttctctttgtaattatataaattcgcttcacattacctctatattctctcctctatacaaggaaaatccaatataagatatcatttttgagattatataaaccaattattaatgtagaaaactatgc |
| 38 | BT2818 | TSS_2222 | tgttccttacaaagcctccttttccattccgtctataatttgtatcaaggtctacccatttgttacagatagtgtttctagctatttgtgcacctaatttggtgaatattgcacgtccatttttcacatcacccctaattttgcatcatcattaaaagataaatgaagttaaccttattgcacataagcaaaggatatatttagaaactgaagagagaataaagagaataaaagtcacccccaacccggttgacctaactataaactaaatatattattaacctatcaaattaaacagt |
| 39 | BT2859 | N/A | actacctcaaaacgcagatcgtatccttgctgtataaccgtcagaaagaacgcgaagcattctccaagcgtccgttcttcccgacgaataagatgcagatgaataaggtggatgaagagtttatgaataaagtgattcgtaccatcgaggaaaatatcatagataccaatctgaacgtagagcatcttgccgagattctgggcatgagtcgttccagtctgctgcggaaaataaagatgctttccaatctgtctcctgtggacttcatccgtttaatccgattgaggaaggctgccgaactgattcatgaaggtaaatacctgattggagatatctgctttatggtaggtatcaactcaccgtcctatttcagcaagcttttcctgaaacagtttggcatgacacctaaagatttcgagaaacaaagtcaggcagacaaagaaaagatagacatgccttcgtgaccgtccgggtgcactataacaaatgatgcacaaaaatgaaacatctgcggaaaatatgcttcagatgtagtctggtgctgcaaaaatggtattttttgcagtgatttttattataccgcctatatatatactcctacttttgccatcgttccggataaggaaagataaccccttttcggtgaacggattgatttatccttaaatgtgaactctaatactaatttaaaattctgatac |
| 40 | BT2896 | TSS_2279 | gttctaacttttaccgcaaactgaaaggagtattggatatgagtccgaatgagtatcttcgtttgatacgtttaaagaaagctgcccagcttctgaaagatggaagatatgggattgtagaaatttcttatatggtaggattcaattcaccttcttatttctcaaactgttttaagaaacaatttggagtgttaccaaaagatttcatgcaatgaacaaattcttataaaattgctctaaaactgattctgactatctctgcactgtttttttggatagaatttaataatatcaggttaatttcttatatccttttcaggtaggttccgataactttgcgaatgtctcgaaggtaggaacttattttctgaccagtctatttggttgggaaaaaactttcttctagatcaattaatatagaagtaacggataaataccagattagaaactggctttataagctattaattattattaatcaatatctaatttttcgcgcgaatatgatttgtatgtattgtttggtagatagggagaagagttgatacataaataaggaattggcaggtattagtttttttataaaactcttcttccacttttcttcgctgaagtatataatccttaaataatacgatttatgaacaattccttttataaggatcatgatataaatgtttaaccaaaataataaagaaa |

| | | | |
|---|---|---|---|
| 41 | BT2909 | N/A | ccgggcgaatggcacacctattatcctaattatcaaacaggttggagtcattattggatcggcttttgcggtccggaagttgatagctggatgacgaatgagtattgttcgaaagaaagtcctgtatttaaggtcggtatcaatgatgagatcgtttctttattcagaaaagctattgatgttgctaacgaagaacctactttatatcaacgcgtattgagtggattggtgacctatctggtagcactaatgtgtagtatcgataagaaccttcaggtagagaatgatgatttctcttctaaaatagattatgcttgtgtgctcatgaaggaattgatagatcaaccagtttctatgcaagagattgccaaaaaagcaggtatgggatattctctttttcgaaagttgtttaaggaacagaagaattatgcacctgtgcaatatttccagaatctgaaaatacagaaagccatagagttgttgactactactactattccggtaaaagagatagcatataggctcgattttgaatccccggcttatttctctgcccggttcaagaaacaaacaggaaaatcacctatagaatatcgagaagaatttcgtatcaaataatgatagtgttctggcaattatcgaaagtctttctcttttaagatgactaagtttgtatcataattaaaaaatgaataatgct |
| 42 | BT2922 | TSS_2293 | gtacatgcaggatctatatccccgttgcattgcgcatatatacacaatgtacagggcatatagatcctgcatattgcatatatatagcttatctgcctctttttttattgcaatattttatgtgcttttatgtggtatatgaaacaattatctcattcctgtttcatttggtatagatttgctattctttcttctatttgagacattattgcgatttattgggatatttatattacccaaaaactatctcatccgctaattttgtctgcgtcaatcagtcgaaacaataatctaattatat |
| 43 | BT2952 | TSS_2321, TSS_2322 | cttcggaattgctaacacctacgggactaaagttacttacaggtacagtaattattcttgccggaatcattttctttgcgatagcagggcacaacagagatagcaaaagccgccagacgcactctgaaacacctattaagaaaaatttcaggaaagggttaattatctgtttgcttgcaggaatattcggacctatgattaactttgctttcgtttatggggcacccttacaagaaaaggctgtagcaacgggagcatcttctctttatgcagccaacgtgatatggagtatcgctttgagtgccggatttattattaatctccttgaatgtatccgtttgttcggcaagaatcaatcttggaaaacttaccgacatcgtacaaccggaggcctgataatggcttctctggcaggggtattatggtatctgagcattatgttctatggtatgggaggcagttttatgggagttttaggagcttctgtgggctgggctaccatgcaatcgacagctattattgcaggtaatgtggccggacttgcttccggcgagtggaaaggagctacccgctatgccatccggatgatggtgataggattggtatgtctcatcggaggagtggtggtgattgccctctaacgactatacttaactatttatattgacttaattattatgttattcaaaaagtctaactt |
| 44 | BT2970 | N/A | gatgtgcactatgtcttggctgatgtaaccttggaacaaggtaagaatacaattaaaaccgttgcaacatataatggaaaagaatacactgacgagattgaatggaactataccggcgaaaagaagcgaagtgccgattggagtgaaaacaaagaagagcacgcaggttggtaatcggagacgtttctgtatgtaatctgaaaaatgttgaatctaaaatataaaagcctatgaatgaaatgagaaagaatctcttttaaatggaaaacaagtatttatcattaactaatctaataattt |
| 45 | BT3011 | TSS_2381 | atcaacgagtatgactggcgctcaagtaaatatcgctttaccaacaagccattacaggaattcatcgattttatcaaccgttcataccatgtaaacatcataattaaagaagaaaagctcaaagagctgaagtttaacggaactatacgtaaagacgagccgcttaccaatatcatcgaaaagatatgtatcagcctcgatttaaaagaaaaacaagaaaataacaacatcatattatattaatacataagaatcaattatcaatatattcactattaaccttttagaaacatgaaaaac |
| 46 | BT3024 | TSS_2400 | aaaaacaatgcaagaaatggaaactgctttcatcatatccggcaagaatattaagaaaggactccggtttgacgatgtaagcatgatgcaatacgatgtggcgtccactatcgcccgcatctttcatttagaacaaccacaagtttggataggtagacccatggagattgttttcaaataatcattcaggatgattaataggatgggaaaaggagaagctaaaataactggcttctcctttttttatttatctatccacgctcataagataggagattataaatactatttcttataagtaacaaaatgagcagtcataatagttttatttcctataagtaatagaaatcacacctccacagctgttaattaccgaataatacttgcatcaacatacatatccttataccttttgtcccccggtaacacaacataatataacttcaatcataaaatgtactcttttgacattcttaataagcaatccaactgattaaagcagcattcaaacattctgataggacatcttatcttttgtttattcaaattaaaatacaatacgatagaattgtattagtaaatgataaaatactacttgtctgacggagcattatcgcctacttttgcatcatcagttattggttgaagtttccatggatcaatcaacaattgaatagagtttataatctaatacaataaatt |
| 47 | BT3047 | N/A | agttgtgtttcttgcagccaacgtgtgtaaatcgtttatgaatgattctctacgtttttttgaatgatttttataatgtaaaatctacactcattcgtatctttgctgtagataacaacgaaataaactaatattatgaaaatgaaaactaatataatctttaaaaaactaaagtcattcgtcaatcgatgtctttttcctgattatatgtgctgttttatcttcctgcgtttataacgctgatatgccaataaagaatttttagtgaaccaataatttaatatgtaaaaagaaaaacgtt |
| 48 | BT3090 | N/A | gctactctgtgaccactatttataattaccggacacgtgtccgcaacaaggcactcggcgaacgggatgagtttgaggctaaagtaatgaaaatagggaaggtggaagagtaagaaatccactactttttttagcacggtcgctattctataatattcttattattagtgttttatgctgttttgtactgtataaaccactatattttttgctatctggcaaaagaagcgcatttaattgtggatacttttgtttcagtgaaaacaaaaagtaatcctaatattaactttaaattgatgtac |
| 49 | BT3108 | TSS_2482 | taatgtagctggcagacatccggaactgcttgagcggttaaagcaggaattttttgtgcaaacagatggattttatcgtagtgaggtggaagaagaaccattaaaataagcccgataaccccttctggaaagcatttgattgattaggtattagattgttttgcaggaattggtgtagaaggatagtttctgaacatgtgtgctcataataaaacatattttctatcgtctacaccttttttcttttacttttgctgcatcacaatcatacataaataacgtttaactttttagaatcat |
| 50 | BT3156 | TSS_2524 | acaaaaaaacattcattcctaaaataaaggatgatggactaatttatatatccatgaataaaggagataatcttaccataatcatcccttaataacaattttgcagtaaatgcgttattttttgctgcgtttatacactattaaaataacacatttacctaattttgactcgttgaatttagaaaattatcacattaaatctaataatatatgaaacgaattcgagatttccccaccactgacaagacaacgatctagttgcctgaaatgtattaacctaatattaattttaaattatcc |
| 51 | BT3174 | TSS_2553 | ctgcaaaacgtcctgtttctaaaaaatgcgggtttttgcctaaaaaacagtttgtttgtcactgcacgggaagtgaatcaccatctattggcatttgtttttgatcacttctctatcaataatgatcatttagcggattgcacagggttcttttgatcattattgaattttaattgatgatttttgaatgtttccttttatgctgtggttgtatttttgtcgaagattgaaacttattgttagctagtgaaagtatgaatctttattattaaatgtagtattatgaaaaaaaggaaacag |
| 52a | BT3239 | TSS_2610 | aaggaagtgtttagatgacataatgattatttgaacagggctaaatttgcacatactgacataaatgtgacattatgatagtttgatgatatatatttatcttcataactatcataatgtatttacaatagttttcaaatttattttgaataattgttttgtatttataaatatgtgcttatatttgcaaaatagcactaagcgaaaataagcgacttactaattcttataataataaagagtgtttatgaaaagaaaattaatgctgttattggcctgcttgctggcaagcataggtctagtgattgcccaaacacctaaaaaggtgacaggtgtcgtcatttctgaggaagatgatcagcctgttgtgggagcatccgtattagtgaaaggtacaacg |
| 52b | BT3240 | TSS_2611 | cctgtcaccttttttaggtgtttgggcaatcactagacctatgcttgccagcaagcaggccaataacagcattaattttcttttcataaacactctttattattataagaattagtaagtcgcttattttcgcttagtgctattttgcaaatataagcacatatttataaatacaaaacaattattcaaaataaatttgaaaactattgtaaatacattatgatagttatgaagataaatatatatcatcaaactatcataatgtcacatttatgtcagtatgtgcaaatttagccctgttcaaataatcattatgtcatctaaacacttccttttttatctttaactaagccttgaataatataaaaattatattagatataaataattagtatgaaaataggttataggagaat |
| 53 | BT3270 | N/A | tccttgcttttgtgggggtgaagtgactgtgatgtggcttatttaagtggatgagccgttatgtttaacaggtaaacaaagcttggttagcatttttagtaagctatatgtgttcgttccatgtctgtgtcttattgcattgtatatcaattagctaagtccgaaagtgacggatgatgataaaatatgagtttcaaactccttagtaaaagctaaaaaatacatcattttaatagcggcaacgaaaaaaaacaaatattttttaagatgcctaattgtcatttattgatggctgaattttgattctgaaaagggaatgatttgttaaatatgtcatttttgttgctgaataatgtcataatgatagtttttcattcttttccgttttcgtacggagagtatctcataaaaaataatcatctttttgtcacccatttatggacgttatgtataattatagtaaagcaaagataaaatcctcattagaaatagataaaatt |
| 54 | BT3278 | N/A | tctcttgaaactgtgaagactcaaaagaagagagcaatgtccttccttagaaaaaaattaggctcctatcattttctattattacaaatgttacttccctaaaaagaagcataacactcaatccgctttacttgttaacacctgtttgcaatgaacaaaatgttcgttgtttttcttgattattgtacaaaaagaacggtcttctaaataaataaagtatcttttgaataaaaaaacttcagaatcctgtccctcatttttttcttcttagtcgtctttgctataaataacctgaaataat |
| 55 | BT3301 | TSS_2664 | ttctcttcgtccattcattgtgtttttttcatatagcattctcttgttttaaatcaagacacaaaaatcgacggaaagatatggaaaatcgggaaaacgtccaagaactgcacaaaaaacattctgatcatctgaaaattaacataacaggagtacgcttttacgacacatatatatatttttgcgacatttcatcgagaagagacatttatgtcatgggtccgtacatcatcttatactaaccataataactttgcagcatcagacaacggaataaggtaaaaagcaaaataaaaaggattgtagtttgaaataggtaatccacaaacaggtagcatattcagctacctgtattttggaaggaaaatctttagttaataatgaatataatattatataga |
| 56 | BT3310 | TSS_2675 | gaatacaatttataattatcgggcgaaagtaaaaaacaaagctcgtggttccagagaagatttcgaagatttagttcggaaaatacgttgatcgatgttgtaacctatccactttttaggcctctttggttgatattaacttgctgtttattagtggtttgtatgtttgtataatccacttattataccctcattgtcaaatcacttcagtttctttatacttttgacctgtgcttctgaaatagtggcgaaagtcctgtcaagcacataccttaattataaactttaaaattaatacgc |
| 57 | BT3332 | TSS_4409 | aaaatggaactgggcaatgacaggcatccccagctgtacgatttatccgccgatccatcggagaaaaacaatgtagcgaaacagcatccggaagttgtcagggagctgtccgaactgcttgaatcggtaaagacaagataaccaatgagaatagcagatttgtattattttgtatcaatcttgaataattctatccatatgagaacaatcaatgcgtatatctttgccatcgtaaattcaacaaaatcacttaatttttaatatatgaaaaaacatctatccaaccagacaagaaaaagg |
| 58 | BT3347 | TSS_2707 | acatttctcccttgaagggcatatttgcacagatattccccaaaagcctgcttaattatttaatgtaattgaaaatgtaattataataatctgactaatatataatatattctggtcatttcaaaaaaaaggcattctaactcttctacatttttattatatttatttgtaattaaccgattgattgttaattataattatcatcatcacacctacataaagttatgcaaactaaataagaaattacttatttagttagctttgccatgtcttttctcagtcaaactcttcgttgatgagaatcagacatatatagatatttgtacttaaatctacaacatattatgagcacaacatgccaacatcaacaaggtgaaggatacctaaaactcctaaagatttagatacatattatgcttataataagaaagataggcaaacgttgtaagcaataatatttattgttcggatgcactttaaccattttattatttatattt |
| 59 | BT3477 | TSS_2826 | aaagttagaagtcatcaattaatagaccttcattttgggttggtaaggataaaatctgctcaaaaaaggataaatactgctcaaacaacgtcagagcccctattccgtatgttaaaagaacgtttgaaatccaaaaaagaatagattttgtcctttcttgaacacagattttccttgtttcttttattaattccattattttgtgtcatcaagatgagatactatggaaaataaagagtttgttaaagctcgatatttgctagaaatacacgatttgttagccgcttctccataacatttcatcctaaatatgacagtactgatattatatattaaaggtataggaaaacgggtataaagattgcaattattcaaaggtgagatagtattaaggtaaaaagtggatttaaaagaatacttagggcttgtttttttatgaggtaaataagattgatggatggataaccttggatttgaagattaatttaaagcaaatattt |
| 60 | BT3518 | TSS_2856 | gaataaatgtcgaatttgactcagcgctaagtaagataaaatgttccggaaaaattgatttgaaagacaactttgagacagtaattaatggattaacctttgtagcaccaatatcttatgcctatgatggacagtataaaacttatcgggtcgtgaaaaaatagacacagaaaaatatagtaattaataaaataaatctcaagaatttatggcgatttttataaagagtggataatactgctgtattatccagatggggctcatatgcctgtaagtattaagttataaaataataatatt |

| | | | |
|---|---|---|---|
| 61 | BT3569 | TSS_2905, TSS_2906, BT_2907 | gtctgcctgatgtaaaagagtagttgcattacctccatcatgacaagcagagagagacaaaatcaggaacaaccctataatataagaaagaagtagcttcatagttcgtttcgttttacaaatgtaaatagaataaaacaaacaagcaacaaaaagacaaataatttttcacacatcataattacaatgttaaaattgccctttctctctctttccctaatctttttgcctttcatttgttcactttgttactttcgccggaagacggcagaataaacttcaacctaaatagtaacga |
| 62 | BT3607 | TSS_2937 | aatgtcgcgatgagcaacagagtcaatcatgacccaacccctgaatcaaacgaaattatatatgatttttttaaatggtacttgcattcagcaaataaatctaccaaagagtagaaagatgcgtttttgaatacactttttcgttttttgcgcaaaaaatcaactctggatgcctaactttgttctatcaattcatccaagtaacctaaattacttgactaaaatgattagcctatgaaaaatatacactagaaaacataaaatgccccatctgcgaatggggcaatattaatgccatattaataaaactaatcataaattatcgcccaattaataaataacacaataataaattcataattaataaattaaaagtgctagctcaaaggaattaataaagctagcagatgattccgagttagtaataaatattaaactattaaaaattcaataggttaaatatatactaaaaagagtagaaactaaaaaaagtaaact |
| 64 | BT3668 | N/A | atttttccagttccaatcggcattatgaatagccaaacctatgttcaaaacaagcgggtgcaatttatcaacaaatgaagccatagcatatagggtttattaatgtgaagataaagatatataaaaaataaatataactcttttgatgtcgataattttaaataatatatcgatattgttttataataaatcctgctatagtcgttcatgtcgatattgtattactctaaatcgtattagtcctacaatgtggactgccgagttgttacatttgcggtatataatctaataggaatattt |
| 65 | BT3679 | N/A | cgtaaagggaactatagtgcatcttgcgtaccactatagtggcacgaactaatatgactaacaaaaaacttaattgttcgtagctaaggacttaagaatgtgagcgataacctgcaaggagtgaactgatttcattaacggtttaccatataacaaattattgttttattgatgttatgttatatgtggtttttattttacagatcagattatccttattccttggtcgaattatagtttaaaaaggaggaatatctataattcgacctcagataactcatattggacctcatctggagtgacttaacttacatttgcacttgttaattggtcgttgaaatcataaatcaattaatgaacggttaatttttatttactaatatcttaaatatttttgct |
| 66 | BT3703 | TSS_2999 | tacttcctgcctcatctgctttcgatttattgacagagaaaaaggttgaattaagaaatgaattaacacttccaagtcgtgaaatatacctttttatcccttttaataagttcacttcaacttatatttatcagaatacagaaaagaaataaactactgtaaaacaaaacattaacattaatacaaacttgtatttttattctagacaacccaaagtctcaagatggttgctatatttgcaatatccggaacagagcggagtaaaaatattagttagcttataatttaataccataaataga |
| 67 | BT3749 | N/A | atcgtctggatggaaaatctcataaagaaatagctgaagaattgggtatcagtgtcaaaggagtggagtatcatataagcaaagctgtgaagctgctgcgtgataatctgaaagagtatgctcctttcctgatattttttatttgatgatcttgaaaatatctttttctcctctttttttaataacagcaaaaaaagttgtatttctcactagggattatttttagtgagtggtattctatatgaaagggggagcaaatatactcctgtttattacttataaagtaactcaacttataat |
| 68 | BT3787 | TSS_3070 | gatttttatcttgattttaatagcggggaatttaacctttaaggagaactggaatactattaaatcatgccaaaatagtaaatatccctgccgtataatataaaacacaaagaaggaaaactaaaaatcttggcaaacacattaattttgctggtgcattaaaaaggtcagaataatataatatgtttaactaaatatctcatgaaactatgaatcaacaacacgatcctccacctgtttttttaagtatgtaatgatccctttttacttttattttatttgaattaaatttaatatctt |
| 69 | BT3854 | TSS_3130, TSS_3131 | atcggtggcggagcctgtcctcatcggtgccggaaataggtctcatcggtacccgaaataaatccttgcggagtaaggcactcattataccatcaaataataattgtttttttttatcatgaaaacattattgataactagatacttatgcgaaaagaatgaaattctttatcgctattaatctaaaattaatatggcattaaggactggaattctatatttgcttcagaaacggacgggatatcctccactatgtttctgatgacttgagcggatcgaactataaccttataaaatgatggaaatatgaaaaacgacacttcttaactactgcttggacttacttgtcagagaaagttttcttgagatgaaaacggtttgaatcaattaatgttaatctataaattatgaaaacaggaaagagcaaagatcatttgcaaagagtttttttaaggttcctttatttgataatggggagcatcctttctttggattctgtt |
| 70 | BT3952 | N/A | atatggaagacctggaaggctgggcagattctattgtttgtaccactacggaacaggcggattttgtcaggaagttcttgcaggtcgtggagcagaatctggatcgtgaagatttgggctccacgtttgtggcagaacaaatgttgatgagttcccgtcagttctaccggaggtttaaggagatttcgggtatgtcacccagcgatttgattaaagattaccggatggagaaggccgcccgtttgttgcagaacgaggaactgtctattcaggatgtaatttccgatgtaggtatatccagtcgtgcttacttctacaaagagtttacccgtaagtttggcgtaactcctaaggtatatagagagaagcttttagggaatgacactcataaaaatcaggagtgaacagccaggaagcggtctggtattagttgccaagaaataaattcccatgctgatctgtgctgctttttaaggttggtctcctttgtactctcttttgttttccattatttattattctgatatatagggtattatccagtactctattttaccatctgtactttaaacagacatttttctcttctctgtttttttggcgcacaggagtatctcatttctttggctattcctttaatttgcgtcaacgtattaaacgtatttggtaaccttattattattaattaaaaacaagtagt |
| 71 | BT3958 | TSS_3220 | agaggaactatagtgcttcttgcgtccaactatagtggaacgaaagggaaactacagtagaacgaatgaatttggactgataagaaattagtttttcaaagtctttcggttaatgattcgggattccttcattccttctgttctttggtcgtttggggtgacagatatatttgaattcatgtctgtatagattttatttattactgcttacgaataaactttttaataaaagtggcgaaaaggcggctattataacttgatacaatgtgcgcaaacttgtgctctgtttgtacgtgaatgaatgctatgtgttgatatatagatgtttataatgtgctttatttctcctttctgttctataaatgagcattttcattttgacataattttgtcacttctgcggaatattcttaatattaaaagtttttaatttgcttccgcttataagaattatataggcgattgataaccttagtattaattaaaaaaaaacaaagtaac |
| 72 | BT3983 | TSS_3253 | atatttgactccggattattttcgctggttgtatacggcatttactcgcgccacgaagacattgtatctggtgaattatcccaaagaacaaattttataaaagatttcttttttttcttcgtaaataattgattcccagtctagatgaatggggagatactcgttgtgaaaaaataaatatgagattatatttattttttttcaaaaacctctaagtgtttttcaaggtatgggttgttatataaatgtgaatcgaaaataataaatggaatatcgattctaatatttatataaaaacta |
| 73 | BT4040 | N/A | agaatataacaacctttatcacgctgtgtactatccaaatagcaattaaatagtttctcccttacacccgtaattaaacgactgccaaaatgctttatatttaagaagtgagtttttcgcccggtctgaccgaaacaatcaaccaattaaaaaggaatagaggaataattaaactaagttcttgtagaacttagcacaaaaaacagtctgttaatccatttgtatctaaactaacgcatctaactaagcaatagatttaccatctttttgctaaaaatcaacgattcgtgcgcaaatatactaattagtttgaaaaagtcatactattttcaataatatttcatatttaaattacaaataagtttttcaattattctcatttaatcaatcaacgggtagaaacagagtgagtatttcactattacacgtttacagcccccaatacttcccattgattatcaacaacatgcattctattccatcttatcatctactatattatagttctacaagaactaagacatagaacctattattttttctgactttaaatcagcatagttcagcacacaaataaaagttttaacttctaattttaatgcctatgagaaagaaataagtattccaaatccaagagtacccacaagtacttcaacttgtttttaccctttttacctttaataaaactaatattaattt |
| 74a | BT4080 | N/A | ctattatcagaagatggagtgataaaagggcaatatagtactatgggagatacttatgttaattggacacgtgatgacaccataacgggagatgtatatcaaaagagttattttgatcgtatgattgaaagaaatcaagtaaatgcttatctgaaagaaaaggaagaagctgttcgtatttggaaggaaatagaactgatatctcaaaaactacattttccctcagaagaattgaatcattttattcgtatttcctgttcgtatggacgcattaaatatgagctttttgctgtttcgtggcagattatgctatgcggttatgtggcagatacaacaaaaaagtcttttaaccggatagagatggataaatatattactgcatttgatgatttgtggaaagaatggaacgacttatctctagaaaacgataattgcccttcaatgtataagatttcttctaactttttttggttttcctgtaggtatacaagagaccattgataaatacaggaaatagttttcatttttttcttcttatatgttatagaatataaaaaaaataaatgaatattacttagtggttagtattagtctagtcgtcttactattaatagtagcaaaaagtatgagaaagaaacataaaataacgaaagtaaaattatgatcattattattatataaattcttaaaaacggtattttct |
| 74b | BT4085 | N/A | attttccgggtagtgctgacttttgtcgtccttatttttatcgtattttcaattgggaaagcatatatgatgaagatgattgatagctactgacggattcatcttggtaaattgatttagaaagtcatttcttcataaaagggagaaatgactttctttttttaatatgttctgagctttcctgcatacaagtaaatttattaataaaaaacatgtaattgttttagtggtatatcatttttaaatcgtcattagataaataacaataaaatagtacacatattaaaaagaactaataata |
| 75a | BT4114 | TSS_4150 | ttgagtaaagaatacaccaatctggaggtatatctgaatgagctagtgaatcatctgtatcctgcggattctaaaaaataaaactaaaagcccgaataaacagttataaaagattaagacgagagtacatttccccataaatcctcgtctttattcattttatgccggaaaaacaaaaaatactatcgtaaaaacaaatattacacatcccacaagccccaaaaacataattttgcaccagtgagaactctaagagtggacacactcgaatattaataatctaatataaaagaacaaaag |
| 75b | BT4119 | N/A | aattgagtgtccatcaggcggctagcatttagagtatgatgaatgctagccgcctgatggacacttttattctaatccaagcaaatactgaaaatgactagtagaaattactatgtaaagtaaaatatggaatttttggtattttaatcccaaaatctcgtatctgatattttagtgataaaatagtgcacctaaacgctcaaaaagtcccctctctttatgttttatataacttcttttgcacatatgaaacaaaaaataagaactataaatcaggaatattaatgaataaacatgagatgattcctgtttcaattttccccatgttttcaacgattgatttgcatgtaacataagtattgattagatagtaatatgtattaactttaaattgaaatga |
| 76 | BT4136 | TSS_3406 | agtacctaccaatggagaattcagcaacattatgcacaacccattgcctcaaacggtctatttcaatcaaaaagttcaggcacgctttatcaaactggaagccaccactcccactgatacagccgcaaaagttgaaccaaatgaaatcggagtaacaatagccccttgatttacattcatcacaagtactaactaaaagcaaaaatcagaatagaaaagacgccatctattttgatttttacactatataaatcttcttattatcaaattaactccaatgaaggcaaagaaagagaaattagtcttaatgaagataacagatctttaatatataaggactgtctatccaaacttatcgaaaaatttaattaaccacaagattcatgcttattgaaaacacaagaatgaattgttaacacttttaatttctatgattaatgaaaaacaaaatagtggtggaagaatactacaaatattaatcctgttttgtttgtatggt |
| 77 | BT4163 | N/A | acaattatatttgaaaagtttgccatattgcttttatatatggttattcttgtttcaagaaaaaccggaataaattgtattatctacactctttcttatttaaaaaacgcgttcagagcatgtattttcgttcaaaaggcggtattttccatgtttatgtacgattcttcattgctatttcttatataagtcgttaagtttgcaacagtttttcacaggtattatgataaagtgcccgaagaaaaacagaatctaacttttgtttaattaaaaaatagataaaatagatttatattgttacttaacagattttaatattatgagaaatgtgatgaatcgtaagcggcattggctgctattattgctcctctcaccttttttttctctcttgtgaggacaag |
| 78 | BT4248 | N/A | ctttaatcaaactcatctttgtatctttatccatcctatcttactttgttattattctatttattattactcgtatattataaatacacttgagttaagtctatccccacatcccttcgataaaaaaaaatagcaaaattattattaaaaccacaacaacttaataatcaatatactaccacccaattaaaaagaagtttttttttcgaaaagtcattttttttatcttagttctaacagaactaagatggcattttgccgtagcttcacttgctcttttactaacaataggaggattttgg |
| 79 | BT4266 | N/A | ctggaattttttttatgtaatatacccttttacttcttattacggcctattcgccagagattgttggtgaaggagaaaatattattatatatggtagaaattaaggtgtgcattttatttcttatgcatttattcttttggggatctactacctttttttatttagtatatttgaaaataaatagttgtgcttggctgttatgattcatttttatatttaacagattgggataagttgctctagaacttggtagttttataccaataagtgtattttttcctgttttcataggctattcaaaggttattaagtttgtgtgctacttaagacggtgtaaagttattaataaaattggtgaacgcaatgtttttttttattttaattatcggcaaacgatgtttttttttaatttttcattctactactcagatcgtaataagatagatataatcttttaattattgtatattgttgattattagtattttatcgctttggataccaagttctagagcgacttattccaatcggtcaattacgactatatttcttaataattatttatgataatgcgaaattataggccgcctccgggctgtgattaaatttttattatcgtttcttcagcatagatcagaacagtgctgaagagagattgaattcattaaaaatttcaataccccccaagaaaaacaaatgaatgataacagtcaaacacaactatatatatattccaaaacttactaaatattaagataaagcatacacacccaaaaacaaacgtattagagtaaaaatgcacatcttatttctaccacattatcataatgctagatatagaatagtaagttttctacttatccatattaacccaactaattattaacctttaatcacagaaaaaca |
| 80 | BT4299 | TSS_3536 | tggagagcaatagagaccttatgccaaattgaataattttgaagaagtcgtaataggacttggtatgtatgtgtataagtgtcttatatatataattgattaattggttgatggatgcttgggatgacaatttatgaactatatagtagaattgagattctaaaaaaatggtgtttgttgataattaaaataaaaaagatattgtatttttcgattttattaataactttacaccgtcttagaatagtacatgcacttaataacctttgaatagcctatggaaacagtagaaaatatactttggggtataaaaataccaagtcctattacgacttagccccaattaatcaatatgattataaccttaaattaacatttatgaaaactcttagcaacagccctccgcctttcaaaatcggagaattcgatctcatttttttacttttatttcagcacagatcagaataggtatgatatgtattctgagttcattatcaacttcaatacccccaatttaagaagatgaataatagcagtcaaacacacccatatatttcaaacttactcatacatacaaaaacaaacgtataccccttgtatgcataccctagttctacccacatagtagacacacaagagtaggtttccctatcacttttttttaaccaatctaattattaaccttttatcacagaaaaaca |

| | | | |
|---|---|---|---|
| 81 | BT4356 | N/A | cgttcacgggaacctttacaaatcaacgtttggaaaagattctggaatattttaaagtatcctccaaaatacgctggaaacatatcaatgatgacaaggatggaagtgacagaaaaaaagagataatagagatttattgattaacctttaacaatttagtagtatgaaagcaa aacccacttagaaaaaagtatgggaagattttggcagatctccccatactaaaatgaatcaattgaaaatgcttgtttggcagacaagcatagttaactcattttcaacttaaatcaatacaaaatt |
| 82 | BT4403 | TSS_3630 | ggaagttgatgctgaagatgttgctcaagatattttctgtaagctttggttacaaccggaaatatggcttgacaatgaaaatgaattagataactacctctttataatgaccagaaacattattctgaatatcttcaagcatcagcaaatcgaacgggaatatcaggaaggatac cttgagaaaacagtacttcatgaattgatagaaggggaagatgcccttaataatatatattatgaagaaatgttgctggttgttcgcctgactttggaaaaaatgcctgaacggcgaaggttaatattcgagctgagtcgttttaagggattaagttataaagaaatcgcagaa aagcttaatgtttccatccgtacggtagagcaccaggtgtatttagcactgatagaactaaaaaaagtgcttctatttttattttttctattttcctttaagtagtgtttttcttttagttgtatttagtgtataaaggaagaaatta |
| 83 | BT4470 | TSS_3675 | tctttgtctccgaaaaacaagagtccgacaatgcgccacatatcattgttcaggttcggataataaaaatccatcgaccaccgttttctttgtggcggaaaacttcccagcatcagcaactttgcttttggtgggagaaagggttttaaaggatgtttttcgatttcaatctccatattg atgtaatttcctacaaaaataacgaatctattcttttccttcaaaatagttgcggaaaaaaacaaatttaaaaatatttggctatttttgcacggagatggtgaaaaacaacttattgaa |
| 84 | BT4635 | N/A | ctcccaataacttattaagttataatgaacaggaaggcatccatatccaatcggtagacgtacaaaaatatatcgcctggaaagatggtttttacctcttccaaacagaaaaactaaaagacattgctacaaaactcagcgattactatggcaaaaagatcatgatagaca gtccattgaagaccattacctgcagtggtaaactagatttgaaagaagatcttgacgaagttcttcagacactaatacggactgtcccggcaagaatagaaaaatcagatggaattatccatatttatgtaaaacattaattaaaatcgaaagcctatgaataaacattatcct taaaaaaataaggccgaataatacccttaatatcatccggcctttgattaaattacttacacgttaatattaattaatctaatcattacaaatgtatgaaaaagcaccgattatttagtcaccaaaggactaaagataaacaactattactaatc |
| 85 | BT4662 | N/A | ttttttcttcttttattgaatgatgacggattttaaaatcatccatccaaacaacctaaattgtacagaacctattccatatcatattacaatgtcataacattatgacacataaaaaactaggatacaggagataaacagctttttaaacaaaaacaacatatgtctcaacaaattt gatatcaaaaaaatcatttttgaacaaagtatgaacaacctaaattaaagaatctaattaatgaggttacatttgcaacagtaaaataacacttttaataattataaaacctaataaagt |
| 86 | BT4672 | N/A | tgtctcccgtggaatatatcaagtccatccgcatgaagaaagctgccatgttattgcaacaaaagaaattcaccgttgccgaagtgatgtatatggtaggattttccaatcattcttatttctccaaatgctttcaggcagagtttggaaaaacaccgcgccaatacttgaatgac gggctgtagaggcattttagggcttttcgtcctatttttatccttatatgtccaatctgtgcttttatgaaaatgccaggcagtctatctttgccgctatcaacaaaccactaaatttgatagc |
| 87 | BT4706 | TSS_3898, TSS_3899 | caggcagatgatacattcacaaagaaaaagataatagtagatttatattagtaaccataaaaatgagaatagcctatgagataacacaaagatcgtaaaaaaaggaagaaagctgctaccaacagctgtcttcccccaatttttcagattcgcggtaaggagtcttgcaa acgctaaccggaatcaaaccacagttcattatagtaataaactttagttaaccgcaaatgtatgaaaaaaaatcattcatttattgcattatatcctaaatatgctttaaatctaaaactacctttagtt |
| 88 | BT4723 | N/A | acttcctaaagaatactttacgggaacgattcctagtggcggcatacagaacgccttgaatattctcatgcttacttcgcctttttattatgaaatggatggctcggtcattgtcttgaaagagaagtagataagcatctttatcatgaaattagatatagccattcctgagttttcccc cggaatggctttttaatttatgaagaaccgttagttaacaagagttaaagatagatagccgattggacgaaaggggagagcagtcggtatacctgtcaaacaacttaaaagttttgatat |
