## Supplementary Table 3 for "A high-throughput microbial glycomics platform for prebiotic development"

| Supplementary Table 3. PUL similarity across *Bacteroides* species | | | |
|---|---|---|---|
| **PUL ID** | **original_species** | **similar_species** | **species_pattern** |
| B. thetaiotaomicron PUL_1 | *Bt* | *Bt* | *Bt* |
| B. thetaiotaomicron PUL_2 | *Bt* | *Bc, Bt, Bu* | *Bc_Bt_Bu* |
| B. thetaiotaomicron PUL_3 | *Bt* | *Bt* | *Bt* |
| B. thetaiotaomicron PUL_4 | *Bt* | *Bt* | *Bt* |
| B. thetaiotaomicron PUL_5 | *Bt* | *Bo, Bt* | *Bo_Bt* |
| B. thetaiotaomicron PUL_6 | *Bt* | *Bt* | *Bt* |
| B. thetaiotaomicron PUL_7 | *Bt* | *Bc, Bo, Bt* | *Bc_Bo_Bt* |
| B. thetaiotaomicron PUL_8 | *Bt* | *Bt* | *Bt* |
| B. thetaiotaomicron PUL_9 | *Bt* | *Bf, Bo, Bt, Bu* | *Bf_Bo_Bt_Bu* |
| B. thetaiotaomicron PUL_10 | *Bt* | *Bt* | *Bt* |
| B. thetaiotaomicron PUL_11 | *Bt* | *Bf, Bo, Bt* | *Bf_Bo_Bt* |
| B. thetaiotaomicron PUL_12 | *Bt* | *Bt* | *Bt* |
| B. thetaiotaomicron PUL_13 | *Bt* | *Bt, Bu* | *Bt_Bu* |
| B. thetaiotaomicron PUL_14 | *Bt* | *Bo, Bt, Bu* | *Bo_Bt_Bu* |
| B. thetaiotaomicron PUL_15 | *Bt* | *Bt* | *Bt* |
| B. thetaiotaomicron PUL_16 | *Bt* | *Bf, Bo, Bt* | *Bf_Bo_Bt* |
| B. thetaiotaomicron PUL_17 | *Bt* | *Bt* | *Bt* |
| B. thetaiotaomicron PUL_18 | *Bt* | *Bt* | *Bt* |
| B. thetaiotaomicron PUL_19 | *Bt* | *Bt* | *Bt* |
| B. thetaiotaomicron PUL_20 | *Bt* | *Bc, Bo, Bt* | *Bc_Bo_Bt* |
| B. thetaiotaomicron PUL_21 | *Bt* | *Bt* | *Bt* |
| B. thetaiotaomicron PUL_22 | *Bt* | *Bo, Bt, Bu* | *Bo_Bt_Bu* |
| B. thetaiotaomicron PUL_23 | *Bt* | *Bc, Bf, Bo, Bt, Bu* | *Bc_Bf_Bo_Bt_Bu* |
| B. thetaiotaomicron PUL_24 | *Bt* | *Bc, Bo, Bt, Bu* | *Bc_Bo_Bt_Bu* |
| B. thetaiotaomicron PUL_25 | *Bt* | *Bt* | *Bt* |
| B. thetaiotaomicron PUL_26 | *Bt* | *Bc, Bo, Bt, Bu* | *Bc_Bo_Bt_Bu* |
| B. thetaiotaomicron PUL_27 | *Bt* | *Bt* | *Bt* |
| B. thetaiotaomicron PUL_28 | *Bt* | *Bc, Bo, Bt, Bu* | *Bc_Bo_Bt_Bu* |
| B. thetaiotaomicron PUL_29 | *Bt* | *Bt* | *Bt* |
| B. thetaiotaomicron PUL_30 | *Bt* | *Bt* | *Bt* |
| B. thetaiotaomicron PUL_31 | *Bt* | *Bt* | *Bt* |
| B. thetaiotaomicron PUL_32 | *Bt* | *Bt* | *Bt* |
| B. thetaiotaomicron PUL_33 | *Bt* | *Bf, Bo, Bt* | *Bf_Bo_Bt* |
| B. thetaiotaomicron PUL_34 | *Bt* | *Bt* | *Bt* |
| B. thetaiotaomicron PUL_35 | *Bt* | *Bt* | *Bt* |
| B. thetaiotaomicron PUL_36 | *Bt* | *Bf, Bo, Bt, Bu* | *Bf_Bo_Bt_Bu* |
| B. thetaiotaomicron PUL_37 | *Bt* | *Bt* | *Bt* |
| B. thetaiotaomicron PUL_38 | *Bt* | *Bt* | *Bt* |
| B. thetaiotaomicron PUL_39 | *Bt* | *Bc, Bf, Bt, Bu* | *Bc_Bf_Bt_Bu* |
| B. thetaiotaomicron PUL_40 | *Bt* | *Bt* | *Bt* |
| B. thetaiotaomicron PUL_41 | *Bt* | *Bt, Bu* | *Bt_Bu* |
| B. thetaiotaomicron PUL_42 | *Bt* | *Bc, Bt, Bu* | *Bc_Bt_Bu* |
| B. thetaiotaomicron PUL_43 | *Bt* | *Bt* | *Bt* |

| | | | |
|---|---|---|---|
| B. thetaiotaomicron PUL_44 | *Bt* | *Bt* | *Bt* |
| B. thetaiotaomicron PUL_45 | *Bt* | *Bt* | *Bt* |
| B. thetaiotaomicron PUL_46 | *Bt* | *Bt* | *Bt* |
| B. thetaiotaomicron PUL_47 | *Bt* | *Bt* | *Bt* |
| B. thetaiotaomicron PUL_48 | *Bt* | *Bo, Bt* | *Bo_Bt* |
| B. thetaiotaomicron PUL_49 | *Bt* | *Bf, Bo, Bt, Bu* | *Bf_Bo_Bt_Bu* |
| B. thetaiotaomicron PUL_50 | *Bt* | *Bt* | *Bt* |
| B. thetaiotaomicron PUL_51 | *Bt* | *Bt* | *Bt* |
| B. thetaiotaomicron PUL_52 | *Bt* | *Bt* | *Bt* |
| B. thetaiotaomicron PUL_53 | *Bt* | *Bt* | *Bt* |
| B. thetaiotaomicron PUL_54 | *Bt* | *Bt* | *Bt* |
| B. thetaiotaomicron PUL_55 | *Bt* | *Bc, Bf, Bt, Bu* | *Bc_Bf_Bt_Bu* |
| B. thetaiotaomicron PUL_56 | *Bt* | *Bo, Bt* | *Bo_Bt* |
| B. thetaiotaomicron PUL_57 | *Bt* | *Bt* | *Bt* |
| B. thetaiotaomicron PUL_58 | *Bt* | *Bt* | *Bt* |
| B. thetaiotaomicron PUL_59 | *Bt* | *Bo, Bt* | *Bo_Bt* |
| B. thetaiotaomicron PUL_60 | *Bt* | *Bf, Bo, Bt, Bu* | *Bf_Bo_Bt_Bu* |
| B. thetaiotaomicron PUL_61 | *Bt* | *Bo, Bt* | *Bo_Bt* |
| B. thetaiotaomicron PUL_62 | *Bt* | *Bt* | *Bt* |
| B. thetaiotaomicron PUL_63 | *Bt* | *Bt* | *Bt* |
| B. thetaiotaomicron PUL_64 | *Bt* | *Bc, Bo, Bt, Bu* | *Bc_Bo_Bt_Bu* |
| B. thetaiotaomicron PUL_65 | *Bt* | *Bc, Bt* | *Bc_Bt* |
| B. thetaiotaomicron PUL_66 | *Bt* | *Bo, Bt* | *Bo_Bt* |
| B. thetaiotaomicron PUL_67 | *Bt* | *Bf, Bo, Bt* | *Bf_Bo_Bt* |
| B. thetaiotaomicron PUL_68 | *Bt* | *Bo, Bt, Bu* | *Bo_Bt_Bu* |
| B. thetaiotaomicron PUL_69 | *Bt* | *Bt* | *Bt* |
| B. thetaiotaomicron PUL_70 | *Bt* | *Bt* | *Bt* |
| B. thetaiotaomicron PUL_71 | *Bt* | *Bc, Bf, Bo, Bt* | *Bc_Bf_Bo_Bt* |
| B. thetaiotaomicron PUL_72 | *Bt* | *Bc, Bf, Bo, Bt, Bu* | *Bc_Bf_Bo_Bt_Bu* |
| B. thetaiotaomicron PUL_73 | *Bt* | *Bt* | *Bt* |
| B. thetaiotaomicron PUL_74 | *Bt* | *Bf, Bo, Bt, Bu* | *Bf_Bo_Bt_Bu* |
| B. thetaiotaomicron PUL_75 | *Bt* | *Bc, Bo, Bt* | *Bc_Bo_Bt* |
| B. thetaiotaomicron PUL_76 | *Bt* | *Bt* | *Bt* |
| B. thetaiotaomicron PUL_77 | *Bt* | *Bo, Bt, Bu* | *Bo_Bt_Bu* |
| B. thetaiotaomicron PUL_78 | *Bt* | *Bc, Bo, Bt, Bu* | *Bc_Bo_Bt_Bu* |
| B. thetaiotaomicron PUL_79 | *Bt* | *Bt* | *Bt* |
| B. thetaiotaomicron PUL_80 | *Bt* | *Bt* | *Bt* |
| B. thetaiotaomicron PUL_81 | *Bt* | *Bt* | *Bt* |
| B. thetaiotaomicron PUL_82 | *Bt* | *Bf, Bo, Bt* | *Bf_Bo_Bt* |
| B. thetaiotaomicron PUL_83 | *Bt* | *Bt* | *Bt* |
| B. thetaiotaomicron PUL_84 | *Bt* | *Bf, Bt* | *Bf_Bt* |
| B. thetaiotaomicron PUL_85 | *Bt* | *Bc, Bo, Bt* | *Bc_Bo_Bt* |
| B. thetaiotaomicron PUL_86 | *Bt* | *Bc, Bo, Bt* | *Bc_Bo_Bt* |
| B. thetaiotaomicron PUL_87 | *Bt* | *Bo, Bt* | *Bo_Bt* |
| B. thetaiotaomicron PUL_88 | *Bt* | *Bt* | *Bt* |

| | | | |
|---|---|---|---|
| B. thetaiotaomicron PUL_89 | *Bt* | *Bf, Bo, Bt* | *Bf_Bo_Bt* |
| B. thetaiotaomicron PUL_90 | *Bt* | *Bf, Bo, Bt, Bu* | *Bf_Bo_Bt_Bu* |
| B. thetaiotaomicron PUL_91 | *Bt* | *Bo, Bt, Bu* | *Bo_Bt_Bu* |
| B. thetaiotaomicron PUL_92 | *Bt* | *Bc, Bf, Bo, Bt* | *Bc_Bf_Bo_Bt* |
| B. thetaiotaomicron PUL_93 | *Bt* | *Bc, Bo, Bt, Bu* | *Bc_Bo_Bt_Bu* |
| B. thetaiotaomicron PUL_94 | *Bt* | *Bf, Bo, Bt* | *Bf_Bo_Bt* |
| B. thetaiotaomicron PUL_95 | *Bt* | *Bt* | *Bt* |
| B. thetaiotaomicron PUL_96 | *Bt* | *Bt* | *Bt* |
| B. fragilis PUL_1 | *Bf* | *Bf* | *Bf* |
| B. fragilis PUL_2 | *Bf* | *Bc, Bf, Bo, Bt, Bu* | *Bc_Bf_Bo_Bt_Bu* |
| B. fragilis PUL_3 | *Bf* | *Bc, Bf, Bo, Bt* | *Bc_Bf_Bo_Bt* |
| B. fragilis PUL_4 | *Bf* | *Bf* | *Bf* |
| B. fragilis PUL_5 | *Bf* | *Bf* | *Bf* |
| B. fragilis PUL_6 | *Bf* | *Bf* | *Bf* |
| B. fragilis PUL_7 | *Bf* | *Bf* | *Bf* |
| B. fragilis PUL_8 | *Bf* | *Bf* | *Bf* |
| B. fragilis PUL_9 | *Bf* | *Bf, Bo, Bt* | *Bf_Bo_Bt* |
| B. fragilis PUL_10 | *Bf* | *Bf* | *Bf* |
| B. fragilis PUL_11 | *Bf* | *Bf* | *Bf* |
| B. fragilis PUL_12 | *Bf* | *Bc, Bf, Bo, Bt, Bu* | *Bc_Bf_Bo_Bt_Bu* |
| B. fragilis PUL_13 | *Bf* | *Bf, Bo* | *Bf_Bo* |
| B. fragilis PUL_14 | *Bf* | *Bf, Bo* | *Bf_Bo* |
| B. fragilis PUL_15 | *Bf* | *Bf* | *Bf* |
| B. fragilis PUL_16 | *Bf* | *Bf* | *Bf* |
| B. fragilis PUL_17 | *Bf* | *Bf* | *Bf* |
| B. fragilis PUL_18 | *Bf* | *Bf* | *Bf* |
| B. fragilis PUL_19 | *Bf* | *Bf* | *Bf* |
| B. fragilis PUL_20 | *Bf* | *Bf* | *Bf* |
| B. fragilis PUL_21 | *Bf* | *Bf* | *Bf* |
| B. fragilis PUL_22 | *Bf* | *Bf, Bo* | *Bf_Bo* |
| B. fragilis PUL_23 | *Bf* | *Bf, Bo* | *Bf_Bo* |
| B. fragilis PUL_24 | *Bf* | *Bf* | *Bf* |
| B. fragilis PUL_25 | *Bf* | *Bf* | *Bf* |
| B. fragilis PUL_26 | *Bf* | *Bf* | *Bf* |
| B. fragilis PUL_27 | *Bf* | *Bf* | *Bf* |
| B. fragilis PUL_28 | *Bf* | *Bf* | *Bf* |
| B. fragilis PUL_29 | *Bf* | *Bf* | *Bf* |
| B. fragilis PUL_30 | *Bf* | *Bc, Bf, Bo, Bt* | *Bc_Bf_Bo_Bt* |
| B. fragilis PUL_31 | *Bf* | *Bf* | *Bf* |
| B. fragilis PUL_32 | *Bf* | *Bf* | *Bf* |
| B. fragilis PUL_33 | *Bf* | *Bf* | *Bf* |
| B. fragilis PUL_34 | *Bf* | *Bf* | *Bf* |
| B. fragilis PUL_35 | *Bf* | *Bf, Bo, Bu* | *Bf_Bo_Bu* |
| B. fragilis PUL_36 | *Bf* | *Bf* | *Bf* |
| B. fragilis PUL_37 | *Bf* | *Bf* | *Bf* |

| | | | |
|---|---|---|---|
| B. fragilis PUL_38 | *Bf* | *Bf, Bo, Bt* | *Bf_Bo_Bt* |
| B. fragilis PUL_39 | *Bf* | *Bf* | *Bf* |
| B. fragilis PUL_40 | *Bf* | *Bf* | *Bf* |
| B. fragilis PUL_41 | *Bf* | *Bf* | *Bf* |
| B. fragilis PUL_42 | *Bf* | *Bf, Bu* | *Bf_Bu* |
| B. fragilis PUL_43 | *Bf* | *Bf, Bo, Bt* | *Bf_Bo_Bt* |
| B. fragilis PUL_44 | *Bf* | *Bf* | *Bf* |
| B. fragilis PUL_45 | *Bf* | *Bf* | *Bf* |
| B. fragilis PUL_46 | *Bf* | *Bf* | *Bf* |
| B. fragilis PUL_47 | *Bf* | *Bf* | *Bf* |
| B. fragilis PUL_48 | *Bf* | *Bf, Bt* | *Bf_Bt* |
| B. fragilis PUL_49 | *Bf* | *Bf* | *Bf* |
| B. fragilis PUL_50 | *Bf* | *Bf* | *Bf* |
| B. fragilis PUL_51 | *Bf* | *Bf* | *Bf* |
| B. fragilis PUL_52 | *Bf* | *Bf, Bu* | *Bf_Bu* |
| B. fragilis PUL_53 | *Bf* | *Bf* | *Bf* |
| B. fragilis PUL_54 | *Bf* | *Bf* | *Bf* |
| B. fragilis PUL_55 | *Bf* | *Bf* | *Bf* |
| B. ovatus PUL_1 | *Bo* | *Bo* | *Bo* |
| B. ovatus PUL_2 | *Bo* | *Bo* | *Bo* |
| B. ovatus PUL_3 | *Bo* | *Bf, Bo, Bt* | *Bf_Bo_Bt* |
| B. ovatus PUL_4 | *Bo* | *Bo* | *Bo* |
| B. ovatus PUL_5 | *Bo* | *Bo* | *Bo* |
| B. ovatus PUL_6 | *Bo* | *Bo* | *Bo* |
| B. ovatus PUL_7 | *Bo* | *Bo* | *Bo* |
| B. ovatus PUL_8 | *Bo* | *Bc, Bf, Bo, Bt, Bu* | *Bc_Bf_Bo_Bt_Bu* |
| B. ovatus PUL_9 | *Bo* | *Bf, Bo, Bt* | *Bf_Bo_Bt* |
| B. ovatus PUL_10 | *Bo* | *Bo* | *Bo* |
| B. ovatus PUL_11 | *Bo* | *Bo* | *Bo* |
| B. ovatus PUL_12 | *Bo* | *Bo* | *Bo* |
| B. ovatus PUL_13 | *Bo* | *Bo, Bt, Bu* | *Bo_Bt_Bu* |
| B. ovatus PUL_14 | *Bo* | *Bo* | *Bo* |
| B. ovatus PUL_15 | *Bo* | *Bo, Bt* | *Bo_Bt* |
| B. ovatus PUL_16 | *Bo* | *Bo* | *Bo* |
| B. ovatus PUL_17 | *Bo* | *Bo* | *Bo* |
| B. ovatus PUL_18 | *Bo* | *Bo, Bt* | *Bo_Bt* |
| B. ovatus PUL_19 | *Bo* | *Bo* | *Bo* |
| B. ovatus PUL_20 | *Bo* | *Bo* | *Bo* |
| B. ovatus PUL_21 | *Bo* | *Bo* | *Bo* |
| B. ovatus PUL_22 | *Bo* | *Bf, Bo, Bt* | *Bf_Bo_Bt* |
| B. ovatus PUL_23 | *Bo* | *Bo* | *Bo* |
| B. ovatus PUL_24 | *Bo* | *Bo* | *Bo* |
| B. ovatus PUL_25 | *Bo* | *Bo* | *Bo* |
| B. ovatus PUL_26 | *Bo* | *Bo* | *Bo* |
| B. ovatus PUL_27 | *Bo* | *Bo* | *Bo* |

| | | | |
|---|---|---|---|
| B. ovatus PUL_28 | *Bo* | *Bo* | *Bo* |
| B. ovatus PUL_29 | *Bo* | *Bo* | *Bo* |
| B. ovatus PUL_30 | *Bo* | *Bf, Bo* | *Bf_Bo* |
| B. ovatus PUL_31 | *Bo* | *Bo* | *Bo* |
| B. ovatus PUL_32 | *Bo* | *Bo* | *Bo* |
| B. ovatus PUL_33 | *Bo* | *Bo* | *Bo* |
| B. ovatus PUL_34 | *Bo* | *Bo* | *Bo* |
| B. ovatus PUL_35 | *Bo* | *Bo* | *Bo* |
| B. ovatus PUL_36 | *Bo* | *Bo* | *Bo* |
| B. ovatus PUL_37 | *Bo* | *Bc, Bo* | *Bc_Bo* |
| B. ovatus PUL_38 | *Bo* | *Bo* | *Bo* |
| B. ovatus PUL_39 | *Bo* | *Bo* | *Bo* |
| B. ovatus PUL_40 | *Bo* | *Bo* | *Bo* |
| B. ovatus PUL_41 | *Bo* | *Bo* | *Bo* |
| B. ovatus PUL_42 | *Bo* | *Bc, Bo* | *Bc_Bo* |
| B. ovatus PUL_43 | *Bo* | *Bc, Bf, Bo, Bt, Bu* | *Bc_Bf_Bo_Bt_Bu* |
| B. ovatus PUL_44 | *Bo* | *Bc, Bf, Bo, Bt, Bu* | *Bc_Bf_Bo_Bt_Bu* |
| B. ovatus PUL_45 | *Bo* | *Bf, Bo, Bt* | *Bf_Bo_Bt* |
| B. ovatus PUL_46 | *Bo* | *Bo, Bu* | *Bo_Bu* |
| B. ovatus PUL_47 | *Bo* | *Bf, Bo* | *Bf_Bo* |
| B. ovatus PUL_48 | *Bo* | *Bo* | *Bo* |
| B. ovatus PUL_49 | *Bo* | *Bc, Bo, Bt, Bu* | *Bc_Bo_Bt_Bu* |
| B. ovatus PUL_50 | *Bo* | *Bo* | *Bo* |
| B. ovatus PUL_51 | *Bo* | *Bf, Bo, Bu* | *Bf_Bo_Bu* |
| B. ovatus PUL_52 | *Bo* | *Bo* | *Bo* |
| B. ovatus PUL_53 | *Bo* | *Bo, Bt* | *Bo_Bt* |
| B. ovatus PUL_54 | *Bo* | *Bc, Bo, Bt, Bu* | *Bc_Bo_Bt_Bu* |
| B. ovatus PUL_55 | *Bo* | *Bo* | *Bo* |
| B. ovatus PUL_56 | *Bo* | *Bo* | *Bo* |
| B. ovatus PUL_57 | *Bo* | *Bo* | *Bo* |
| B. ovatus PUL_58 | *Bo* | *Bo* | *Bo* |
| B. ovatus PUL_59 | *Bo* | *Bo* | *Bo* |
| B. ovatus PUL_60 | *Bo* | *Bc, Bf, Bo* | *Bc_Bf_Bo* |
| B. ovatus PUL_61 | *Bo* | *Bo* | *Bo* |
| B. ovatus PUL_62 | *Bo* | *Bo* | *Bo* |
| B. ovatus PUL_63 | *Bo* | *Bo* | *Bo* |
| B. ovatus PUL_64 | *Bo* | *Bf, Bo, Bt* | *Bf_Bo_Bt* |
| B. ovatus PUL_65 | *Bo* | *Bo* | *Bo* |
| B. ovatus PUL_66 | *Bo* | *Bo* | *Bo* |
| B. ovatus PUL_67 | *Bo* | *Bo, Bt* | *Bo_Bt* |
| B. ovatus PUL_68 | *Bo* | *Bo* | *Bo* |
| B. ovatus PUL_69 | *Bo* | *Bo* | *Bo* |
| B. ovatus PUL_70 | *Bo* | *Bf, Bo, Bt, Bu* | *Bf_Bo_Bt_Bu* |
| B. ovatus PUL_71 | *Bo* | *Bo, Bt* | *Bo_Bt* |
| B. ovatus PUL_72 | *Bo* | *Bo* | *Bo* |

| | | | |
|---|---|---|---|
| B. ovatus PUL_73 | *Bo* | *Bc, Bo, Bt, Bu* | *Bc_Bo_Bt_Bu* |
| B. ovatus PUL_74 | *Bo* | *Bo* | *Bo* |
| B. ovatus PUL_75 | *Bo* | *Bo* | *Bo* |
| B. ovatus PUL_76 | *Bo* | *Bo, Bt* | *Bo_Bt* |
| B. ovatus PUL_77 | *Bo* | *Bf, Bo, Bt* | *Bf_Bo_Bt* |
| B. ovatus PUL_78 | *Bo* | *Bo* | *Bo* |
| B. ovatus PUL_79 | *Bo* | *Bo* | *Bo* |
| B. ovatus PUL_80 | *Bo* | *Bo* | *Bo* |
| B. ovatus PUL_81 | *Bo* | *Bo, Bu* | *Bo_Bu* |
| B. ovatus PUL_82 | *Bo* | *Bo, Bt, Bu* | *Bo_Bt_Bu* |
| B. ovatus PUL_83 | *Bo* | *Bc, Bf, Bo, Bt* | *Bc_Bf_Bo_Bt* |
| B. ovatus PUL_84 | *Bo* | *Bo* | *Bo* |
| B. ovatus PUL_85 | *Bo* | *Bf, Bo, Bt* | *Bf_Bo_Bt* |
| B. ovatus PUL_86 | *Bo* | *Bc, Bo* | *Bc_Bo* |
| B. ovatus PUL_87 | *Bo* | *Bo* | *Bo* |
| B. ovatus PUL_88 | *Bo* | *Bo* | *Bo* |
| B. ovatus PUL_89 | *Bo* | *Bo* | *Bo* |
| B. ovatus PUL_90 | *Bo* | *Bo* | *Bo* |
| B. ovatus PUL_91 | *Bo* | *Bc, Bf, Bo, Bt* | *Bc_Bf_Bo_Bt* |
| B. ovatus PUL_92 | *Bo* | *Bo* | *Bo* |
| B. ovatus PUL_93 | *Bo* | *Bo* | *Bo* |
| B. ovatus PUL_94 | *Bo* | *Bo* | *Bo* |
| B. ovatus PUL_95 | *Bo* | *Bo* | *Bo* |
| B. ovatus PUL_96 | *Bo* | *Bo* | *Bo* |
| B. ovatus PUL_97 | *Bo* | *Bo, Bt* | *Bo_Bt* |
| B. ovatus PUL_98 | *Bo* | *Bc, Bf, Bo, Bt, Bu* | *Bc_Bf_Bo_Bt_Bu* |
| B. ovatus PUL_99 | *Bo* | *Bc, Bo, Bt, Bu* | *Bc_Bo_Bt_Bu* |
| B. ovatus PUL_100 | *Bo* | *Bc, Bo, Bt* | *Bc_Bo_Bt* |
| B. ovatus PUL_101 | *Bo* | *Bo, Bt, Bu* | *Bo_Bt_Bu* |
| B. ovatus PUL_102 | *Bo* | *Bo, Bt* | *Bo_Bt* |
| B. ovatus PUL_103 | *Bo* | *Bf, Bo, Bt* | *Bf_Bo_Bt* |
| B. ovatus PUL_104 | *Bo* | *Bf, Bo* | *Bf_Bo* |
| B. ovatus PUL_105 | *Bo* | *Bo* | *Bo* |
| B. ovatus PUL_106 | *Bo* | *Bo* | *Bo* |
| B. ovatus PUL_107 | *Bo* | *Bc, Bo, Bt* | *Bc_Bo_Bt* |
| B. ovatus PUL_108 | *Bo* | *Bc, Bo, Bt* | *Bc_Bo_Bt* |
| B. ovatus PUL_109 | *Bo* | *Bo* | *Bo* |
| B. ovatus PUL_110 | *Bo* | *Bf, Bo, Bt* | *Bf_Bo_Bt* |
| B. ovatus PUL_111 | *Bo* | *Bo* | *Bo* |
| B. ovatus PUL_112 | *Bo* | *Bo* | *Bo* |
| B. ovatus PUL_113 | *Bo* | *Bf, Bo, Bu* | *Bf_Bo_Bu* |
| B. ovatus PUL_114 | *Bo* | *Bc, Bf, Bo, Bt, Bu* | *Bc_Bf_Bo_Bt_Bu* |
| B. ovatus PUL_115 | *Bo* | *Bc, Bo, Bt* | *Bc_Bo_Bt* |
| B. cellulosilyticus PUL_1 | *Bc* | *Bc* | *Bc* |
| B. cellulosilyticus PUL_2 | *Bc* | *Bc* | *Bc* |

| | | | |
|---|---|---|---|
| B. cellulosilyticus PUL_3 | *Bc* | *Bc* | *Bc* |
| B. cellulosilyticus PUL_4 | *Bc* | *Bc* | *Bc* |
| B. cellulosilyticus PUL_5 | *Bc* | *Bc* | *Bc* |
| B. cellulosilyticus PUL_6 | *Bc* | *Bc, Bo, Bt* | *Bc_Bo_Bt* |
| B. cellulosilyticus PUL_7 | *Bc* | *Bc* | *Bc* |
| B. cellulosilyticus PUL_8 | *Bc* | *Bc* | *Bc* |
| B. cellulosilyticus PUL_9 | *Bc* | *Bc* | *Bc* |
| B. cellulosilyticus PUL_10 | *Bc* | *Bc, Bt* | *Bc_Bt* |
| B. cellulosilyticus PUL_11 | *Bc* | *Bc, Bf, Bo, Bt* | *Bc_Bf_Bo_Bt* |
| B. cellulosilyticus PUL_12 | *Bc* | *Bc* | *Bc* |
| B. cellulosilyticus PUL_13 | *Bc* | *Bc* | *Bc* |
| B. cellulosilyticus PUL_14 | *Bc* | *Bc* | *Bc* |
| B. cellulosilyticus PUL_15 | *Bc* | *Bc* | *Bc* |
| B. cellulosilyticus PUL_16 | *Bc* | *Bc, Bo, Bt* | *Bc_Bo_Bt* |
| B. cellulosilyticus PUL_17 | *Bc* | *Bc, Bo, Bt, Bu* | *Bc_Bo_Bt_Bu* |
| B. cellulosilyticus PUL_18 | *Bc* | *Bc* | *Bc* |
| B. cellulosilyticus PUL_19 | *Bc* | *Bc* | *Bc* |
| B. cellulosilyticus PUL_20 | *Bc* | *Bc* | *Bc* |
| B. cellulosilyticus PUL_21 | *Bc* | *Bc* | *Bc* |
| B. cellulosilyticus PUL_22 | *Bc* | *Bc* | *Bc* |
| B. cellulosilyticus PUL_23 | *Bc* | *Bc* | *Bc* |
| B. cellulosilyticus PUL_24 | *Bc* | *Bc, Bu* | *Bc_Bu* |
| B. cellulosilyticus PUL_25 | *Bc* | *Bc* | *Bc* |
| B. cellulosilyticus PUL_26 | *Bc* | *Bc, Bt* | *Bc_Bt* |
| B. cellulosilyticus PUL_27 | *Bc* | *Bc* | *Bc* |
| B. cellulosilyticus PUL_28 | *Bc* | *Bc* | *Bc* |
| B. cellulosilyticus PUL_29 | *Bc* | *Bc, Bf, Bo, Bt* | *Bc_Bf_Bo_Bt* |
| B. cellulosilyticus PUL_30 | *Bc* | *Bc* | *Bc* |
| B. cellulosilyticus PUL_31 | *Bc* | *Bc, Bu* | *Bc_Bu* |
| B. cellulosilyticus PUL_32 | *Bc* | *Bc* | *Bc* |
| B. cellulosilyticus PUL_33 | *Bc* | *Bc* | *Bc* |
| B. cellulosilyticus PUL_34 | *Bc* | *Bc* | *Bc* |
| B. cellulosilyticus PUL_35 | *Bc* | *Bc, Bo, Bt, Bu* | *Bc_Bo_Bt_Bu* |
| B. cellulosilyticus PUL_36 | *Bc* | *Bc* | *Bc* |
| B. cellulosilyticus PUL_37 | *Bc* | *Bc* | *Bc* |
| B. cellulosilyticus PUL_38 | *Bc* | *Bc, Bf, Bo, Bt, Bu* | *Bc_Bf_Bo_Bt_Bu* |
| B. cellulosilyticus PUL_39 | *Bc* | *Bc* | *Bc* |
| B. cellulosilyticus PUL_40 | *Bc* | *Bc* | *Bc* |
| B. cellulosilyticus PUL_41 | *Bc* | *Bc* | *Bc* |
| B. cellulosilyticus PUL_42 | *Bc* | *Bc, Bt* | *Bc_Bt* |
| B. cellulosilyticus PUL_43 | *Bc* | *Bc* | *Bc* |
| B. cellulosilyticus PUL_44 | *Bc* | *Bc* | *Bc* |
| B. cellulosilyticus PUL_45 | *Bc* | *Bc* | *Bc* |
| B. cellulosilyticus PUL_46 | *Bc* | *Bc* | *Bc* |
| B. cellulosilyticus PUL_47 | *Bc* | *Bc* | *Bc* |

| | | | |
|---|---|---|---|
| B. cellulosilyticus PUL_48 | *Bc* | *Bc* | *Bc* |
| B. cellulosilyticus PUL_49 | *Bc* | *Bc* | *Bc* |
| B. cellulosilyticus PUL_50 | *Bc* | *Bc, Bf, Bo, Bt, Bu* | *Bc_Bf_Bo_Bt_Bu* |
| B. cellulosilyticus PUL_51 | *Bc* | *Bc* | *Bc* |
| B. cellulosilyticus PUL_52 | *Bc* | *Bc* | *Bc* |
| B. cellulosilyticus PUL_53 | *Bc* | *Bc, Bo, Bu* | *Bc_Bo_Bu* |
| B. cellulosilyticus PUL_54 | *Bc* | *Bc, Bo, Bt* | *Bc_Bo_Bt* |
| B. cellulosilyticus PUL_55 | *Bc* | *Bc, Bo, Bu* | *Bc_Bo_Bu* |
| B. cellulosilyticus PUL_56 | *Bc* | *Bc, Bu* | *Bc_Bu* |
| B. cellulosilyticus PUL_57 | *Bc* | *Bc, Bu* | *Bc_Bu* |
| B. cellulosilyticus PUL_58 | *Bc* | *Bc* | *Bc* |
| B. cellulosilyticus PUL_59 | *Bc* | *Bc* | *Bc* |
| B. cellulosilyticus PUL_60 | *Bc* | *Bc* | *Bc* |
| B. cellulosilyticus PUL_61 | *Bc* | *Bc, Bf, Bo, Bt* | *Bc_Bf_Bo_Bt* |
| B. cellulosilyticus PUL_62 | *Bc* | *Bc* | *Bc* |
| B. cellulosilyticus PUL_63 | *Bc* | *Bc* | *Bc* |
| B. cellulosilyticus PUL_64 | *Bc* | *Bc* | *Bc* |
| B. cellulosilyticus PUL_65 | *Bc* | *Bc* | *Bc* |
| B. cellulosilyticus PUL_66 | *Bc* | *Bc* | *Bc* |
| B. cellulosilyticus PUL_67 | *Bc* | *Bc* | *Bc* |
| B. cellulosilyticus PUL_68 | *Bc* | *Bc* | *Bc* |
| B. cellulosilyticus PUL_69 | *Bc* | *Bc, Bo* | *Bc_Bo* |
| B. cellulosilyticus PUL_70 | *Bc* | *Bc, Bt* | *Bc_Bt* |
| B. cellulosilyticus PUL_71 | *Bc* | *Bc* | *Bc* |
| B. cellulosilyticus PUL_72 | *Bc* | *Bc* | *Bc* |
| B. cellulosilyticus PUL_73 | *Bc* | *Bc* | *Bc* |
| B. cellulosilyticus PUL_74 | *Bc* | *Bc* | *Bc* |
| B. cellulosilyticus PUL_75 | *Bc* | *Bc* | *Bc* |
| B. cellulosilyticus PUL_76 | *Bc* | *Bc* | *Bc* |
| B. cellulosilyticus PUL_77 | *Bc* | *Bc* | *Bc* |
| B. cellulosilyticus PUL_78 | *Bc* | *Bc* | *Bc* |
| B. cellulosilyticus PUL_79 | *Bc* | *Bc* | *Bc* |
| B. cellulosilyticus PUL_80 | *Bc* | *Bc* | *Bc* |
| B. cellulosilyticus PUL_81 | *Bc* | *Bc* | *Bc* |
| B. cellulosilyticus PUL_82 | *Bc* | *Bc* | *Bc* |
| B. cellulosilyticus PUL_83 | *Bc* | *Bc* | *Bc* |
| B. cellulosilyticus PUL_84 | *Bc* | *Bc* | *Bc* |
| B. cellulosilyticus PUL_85 | *Bc* | *Bc* | *Bc* |
| B. cellulosilyticus PUL_86 | *Bc* | *Bc* | *Bc* |
| B. cellulosilyticus PUL_87 | *Bc* | *Bc* | *Bc* |
| B. cellulosilyticus PUL_88 | *Bc* | *Bc* | *Bc* |
| B. cellulosilyticus PUL_89 | *Bc* | *Bc* | *Bc* |
| B. cellulosilyticus PUL_90 | *Bc* | *Bc* | *Bc* |
| B. cellulosilyticus PUL_91 | *Bc* | *Bc* | *Bc* |
| B. cellulosilyticus PUL_92 | *Bc* | *Bc* | *Bc* |

| | | | |
|---|---|---|---|
| B. cellulosilyticus PUL_93 | *Bc* | *Bc, Bo* | *Bc_Bo* |
| B. cellulosilyticus PUL_94 | *Bc* | *Bc* | *Bc* |
| B. cellulosilyticus PUL_95 | *Bc* | *Bc* | *Bc* |
| B. cellulosilyticus PUL_96 | *Bc* | *Bc* | *Bc* |
| B. cellulosilyticus PUL_97 | *Bc* | *Bc* | *Bc* |
| B. cellulosilyticus PUL_98 | *Bc* | *Bc* | *Bc* |
| B. cellulosilyticus PUL_99 | *Bc* | *Bc, Bo, Bt* | *Bc_Bo_Bt* |
| B. cellulosilyticus PUL_100 | *Bc* | *Bc* | *Bc* |
| B. cellulosilyticus PUL_101 | *Bc* | *Bc* | *Bc* |
| B. cellulosilyticus PUL_102 | *Bc* | *Bc* | *Bc* |
| B. uniformis PUL_1 | *Bu* | *Bu* | *Bu* |
| B. uniformis PUL_2 | *Bu* | *Bu* | *Bu* |
| B. uniformis PUL_3 | *Bu* | *Bu* | *Bu* |
| B. uniformis PUL_4 | *Bu* | *Bu* | *Bu* |
| B. uniformis PUL_5 | *Bu* | *Bu* | *Bu* |
| B. uniformis PUL_6 | *Bu* | *Bu* | *Bu* |
| B. uniformis PUL_7 | *Bu* | *Bc, Bf, Bo, Bt, Bu* | *Bc_Bf_Bo_Bt_Bu* |
| B. uniformis PUL_8 | *Bu* | *Bu* | *Bu* |
| B. uniformis PUL_9 | *Bu* | *Bu* | *Bu* |
| B. uniformis PUL_10 | *Bu* | *Bu* | *Bu* |
| B. uniformis PUL_11 | *Bu* | *Bc, Bo, Bu* | *Bc_Bo_Bu* |
| B. uniformis PUL_12 | *Bu* | *Bc, Bo, Bt, Bu* | *Bc_Bo_Bt_Bu* |
| B. uniformis PUL_13 | *Bu* | *Bc, Bf, Bu* | *Bc_Bf_Bu* |
| B. uniformis PUL_14 | *Bu* | *Bu* | *Bu* |
| B. uniformis PUL_15 | *Bu* | *Bu* | *Bu* |
| B. uniformis PUL_16 | *Bu* | *Bc, Bf, Bo, Bt, Bu* | *Bc_Bf_Bo_Bt_Bu* |
| B. uniformis PUL_17 | *Bu* | *Bt, Bu* | *Bt_Bu* |
| B. uniformis PUL_18 | *Bu* | *Bu* | *Bu* |
| B. uniformis PUL_19 | *Bu* | *Bu* | *Bu* |
| B. uniformis PUL_20 | *Bu* | *Bu* | *Bu* |
| B. uniformis PUL_21 | *Bu* | *Bo, Bu* | *Bo_Bu* |
| B. uniformis PUL_22 | *Bu* | *Bc, Bu* | *Bc_Bu* |
| B. uniformis PUL_23 | *Bu* | *Bu* | *Bu* |
| B. uniformis PUL_24 | *Bu* | *Bu* | *Bu* |
| B. uniformis PUL_25 | *Bu* | *Bu* | *Bu* |
| B. uniformis PUL_26 | *Bu* | *Bu* | *Bu* |
| B. uniformis PUL_27 | *Bu* | *Bu* | *Bu* |
| B. uniformis PUL_28 | *Bu* | *Bc, Bf, Bo, Bt, Bu* | *Bc_Bf_Bo_Bt_Bu* |
| B. uniformis PUL_29 | *Bu* | *Bu* | *Bu* |
| B. uniformis PUL_30 | *Bu* | *Bf, Bt, Bu* | *Bf_Bt_Bu* |
| B. uniformis PUL_31 | *Bu* | *Bu* | *Bu* |
| B. uniformis PUL_32 | *Bu* | *Bf, Bo, Bt, Bu* | *Bf_Bo_Bt_Bu* |
| B. uniformis PUL_33 | *Bu* | *Bu* | *Bu* |
| B. uniformis PUL_34 | *Bu* | *Bf, Bo, Bt, Bu* | *Bf_Bo_Bt_Bu* |
| B. uniformis PUL_35 | *Bu* | *Bt, Bu* | *Bt_Bu* |

| | | | |
|---|---|---|---|
| B. uniformis PUL_36 | *Bu* | *Bf, Bo, Bt, Bu* | *Bf_Bo_Bt_Bu* |
| B. uniformis PUL_37 | *Bu* | *Bu* | *Bu* |
| B. uniformis PUL_38 | *Bu* | *Bu* | *Bu* |
| B. uniformis PUL_39 | *Bu* | *Bc, Bo, Bu* | *Bc_Bo_Bu* |
| B. uniformis PUL_40 | *Bu* | *Bu* | *Bu* |
| B. uniformis PUL_41 | *Bu* | *Bu* | *Bu* |
| B. uniformis PUL_42 | *Bu* | *Bu* | *Bu* |
| B. uniformis PUL_43 | *Bu* | *Bc, Bf, Bo, Bt, Bu* | *Bc_Bf_Bo_Bt_Bu* |
| B. uniformis PUL_44 | *Bu* | *Bc, Bf, Bu* | *Bc_Bf_Bu* |
| B. uniformis PUL_45 | *Bu* | *Bu* | *Bu* |
| B. uniformis PUL_46 | *Bu* | *Bu* | *Bu* |
| B. uniformis PUL_47 | *Bu* | *Bu* | *Bu* |
| B. uniformis PUL_48 | *Bu* | *Bc, Bf, Bo, Bt, Bu* | *Bc_Bf_Bo_Bt_Bu* |
| B. uniformis PUL_49 | *Bu* | *Bc, Bo, Bu* | *Bc_Bo_Bu* |
| B. uniformis PUL_50 | *Bu* | *Bu* | *Bu* |
| B. uniformis PUL_51 | *Bu* | *Bu* | *Bu* |
| B. uniformis PUL_52 | *Bu* | *Bu* | *Bu* |
| B. uniformis PUL_53 | *Bu* | *Bu* | *Bu* |
| B. uniformis PUL_54 | *Bu* | *Bc, Bo, Bt, Bu* | *Bc_Bo_Bt_Bu* |
| B. uniformis PUL_55 | *Bu* | *Bu* | *Bu* |
