## Supplementary Table 4 for "A high-throughput microbial glycomics platform for prebiotic development"

| Supplementary Table 4. Carbohydrates examined in this study. | | | |
|---|---|---|---|
| **Name** | **Vendor** | **Cat#** | **Lot#** |
| Amylopectin (from maize) | Sigma | 10120-250G | BCBV5057 |
| Arabinan (sugar beet) | Megazyme | P-ARAB | 90901 |
| Chondroitin disaccharide (∆di-0S) sodium salt | Sigma | C3920-5MG | SLCF3976 |
| sulfate C) | Sigma | C6737-5G | 0000338962 |
| Chondroitin sulfate sodium salt (from shark cartilage) | Sigma | C4384-5G | BCCC4066 |
| D-(-)-Fructose | Sigma | F0127-500G | SLBV8796 |
| D-(-)-Ribose (≥99%) | Sigma | R7500-100G | SLBW8877 |
| D-(+)-Cellobiose, ≥99% | Sigma | 22150-10G | 102482957 |
| D-(+)-Galactose | Sigma | G0750-500G | BCBV3624 |
| D-(+)-Glucose | Sigma | G8270-1KG | SLBX1648 |
| D-(+)-Maltose monohydrate (from potato), ≥99% | Sigma | M5885-1KG | SLCC1607 |
| D-(+)-Mannose | Sigma | M6020-100G | BCBV4824 |
| D-(+)-Raffinose pentahydrate | Sigma | 83400-25G | 102539145 |
| D-(+)-Trehalose dihydrate (from Saccaromyces cerevisiae) | Sigma | T9531-10G | SLCJ3532 |
| D-(+)-Turanose | Sigma | T2754-1G | 102589729 |
| D-(+)-Xylose | Sigma | X1500-500G | WXBC6170V |
| Dermatan Sulfate | Biosynth. | YD30120 | 0000210359 |
| Dextran (from Leuconostoc spp) Mr~40,000 | Sigma | 31389-25G | BCCD2958 |
| Dextran (MW=~64000) | Megazyme | P-DEXT | 150401 |
| D-Galacturonic acid sodium salt | Sigma | T3960-25G-F | BCBX9550 |
| D-Glucuronic acid sodium salt monohydrate | Sigma | G8645-25G | SHBK4741 |
| Fructooligosaccharides (FOS) (from chicory root; DP2-8) | Megazyme | P-FOS28 | 140902a |
| Galacturonan oligosaccharides Ave DP10/15 | Elicityl | GAT114 | E1211-07 EC09 |
| Glycogen (from bovine liver-Type1X) | Sigma | G0885-5G | SLBZ0559 |
| Glycogen (from rabbit liver, ≥85%) | Sigma | G8876-5G | 1003393684 |
| Gum arabic (from acacia tree) | Sigma | G9752-500G | 102640442 |
| Heparin Disaccharide (∆UA-GLCN) | Dextra | H1012 | CCDX42-26 |
| Heparin sodium salt (from porcine intestinal mucosa) | Sigma | H4784-1G | SLCC4685 |
| Hyaluronic Acid disaccharide sodium salt | Biosynth. | OH11537 | 0000051086 |
| Hyaluronic Acid sodium salt (from Streptococcus equi) | Sigma | 53747-10G | 102437226 |
| Isolichenan polysaccharide (from Cetraria islandica) | Elicityl | GLU601 | E1308-02 EC02 |
| Isomalto Oligomers (DP4-8) | Biosynth. | OI171851 | 0000207905 |
| Isomaltose (~98%) | Sigma | I7253-500MG | 0000215448 |
| Isomaltotetraose DP4 (>95%) (from dextran) | Elicityl | GLU354-95% | EF0825-MF02 |
| Isomaltotriose DP3 (>95%) | Elicityl | GLU353-95% | EF0825-MF01 |
| Isomaltotriose, 97% | Sigma | J64186.ME | Q18K011 |
| Karaya Gum (from Sterculia tree) | Sigma | G0503-500G | 283934 |
| Kojibiose | Santa Cruz | SC-218616B | A0625 |
| L-(-)-Fucose | Sigma | F2252-25G | SLBX2465 |
| L-(+)-Arabinose | Sigma | A3256-100G | BCBW8082 |
| Laminarin (from Laminaria digitata) | Sigma | L9634-1G | 1003510678 |
| Larch Arabinogalactan | Megazyme | P-ARGAL | 20701c |
| Levan (from Timothy Grass) | Megazyme | P-LEVAN | 180620a |
| Lichenan polysaccharide (from Cetraria islandica) | Elicityl | GLU600 | E1308-02 EC01 |
| Locust bean gum (from Ceratonia siliqua seeds)-a galactomannan polysaccharide) | Sigma | G0753-500G | 274283 |
| L-Rhamnose monohydrate | Sigma | R3875-100G | BCBW9970 |
| Maltotriose hydrate, 95% | Sigma | 851493-1G | MKCJ6385 |
| Man α-2 Man | Santa Cruz | SC-256377 | A0625 |
| Man α-2 Man α-2 Man | Omicron Biochemicals | TRI-007 | H01-N0113 |
| Man α-2 Man α-3 Man | Omicron Biochemicals | TRI-008 | H02-N1813 |
| Man α-2 Man α-6 Man | Omicron Biochemicals | TRI-009 | H01-N1813 |
| Man α-3 [Man α-6] Man | Omicron Biochemicals | TRI-006 | W02-31425 |
| Man α-3 Man | Santa Cruz | SC-216644A | L1224 |
| Man α-3 Man α-6 Man | Omicron Biochemicals | TRI-010 | R01-D1313 |
| Man α-4 [Man α-6] Man | Omicron Biochemicals | TRI-011 | W02-30525/hgh5 |
| Man α-4 Man | Biosynth. | OM04762 | 000021983 |
| Man α-6 Man | Omicron Biochemicals | DIS-012 | S02-92922 |
| Man α-6 Man α-6 Man | Omicron Biochemicals | TRI-005 | B01-N1313 |
| Man β-4 Man | Biosynth. | OM02505 | 0000044718 |
| Mannan (from Saccharomyces cerevisiae) | Sigma | M7504-5G | SLBT8710 |
| Mannuronate oligosaccharides Ave DP20 | Elicityl | ALG500 | E0910-14 EC01 |
| Melibiose, ≥98% | Sigma | M5500-5G | 102505879 |
| Mucin (from porcine stomach-Type III, bound sialic acid 0.5-1.5% partially purified powder) | Sigma | M1778-100G | 0000337154 |
| N-Acetyl-D-galactosamine (~98%) | Sigma | A2795-1G | BCBW3620 |

| | | | |
|---|---|---|---|
| N-Acetyl-D-glucosamine | Sigma | A3286-100G | SLBZ73647 |
| N-Acetyl-D-lactosamine (≥98%) | Sigma | A7791-500MG | BCCJ2846 |
| Palatinose hydrate, ≥99% | Sigma | P2007-1G | 260484 |
| Pectic Galactan (Lupin) | Megazyme | P-PGALU | 20401c |
| Polygalacturonic Acid (PGA) (prepared from citrus pectin) | Megazyme | P-PGACT | 30402d |
| Pullulan | Megazyme | P-PULLN | 130701a |
| Pustulan polysaccharide (from Lasallia pustulata) | Elicityl | GLU900 | EC0467-EC01 |
| Rhamnogalacturonan I (from potato pectic fibre) | Megazyme | P-RHAM1 | 141102b |
| Stachyose hydrate | Sigma | 851787-1G | 1003634144 |
| Starch (from potato) | Sigma | S4251-2KG | BCBX0477 |
| Sucrose | Sigma | S03089-5KG | 1003278867 |
| Xanthum gum (from Xanthomonas campestris) | Sigma | G1253-100 | 1003504905 |
| Zymosan | Santa Cruz | SC-296863A | B1025 |
| β-Gentiobiose (≥85% and remainder primarily α-anomer) | Sigma | G3000-1G | 102403061 |
| κ-Carrageenan (sulfated plant polysaccharide) | Sigma | 22048-25G-F | 102601054 |
| **Biologically-derived mixtures** | | | |
| **Name** | **Manufacturer** | **Cat#** | **Extraction Metho** |
| 5021 Rodent Chow | LabDiet | N/A | ORNG |
| All-purpose flour | Gold Medal | N/A | ORNG |
| Almond Flour | Blue Diamond | N/A | ORNG |
| Avocado | N/A | N/A | ORNG |
| Bell Pepper | N/A | N/A | ORNG |
| Blueberries | Natures Partner | N/A | ORNG |
| Boar Cecal Glycans | Penn State Meats Lab | N/A | ORNG |
| Broccoli | N/A | N/A | ORNG |
| Buckwheat flour | Anthony's | N/A | ORNG |
| Bud Light | Anheuser-Busch | N/A | ProK/ORNG |
| Butternut Squash | N/A | N/A | ORNG |
| Carrots | N/A | N/A | ORNG |
| Cassava Flour | Pereg Natural Foods | N/A | ORNG |
| Cocoa Powder | Fair Trade | N/A | ORNG |
| Coconut Flour | Pereg Natural Foods | N/A | ORNG |
| Coffee Beans | Dunkin | N/A | ORNG |
| Corn Flakes | Kellogs | N/A | ORNG |
| Corn Meal Flour | Indian Head | N/A | ORNG |
| Egg Whites | N/A | N/A | ORNG |
| Egg Yolks | N/A | N/A | ORNG |
| Fig | Herman Produce | N/A | ORNG |
| Garbanzo Beans | Fair Trade | N/A | ORNG |
| Grape Juice Glycans | Fair Trade | N/A | ProK/ORNG |
| Guinness Draught Stout | Guinness | N/A | ProK/ORNG |
| Honey Crisp Apple Flesh | N/A | N/A | ORNG |
| Kiwi | N/A | N/A | ORNG |
| Lentils | Wild Harvest | N/A | ORNG |
| Mannooligosaccharides (MOS) | Alltech | N/A | ORNG |
| Oyster Mushrooms | N/A | N/A | ORNG |
| Papaya Flesh | La Carreta | N/A | ORNG |
| Plantains | N/A | N/A | ORNG |
| PMG Mucin (from porcine stomach-Type III, bound sialic acid 0.5-1.5% partially purified powder) | Sigma | N/A | ORNG |
| PMOG Mucin (from porcine stomach-Type III, bound sialic acid 0.5-1.5% partially purified powder) | Sigma | N/A | Alkaline β- |
| Portobello Mushroom | N/A | N/A | ORNG |
| Pumpkin Flesh | N/A | N/A | ORNG |
| Pumpkin Seeds | N/A | N/A | ORNG |
| Red Beets | N/A | N/A | ORNG |
| Red Wine | Woodbridge Cabernet Sauvignon | N/A | ProK/ORNG |
| Single Donor Human Milk | Innovative Research | IRHUBMKS50ML-52098 | ProK |
| Single Donor Human Milk | Innovative Research | IRHUBMKS50ML-52098 | ORNG |
| Soybeans | N/A | N/A | ORNG |
| Strawberries | Spivey Farms | N/A | ORNG |
| Sweet Peas (canned) | Delmonte | N/A | ORNG |
| Perpetual IPA | Troegs | N/A | ProK/ORNG |
| Vidalia Onion | N/A | N/A | ORNG |
| White Rice Flour | Naturevibe Botanicals | N/A | ORNG |
| White Bread | Old Tyme Classic | N/A | ORNG |
| Whole grain bread | Arnold | N/A | ORNG |
| Yeast Extract | Difco | N/A | ORNG |
