## Supplementary Table 5 for "A high-throughput microbial glycomics platform for prebiotic development"

| Supplementary Table 5. Strains and plasmids used in this study | | | | |
|---|---|---|---|---|
| **Plasmid** | **Purpose** | | | **Citation** |
| pNBU2-$P_{rpoD}$-rpiL* | Cloning ECF-class PUL sensors for over-expression and constitutive activity | | | this work |
| pNBU2-$P_{rpoD}$-rpiL*-BT3172HK | Cloning HTCS-class PUL sensor response regulator domains for constitutive activity | | | this work |
| pBolux | A promoterless plasmid containing the *Bacteroides*-optimized luciferase cassette | | | PMCID: PMC9845299 |
| pEXCHANGE | A plasmid to manipulate the Bt genome using allelic exchange | | | PMCID: PMC2563962 |
| pT7-7-6H4A | A plasmid used to express N-terminal hexahistidine tagged proteins from the T7 promoter. | | | PMCID: PMC9845299 |
| **Strain ID** | **genotype** | **plasmid** | **purpose** | **generated in** |
| ***E. coli* strains** | | | | |
| GT3664 | *λpir* | *att-1::pNBU2-ermGb-$P_{rpoD}$-rpiL*-BT1278* | a strain expressing the PUL16 sigma-factor, BT1278 | this work |
| GT3665 | *λpir* | *att-1::pNBU2-ermGb-$P_{rpoD}$-rpiL*-BT1617* | a strain expressing the PUL19 sigma-factor, BT1617 | this work |
| GT3802 | *λpir* | *att-1::pNBU2-ermGb-$P_{rpoD}$-rpiL*-BT3097** | a strain expressing the constitutively active PUL49 HTCS, BT3097* | this work |
| GT3669 | *λpir* | *att-1::pNBU2-ermGb-$P_{rpoD}$-rpiL*-BT3269* | a strain expressing the constitutively active PUL53 HTCS, BT3269* | this work |
| GT3184 | *λpir* | *att-1::pNBU2-ermGb-$P_{rpoD}$-rpiL*-BT3334** | a strain expressing the constitutively active PUL57 HTCS, BT3334* | this work |
| GT1675 | *λpir* | *pBolux* | plasmid encoding a *Bacteroides*-optimiated luciferase cassette lacking a promoter | PMCID: PMC9845299 |
| GT4073 | *λpir* | *P-PUL1* | p*Bolux* harboring the promoter identified for PUL1 | this work |
| GT4074 | *λpir* | *P-PUL2* | p*Bolux* harboring the promoter identified for PUL2 | this work |
| GT3697 | *λpir* | *P-PUL3* | p*Bolux* harboring the promoter identified for PUL3 | this work |
| BT4075 | *λpir* | *P-PUL4* | p*Bolux* harboring the promoter identified for PUL4 | this work |
| GT1772 | *λpir* | *P-PUL5* | p*Bolux* harboring the promoter identified for PUL5 | this work |
| GT1774 | *λpir* | *P-PUL6* | p*Bolux* harboring the promoter identified for PUL6 | this work |
| GT1679 | *λpir* | *P-PUL7* | p*Bolux* harboring the promoter identified for PUL7 | this work |
| GT4094 | *λpir* | *P-PUL8* | p*Bolux* harboring the promoter identified for PUL8 | this work |
| GT1969 | *λpir* | *P-PUL9* | p*Bolux* harboring the promoter identified for PUL9 | this work |
| GT1776 | *λpir* | *P-PUL10* | p*Bolux* harboring the promoter identified for PUL10 | this work |
| GT3684 | *λpir* | *P-PUL11* | p*Bolux* harboring the promoter identified for PUL11 | this work |
| GT4077 | *λpir* | *P-PUL12* | p*Bolux* harboring the promoter identified for PUL12 | this work |
| GT3951 | *λpir* | *P-PUL13* | p*Bolux* harboring the promoter identified for PUL13 | this work |
| GT1781 | *λpir* | *P-PUL14a* | p*Bolux* harboring the promoter identified for PUL14a | this work |
| GT1782 | *λpir* | *P-PUL14b* | p*Bolux* harboring the promoter identified for PUL14b | this work |
| GT4116 | *λpir* | *P-PUL15* | p*Bolux* harboring the promoter identified for PUL15 | this work |
| GT3685 | *λpir* | *P-PUL16* | p*Bolux* harboring the promoter identified for PUL16 | this work |
| GT1785 | *λpir* | *P-PUL17* | p*Bolux* harboring the promoter identified for PUL17 | this work |
| GT1786 | *λpir* | *P-PUL18* | p*Bolux* harboring the promoter identified for PUL18 | this work |
| GT3686 | *λpir* | *P-PUL19* | p*Bolux* harboring the promoter identified for PUL19 | this work |
| GT1788 | *λpir* | *P-PUL20* | p*Bolux* harboring the promoter identified for PUL20 | this work |
| GT4095 | *λpir* | *P-PUL21* | p*Bolux* harboring the promoter identified for PUL21 | this work |
| GT1790 | *λpir* | *P-PUL22* | p*Bolux* harboring the promoter identified for PUL22 | PMCID: PMC9845299 |
| GT4096 | *λpir* | *P-PUL23* | p*Bolux* harboring the promoter identified for PUL23 | this work |
| GT3905 | *λpir* | *P-PUL24* | p*Bolux* harboring the promoter identified for PUL24 | this work |
| GT1794 | *λpir* | *P-PUL25* | p*Bolux* harboring the promoter identified for PUL25 | this work |
| GT1795 | *λpir* | *P-PUL26* | p*Bolux* harboring the promoter identified for PUL26 | this work |
| GT3688 | *λpir* | *P-PUL27* | p*Bolux* harboring the promoter identified for PUL27 | this work |
| GT3687 | *λpir* | *P-PUL28* | p*Bolux* harboring the promoter identified for PUL28 | this work |
| GT3690 | *λpir* | *P-PUL29* | p*Bolux* harboring the promoter identified for PUL29 | this work |
| GT1800 | *λpir* | *P-PUL30* | p*Bolux* harboring the promoter identified for PUL30 | this work |
| GT1801 | *λpir* | *P-PUL31* | p*Bolux* harboring the promoter identified for PUL31 | this work |
| GT1803 | *λpir* | *P-PUL32* | p*Bolux* harboring the promoter identified for PUL32 | this work |
| GT3906 | *λpir* | *P-PUL33* | p*Bolux* harboring the promoter identified for PUL33 | this work |
| GT4132 | *λpir* | *P-PUL34* | p*Bolux* harboring the promoter identified for PUL34 | this work |
| GT2355 | *λpir* | *P-PUL35* | p*Bolux* harboring the promoter identified for PUL35 | this work |
| GT1807 | *λpir* | *P-PUL36* | p*Bolux* harboring the promoter identified for PUL36 | this work |
| GT2358 | *λpir* | *P-PUL37* | p*Bolux* harboring the promoter identified for PUL37 | this work |
| GT1809 | *λpir* | *P-PUL38* | p*Bolux* harboring the promoter identified for PUL38 | PMCID: PMC9845299 |
| GT4133 | *λpir* | *P-PUL39* | p*Bolux* harboring the promoter identified for PUL39 | this work |
| GT4099 | *λpir* | *P-PUL40* | p*Bolux* harboring the promoter identified for PUL40 | this work |
| GT4100 | *λpir* | *P-PUL41* | p*Bolux* harboring the promoter identified for PUL41 | this work |
| GT1814 | *λpir* | *P-PUL42* | p*Bolux* harboring the promoter identified for PUL42 | this work |
| GT1815 | *λpir* | *P-PUL43* | p*Bolux* harboring the promoter identified for PUL43 | this work |
| GT3955 | *λpir* | *P-PUL44* | p*Bolux* harboring the promoter identified for PUL44 | this work |
| GT2348 | *λpir* | *P-PUL45* | p*Bolux* harboring the promoter identified for PUL45 | this work |
| GT4101 | *λpir* | *P-PUL46* | p*Bolux* harboring the promoter identified for PUL46 | this work |
| GT1819 | *λpir* | *P-PUL47* | p*Bolux* harboring the promoter identified for PUL47 | this work |
| GT1820 | *λpir* | *P-PUL48* | p*Bolux* harboring the promoter identified for PUL48 | this work |
| GT1821 | *λpir* | *P-PUL49* | p*Bolux* harboring the promoter identified for PUL49 | this work |
| GT1822 | *λpir* | *P-PUL50* | p*Bolux* harboring the promoter identified for PUL50 | this work |
| GT1823 | *λpir* | *P-PUL51* | p*Bolux* harboring the promoter identified for PUL51 | this work |
| GT1824 | *λpir* | *P-PUL52a* | p*Bolux* harboring the promoter identified for PUL52a | this work |
| GT1825 | *λpir* | *P-PUL52b* | p*Bolux* harboring the promoter identified for PUL52b | this work |
| GT1826 | *λpir* | *P-PUL53* | p*Bolux* harboring the promoter identified for PUL53 | this work |
| GT1827 | *λpir* | *P-PUL54* | p*Bolux* harboring the promoter identified for PUL54 | this work |
| GT1829 | *λpir* | *P-PUL55* | p*Bolux* harboring the promoter identified for PUL55 | this work |
| GT1830 | *λpir* | *P-PUL56* | p*Bolux* harboring the promoter identified for PUL56 | this work |
| GT1831 | *λpir* | *P-PUL57* | p*Bolux* harboring the promoter identified for PUL57 | PMCID: PMC9845299 |
| GT1832 | *λpir* | *P-PUL58* | p*Bolux* harboring the promoter identified for PUL58 | this work |
| GT1833 | *λpir* | *P-PUL59* | p*Bolux* harboring the promoter identified for PUL59 | this work |
| GT3692 | *λpir* | *P-PUL60* | p*Bolux* harboring the promoter identified for PUL60 | this work |
| GT1837 | *λpir* | *P-PUL61* | p*Bolux* harboring the promoter identified for PUL61 | this work |
| GT1838 | *λpir* | *P-PUL62* | p*Bolux* harboring the promoter identified for PUL62 | this work |
| GT1839 | *λpir* | *P-PUL64* | p*Bolux* harboring the promoter identified for PUL64 | this work |
| GT1843 | *λpir* | *P-PUL65* | p*Bolux* harboring the promoter identified for PUL65 | this work |
| GT4078 | *λpir* | *P-PUL66* | p*Bolux* harboring the promoter identified for PUL66 | this work |
| GT3949 | *λpir* | *P-PUL67* | p*Bolux* harboring the promoter identified for PUL67 | this work |
| GT1840 | *λpir* | *P-PUL68* | p*Bolux* harboring the promoter identified for PUL68 | this work |
| GT1841 | *λpir* | *P-PUL69* | p*Bolux* harboring the promoter identified for PUL69 | this work |
| GT4135 | *λpir* | *P-PUL70* | p*Bolux* harboring the promoter identified for PUL70 | this work |
| GT3950 | *λpir* | *P-PUL71* | p*Bolux* harboring the promoter identified for PUL71 | this work |
| GT1846 | *λpir* | *P-PUL72* | p*Bolux* harboring the promoter identified for PUL72 | this work |
| GT4105 | *λpir* | *P-PUL73* | p*Bolux* harboring the promoter identified for PUL73 | this work |
| GT4136 | *λpir* | *P-PUL74a* | p*Bolux* harboring the promoter identified for PUL74a | this work |
| GT1849 | *λpir* | *P-PUL74b* | p*Bolux* harboring the promoter identified for PUL74b | this work |
| GT1850 | *λpir* | *P-PUL75a* | p*Bolux* harboring the promoter identified for PUL75a | this work |
| GT1851 | *λpir* | *P-PUL75b* | p*Bolux* harboring the promoter identified for PUL75b | this work |
| GT2102 | *λpir* | *P-PUL76* | p*Bolux* harboring the promoter identified for PUL76 | this work |
| GT1853 | *λpir* | *P-PUL77* | p*Bolux* harboring the promoter identified for PUL77 | this work |
| GT1855 | *λpir* | *P-PUL78* | p*Bolux* harboring the promoter identified for PUL78 | this work |
| GT1856 | *λpir* | *P-PUL79* | p*Bolux* harboring the promoter identified for PUL79 | this work |
| GT1857 | *λpir* | *P-PUL80* | p*Bolux* harboring the promoter identified for PUL80 | this work |
| GT3694 | *λpir* | *P-PUL81* | p*Bolux* harboring the promoter identified for PUL81 | this work |
| GT2639 | *λpir* | *P-PUL82* | p*Bolux* harboring the promoter identified for PUL82 | this work |
| GT1860 | *λpir* | *P-PUL83* | p*Bolux* harboring the promoter identified for PUL83 | this work |
| GT2350 | *λpir* | *P-PUL84* | p*Bolux* harboring the promoter identified for PUL84 | this work |
| GT1862 | *λpir* | *P-PUL85* | p*Bolux* harboring the promoter identified for PUL85 | this work |

| | | | | |
|---|---|---|---|---|
| GT1863 | *λpir* | *P-PUL86* | p*Bolux* harboring the promoter identified for PUL86 | this work |
| GT3695 | *λpir* | *P-PUL87* | p*Bolux* harboring the promoter identified for PUL87 | this work |
| GT3696 | *λpir* | *P-PUL88* | p*Bolux* harboring the promoter identified for PUL88 | this work |
| GT1678 | *λpir* | *P-rpoD* ( *Bt*) | p*Bolux* harboring the *B. thetaiotaomicron rpoD* promoter | this work |
| GT3425 | *λpir* | *P-rpoD (Bf)* | p*Bolux* harboring the *B. fragilis rpoD* promoter | this work |
| GT3427 | *λpir* | *P-rpoD (Bo)* | p*Bolux* harboring the *B. ovatus rpoD* promoter | this work |
| GT5387 | *λpir* | *P-rpoD (Bc)* | p*Bolux* harboring the *B. cellulosilyticus rpoD* promoter | this work |
| GT5359 | *λpir* | *P-rpoD (Bu)* | p*Bolux* harboring the *B. uniformis rpoD* promoter | this work |
| GT3563 | *λpir* | *pT7-7-6H4A-BT3330* | pT7-7::6H4A expressing SGBP$^{PUL57}$ | PMCID: PMC9845299 |
| GT4463 | *λpir* | *pT7-7-6H4A-BT3961* | pT7-7::6H4A expressing SGBP$^{PUL71}$ | this work |
| GT4414 | *λpir* | *pT7-7-6H4A-BT3313* | pT7-7::6H4A expressing SGBP$^{PUL56}$ | this work |
| GT4967 | *λpir* | *pT7-7-6H4A-BT3312* | pT7-7::6H4A expressing GH20$^{PUL56}$ | this work |
| GT4968 | *λpir* | *pT7-7-6H4A-BT3962* | pT7-7::6H4A expressing GH92-1$^{PUL71}$ | this work |
| GT4969 | *λpir* | *pT7-7-6H4A-BT3963* | pT7-7::6H4A expressing GH92-2$^{PUL71}$ | this work |
| GT4980 | *λpir* | *pT7-7-6H4A-BT3965* | pT7-7::6H4A expressing GH92-3$^{PUL71}$ | this work |
| GT5421 | *λpir* | *pEXCHANGE-ΔBT2160* | Plasmid used to generate a *sensor*$^{PUL58}$-deficient strain | this work |
| GT29 | *λpir* | *pEXCHANGE-ΔBT2628* | Plasmid used to generate a *sensor*$^{PUL36}$-deficient strain | this work |
| GT87 | *λpir* | *pEXCHANGE-ΔBT3786* | Plasmid used to generate a *sensor*$^{PUL68}$-deficient strain | this work |
| GT4558 | *λpir* | *pEXCHANGE-ΔBT3853* | Plasmid used to generate a *sensor*$^{PUL69}$-deficient strain | this work |
| GT4151 | *λpir* | *pEXCHANGE-ΔBT3957* | Plasmid used to generate a *sensor*$^{PUL71}$-deficient strain | this work |
| GT5505 | *λpir* | *pEXCHANGE-ΔBT3957-BT3965* | Plasmid used to generate a PUL71-deficient strain | this work |
| GT4485 | *λpir* | *pEXCHANGE-ΔBT3309* | Plasmid used to generate a *sensor*$^{PUL56}$-deficient strain | this work |
| | | ***B. thetaiotaomicron* strains** | | |
| GT23 | *Δtdk* | | a *tdk*-deficient strain used as *wild-type* for all experiments | PMCID: PMC3545799 |
| GT2111 | *Δtdk* | *att-1::pNBU2-ermGb* | a strain harboring an empty pNBU2 vector | this work |
| GT3664 | *Δtdk* | *att-1::pNBU2-ermGb::P*$_{rpoD}$*-rpiL*-BT1278* | a strain expressing the PUL16 sigma-factor, BT1278 | this work |
| GT3665 | *Δtdk* | *att-1::pNBU2-ermGb::P*$_{rpoD}$*-rpiL*-BT1617* | a strain expressing the PUL19 sigma-factor, BT1617 | this work |
| GT3802 | *Δtdk* | *att-1::pNBU2-ermGb::P*$_{rpoD}$*-rpiL*-BT3097** | a strain expressing the constitutively active PUL49 HTCS, BT3097* | this work |
| GT3669 | *Δtdk* | *att-1::pNBU2-ermGb::P*$_{rpoD}$*-rpiL*-BT3269* | a strain expressing the constitutively active PUL53 HTCS, BT3269* | this work |
| GT3184 | *Δtdk* | *att-1::pNBU2-ermGb::P*$_{rpoD}$*-rpiL*-BT3334** | a strain expressing the constitutively active PUL57 HTCS, BT3334* | this work |
| GT1867 | *Δtdk* | p*Bolux* | *wild-type Bt* harboring p*Bolux* | this work |
| GT4079 | *Δtdk* | *P-PUL1* | *wild-type Bt* harboring P-*PUL1* | this work |
| GT4080 | *Δtdk* | *P-PUL2* | *wild-type Bt* harboring P-*PUL2* | this work |
| GT3787 | *Δtdk* | *P-PUL3* | *wild-type Bt* harboring P-*PUL3* | this work |
| GT4081 | *Δtdk* | *P-PUL4* | *wild-type Bt* harboring P-*PUL4* | this work |
| GT1873 | *Δtdk* | *P-PUL5* | *wild-type Bt* harboring P-*PUL5* | this work |
| GT1875 | *Δtdk* | *P-PUL6* | *wild-type Bt* harboring P-*PUL6* | this work |
| GT1876 | *Δtdk* | *P-PUL7* | *wild-type Bt* harboring P-*PUL7* | this work |
| GT4117 | *Δtdk* | *P-PUL8* | *wild-type Bt* harboring P-*PUL8* | this work |
| GT1878 | *Δtdk* | *P-PUL9* | *wild-type Bt* harboring P-*PUL9* | this work |
| GT1879 | *Δtdk* | *P-PUL10* | *wild-type Bt* harboring P-*PUL10* | this work |
| GT3779 | *Δtdk* | *P-PUL11* | *wild-type Bt* harboring P-*PUL11* | this work |
| GT4083 | *Δtdk* | *P-PUL12* | *wild-type Bt* harboring P-*PUL12* | this work |
| GT3972 | *Δtdk* | *P-PUL13* | *wild-type Bt* harboring P-*PUL13* | this work |
| GT1884 | *Δtdk* | *P-PUL14a* | *wild-type Bt* harboring P-*PUL14* a | this work |
| GT1885 | *Δtdk* | *P-PUL14b* | *wild-type Bt* harboring P-*PUL14* b | this work |
| GT4131 | *Δtdk* | *P-PUL15* | *wild-type Bt* harboring P-*PUL15* | this work |
| GT3780 | *Δtdk* | *P-PUL16* | *wild-type Bt* harboring P-*PUL16* | this work |
| GT1888 | *Δtdk* | *P-PUL17* | *wild-type Bt* harboring P-*PUL17* | this work |
| GT1889 | *Δtdk* | *P-PUL18* | *wild-type Bt* harboring P-*PUL18* | this work |
| GT3781 | *Δtdk* | *P-PUL19* | *wild-type Bt* harboring P-*PUL19* | this work |
| GT1891 | *Δtdk* | *P-PUL20* | *wild-type Bt* harboring P-*PUL20* | this work |
| GT4118 | *Δtdk* | *P-PUL21* | *wild-type Bt* harboring P-*PUL21* | this work |
| GT1893 | *Δtdk* | *P-PUL22* | *wild-type Bt* harboring P-*PUL22* | PMCID: PMC9845299 |
| GT4119 | *Δtdk* | *P-PUL23* | *wild-type Bt* harboring P-*PUL23* | this work |
| GT3908 | *Δtdk* | *P-PUL24* | *wild-type Bt* harboring P-*PUL24* | this work |
| GT1897 | *Δtdk* | *P-PUL25* | *wild-type Bt* harboring P-*PUL25* | this work |
| GT1898 | *Δtdk* | *P-PUL26* | *wild-type Bt* harboring P-*PUL26* | this work |
| GT3783 | *Δtdk* | *P-PUL27* | *wild-type Bt* harboring P-*PUL27* | this work |
| GT3782 | *Δtdk* | *P-PUL28* | *wild-type Bt* harboring P-*PUL28* | this work |
| GT3878 | *Δtdk* | *P-PUL29* | *wild-type Bt* harboring P-*PUL29* | this work |
| GT1903 | *Δtdk* | *P-PUL30* | *wild-type Bt* harboring P-*PUL30* | this work |
| GT1904 | *Δtdk* | *P-PUL31* | *wild-type Bt* harboring P-*PUL31* | this work |
| GT1906 | *Δtdk* | *P-PUL32* | *wild-type Bt* harboring P-*PUL32* | this work |
| GT3911 | *Δtdk* | *P-PUL33* | *wild-type Bt* harboring P-*PUL33* | this work |
| GT4137 | *Δtdk* | *P-PUL34* | *wild-type Bt* harboring P-*PUL34* | this work |
| GT2372 | *Δtdk* | *P-PUL35* | *wild-type Bt* harboring P-*PUL35* | this work |
| GT1910 | *Δtdk* | *P-PUL36* | *wild-type Bt* harboring P-*PUL36* | this work |
| GT2375 | *Δtdk* | *P-PUL37* | *wild-type Bt* harboring P-*PUL37* | this work |
| GT1912 | *Δtdk* | *P-PUL38* | *wild-type Bt* harboring P-*PUL38* | PMCID: PMC9845299 |
| GT4138 | *Δtdk* | *P-PUL39* | *wild-type Bt* harboring P-*PUL39* | this work |
| GT4122 | *Δtdk* | *P-PUL40* | *wild-type Bt* harboring P-*PUL40* | this work |
| GT4123 | *Δtdk* | *P-PUL41* | *wild-type Bt* harboring P-*PUL41* | this work |
| GT1917 | *Δtdk* | *P-PUL42* | *wild-type Bt* harboring P-*PUL42* | this work |
| GT1918 | *Δtdk* | *P-PUL43* | *wild-type Bt* harboring P-*PUL43* | this work |
| GT3976 | *Δtdk* | *P-PUL44* | *wild-type Bt* harboring P-*PUL44* | this work |
| GT2365 | *Δtdk* | *P-PUL45* | *wild-type Bt* harboring P-*PUL45* | this work |
| GT4124 | *Δtdk* | *P-PUL46* | *wild-type Bt* harboring P-*PUL46* | this work |
| GT1922 | *Δtdk* | *P-PUL47* | *wild-type Bt* harboring P-*PUL47* | this work |
| GT1923 | *Δtdk* | *P-PUL48* | *wild-type Bt* harboring P-*PUL48* | this work |
| GT1924 | *Δtdk* | *P-PUL49* | *wild-type Bt* harboring P-*PUL49* | this work |
| GT1925 | *Δtdk* | *P-PUL50* | *wild-type Bt* harboring P-*PUL50* | this work |
| GT1926 | *Δtdk* | *P-PUL51* | *wild-type Bt* harboring P-*PUL51* | this work |
| GT1927 | *Δtdk* | *P-PUL52a* | *wild-type Bt* harboring P-*PUL52* a | this work |
| GT1928 | *Δtdk* | *P-PUL52b* | *wild-type Bt* harboring P-*PUL52b* | this work |
| GT1929 | *Δtdk* | *P-PUL53* | *wild-type Bt* harboring P-*PUL53* | this work |
| GT1930 | *Δtdk* | *P-PUL54* | *wild-type Bt* harboring P-*PUL54* | this work |
| GT1932 | *Δtdk* | *P-PUL55* | *wild-type Bt* harboring P-*PUL55* | this work |
| GT1933 | *Δtdk* | *P-PUL56* | *wild-type Bt* harboring P-*PUL56* | this work |
| GT1934 | *Δtdk* | *P-PUL57* | *wild-type Bt* harboring P-*PUL57* | PMCID: PMC9845299 |
| GT1935 | *Δtdk* | *P-PUL58* | *wild-type Bt* harboring P-*PUL58* | this work |
| GT1936 | *Δtdk* | *P-PUL59* | *wild-type Bt* harboring P-*PUL59* | this work |
| GT3784 | *Δtdk* | *P-PUL60* | *wild-type Bt* harboring P-*PUL60* | this work |
| GT1940 | *Δtdk* | *P-PUL61* | *wild-type Bt* harboring P-*PUL61* | this work |
| GT1941 | *Δtdk* | *P-PUL62* | *wild-type Bt* harboring P-*PUL62* | this work |
| GT1942 | *Δtdk* | *P-PUL64* | *wild-type Bt* harboring P-*PUL64* | this work |
| GT1946 | *Δtdk* | *P-PUL65* | *wild-type Bt* harboring P-*PUL65* | this work |
| GT4084 | *Δtdk* | *P-PUL66* | *wild-type Bt* harboring P-*PUL66* | this work |
| GT3970 | *Δtdk* | *P-PUL67* | *wild-type Bt* harboring P-*PUL67* | this work |
| GT1943 | *Δtdk* | *P-PUL68* | *wild-type Bt* harboring P-*PUL68* | this work |
| GT1944 | *Δtdk* | *P-PUL69* | *wild-type Bt* harboring P-*PUL69* | this work |

| | | | | |
|---|---|---|---|---|
| GT4140 | *Δtdk* | *P-PUL70* | *wild-type Bt* harboring P-*PUL70* | this work |
| GT3971 | *Δtdk* | *P-PUL71* | *wild-type Bt* harboring P-*PUL71* | this work |
| GT1949 | *Δtdk* | *P-PUL72* | *wild-type Bt* harboring P-*PUL72* | this work |
| GT4128 | *Δtdk* | *P-PUL73* | *wild-type Bt* harboring P-*PUL73* | this work |
| GT4141 | *Δtdk* | *P-PUL74a* | *wild-type Bt* harboring P-*PUL74* a | this work |
| GT1952 | *Δtdk* | *P-PUL74b* | *wild-type Bt* harboring P-*PUL74b* | this work |
| GT1953 | *Δtdk* | *P-PUL75a* | *wild-type Bt* harboring P-*PUL75* a | this work |
| GT1954 | *Δtdk* | *P-PUL75b* | *wild-type Bt* harboring P-*PUL75b* | this work |
| GT2134 | *Δtdk* | *P-PUL76* | *wild-type Bt* harboring P-*PUL76* | this work |
| GT1956 | *Δtdk* | *P-PUL77* | *wild-type Bt* harboring P-*PUL77* | this work |
| GT1958 | *Δtdk* | *P-PUL78* | *wild-type Bt* harboring P-*PUL78* | this work |
| GT1959 | *Δtdk* | *P-PUL79* | *wild-type Bt* harboring P-*PUL79* | this work |
| GT1960 | *Δtdk* | *P-PUL80* | *wild-type Bt* harboring P-*PUL80* | this work |
| GT3785 | *Δtdk* | *P-PUL81* | *wild-type Bt* harboring P-*PUL81* | this work |
| GT2850 | *Δtdk* | *P-PUL82* | *wild-type Bt* harboring P-*PUL82* | this work |
| GT1963 | *Δtdk* | *P-PUL83* | *wild-type Bt* harboring P-*PUL83* | this work |
| GT2367 | *Δtdk* | *P-PUL84* | *wild-type Bt* harboring P-*PUL84* | this work |
| GT1965 | *Δtdk* | *P-PUL85* | *wild-type Bt* harboring P-*PUL85* | this work |
| GT1966 | *Δtdk* | *P-PUL86* | *wild-type Bt* harboring P-*PUL86* | this work |
| GT3786 | *Δtdk* | *P-PUL87* | *wild-type Bt* harboring P-*PUL87* | this work |
| GT3879 | *Δtdk* | *P-PUL88* | *wild-type Bt* harboring P-*PUL88* | this work |
| GT3432 | | *pBolux* | *Bt type strain* harboring corresponding p*Bolux* | PMCID: PMC9845299 |
| GT3433 | *VPI-5482* | *P-rpoD* | *Bt type strain* harboring corresponding P-*rpoD* | PMCID: PMC9845299 |
| GT5607 | | *pBolux* | A *sensor*$^{PUL58}$-deficient strain harboring p*Bolux* | this work |
| GT5608 | | P-*PUL58* | A *sensor*$^{PUL58}$-deficient strain harboring P-*PUL58* | this work |
| GT5773 | *Δtdk ΔBT2160* | P-*PUL46* | | this work |
| GT2619 | | *pBolux* | A *sensor*$^{PUL22}$*-deficient* strain harboring p*Bolux* | PMCID: PMC9845299 |
| GT2620 | | P-*PUL22* | A *sensor*$^{PUL22}$*-deficient* strain harboring P-*PUL22* | PMCID: PMC9845299 |
| GT3915 | *Δtdk ΔBT1754* | P-*PUL37* | A *sensor*$^{PUL22}$*-deficient* strain harboring P-*PUL37* | this work |
| GT5610 | | *pBolux* | A *sensor*$^{PUL37}$*-deficient* strain harboring p*Bolux* | this work |
| GT5611 | | P-*PUL22* | A *sensor*$^{PUL37}$*-deficient* strain harboring P-*PUL22* | this work |
| GT5612 | *Δtdk ΔBT2802* | P-*PUL37* | A *sensor*$^{PUL37}$*-deficient* strain harboring P-*PUL37* | this work |
| GT104 | *Δtdk ΔBT2628 ΔBT3786* | | A strain lacking the PUL36 and PUL68 sensors | this work |
| GT4572 | *Δtdk ΔBT3853* | | A strain lacking the PUL69 sensors | this work |
| GT5542 | *Δtdk ΔBT2628 ΔBT3786 ΔBT3853* | | A strain lacking the PUL36, PUL68, and PUL69 sensors | this work |
| GT4487 | *Δtdk ΔBT2628 ΔBT3786* | *pBolux* | A strain lacking the PUL36 and PUL68 sensors harboring p*Bolux* | this work |
| GT4488 | *Δtdk ΔBT2628 ΔBT3786* | P-PUL36 | A strain lacking the PUL36 and PUL68 sensors harboring P-PUL36 | this work |
| GT4490 | *Δtdk ΔBT2628 ΔBT3786* | P-PUL68 | A strain lacking the PUL36 and PUL68 sensors harboring P-PUL68 | this work |
| GT4592 | *Δtdk ΔBT3853* | *pBolux* | A *sensor*$^{PUL69}$*-deficient* strain harboring p*Bolux* | this work |
| GT4596 | *Δtdk ΔBT3853* | P-PUL69 | A *sensor*$^{PUL69}$*-deficient* strain harboring P-*PUL69* | this work |
| GT2615 | *Δtdk ΔBT0366* | *pBolux* | A *sensor*$^{PUL7}$-deficient strain harboring p*Bolux* | this work |
| GT5869 | *Δtdk ΔBT0366* | *P-PUL5* | A *sensor*$^{PUL7}$-deficient strain harboring P-*PUL5* | this work |
| GT5870 | *Δtdk ΔBT0366* | *P-PUL7* | A *sensor*$^{PUL7}$-deficient strain harboring P-*PUL7* | this work |
| GT5880 | *Δtdk ΔBT0366* | *P-PUL13* | A *sensor*$^{PUL7}$-deficient strain harboring P-*PUL13* | this work |
| GT5871 | *Δtdk ΔBT0366* | *P-PUL22* | A *sensor*$^{PUL7}$-deficient strain harboring P-*PUL22* | this work |
| GT5872 | *Δtdk ΔBT0366* | *P-PUL47* | A *sensor*$^{PUL7}$-deficient strain harboring P-*PUL47* | this work |
| GT5873 | *Δtdk ΔBT0366* | *P-PUL48* | A *sensor*$^{PUL7}$-deficient strain harboring P-*PUL48* | this work |
| GT5874 | *Δtdk ΔBT0366* | *P-PUL65* | A *sensor*$^{PUL7}$-deficient strain harboring P-*PUL65* | this work |
| GT5875 | *Δtdk ΔBT0366* | *P-PUL77* | A *sensor*$^{PUL7}$-deficient strain harboring P-*PUL77* | this work |
| GT5876 | *Δtdk ΔBT0366* | *P-PUL86* | A *sensor*$^{PUL7}$-deficient strain harboring P-*PUL86* | this work |
| GT4253 | *Δtdk ΔBT3957* | | A strain lacking the PUL71 sensor | this work |
| GT5596 | *Δtdk ΔBT2628 ΔBT3786 ΔBT3853 ΔBT3957* | | A strain lacking the PUL36, PUL68, PUL69, and PUL71 sensors | this work |
| GT5599 | *Δtdk ΔBT3957-BT3965* | | A strain lacking the entire PUL57 | this work |
| GT4501 | *Δtdk ΔBT3309* | | A strain lacking the PUL56 sensor | this work |
| | | ***B. fragilis* strains** | | |
| GT3428 | *NCTC 9343* | *pBolux* | the *Bf* type strain harboring p*Bolux* | this work |
| GT3429 | *NCTC 9343* | *P-rpoD (Bf)* | the *Bf* type strain harboring the corresponding P-*rpoD* | this work |
| | | ***B. ovatus* strains** | | |
| GT3489 | *ATCC 8483* | *pBolux* | the *Bo* type strain harboring p*Bolux* | PMCID: PMC9845299 |
| GT3490 | *ATCC 8483* | *P-rpoD (Bo)* | the *Bo* type strain harboring the corresponding P-*rpoD* | PMCID: PMC9845299 |
| | | ***B. uniformis* strains** | | |
| GT5191 | *8492* | *pBolux* | the *Bu* type strain harboring p*Bolux* | this work |
| GT5383 | *8492* | *P-rpoD (Bu)* | the *Bu* type strain harboring the corresponding P-*rpoD* | this work |
| | | ***B. cellulosilyticus* strains** | | |
| GT4559 | *DSM 14838* | *pBolux* | the *Bc* type strain harboring p*Bolux* | this work |
| GT5453 | *DSM 14838* | *P-rpoD (Bce)* | the *Bc* type strain harboring the corresponding P-*rpoD* | this work |
