## Supplementary Table 6 for "A high-throughput microbial glycomics platform for prebiotic development"

| Supplementary Table 6. Oligonucelotides used in this study | | |
|---|---|---|
| **Constitutively-active sensors** | | |
| **Sensor** | **Primer direction** | **Primer Sequence** |
| PUL16 (BT1278) | forward | tatttatttaattaaacgaatccatgaccccaaaccaagacg |
| | reverse | aagataggcaattagtcgacttaagggaacagcaaaaaaaatgca |
| PUL19 (BT1617) | forward | tatttatttaattaaacgaatccatggaaacctttgacgaaaaacaac |
| | reverse | aagataggcaattagtcgacttagaaaaatagaataaaggttatgaataatatttttcgg |
| PUL49 (BT3269*) | forward | tatttatttaattaaacgaatccatgagcgataaagatgatttgatagtagc |
| | reverse | aagataggcaattagtcgacctaataaaatataagagcaagaagtacagaataactgg |
| PUL 53 (BT3097*) | forward | gtatctgaacgaactgcagcggatgaaagaaacgatcgaact |
| | reverse | aagataggcaattagtcgacttacagattcagttgtttgataaattcagt |
| PUL57 (BT3334*) | forward | gtatctgaacgaactgataatcgttccaaccgtagaaatactgc |
| | reverse | agataggcaattagtcgacttattgatccgtttcttcgttcccttc |
| **Reporter plasmid construction** | | |
| **P-*PUL*** | **Primer direction** | **Primer Sequence** |
| 1 | forward | agctcggtacccggggatccaggacgaaaatgtaactttgcca |
| | reverse | atgcgggagtgactagtacttactatttttctcgctgcaaaattcca |
| 2 | forward | gctcggtacccggggatcccttccaatgatactgaagagaaaatcattgctg |
| | reverse | atgcgggagtgactagtaacattaaatttgaggtttaaaaatagtattaaattc |
| 3 | forward | agctcggtacccggggatccttggaacaattgactccccg |
| | reverse | atgcgggagtgactagtcatatattcaaataagggtgctgatttctcc |
| 4 | forward | agctcggtacccggggatcccgtgtggcacggctgcagct |
| | reverse | aaatgcgggagtgactagtctttgatacttgaatttaaattaatgaattgatttatttttacagatttatctac |
| 5 | forward | gctcggtacccggggatccgtctaagtcttctctttataggaaaatgaaattagc |
| | reverse | aaatgcgggagtgactagtactattagatttataaagttatttagaccaaataagtattagttgtgc |
| 6 | forward | gctcggtacccggggatccttcaatttcaataataacttttgtgtgttagagtccct |
| | reverse | aaatgcgggagtgactagtaatactgctcttttaaggttaataaaaattgtctgtgtg |
| 7 | forward | gctcggtacccggggatcctcaatgtgacaccaagcgcactg |
| | reverse | aaatgcgggagtgactagtaagtagtaacggccatttccttgtcttc |
| 8 | forward | agctcggtacccggggatccaagtggattgacgatgcacg |
| | reverse | atgcgggagtgactagtgtttactagtgataataaacagaatgtcgt |
| 9 | forward | gctcggtacccggggatccaaaataggaatttgcgtctgatgtacaaaaacg |
| | reverse | aaatgcgggagtgactagtccccgaaactccgtaatagacca |
| 10 | forward | gctcggtacccggggatccgctatggctacggatacggttacg |
| | reverse | aaatgcgggagtgactagtaattatcatttttaagggataatacaaatataacaaaaaaaacggg |
| 11 | forward | gctcggtacccggggatcctattgcccatcgtctggaacgt |
| | reverse | aaatgcgggagtgactagtttttatcactattaaaggttaacatatagttatttatcgtagctgc |
| 12 | forward | agctcggtacccggggatccgaatattgcaaaggtaatttttaaaattgtttttccc |
| | reverse | aaatgcgggagtgactagtaattttgattttaaaataataactagtttaccggttttgtttgt |
| 13 | forward | agctcggtacccggggatccctcttcctttcaatgagcggc |
| | reverse | aaatgcgggagtgactagtagattaaattttaatttatataaaatgatatattacatcatcagacatctattcacat |
| 14a | forward | gctcggtacccggggatcctggtgtttcctttaaaaccaaaatgcc |
| | reverse | aaatgcgggagtgactagttgtgacttttaggggttgggc |
| 14b | forward | gctcggtacccggggatcctagtttttcagaactacttaatgctctatttatcaaatgattatg |
| | reverse | aaatgcgggagtgactagtcttattctttttaattagaaatagtttttaggtagttaacaaacatgtg |
| 15 | forward | agctcggtacccggggatccattaaggggggggctatgctg |
| | reverse | aaatgcgggagtgactagtcttgattcgttaaatctctgtagattccccag |
| 16 | forward | gctcggtacccggggatccagaccaggcaccactatctgag |
| | reverse | aaatgcgggagtgactagttttcttagtattaggttattaactaaattatttcacggtaattgaatag |
| 17 | forward | gctcggtacccggggatcctcttaaaagcgcttggtttggcc |
| | reverse | aaatgcgggagtgactagtaattttgtggcttaggatttatttgcttgttctaac |
| 18 | forward | gctcggtacccggggatccaaggggaaagtcgaaaggtgg |
| | reverse | aaatgcgggagtgactagttaagtccttttacttttaattattattaaaaacagccgca |
| 19 | forward | gctcggtacccggggatcccatacacatcatcgtccctccgc |
| | reverse | aaatgcgggagtgactagtacttttgtgagttaatcattaatactaatttaagtgtacacaac |
| 20 | forward | gctcggtacccggggatccgggcggaagagtttttaagagagaatatag |
| | reverse | aaatgcgggagtgactagtaagtttattaattaagattaccttttatataagtagcgaagtctttcc |
| 21 | forward | agctcggtacccggggatccgtactatgcagagagttacg |
| | reverse | aaatgcgggagtgactagttcgttttaatttaaagttaatattaattttcctttgtttagtaaaatacaatt |
| 22 | forward | gctcggtacccggggatcctatcattcagttttctgttggttactttgagtga |
| | reverse | aaatgcgggagtgactagttagtttaatgttattaatttaaaagtacgaattttctcttttcgatg |
| 23 | forward | agctcggtacccggggatccgcttaattttcgcttaattgattattaa |
| | reverse | aaatgcgggagtgactagtcgtgattttcaaataatttttcacggtctctttttttatattatac |
| 24 | forward | agctcggtacccggggatcctggaacgtgaacttgaaaac |
| | reverse | atgcgggagtgactagtatttatgctttctataaaggagac |
| 25 | forward | gctcggtacccggggatcctttagcagcatttaaacattattacctaacagaaagaaag |
| | reverse | aaatgcgggagtgactagtggccgttactacgacttcgtctag |
| 26 | forward | gctcggtacccggggatcctgtaatctatctaattttatgttcgtttcataaatttaaaaccataca |
| | reverse | aaatgcgggagtgactagtactctgtgattttaaaaagtaagtcataagggttttatattaga |
| 27 | forward | agctcggtacccggggatcccgaccggcttccggaacgtt |
| | reverse | atgcgggagtgactagtcattcaatctttttttagtgattaatcggt |
| 28 | forward | agctcggtacccggggatccttgcttataaatctattcgtctgacag |
| | reverse | atgcgggagtgactagtattgcgcgcttttaataagaagacac |

| | | |
|---|---|---|
| 29 | forward | agctcggtacccggggatccgctggcactggcctatggta |
| | reverse | atgcgggagtgactagtacgtattaggtttttaaataaaacaataagtaaataac |
| 30 | forward | gctcggtacccggggatcccaaaaagaaacgtttgcagggtagttttatatcc |
| | reverse | aaatgcgggagtgactagttaattttcttttcataaattctctcttaaagttcaactttacattaatt |
| 31 | forward | gctcggtacccggggatccaaagatagtaaaaatcaacccattccgattcaaag |
| | reverse | aaatgcgggagtgactagtgtttttgtttaattgtatgttcactaattggtttttgaatagt |
| 32 | forward | gctcggtacccggggatccgcaaatgataaaaaaagcaattttgaatgaaaagatatacatg |
| | reverse | aaatgcgggagtgactagtgtttttctataagattttattagtttgaaaaaaggtcttttaggc |
| 33 | forward | agctcggtacccggggatccttgcaatcattacctgaacaaacct |
| | reverse | atgcgggagtgactagtctttttatcgttttcatttatataacgattc |
| 34 | forward | agctcggtacccggggatccgaaacgccagacgaatcagg |
| | reverse | atgcgggagtgactagtcaatactttttaattttcatttcttttgc |
| 35 | forward | gctcggtacccggggatccgtacagaaagcattgaaacgtatgcc |
| | reverse | aaatgcgggagtgactagtagatattggcttttcctatttatatttatagaaaggc |
| 36 | forward | gctcggtacccggggatccagatagatattgggagtgcatatgtttgtttttac |
| | reverse | aaatgcgggagtgactagtttctatattctttatatttgattagtttctattatatgattaaacagcggatatc |
| 37 | forward | gctcggtacccggggatcctatgaaagtcagcttttacgcctctttg |
| | reverse | aaatgcgggagtgactagtgcatagttttctacattaataattggtttatataatctcaaaaatgatatcttata |
| 38 | forward | gctcggtacccggggatcctgttccttacaaagcctccttttcca |
| | reverse | aaatgcgggagtgactagtactgtttaatttgataggttaataatatatttagtttatagttaggtcaac |
| 39 | forward | agctcggtacccggggatccactacctcaaaacgcagatc |
| | reverse | aaatgcgggagtgactagtgtatcagaattttaaattagtattagagttcacatttaaggataaatca |
| 40 | forward | agctcggtacccggggatccgttctaacttttaccgcaaac |
| | reverse | aaatgcgggagtgactagttttctttattattttggttaaacatttatatcatgatccttataaaagg |
| 41 | forward | agctcggtacccggggatccccgggcgaatggcacaccta |
| | reverse | atgcgggagtgactagtagcattattcatttttaattatgatac |
| 42 | forward | gctcggtacccggggatccgtacatgcaggatctatatccccg |
| | reverse | aaatgcgggagtgactagtatataattagattattgtttcgactgattgacgcaga |
| 43 | forward | gctcggtacccggggatcccttcggaattgctaacacctacg |
| | reverse | aaatgcgggagtgactagtaagttagactttttgaataacataataattaagtcaatataaatagttaagtatagt |
| 44 | forward | agctcggtacccggggatccctattaatatccgacaattcagttgttaatacct |
| | reverse | atgcgggagtgactagtacttattttattgtctttgattctatgtattaaaacag |
| 45 | forward | gctcggtacccggggatccacaatcggctaaagtcataaacctgac |
| | reverse | aaatgcgggagtgactagtctttttataatattaatgagttctattgaattcacgtc |
| 46 | forward | agctcggtacccggggatccaaaaacaatgcaagaaatggaaac |
| | reverse | aaatgcgggagtgactagtaatttattgtattagattataaactctattcaattgttgattgatcca |
| 47 | forward | gctcggtacccggggatccagttgtgtttcttgcagccaac |
| | reverse | aaatgcgggagtgactagtaacgtttttctttttacatattaaattattggttcactaaaaattc |
| 48 | forward | gctcggtacccggggatccgctactctgtgaccactatttataattaccgg |
| | reverse | aaatgcgggagtgactagtgtacatcaatttaaagttaatattaggattactttttgttttcactg |
| 49 | forward | gctcggtacccggggatcctaatgtagctggcagacatccg |
| | reverse | aaatgcgggagtgactagtatgattctaaaaagttaaacgttatttatgtatgattgtgatgc |
| 50 | forward | gctcggtacccggggatccacaaaaaaacattcattcctaaaataaaggatgatggac |
| | reverse | aaatgcgggagtgactagtggataatttaaaattaatattaggttaatacatttcaggcaactagatc |
| 51 | forward | gctcggtacccggggatccctgcaaaacgtcctgtttctaaaaaatg |
| | reverse | aaatgcgggagtgactagtctgtttcctttttttcataatactacatttaataataaagattcatact |
| 52a | forward | gctcggtacccggggatccaaggaagtgtttagatgacataatgattatttgaacag |
| | reverse | aaatgcgggagtgactagtcgttgtacctttcactaatacggatgc |
| 52b | forward | gctcggtacccggggatcccctgtcaccttttaggtgtttgg |
| | reverse | aaatgcgggagtgactagtattctcctataacctattttcatactaattatttatatctaatat |
| 53 | forward | gctcggtacccggggatcctccttgcttttgtgggggtg |
| | reverse | aaatgcgggagtgactagtaattttatctatttctaatgaggattttatctttgctttactataattatac |
| 54 | forward | gctcggtacccggggatcctctcttgaaactgtgaagactcaaaagaag |
| | reverse | aaatgcgggagtgactagtattatttcaggttatttatagcaaagacgactaagaag |
| 55 | forward | gctcggtacccggggatccttctcttcgtccattcattgtgtttttcatatag |
| | reverse | aaatgcgggagtgactagttctatataatattatattcattattaactaaagattttccttccaaaatacaggtagct |
| 56 | forward | gctcggtacccggggatccgaatacaatttataattatcgggcgaaagtaaaaaacaaagc |
| | reverse | aaatgcgggagtgactagtgcgtattaattttaaagtttataattaaggtatgtgcttgacag |
| 57 | forward | gctcggtacccggggatccaaaatggaactgggcaatgacagg |
| | reverse | aaatgcgggagtgactagtccttttcttgtctggttggatagatgtttttt |
| 58 | forward | gctcggtacccggggatccacatttctcccttgaagggc |
| | reverse | aaatgcgggagtgactagtaaatataaataataaaatggttaaagtgcatccgaacaataaatattattgc |
| 59 | forward | gctcggtacccggggatccaaagttagaagtcatcaattaatagaccttcattttgg |
| | reverse | aaatgcgggagtgactagtgattttgtttttttcatacgtaaaaaattaatgattaataaaatattatattgttgg |
| 60 | forward | gctcggtacccggggatccgaataaatgtcgaatttgactcagcgctaag |
| | reverse | aaatgcgggagtgactagtaatattattattttataacttaatacttacaggcatatgagcccc |
| 61 | forward | gctcggtacccggggatccgtctgcctgatgtaaaagagtagttgca |
| | reverse | aaatgcgggagtgactagttcgttactatttaggttgaagtttattctgccg |
| 62 | forward | gctcggtacccggggatccaatgtcgcgatgagcaacag |
| | reverse | aaatgcgggagtgactagtagtttactttttttagtttctactctttttagtatatatttaacctattg |
| 65 | forward | gctcggtacccggggatcccgtaaagggaactatagtgcatcttgc |
| | reverse | aaatgcgggagtgactagtagcaaaaatatttaagatattagtaaataaaaaattaaccgttcattaattga |
| 64 | forward | gctcggtacccggggatccatttttccagttccaatcggcattatg |
| | reverse | aaatgcgggagtgactagtaaatattcctattagattatataccgcaaatgtaacaac |
| 66 | forward | agctcggtacccggggatcctacttcctgcctcatctgctttc |

| | | |
|---|---|---|
| [illegible] | reverse | atgcgggagtgactagttctatttatggtattaaattataagctaac |
| 67 | forward | agctcggtacccggggatccatcgtctggatggaaaatctcataaag |
| | reverse | atgcgggagtgactagtattataagttgagttactttataagtaataaacag |
| 68 | forward | gctcggtacccggggatccgatttttatcttgattttaatagcggggaatttaac |
| | reverse | aaatgcgggagtgactagtaagatattaaatttaattcaaataaaataaaaagtaaaaagggatcattacatac |
| 69 | forward | gctcggtacccggggatccatcggtggcggagcctgtcc |
| | reverse | aaatgcgggagtgactagtaacagaatccaaagaaaggatgctccc |
| 70 | forward | agctcggtacccggggatccttatatggaagacctggaagg |
| | reverse | aaatgcgggagtgactagtactacttgtttttaattaataataataaggttaccaaatacgtttaatacg |
| 71 | forward | gctcggtacccggggatccagaggaactatagtgcttcttgcg |
| | reverse | aaatgcgggagtgactagtgttactttgttttttttaattaatactaaggttatcaatcgcct |
| 72 | forward | gctcggtacccggggatccatatttgactccggattattttcgctggt |
| | reverse | aaatgcgggagtgactagttagtttttatataaatattagaatcgatattccatttattattttcgattcaca |
| 73 | forward | agctcggtacccggggatccagaatataacaacctttatcacg |
| | reverse | aaatgcgggagtgactagtaaattaatattagttttattaaaggtaaaaagggtaaaaacaagttgaag |
| 74a | forward | agctcggtacccggggatccctattatcagaagatggagtga |
| | reverse | aaatgcgggagtgactagtagaaaataccgtttttaagaatttatataataataatgatcataattttactttcg |
| 74b | forward | gctcggtacccggggatccattttccgggtagtgctgacttt |
| | reverse | aaatgcgggagtgactagttattattagttctttttaatatgtgtactattttattgttatttatctaatgacga |
| 75a | forward | gctcggtacccggggatccttgagtaaagaatacaccaatctggaggtatatct |
| | reverse | aaatgcgggagtgactagtcttttgttcttttatattagattattaatattcgagtgtgtccac |
| 75b | forward | gctcggtacccggggatccaattgagtgtccatcaggcg |
| | reverse | aaatgcgggagtgactagtcatttcaatttaaagttaatacatattactatctaatcaatacttatgttacatgc |
| 76 | forward | gctcggtacccggggatccagtacctaccaatggagaattcagc |
| | reverse | aaatgcgggagtgactagtaccatacaaacaaaacaggattaatatttgtagtattcttc |
| 77 | forward | gctcggtacccggggatccacaattatatttgaaaagtttgccatattgcttttatatatgg |
| | reverse | aaatgcgggagtgactagtcttgtcctcacaagagagaaaaaaagg |
| 78 | forward | gctcggtacccggggatccctttaatcaaactcatctttgtatctttatccatcctatc |
| | reverse | aaatgcgggagtgactagtccaaaatcctcctattgttagtaaaagagcaag |
| 79 | forward | gctcggtacccggggatccctggaatttttttatgtaatatacccttttacttcttattacgg |
| | reverse | aaatgcgggagtgactagttgtttttctgtgattaaaggttaataattagttgggttaatatg |
| 80 | forward | gctcggtacccggggatcctggagagcaatagagaccttatgc |
| | reverse | aaatgcgggagtgactagttgtttttctgtgataaaaggttaataattagattggttaaaaaaaagtg |
| 81 | forward | agctcggtacccggggatccagaagggagtcttgcactatttc |
| | reverse | atgcgggagtgactagtcccttattccaccttttattatatgacaag |
| 82 | forward | gctcggtacccggggatccggaagttgatgctgaagatgtt |
| | reverse | aaatgcgggagtgactagtcattaatttcttcctttatacactaaatac |
| 83 | forward | gctcggtacccggggatcctctttgtctccgaaaaacaagagtccg |
| | reverse | aaatgcgggagtgactagtttcaataagttgtttttcaccatctccgtgc |
| 84 | forward | gctcggtacccggggatcctaaagccaaaaagaaacttactgtacctc |
| | reverse | aaatgcgggagtgactagttttcaatgttgatttatcttaggacaatctcaa |
| 85 | forward | gctcggtacccggggatcctttttttcttcttttattgaatgatgacggattttaaaatcatc |
| | reverse | aaatgcgggagtgactagtactttattaggttttataattattaaaagtgttattttactgttgc |
| 86 | forward | gctcggtacccggggatcctgtctcccgtggaatatatcaagtcc |
| | reverse | aaatgcgggagtgactagtgctatcaaatttagtggtttgttgatagcg |
| 87 | forward | gctcggtacccggggatcccaggcagatgatacattcacaaagaaaaagat |
| | reverse | aaatgcgggagtgactagtaactaaaggtagttttagatttaaagcatatttaggatataatgca |
| 88 | forward | gctcggtacccggggatccacttcctaaagaatactttacgggaacgattc |
| | reverse | aaatgcgggagtgactagtatatcaaaacttttaagttgtttgacaggtataccga |
| **rpoD-reporter plasmids** | | |
| **Species** | **Primer direction** | **Primer Sequence** |
| *B. fragilis* | forward | gctcggtacccggggatccagcagagtatcacagagcagag |
| | reverse | aaatgcgggagtgactagtcaaagttacgactatttatttgttaacgtataaattttc |
| *B. cellulosilyticus* | forward | agctcggtacccggggatccctaagactcaggaacaggttttgttt |
| | reverse | atgcgggagtgactagttcttctaaaacagatttgggttgcaaag |
| *B. uniformis* | forward | agctcggtacccggggatccaaggcgatgaaagaggcaaga |
| | reverse | atgcgggagtgactagttcttataatacagatttgggttgcaaaggt |
| **Engineering genetic deletions by allelic exchange** | | |
| **Amplicon** | **Primer direction** | **Primer Sequence** |
| ΔBT2160_UP | forward | gctctagaactagtggatccaaatcagatatttaactccgccctcag |
| | reverse | cagaacaaatcgctttttaatatatgaagtaaatttgttc |
| ΔBT2160_DOWN | forward | catatattaaaaaagcgatttgttctgccaatataaacgaatttgtgccattcg |
| | reverse | agataacattcgagtcgactgccagtcccggagtcatgt |
| ΔBT2628_UP | forward | caggatcctttcacataagaacctttgt |
| | reverse | gccccgggagtgtattattcttagattt |
| ΔBT2628_DOWN | forward | gccccgggtaaatttgaaatgttgtaaa |
| | reverse | gccgtcgacataatactatttttcacatt |
| ΔBT3786_UP | forward | gcggatcccctttacataagagcctttgtcgaa |
| | reverse | gacccgggggcatagaatccttagatttgcaat |
| ΔBT3786_DOWN | forward | gacccgggttattggagtggtatggcaacaatt |
| | reverse | gccgtcgacgataggcagtatccgtacca |

| | | |
|---|---|---|
| ΔBT3853_UP | forward | gctctagaactagtggatccatcttaggtgctactctcggtg |
| | reverse | attattaatagaggattaataaaaaccggaatccttgttcaaaagtaag |
| ΔBT3853_DOWN | forward | tgaacaaggattccggtttttattaatcctctattaataattttccggtgcttcc |
| | reverse | gaagataacattcgagtcgactctcttcggattatagccgacatatcc |
| ΔBT3957_UP | forward | gctctagaactagtggatccggaagaacgacggtcgtgta |
| | reverse | tatattaatagcaagtgttattgttcatgtcttattag |
| ΔBT3957_DOWN | forward | catgaacaataacacttgctattaatatatgctatccccactatcttattctttact |
| | reverse | gaagataacattcgagtcgactcagacacacgccctgttac |
| ΔBT3957-65_DOWN | forward | catgaacaataacacttgctattaatatataaggttaggtattaagataaaaatggaaaggatg |
| | reverse | gaagataacattcgagtcgacatctggcagacggttctgcc |
| ΔBT3309_UP | forward | gctctagaactagtggatcctttacccgtcgcttttatcaaccg |
| | reverse | tagagtattaaagttttgtaggaaatatgacattcttc |
| ΔBT3309_DOWN | forward | tcatatttcctacaaaactttaatactctatcgatgttgtaacctatccactttttagg |
| | reverse | aagataacattcgagtcgacttccgttcatagaagtagtaccacg |
| **Engineering protein expression** | | |
| **Recombinant Protein** | **Primer direction** | **Primer Sequence** |
| N6H4A-BT3312 | forward | catcacgccgcggccgcttgtagcaatagtgatgatgcg |
| | reverse | gcttatcatcgataagcttactttgactttgcccaacggt |
| N6H4A-BT3313 | forward | catcacgccgcggccgcttgtaatgatgacgagttcct |
| | reverse | agcttatcatcgataagcttacttcttcacaacagaaag |
| N6H4A-BT3961 | forward | acggaggtggtgcggccgctggcatgacggaaccttggaa |
| | reverse | agcttatcatcgataagcttacttcttcattctaaaccaacgctcg |
| N6H4A-BT3962 | forward | catcacgccgcggccgcgcaaaagcagccggtggat |
| | reverse | gcttatcatcgataagcttttacttgttatcacgagagaaggaataggg |
| N6H4A-BT3963 | forward | catcacgccgcggccgctcagcaaccggtagattacg |
| | reverse | gcttatcatcgataagcttaattgattccattagttaatgaataaggtttgtcctg |
| N6H4A-BT3965 | forward | catcacgccgcggccgctcaaactgaaaagctgacgg |
| | reverse | gcttatcatcgataagcttaaaacccaattgatgatttatacttcatcgtcttc |
| **qPCR** | | |
| **target gene** | **Primer direction** | **Primer Sequence** |
| BT0029 | forward | atccggagcgtatgaataccg |
| | reverse | tcgcgcccattttgtaacag |
| BT0140 | forward | tcgctggcaattcttttggg |
| | reverse | agataacggctgcatgaagc |
| BT0190 | forward | ttaaaccctgtggcaatggc |
| | reverse | agttccagttcggcaaagac |
| BT0206 | forward | atggcgatgctttgtggtac |
| | reverse | aacgatttgccggtcatcac |
| BT0439 | forward | aatgcaaccccaaacagcag |
| | reverse | atggcaccaataacgcttgc |
| BT0452 | forward | ttgtgcctggaacggatttg |
| | reverse | tgaatgcttgtgtccagacc |
| BT0867 | forward | acaagtcgccggtttgattg |
| | reverse | acgcctcgaatccaaaactc |
| BT1119 | forward | accatatatcccgcagacatcg |
| | reverse | aaccgtacattgccgtttcg |
| BT1440 | forward | aattgaagcccctgccaatg |
| | reverse | ttgttcaccgcttgttccac |
| BT1552 | forward | actgtggcgtatctcaaacg |
| | reverse | aagacaaacggagagcaagg |
| BT1774 | forward | atgctggtttcgaccgtttg |
| | reverse | aaccacgccatgtcattacg |
| BT1875 | forward | cggcaaatggatggaagatcc |
| | reverse | tgttgccagaaaacgcttgg |
| BT2107 | forward | tgaccagccatctaggtatgtc |
| | reverse | tgcattccgtcgtttccttg |
| BT2362 | forward | acgggacattgaaccattcc |
| | reverse | tccatgcgttgtcattggtg |
| BT2364 | forward | tcgttcgacaaactgggaac |
| | reverse | tcatgtatgcgggcaagttg |
| BT2531 | forward | ttaccggacgtttcaacacc |
| | reverse | tggcttcgtttgctaactgc |
| BT2952 | forward | tctttcttcacctgccaacg |

| | | |
|---|---|---|
| [illegible] | reverse | ttaccggcactgttgaaacc |
| BT2968 | forward | tttcccgctgtatatcctaccg |
| | reverse | acttccgtcatcgctgtttc |
| BT3156 | forward | ttctgaacggtttgctgctc |
| | reverse | atggctcccaaaagctttcg |
| BT3174 | forward | ggtaagttcggaagcaatcgc |
| | reverse | cctgaagggcactggtcaag |
| BT3240 | forward | tgtgtcagtccagaacgtatcc |
| | reverse | aaatggaggtagcaccacgtac |
| BT3297 | forward | tgaaagtaggagtcgcaggttc |
| | reverse | attgttgcgcatggctgatg |
| BT3494 | forward | atcgtgtggaccaatgcaag |
| | reverse | actttgtgtcgggctattgc |
| BT3505 | forward | tggcaaacgtcgtgaattcc |
| | reverse | atgtctactgtccgcttctcac |
| BT3670 | forward | atttgcacacttgggacagc |
| | reverse | cgattgcctggaaactaacgc |
| BT3952 | forward | tgacgccaacacgaacaatg |
| | reverse | ttgtttcagacgcaggaagc |
| BT3958 | forward | tcgtttgacgcaagaccttg |
| | reverse | tatgcacgtcgaaagccaac |
| BT4039 | forward | ttgcacgcaacgaaatcctc |
| | reverse | ggcgagttcctttcttttcgag |
| BT4081 | forward | tcgtgttgcaggtgtgattg |
| | reverse | aattccgcgcacccaaaac |
| BT4247 | forward | tgccaacgtcaacatttccg |
| | reverse | tttccctgtgttcagcaagc |
| BT4267 | forward | tcatactttcggtgcgttgc |
| | reverse | atgctgctgatgcccaaatc |
| BT4470 | forward | actccactcgcattgattcg |
| | reverse | tgcagttggtaatgggcatc |
